# Structures of DPAGT1 explain glycosylation disease mechanisms and advance TB antibiotic design

**DOI:** 10.1101/291278

**Authors:** Yin Yao Dong, Hua Wang, Ashley C.W. Pike, Stephen A. Cochrane, Sadra Hamedzadeh, Filip J. Wyszyński, Simon R. Bushell, Sylvain F. Royer, David A. Widdick, Andaleeb Sajid, Helena I. Boshoff, Ricardo Lucas, Wei-Min Liu, Seung Seo Lee, Takuya Machida, Shahid Mehmood, Katsiaryna Belaya, Wei-Wei Liu, Amy Chu, Leela Shrestha, Shubhashish M.M. Mukhopadhyay, Nicola A. Burgess-Brown, Mervyn J. Bibb, Clifton E. Barry, Carol V. Robinson, David Beeson, Benjamin G. Davis, Elisabeth P. Carpenter

**Affiliations:** Structural Genomics Consortium, University of Oxford, Oxford, OX3 7DQ, UK.; Chemistry Research Laboratory, University of Oxford, 12 Mansfield Road, Oxford OX1 3TA, UK.; School of Chemistry and Chemical Engineering, Queen’s University Belfast, UK.; Department of Molecular Microbiology, John Innes Centre, Norwich Research Park, Norwich NR4 7UH, UK.; Tuberculosis Research Section, Laboratory of Clinical Immunology and Microbiology, National Institute of Allergy and Infectious Diseases, National Institutes of Health, Bethesda, Maryland 20892, USA.; Department of Chemistry, South Parks Road, Oxford OX1 3QZ UK.; Neurosciences Group, Nuffield Department of Clinical Neuroscience, Weatherall Institute of Molecular Medicine, University of Oxford, Oxford, OX3 9DS, UK.

**Keywords:** DPAGT1, GPT, Protein N-glycosylation, congenital myasthenic syndrome, congenital disorders of glycosylation, antibiotic design, tunicamycin

## Abstract

Protein glycosylation is a widespread post-translational modification. The first committed step to the lipid-linked glycan used for this process is catalysed by dolichyl-phosphate N-acetylglucosamine-phosphotransferase DPAGT1 (GPT/E.C. 2.7.8.15). Missense DPAGT1 variants cause congenital myasthenic syndrome and congenital disorders of glycosylation. In addition, naturally-occurring bactericidal nucleoside analogues such as tunicamycin are toxic to eukaryotes due to DPAGT1 inhibition, preventing their clinical use as antibiotics. However, little is known about the mechanism or the effects of disease-associated mutations in this essential enzyme. Our structures of DPAGT1 with the substrate UDP-GlcNAc and tunicamycin reveal substrate binding modes, suggest a mechanism of catalysis, provide an understanding of how mutations modulate activity (and thus cause disease) and allow design of non-toxic ‘lipid-altered’ tunicamycins. The structure-tuned activity of these analogues against several bacterial targets allowed design of potent antibiotics for *Mycobacterium tuberculosis*, enabling treatment *in vitro*, *in cellulo* and *in vivo* thereby providing a promising new class of antimicrobial drug.

**Highlights:**

- Structures of DPAGT1 with UDP-GlcNAc and tunicamycin reveal mechanisms of catalysis
- DPAGT1 mutants in patients with glycosylation disorders modulate DPAGT1 activity
- Structures, kinetics and biosynthesis reveal role of lipid in tunicamycin
- Lipid-altered, tunicamycin analogues give non-toxic antibiotics against TB

## Introduction

N-glycosylation of asparagine residues is a common post-translational modification of eukaryotic proteins, required for protein stability, processing and function, and many diseases are associated with incorrect glycosylation (Freeze et al., 2012). This process requires dolichol-PP-oligosaccharides, that provide the oligosaccharides that are transferred onto asparagine residues (Helenius and Aebi, 2004). The first step in production of dolichol-PP-oligosaccharides, involves the ER integral membrane enzyme dolichyl-phosphate alpha-N-acetyl-glucosaminyl-phosphotransferase (DPAGT1, E.C. 2.7.8.15, also known as GlcNAc-1-P Transferase (GPT)). It catalyses the transfer of an N-acetyl-D-glucosamine-1-phosphoryl unit (GlcNAc-1-P) from UDP-N-acetyl glucosamine (UDP-GlcNAc) onto dolichyl phosphate (Dol-P) (Figure 1A) (Heifetz and Elbein, 1977; Lehrman, 1991). The product GlcNAc-PP-Dol is anchored to the ER membrane by its dolichyl moiety and then sugar units are added sequentially to build the N-glycan that is then transferred onto Asn residues.

**Figure 1.**
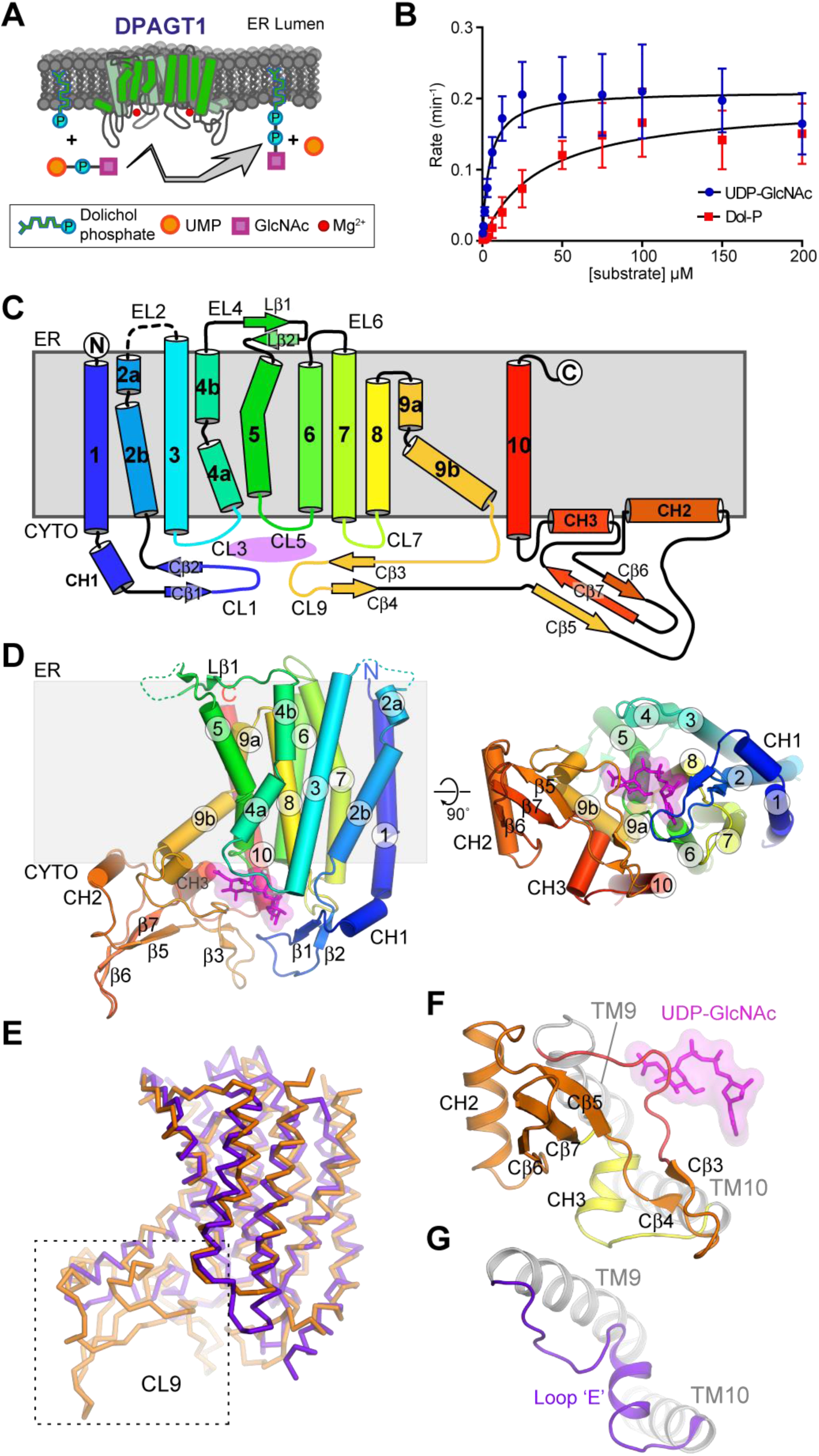
The structure of DPAGT1 reveals a cytoplasmic facing active site with a CL9 domain forming one face of the active site. (A) Cartoon of the DPAGT1 reaction. (B) Michaelis-Menten kinetics (C) Topology of DPAGT1 with helices shown as cylinders, strands as arrows and the active site indicated in magenta. (D) Schematic of the DPAGT1 structure. Two perpendicular views are shown looking along the membrane plane and onto the cytoplasmic face, with UDP-GlcNAc in magenta. (E) Comparison of the DPAGT1 (orange) and MraY (PDB: 5CKR; purple) folds. (F) CL9 domain in DPAGT1. (G) Short CL9 strand (loop ‘E’) and single helix in MraY.

Mutations in DPAGT1 impair protein N-glycosylation, leading to at least two syndromes, depending on the extent of loss of activity. Congenital myasthenic syndrome (DPAGT1-CMS OMIM ref: 614750) is a disorder of neuromuscular transmission characterised by fatigable weakness of proximal muscles (Basiri et al., 2013; Belaya et al., 2012; Iqbal et al., 2013). A reduction in endplate acetylcholine receptors (AChR) and abnormal synaptic structure are thought to be the result of incorrect glycosylation of the AChR and other synaptic proteins. Mutations in DPAGT1 also cause congenital disorder of glycosylation type Ij (CDG-Ij, OMIM ref: 608093) (Carrera et al., 2012; Selcen et al., 2014; Wu et al., 2003; Wurde et al., 2012), a more severe multisystem syndrome that may involve intellectual disability, epilepsy, microcephaly, severe hypotonia, structural brain anomalies.

Inhibition of polyisoprenyl-phosphate N-acetylaminosugar-1-phosphoryl transferases (PNPTs), such as DPAGT1 and the bacterial enzyme MraY, by small molecules is lethally toxic to many higher and lower organisms. *Streptomyces* bacteria have exploited this toxicity by producing the PNPT inhibitor tunicamycin, which blocks MraY, a critical enzyme in biosynthesis of cell walls in many bacterial pathogens (Figure S1A,B) (Dini, 2005). Tunicamycin has antibacterial, anti-fungal and anti-viral activities (Takatsuki et al., 1971; Takatsuki and Tamura, 1971). Unfortunately, it also inhibits eukaryotic PNPTs, such as DPAGT1 (Heifetz et al., 1979) leading to severe toxicity in eukaryotic cells. However, although bacterial (e.g. MraY) and human (e.g. DPAGT1) PNPTs are similar, it should be possible to design unnatural tunicamycin analogues that specifically inhibit bacterial proteins.

Here we present structures of human DPAGT1 with and without ligands. The protein production methods, structures, assays and complexes with substrates and inhibitors are components of a “target enabling package” developed at the Structural Genomics Consortium and released in June 2017 (http://www.thesgc.org/tep/DPAGT1), which has already been used by others (Yoo et al., 2018). These structures, combined with site-directed mutagenesis and activity analysis, reveal both the mechanism of catalysis by DPAGT1 and the molecular basis of DPAGT1-related diseases. In order to improve the effectiveness of tunicamycin as a drug, we modified the tunicamycin core scaffold, **TUN**, using a scalable, semi-synthetic strategy that enabled selective lipid chain addition. These tunicamycin analogues show nanomolar antimicrobial potency, ablated inhibition of DPAGT1 and much reduced toxicity. These non-toxic tunicamycin analogues allowed rapid and effective clearance of *Mycobacterium tuberculosis* (*Mtb*) from mammals, thus providing leads for tuberculosis (TB) antibiotic development.

## Results

### DPAGT1 activity and architecture

To determine the structure of DPAGT1, we expressed both full-length wild type (WT) DPAGT1 and the missense mutant Val264Gly in the baculovirus/insect cell system. The Val264Gly mutation is a variant found in some CMS patients (Belaya et al., 2012), which improved crystallisation behaviour, compared to the WT protein. We tested the enzymatic activity of WT and Val264Gly DPAGT1, to confirm that the protein was functional. The identity of the product GlcNAc-PP-Dol was confirmed by mass spectrometry (Figure S1C). WT DPAGT1 has an apparent *K*_m_ of 4.5 ± 0.8 μM and a k_cat_ of 0.21 ± 0.007 min^−1^ towards the UDP-GlcNAc substrate (Figure 1B, STAR Methods); Dol-P displayed an apparent *K*_*m*_ of 36.3 ± 7.2 μM and a k_cat_ of 0.20 ± 0.012 min^−1^ (Figure 1B). Notably, the Val264Gly mutant showed 2.5-fold higher activity (Figure S1D) and similar thermostability to the WT protein (Tm_1/2_ of 51.7 ± 0.2 °C for the WT and 50.4 ± 0.3 °C for the mutant, Figure S1E, STAR Methods, Supplemental Information SI 1). Whilst a 3-fold reduction in activity was seen in the presence of product analogue GlcNAc-PP-Und (equimolar to Dol-P and UDP-GlcNAc), the addition of the second product, UMP, did not inhibit the reaction (Figure S1F). Tunicamycin gave complete inhibition at a 1:1 molar ratio with DPAGT1 (Figure S1G). While both the substrates thermostabilised WT and mutant DPAGT1 by 3-7 °C, tunicamycin thermostabilised both by more than 30 °C (Figure S1E). Interestingly, phosphatidylglycerol (PG) co-purified with the DPAGT1, in agreement with previous observations that PG increased the activity of DPAGT1 extracts (Kaushal and Elbein, 1985) (Figure S1H,I).

The crystal structures of the WT DPAGT1 and Val264Gly mutant were solved by X-ray crystallography using molecular replacement, with the bacterial homologue MraY ((Chung et al., 2013), PDB: 4J72, 19 % identity) as an initial model (STAR Methods section). The WT- and Val264Gly-DPAGT1 unliganded proteins gave structures to 3.6 Å and 3.2 Å resolution (Table S1). Complexes with UDP-GlcNAc and tunicamycin gave data to 3.1 Å and 3.4 Å resolution, respectively (Figure 1, Figure S2A-E and Table S1; Methods section). In the crystals DPAGT1 is a dimer (Figure S2F), with an 1850 A^2^ interaction surface, although this dimer interface differs from that seen in MraY (Figure S2G). However, unlike the DPAGT1/tunicamycin structures presented by (Yoo et al., 2018), no intermolecular disulphide bond was observed at the dimer interface. In solution DPAGT1 exists predominantly as a dimer, although the monomer was also detected by native mass spectroscopy (Figure S2H, I).

The DPAGT1 structure (as reported here and as a complex with tunicamycin in (Yoo et al., 2018)) consists of 10 transmembrane helices (TMH1 to 10), with both termini in the ER lumen (Figure 1C, D). Five loops connect the TMHs on the cytoplasmic side of the membrane (CL1, −3, −5, −7 and −9), which form part of the active site, three loops on the ER side of the membrane (EL2, −4, −6) and one (EL8) embedded in the membrane on the ER side. DPAGT1 has a similar overall fold to MraY (Chung et al., 2013; Yoo et al., 2018) (Figure 1E). A characteristic feature of the eukaryotic DPAGT1 PNPT family, not found in prokaryotic PNPTs, is a 52-residue insertion between Arg306-Cys358 in CL9, following TMH9. This motif adopts a mixed α/β fold with an extended structure with two β-hairpins, a three-stranded β-sheet (Cβ5-Cβ7) and two amphipathic α-helices (CH2/CH3). This CL9 domain (Figure 1F) forms part of the substrate recognition site in human DPAGT1 but not in the bacterial MraY (Figure 1G).

The active site is on the cytoplasmic face of the membrane, formed by the four cytoplasmic loops between the TMHs (Figure 2A). The long CL1 loop forms the ‘back wall’; CL5 and CL7 form the base and the ‘side walls’ are formed by TMH3-CL3-TMH4, TMH9b and the extended loop at the start of the CL9 domain (Ile297-Pro305). The entrance to the catalytic site, between TMH4 and TMH9b, is open and accessible from the lipid bilayer via a 10 Å wide cleft. Within the membrane, adjacent to the active site, there is a hydrophobic concave region (Figure 2B), created by a 60° bend in TMH9 midway through the bilayer, which creates a groove in the DPAGT1 surface (Figure 1D).

**Figure 2.**
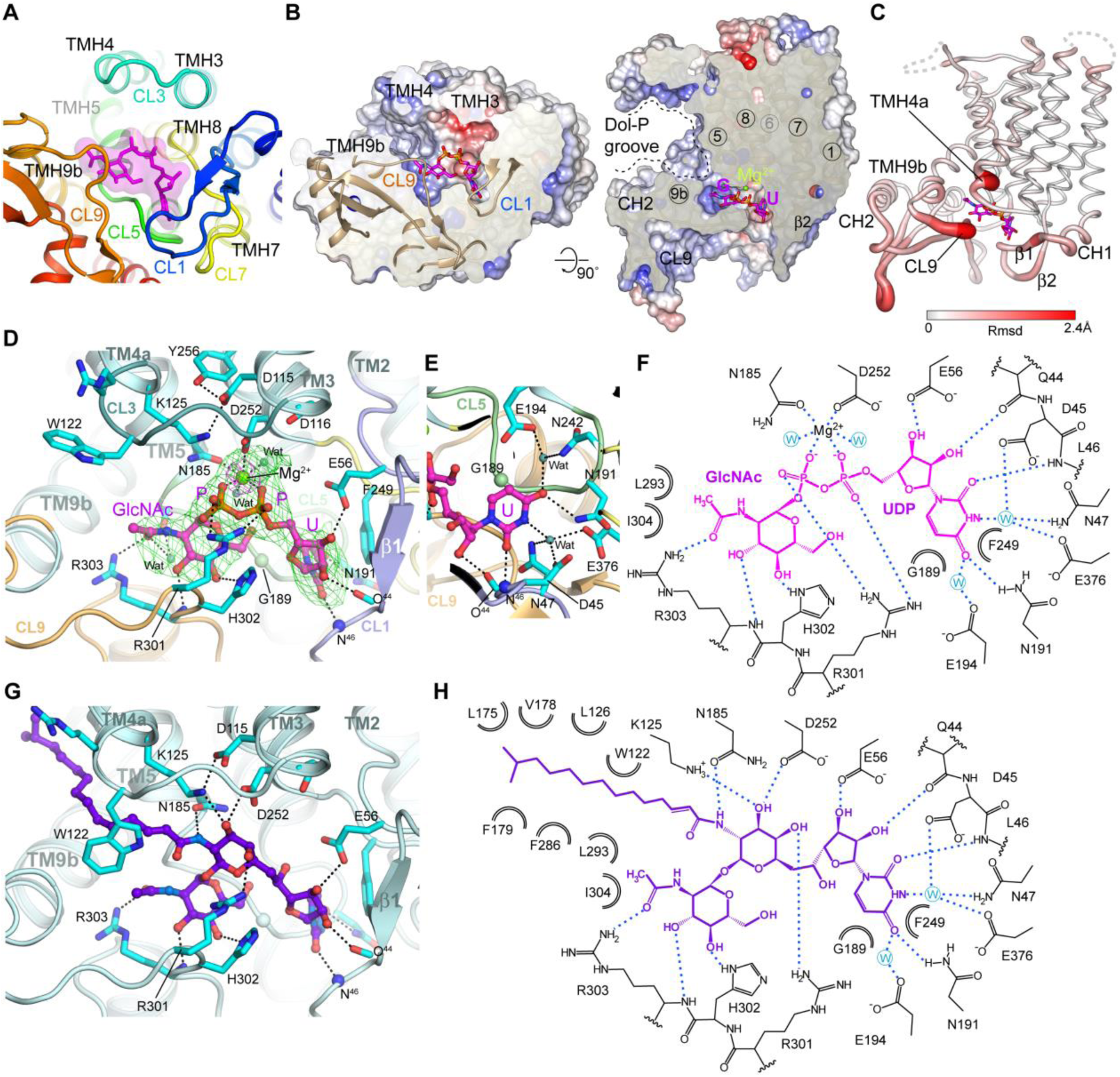
The DPAGT1 active site, and complexes with UDP-GlcNAc and Tunicamycin. (A) Overview of the DPAGT1 active site, showing the loops that form the active site, with UDP-GlcNAc in magenta. (B) Sliced molecular surface showing the occluded active site cleft and putative Dol-P recognition groove. The surface is coloured by electrostatic potential and bound UDP-GlcNAc in magenta. (C) Conformational changes with UDP-GlcNAc binding. The protein is depicted in tube form with the tube thickness and colouring reflecting the rmsd in mainchain atomic positions between the unbound and UDP-GlcNAc-bound structures. (D) Binding mode of UDP-GlcNAc within active site cleft. Omit Fo-Fc difference electron density is shown for UDP-GlcNAc (green mesh, contoured at 3σ) and 4Å anomalous difference Fourier electron density (magenta mesh, contoured at 15σ) from a dataset with MnCl_2_. Recognition of uridine moiety of UDP-GlcNAc. (E) Schematic representation of interactions made by UDP-GlcNAc. (F) Binding mode of tunicamycin within active site cleft. (G) Schematic representation of interactions made by tunicamycin.

### Binding mode for the UDP-GlcNAc and the metal ion

The structure of the DPAGT1/UDP-GlcNAc complex reveals an overall stabilisation of the active site, due to movements of CL1 and CL9, and the N-terminus of TMH4 (Figure 2C), without any global changes in conformation. The C-terminal end of TMH9b and the following loop region (Phe286-Ile304) display the largest conformational change with an induced fit motion around the GlcNAc-PP (Figure 2C).

The uridyl moiety of UDP-GlcNAc is buried in a narrow cleft at the back of the active site formed by CL5 and CL7. The uracil ring is sandwiched between Gly189 and Phe249 with additional recognition conferred by hydrogen bonds between the Leu46 backbone amide, the Asn191 sidechain to the uracil carbonyls (Figure 2D, E) and an extensive hydrogen bond network involving two waters, links the uracil ring to five residues (Figure 2D, E). Hydrogen-bonding of the ribosyl hydroxyls to the Gln44 mainchain carbonyl and Glu56 sidechain carboxylate complete the recognition of the uridyl nucleotide.

The pyrophosphate bridge is stabilised by interactions with Arg301 and by the catalytic Mg^2+^ ion (Figure 2D). The Arg301 sidechain coordinates one pair of α and β phosphate oxygen atoms, whilst the Mg^2+^ ion is chelated by the second pair of α and β phosphate oxygens. Each oxygen atom is thus singly coordinated in an elegant Arg-Mg^2+^-‘pyrophosphate pincer’. The rest of the octahedral coordination sphere of the Mg^2+^ ion comes from the sidechains of the highly conserved residues Asn185 and Asp252, as well as two water molecules. Data from DPAGT1 co-crystallised with UDP-GlcNAc and Mn^2+^ gave a single anomalous difference peak at the metal ion binding site, confirming the presence of a single Mg^2+^ ion in the active site (Figure 2D). The position of the Mg^2+^ ion differs by 4Å from that observed in MraY unliganded structure and the co-ordination differs (Chung et al., 2013).

The GlcNAc moiety-binding site is formed by the CL9 domain and the CL5 loop, although all the direct hydrogen bond interactions are with the CL9 domain. The OH3 and OH4 hydroxyls of GlcNAc form hydrogen bonds with the sidechain of His302 and the mainchain amide of Arg303, respectively. Critically, the mainchain of residues 300–303 and, in particular, the sidechain of Arg303 define the GlcNAc recognition pocket by specifically recognising the N-acetyl substituent, forming a wall to the sugar-recognition pocket that appears intolerant of larger substituents, thereby ‘gating’ substrate. This structure is absent in MraY, which has a much smaller CL9 loop.

To our surprise, this structure does not support prior predictions that the highly conserved ‘aspartate rich’ D^115^Dxx(D/N/E)^119^ motif is directly involved in Mg^2+^ binding and/or catalysis (Lloyd et al., 2004). This sequence is adjacent to the active site, but these residues do not directly coordinate the Mg^2+^ ion or the substrate (Figure 2D). Instead, Asp115 is hydrogen-bonded to the sidechains of Lys125 and Tyr256. Lys125 lies adjacent to the phosphates (Figure 2D) and has been implicated in catalysis (Al-Dabbagh et al., 2008). Asp116 forms hydrogen-bonds to the Ser57 and Thr253 sidechains and N-caps TMH8, thus stabilising the residues that interact with the UDP ribosyl moiety (Glu56) and the Mg^2+^ ion (Asp252). DPAGT1 with residues Asp115 and Asp116 mutated to Asn, Glu or Ala retained at least 10% of WT activity (Figure 3A), suggesting that they are not essential for catalysis. The third, less well-conserved position in the motif, Asn119, does not make any significant interactions. Thus, two of the three conserved acidic residues perform structural roles; none appear directly involved in Mg^2+^-binding or catalysis.

**Figure 3.**
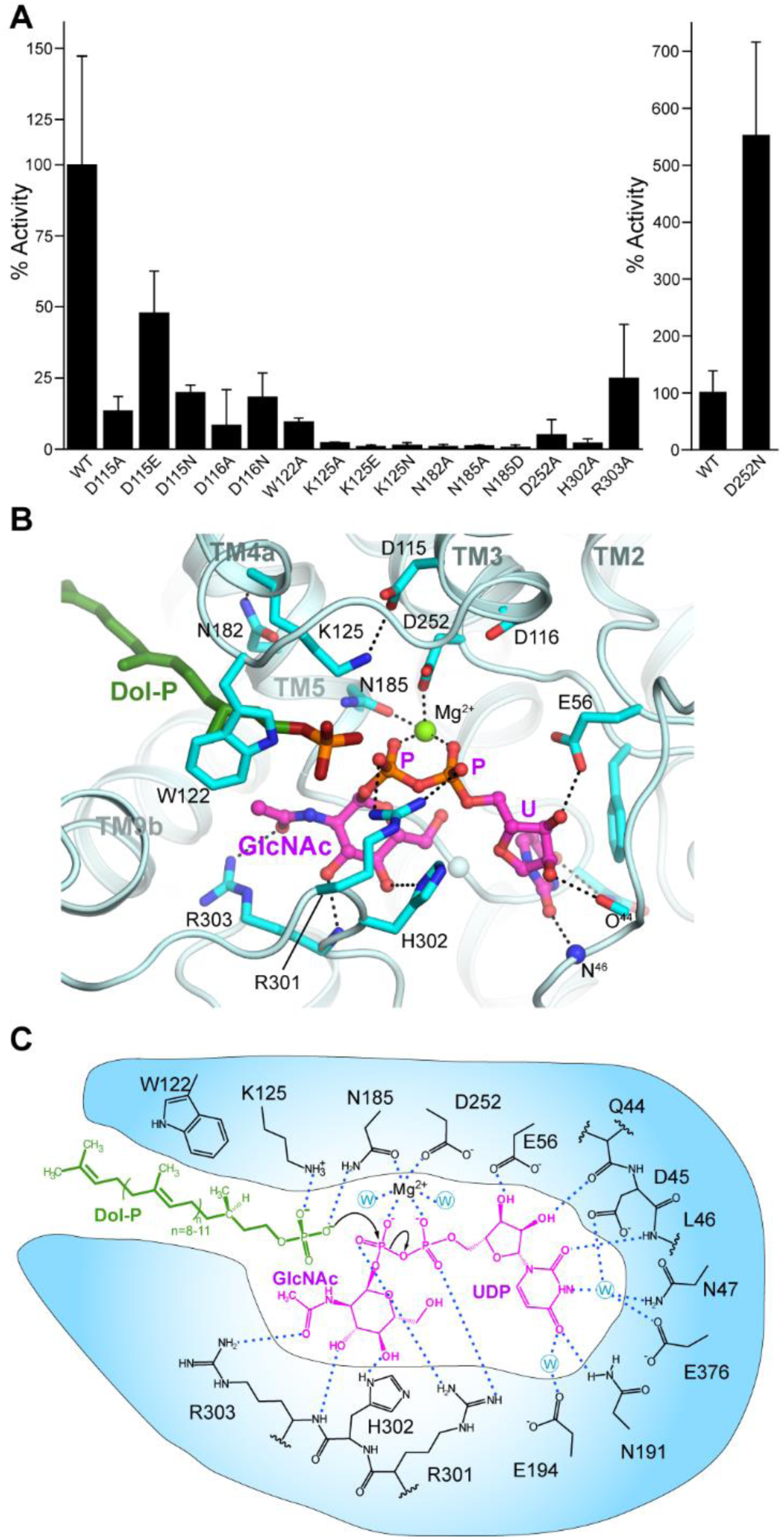
Proposed mechanism of catalysis for DPAGT1. (A) Relative activity for mutants of catalytic site residues. (B) UDP-GlcNAc complex active site structure with Dol-P modelled based on the tunicamycin complex lipid chain position. (C) Proposed catalytic mechanism for DPAGT1.

### Comparison of the UDP-GlcNAc and tunicamycin complexes reveals tunicamycin Michaelis-complex mimicry and DolP substrate binding

The structure of the complex between tunicamycin and DPAGT1 (this work and (Yoo et al., 2018)) shows that inhibition is achieved through its partial mimicry of the Michaelis complex that is formed during catalysis between acceptor phospholipid Dol-P and UDP-GlcNAc. The uridyl and GlcNAc moieties in tunicamycin and UDP-GlcNAc occupy essentially identical binding sites (Figure 2G,H). In tunicamycin the pyrophosphate bridge of UDP-GlcNAc is replaced by a galactosaminyl moiety, which displaces the Mg^2+^ ion and interacts with the sidechains of Arg301, Asp252 and Asn185, thus partially mimicking the pyrophosphate-to-metal chelate arrangement found with UDP-GlcNAc•Mg^2+^.

As well as this UDP-GlcNAc mimicry, tunicamycin’s mimicry of Dol-P gave critical insight into potential binding of co-substrate Dol-P. The lipid chain of tunicamycin occupies the concave groove that runs along TMH5, between helices TMH4 and TMH9a, (see above). The sidechain of Trp122 pivots around its Cβ–Cγ bond to lie over the lipid chain, trapping it in an enclosed tunnel. The surface beyond this hydrophobic tunnel, up to the EL4 loop on the ER lumen face of the membrane, is highly conserved and so it seems likely that the lipid moiety of Dol-P could bind to this surface. At the other end of the tunicamycin lipid moiety, the amide forms polar interactions with Asn185 and lies close to Lys125, suggesting that the amide moiety partially mimics the phosphate head-group of Dol-P (Figure 2G,H).

### The DPAGT1 catalytic mechanism

Several alternative mechanisms have been proposed for the PNPT family including a one-step, simple nucleophilic attack (Al-Dabbagh et al., 2008) or a two-step, double displacement reaction via a covalent intermediate (Lloyd et al., 2004). We did not observe any covalent modification of DPAGT1 in the presence of UDP-GlcNAc, nor did we see release of UMP in the absence of Dol-P as would be predicted for a two-step mechanism. In addition, there are no suitably placed residues in the active site to act as a nucleophile. Therefore the most probable mechanism involves direct nucleophilic attack by a Dol-P phosphate oxygen atom on the phosphorus atom of the β-phosphate of UDP-GlcNAc, causing phosphate inversion and loss of UMP (Figure 3B, C).

When bound to DPAGT1, UDP-GlcNAc adopts a bent-back conformation, with the donor sugar lying below the phosphates, rotated towards the uridine (Figure 2D). The pyranose ring is inclined so that the O6 hydroxyl of the GlcNAc is within 3.1Å of the O5B atom of the α-phosphate. This orientation of the sugar presents the β-phosphate of the UDP-GlcNAc to the position that would be occupied by the phosphate of the Dol-P, exposing the lowest unoccupied molecular orbital (LUMO) of its β-phosphate electrophile for reaction with the Dol-P phosphate *O*-nucleophile (Figure 3B, C).

Providing the correct geometry for this one-step phosphoryl transfer appears from our structural analyses to be key to the catalysis provided by DPAGT1. Analyses of other enzymatic phosphoryl transfer reactions suggest that a bridging Arg (in a very similar position to Arg301) and bridging metals (in a very similar position to the Mg^2+^ ion) do not tighten the transition state but instead provide binding energy that optimizes geometry and alignment for attack (Lassila et al., 2011). They might also preferentially favour the formation of trigonal bipyramidal geometry during nucleophilic attack. Similarly, despite classical emphasis on reducing electrostatic repulsions between anionic nucleophiles with anionic electrophiles, such as those present in phosphoryl transfer (Westheimer, 1987), such effects are small in model systems (Lassila et al., 2011). This suggests that the role here of Lys125, which would be close to the phosphate oxygens in Dol-P, would be mainly to act as a guide to position the phosphate. The correct alignment of Dol-P for attack would be further facilitated by the ‘grip’ provided by Trp122 holding the Dol-chain into the tunnel observed in the tunicamycin•DPAGT1 complex.

Representative residues proposed to bind Dol-P, sugar and pyrophosphate were probed by mutagenesis. Mutation of Mg^2+^-chelating residues to Ala in Asn185Ala and Asp252Ala reduced the DPAGT1 activity to 1.2% and 7%, respectively (Figure 3A). The more conservative Asn185Asp mutation, which would be expected to retain Mg^2+^-binding activity, also ablated activity (0.7% of WT) suggesting an additional role for Asn185 in catalysis. The amide group of Asn185 lies within 4 Å of the predicted Dol-P phosphate-binding site, forming hydrogen bonds with the nucleophilic oxygen of Dol-P to guide it towards the β-phosphate. Mutations of Lys125, which also lies near the Dol-P phosphate binding site, to Lys125Ala, Lys125Glu and Lys125Gln, all reduced the activity to below 2.2%, consistent with a critical guiding role for Lys125. Interestingly, an Asp252Asn mutation increased activity 5-fold (Figure 3A). This mutation removes a coordinating negative charge from the Mg^2+^, making the Mg^2+^ more electropositive and the β-phosphorus more electrophilic, thus potentially increasing its susceptibility to nucleophilic attack.

Mutation of His302, which hydrogen bonds to the O4 oxygen of GlcNAc in UDP-GlcNAc to hold it in its bent-back conformation, causes 98% loss of activity, again consistent with the guiding role that active site residues play in aligning access of the nucleophile to the β-phosphate. Finally, mutation of the Arg301 sidechain that, along with the Mg^2+^ ion, completes the ‘pincer’ of the pyrophosphate caused almost complete loss of activity and this mutation has been found in patients with CDG-Ij (Imtiaz et al., 2012).

### Mutations in DPAGT1 in CMS and CDG

DPAGT1-CMS and CDG-Ij are recessive disorders, caused by either homozygous or compound heterozygous mutations (Supplemental Information SI 1). A variety of mechanisms can cause loss of DPAGT1 function, including RNA splicing errors resulting in exon skipping, protein truncation, instability and loss of enzymatic activity. The structures of DPAGT1, together with data on catalytic activity, thermostability and purification yields of Sf9-expressed, purified protein (STAR Methods, Figure 4A, B, C, Figure S3, and Supplemental Information SI 1) allowed us to explain how many DPAGT1 variants are involved in disease.

**Figure 4.**
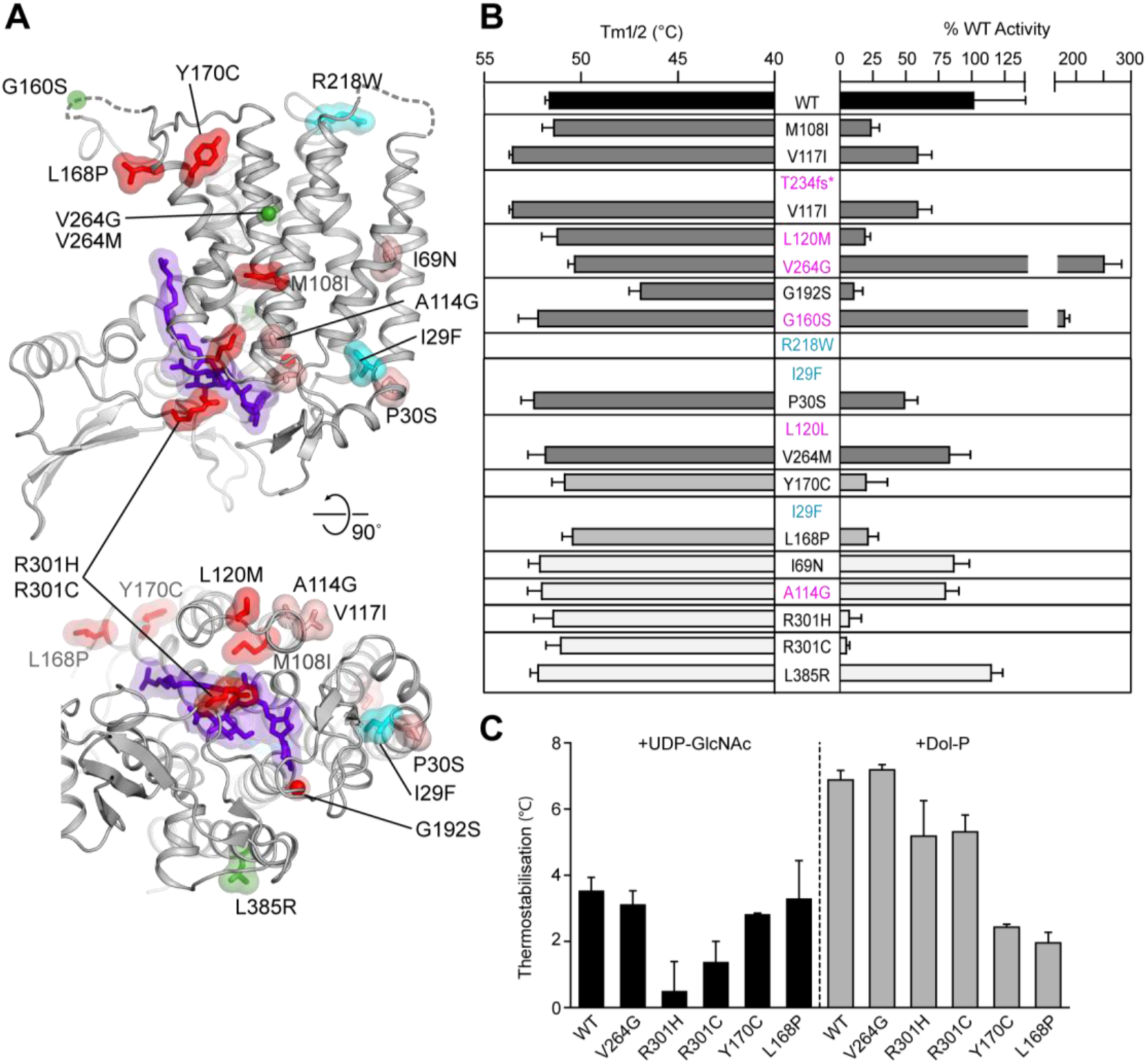
Location, relative activity and stability of missense variants in DPAGT1 found in patients with CMS and CDG-Ij. (A) Location of missense variants in the DPAGT1 protein of patients with CMS and CDG-Ij. Residues are coloured by relative catalytic activity (green, increase in activity; pink (>40%) and red (<40%) residual activity) or instability on purification (cyan). (B) Thermostability (left bars) and relative activity (right bars) for DPAGT1 missense variants found in CMS (dark grey) or CDG-Ij (mid/light grey). Mutations involving truncation (Thr234Hisfs*116) or abnormal splicing (Ala114Gly, Leu120Met, Leu120Leu, Gly160Ser, Val264Gly) are indicated in magenta (although in these cases the pathogenic effect is at the mRNA level). (C) Changes in thermostability of the DPAGT1 disease variants upon the addition of the substrates UDP-GlcNAc and Dol-P.

For mutations found in patients with DPAGT1 CMS, in general we found that one allele had either a relatively large reduction in the catalytic activity of the expressed protein (Met108Ile, Leu120Met, Gly192Ser, Figure 4B, Supplemental Information SI 1) or protein quantity (e.g. truncation in Thr234Hisfs*116 (Belaya et al., 2012), low protein yield for Ile29Phe or exon skipping in Leu120Leu (Selcen et al., 2014)). However, expressed protein from the second allele had catalytic activity that was closer to WT (Val117Ile, Pro30Ser, Val264Met) or surprisingly, even an increase in activity (Val264Gly and Gly160Ser) (Figure 4B, Supplemental Information SI 1).

Missense variants that cause changes in enzyme activity are generally near the active site or in the core of the protein. Met108 lies at the centre of a hydrophobic cluster of residues between TMHs 3, 4, 5 and 8 (Figure S3B). The CL3 loop, between TMHs 3 and 4, forms one side of the active site and TMHs 4 and 5 interact with the Dol-P lipid chain. Therefore replacement of a Met sidechain with the branched Ile sidechain could sterically hinder both the active site and Dol-P binding. The two other activity reducing mutations, Leu120Met and Gly192Ser, are both on loops directly adjacent to the active site and are likely to affect catalysis or UDP-GlcNAc binding: Leu120 lies on the CL3 loop close to the predicted Dol-P/UDP-GlcNAc interface (Figure S3C) and Gly192 lies within the CL5 loop in the uridyl recognition pocket (Figure S3D). CL5 forms a sharp turn at the highly conserved Gly192, allowing the correct positioning of UDP-GlcNAc-interacting residues such as Asn191 and Glu194, suggesting that the CL5 loop conformation at Gly194 is critical for uridine binding. Two other CMS mutants (Ile29Phe, Arg218Trp) expressed protein that was too unstable to purify.

In contrast to missense variants that cause major loss of activity, the mutations that give close to WT activity (Val117Ile, Pro30Ser, Val264Met) are much less disruptive. For example, although Val117 is on the CL3 loop near the active site (Figure S3C), its sidechain projects into the lipid environment of the membrane, where this conservative substitution is easily tolerated. The Pro30Ser mutation lies at the site of a kink in TMH1 as it emerges on the cytoplasmic face of the membrane and a reduction in the kink at this point would alter the positioning of the CL1 loop, causing some disruption to the uracil base-binding site (Figure S3E).

Given that DPAGT1-CMS is a recessive disorder we were surprised to find an increase in enzymatic activity with the Val264Gly and Gly160Ser variants. Val264 is mutated to either Gly or Met in CMS patients, giving either a 2.5-fold increase or a slight (18%) decrease in catalytic activity for purified protein. Val264 is located on TMH8 in the core of the protein adjacent to TMH3/4 (Figure S3F). Comparison of the WT and Val264Gly DPAGT1 structures showed a 1-1.5 Å inward movement at the C-terminal end of TMH4b towards TMH8 (Figure S3G). This movement could affect both the DPAGT1 dimer interface and the exact position of EL4, which lies above this site, forming the top of the Dol-P lipid-binding site. Conversely, the Val264Met variant would be poorly accommodated at this buried site. Since the Gly160Ser mutation lies in the disordered EL4 luminal loop, it is unclear why the activity increases. Since these missense mutations cause an unexpected increase in enzyme activity, we explored other causes of pathogenicity. Tissue from muscle biopsies was not available, so we used the ‘exon trap’ system to detect abnormal RNA splicing. Both mutations, c.478G>A, p.Gly160Ser and c.791T>G, p.Val264Gly, give rise to abnormal RNA splicing of their respective RNA transcripts (Figure S4) thus explaining the pathogenicity of these variants.

Mutations in DPAGT1 can also lead to a more severe or lethal disease, CDG-Ij. In cases of CDG-Ij where the patients survive to adulthood, they suffer severe multisystem disorders that may include hypotonia, medically intractable seizures and mental retardation (Iqbal et al., 2013; Wu et al., 2003). In these cases one allele has approximately 20% activity (Leu168Pro, Tyr170Cys), whereas the second allele either gives protein that was too unstable to purify (Ile29Phe) or it has a splicing defect that reduces WT mRNA levels by 90% (Wu et al., 2003). The Leu168Pro and Tyr170Cys mutations both affect Dol-P binding. The long (85-105 carbon) chain of Dol-P is predicted to bind on a groove between TMH4, TMH5 and TMH9 within the membrane, extending towards the ER, below the EL4 loop. Leu168 and Tyr170 lie at of the N-terminal end of TMH5, below the EL4 loop (Figure 4A and Figure S3H). The Leu168Pro mutation would disrupt the start of TMH5, following EL4 and the Tyr170Cys mutation would disrupt the packing between the N-terminal end of TMHs 4 and 5 thereby disrupting the ‘far end’ of the critical Dol-P binding site. This suggestion is consistent with the changes in thermostability of DPAGT1 with Dol-P. With WT DPAGT1, addition of Dol-P increased the Tm_1/2_ by 6.5 °C, whereas for the Leu148Pro and Tyr170Cys mutations, the change in Tm_1/2_ was only 2 °C, suggesting almost ablated binding for the Dol-P (Figure 4C, Supplemental Information SI 1). Interestingly, tunicamycin has a shorter lipid tail, so it would not extend to the ER end of the Dol-P binding site and the mutations did not affect stabilisation with tunicamycin.

For the most severe cases of CDG-Ij, in which the patients died in early infancy, the causes of loss of DPAGT1 activity are more complex. In two cases, there is either a homozygous Arg301His mutation, or compound heterozygous Arg301Cys and Leu385Arg mutations. The Arg301 mutations both cause a 20-fold drop in enzyme activity, which is easily explained as Arg301 lies in the active site, where it plays a critical role in the bifurcated pyrophosphate binding ‘pincer’ (Figures 2D, F). The Leu385Arg mutation places a hydrophilic Arg sidechain inside the membrane, which we would expect to be destabilising, although in the short chain detergent OGNG, it does retain some catalytic activity (Figure 4A). There are two missense variants found in patients with CDG-Ij where the mutations (Ala114Gly (Wurde et al., 2012) or Ile69Asn (Timal et al., 2012)) do not appear to have a significant effect on protein stability or catalytic activity (Figure 4B), although in cells the overall activity is reported to be reduced to 18 or 22%. For these mutations, a reduction in the amount of the correct mRNA was reported ((Timal et al., 2012; Wurde et al., 2012)), which may explain the loss of in-cell activity.

### Development of non-toxic ‘TUN-X,X’ analogues of tunicamycin

Not only is DPAGT1 clinically important due to its role in disorders of glycosylation, it also plays a significant, albeit negative, role in another important clinical context, namely antimicrobial development. The potent ‘off-target’ inhibitory effects of tunicamycin on DPAGT1 (see above) cause toxicity for a potentially highly valuable antimicrobial tunicamycin that displays a different mode-of-action, that would be complementary to all current clinically-used antibiotics. We used the structural data and coupled this with a genetic approach to pinpoint molecular features in tunicamycin that allowed design of analogues (**TUN-X,X**) that retain anti-microbial activity yet no longer inhibit DPAGT1.

Tunicamycin is synthesized by *Streptomyces chartreusis* but not by *Streptomyces coelicolor*. We have previously cloned and sequenced the tunicamycin biosynthetic cluster (*tun*) from *S. chartreusis* and expressed it heterologously in *S. coelicolor*. A proposed biosynthetic pathway has been based largely on homology of the encoded gene products with proteins of known function, supported by partial *in vitro* studies of TunA and TunF (Wyszynski et al., 2010; Wyszynski et al., 2012). The cluster contains 14 genes, *tunA-N*. In-frame deletion mutations in all 14 of the cloned *tun* genes in *S. coelicolor* (Widdick et al., 2018) revealed interesting responses in that host. Attempts to delete *tunI* and *tunJ*, encoding the components of an ABC transporter, proved difficult – putative deletion mutants arose only after prolonged incubation, suggesting a role for TunIJ in immunity to tunicamycin. Notably, sequencing of the cloned *tun* gene cluster in one of the emergent Δ*tunI* mutants revealed a G-to-A missense suppressor mutation in *tunC.* This mutation would result in a Gly70Asp substitution in the N-acyltransferase that attaches the key (see above) lipid chain of tunicamycin, presumably resulting in loss of enzyme function – vitally, this led to a loss of tunicamycin activity. This pinpointed a key role for the lipid chain moieties in determining the biological activity of tunicamycin.

### Redesign of tunicamycin Lipidation – Creation of TUN-X,X analogues that do not inhibit DPAGT1

These genetic observations suggested a vital dependency of the toxic action of tunicamycin upon its lipid moiety that, in turn, suggested the creation of analogues through the tailoring of the lipidation state of tunicamycin. We designed a semi-synthetic strategy to access systematically ‘lipid-altered’ variants based on the tunicamine scaffold **TUN** that is at the core of tunicamycin (Figure 5A). Large-scale fermentation of *Streptomyces chartreusis* NRRL 3882 (see Supplemental Information SI 2) allowed access to crude tunicamycin on a multi-gram scale; methanol extraction; optimized growth and extraction procedures allowed yields of 42 ± 5 mg per litre of culture (Supplemental Information SI 2). This allowed degradative (Ito et al., 1979) conversion of tunicamycin to its unfunctionalised core scaffold **TUN**. Critically, since the nucleobase of tunicamycin is hydrolytically sensitive, the creation of mixed Boc-imides at positions 10ʹ and 2ʹʹ allowed mild selective deamidation on a gram-scale (see Supplemental Information SI 2). Chemoselective carbodiimide-or uronate-mediated acylation allowed direct lipid-tuning in a systematic, divergent manner through dual modification at 10ʹ-N and/or 2ʹʹ-N, yielding a logically, lipid-altered library of novel analogues, **TUN-X,X** varying in chain length by one carbon, from C7 to C12 (**TUN-7,7** to **TUN-12,12**, Figure 5A).

**Figure 5.**
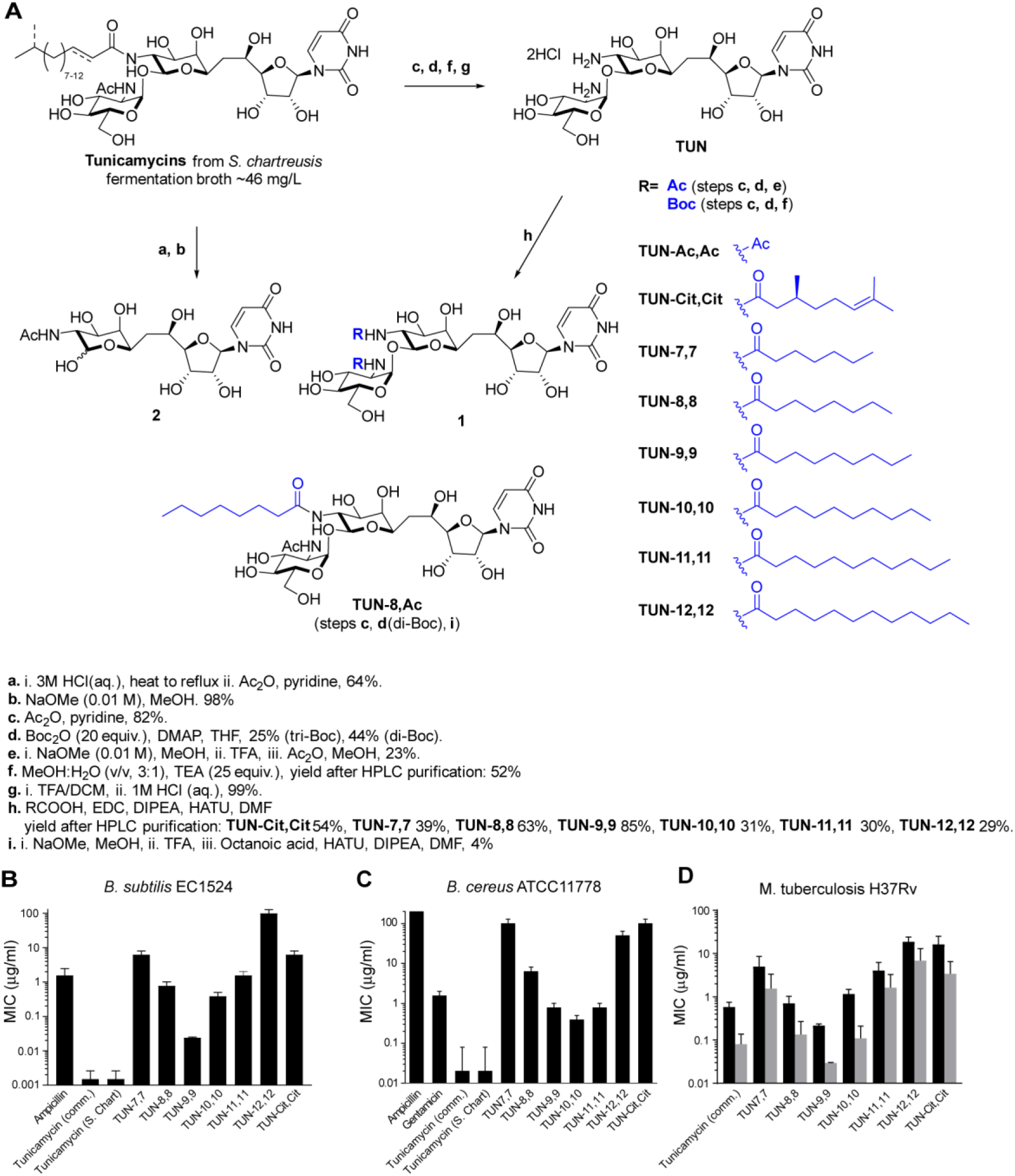
Design, semi-synthetic synthesis and antibacterial effects of TUN-X,X analogues. (A) Semi-synthetic strategy to access TUN mimics. (B-D) MIC values obtained from micro-broth dilution antimicrobial susceptibility tests of (B) *B. subtilis* EC1524, (C) *B. cereus* ATCC11778, (D) *Mtb* H37Rv (ATCC27294) cultured in 7H9/ADC/Tw (black), or GAST/Fe (grey) media.

### TUN Analogues Show Potent Antimicrobial Activity against a Range of Bacteria

We evaluated the analogues (**TUN-7,7, −8,8, −9,9, −10,10, −11,11, −12,12**) for potency against a representative range of Gram-negative and Gram-positive bacteria. First, Kirby-Bauer disc diffusion susceptibility tests (Figure S5A-E), revealed potent activity against the model species *Bacillus subtilis* (EC1524) and opportunistic pathogen *Bacillus cereus* (ATCC 11778). In addition, there was a weaker but significant effect on the pathogenic bacteria *Staphylococcus aureus* (ATCC 29219) and *Pseudomonas aeruginosa* (ATCC 27853); notably the latter is a strain resistant to natural tunicamycin. No activity was seen against *Micrococcus luteus*. Consistent with the critical role of lipid suggested by the genetic experiments, none of the non-lipidated analogues (e.g. **TUN** or **TUN-Ac,Ac**) or synthetic intermediates showed any activity. Vitally, lipid-length (**X** = 7, 8,….12) in the **TUN-X,X** analogues systematically modulated activity; the most potent analogues **TUN-8,8** and **TUN-9,9** were those with C8 and C9 chain lengths.

These promising initial screens of activity, were confirmed through the determination of minimal and half maximal inhibitory concentrations (MIC and IC_50_) and minimal bactericidal concentrations (MBC) by both a micro-broth dilution test and drop plate test, respectively (Figure 5B,C, Figures S5F-J, Table S2). Again, only lipidated variants (tunicamycin and **TUN-X,X**) displayed anti-bacterial activity, with MICs down to 0.02 ± 0.01 µg/ml for **TUN-9,9** against *B. subtilis* and 0.33 ±0.11 µg/ml against *B. cereus*, with **TUN-10,10**.

One of the most pernicious pathogens of global concern http://www.who.int/tb/publications/global_report)/en/) is *Mycobacterium tuberculosis* (*Mtb*), the etiological agent of TB. Testing of the lipid-altered analogues (**TUN-7,7** to **-12,12**) against the pathogenic *Mtb* strain *Mycobacterium tubercu*losis H37Rv again revealed lipid-tuned activity down to the striking MIC values (0.03 ± 0.001 μg/mL in minimal growth medium and 0.22 ± 0.02 μg/mL in rich 7H9-based growth medium) for **TUN-9,9**: some 5-fold more potent than even tunicamycin itself (Figure 5D, Table S3).

### Lipid-altered TUN-X,X analogues are non-toxic to eukaryotic cells

The cytotoxicity of tunicamycin towards eukaryotic cells has until now rendered it unsuitable for clinical use. To probe the effect of lipid-alteration upon such toxicity, we evaluated the effect of analogues **TUN-7,7** to **-12,12** (as well as corresponding synthetic intermediates) on representative human cell lines from the liver (HepG2), kidney (HEK293) and blood (Raji) cells. These were examined both by proliferation dose-response curve and by analysis of morphological or phenotypic changes by microscopy. Consistent with prior observations (Takatsuki et al., 1972), 24 hour incubation with tunicamycin showed both clear cytotoxicity (Figure 6A-C, Figure S6) and morphological changes (Figure 6D, Figure S6B-D).

**Figure 6.**
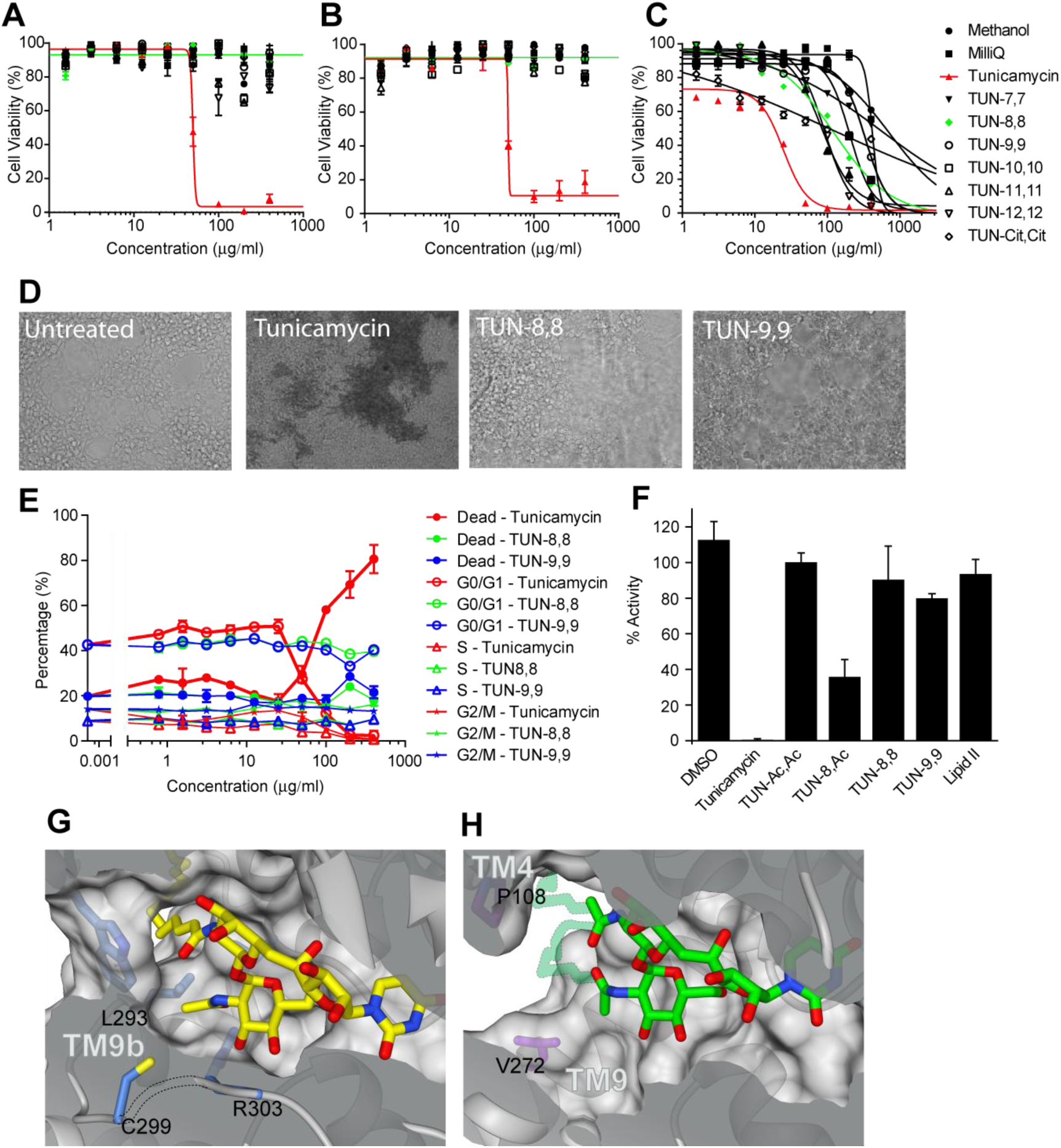
The TUN-X,X analogues are not toxic to cultured human cells HEK293 and HepG2 due to the restrictive tunicamycin-binding site in DPAGT1 compared to MraY. (A-C) Dose response curves from cell proliferation assays with (A) HEK293, (B) HepG2 and (C) Raji cells with tunicamycin and the analogues. (D) Effect of 400 µg/ml tunicamycin, **TUN-8,8** and **TUN-9,9** on the morphology of HEK293 cells. (E) Effects of tunicamycin, **TUN-8,8** and **TUN-9,9** on the cell cycle of HEK293 cells. (F) Effects of tunicamycin and **TUN-X,X** analogues on the DPAGT1 catalytic activity. (G) The DPAGT1 lipid-binding sites, showing the restrictive tunnel with tunicamycin bound. (H) The more open MraY lipid-binding site, with the additional modelled lipid chain shown with lower contrast.

Cell-cycle analysis (Figure 6E and S6A) suggested that cell death coincided with a dramatic decline in G0/1 phase populations with an LC_50_ ~100 μg/ml.

Consistent with a mode of action that requires lipidation for toxicity, all non-lipidated variants (**TUN** core and synthetic intermediates) displayed no significant adverse effects; these variants therefore do not act upon *either* bacteria or mammalian cells in any potent manner.

However, and in contrast to tunicamycin’s toxicity (LD_50_ = 51.25 ± 31.27, 44.74 ± 4.73, 26.82 ±11.46 μg/mL for HEK293, HepG2 and Raji cells, respectively, Figure 6A-C and Table S4) the designed **TUN-X,X** variants **TUN-7,7** to **-12,12**, with their *altered* lipids, showed either mild or negligible toxicity (LD_50_ > 400 μg/mL) towards mammalian cells. Moreover, a high level (>75%) of viable cells with no morphological changes were observed after 24 hours (Figure S6A) when HepG2 or HEK293 cells were incubated with this same high dosage (400 μg/mL). Moreover, no variation in cell cycle was observed, with healthy G0/1 populations being maintained even at the highest concentrations (Figure 6E, Figure S6A).

The mechanistic origin of this reduced toxicity was tested *in vitro* with purified DPAGT1 enzyme. We measured the activity of DPAGT1 in the presence of the **TUN-8,8** and **TUN-9,9** analogues. Whilst native tunicamycin completely inhibited DPAGT1, these analogues had negligible effect on DPAGT1 activity (Figure 6F). This is consistent with the observation that while tunicamycin inhibits the glycosylation of a model protein, it is not affected by **TUN-8,8** or **TUN-9,9** (Figure S6E-G). Notably, synthetic reinstallation of only a *single* C8 lipid into analogue **TUN-8,Ac** restored inhibitory activity towards DPAGT1 (Figure 6F), wholly consistent with our design and the critical role of the second lipid preventing binding to DPAGT1. Given that in patients with CMS, a loss of the activity of one gene, reducing activity by 50%, is not sufficient to cause significant disease, it seems likely that a reduction in activity by <10%, caused by the **TUN-X,X** analogues, during a short-term treatment is unlikely to cause significant toxicity. Together these results confirmed our hypothesis that systematic ‘lipid alteration’ could create tunicamycin analogues in which mammalian cytotoxicity is separated from antibacterial effects for the first time.

### A molecular explanation for the differences in the TUN-X,X lipid analogue binding to DPAGT1 and MraY

Comparison of the structures of the complexes of tunicamycin with DPAGT1 and MraY (Hakulinen et al., 2017; Yoo et al., 2018); this work) gave a clear explanation for the preferential effects of the analogues on MraY over DPAGT1. Overall the MraY tunicamycin binding site has a much more open, shallow surface than in DPAGT1; in the latter the lipid tail is completely enclosed by Trp122 adjacent to the active site (Figure 6G). The open MraY binding site has a disordered loop CL1, a longer TMH9 and a relatively short CL9 region, with only one short α-helix (Figure 1G). In contrast, DPAGT1 has an ordered CL1 which folds over the UDP-binding site. It also has a shorter TMH9, followed by a loop and extended strand (residues Gln292 to Arg306) (Figure 1F), which folds over tunicamycin forming numerous interactions, including those with Arg301, His302, Arg303 (Figure 2G, H). This extended structure is stabilised by its interactions with the rest of the CL9 domain, a feature found only in eukaryotes.

The amino acetyl group on GlcNAc is the attachment site of the second lipid chain in **TUN-X,X** analogues – it occupies clearly distinct environments in the two proteins. In DPAGT1 it is enclosed by the loop at the end of TMH9, and by a tight ‘gating’ cluster of side chains from Trp122, Ile186, Leu293, Cys299 and Arg303 (Figure 6G). By contrast, in MraY, there is a 10 Å gap between Pro108 on TMH4a and Val272 on TMH9, providing ample space for more than one lipid chain to be attached to the amines in **TUN-X,X** analogues (Figures 6G, H).

### Lipid-altered TUNs show efficacy against *Mtb* in mice

The greatly enhanced therapeutic index of the **TUN-X,X** analogues in culture (Table S5) suggested their strong potential application in treating pathogen infections in mammals. Toxicity and efficacy were probed in infection models of *Mtb* both *in cellulo* and *in vivo*. *Mtb* resides in macrophages following infection of mammals. First, as a stringent *in cellulo* test of the ability to treat infection, **TUN-8,8**, **TUN-9,9**, **TUN-10,10** and **TUN-11,11** were used to treat *Mtb*-infected macrophages (Figure 7A) which showed that these analogues were effective at reducing intracellular bacterial burdens by 1- and 2-logs at 1 × and 10 × MIC, respectively. Notably, no toxicity was observed against the host macrophages during treatment of the intracellular infection.

**Figure 7.**
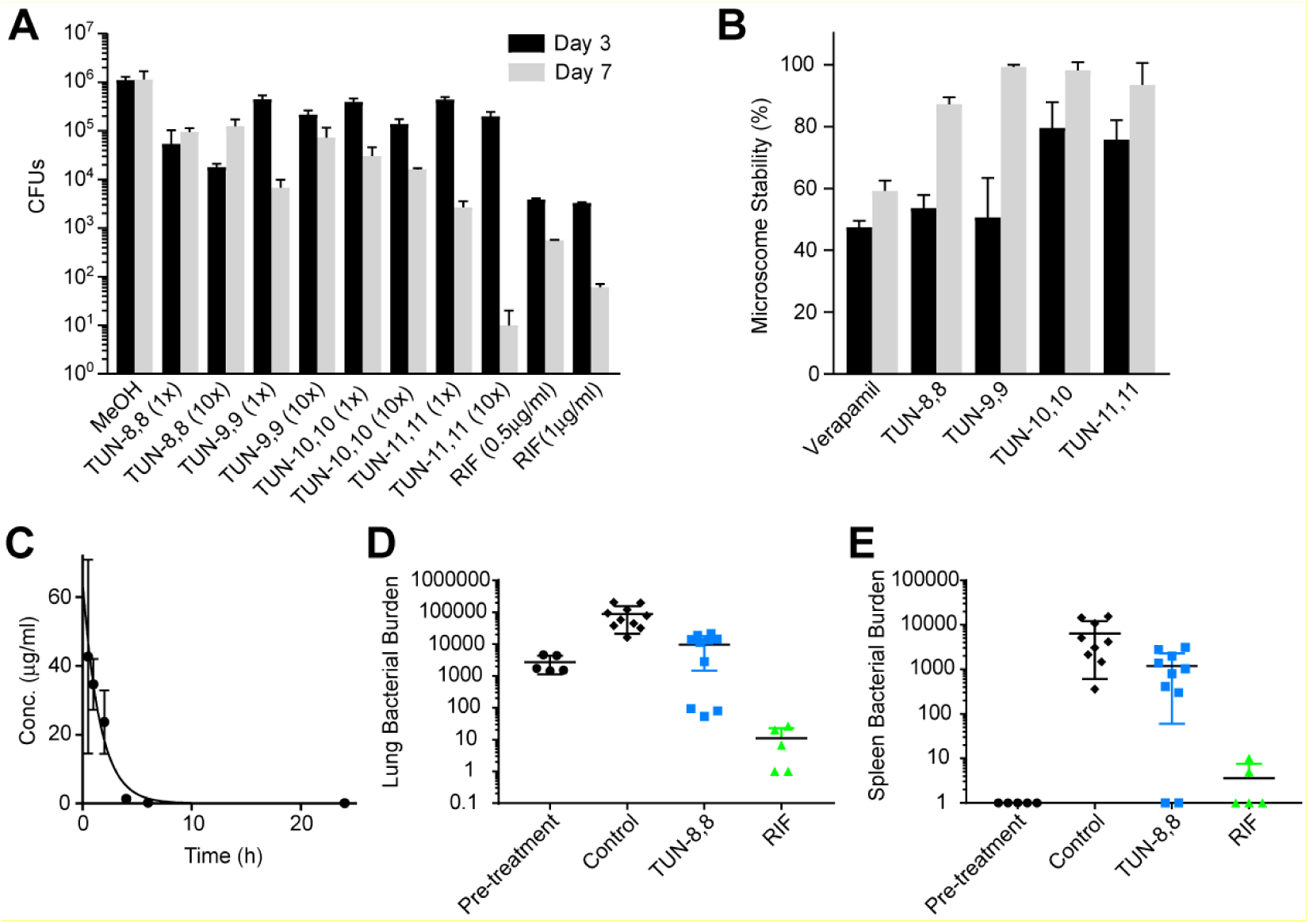
Tunicamycin analogues eradicate *Mtb* during host pathogenesis. (A) Efficacy of tunicamycin analogues in macrophages. Infected J774A.1 macrophages were treated with compounds (1x and 10x MIC for tunicamycin) for 3 or 7 days after which bacterial burdens were enumerated. (B) Microsomal stability of the tunicamycin analogues in human (black) and mice (grey) liver microsomes. (C) Blood serum **TUN-8,8** concentrations after a single intra-peritoneal injection of 30 mg/kg of the analogue. Sixteen mice were dosed and blood collected from the tail vein at indicated times. **TUN-8,8** concentrations in serum were analyzed by LC-MS. (D) Efficacy of **TUN-8,8** in reducing lung bacterial burdens in *Mtb*-infected mice after 2 weeks of treatment. (E) Efficacy of **TUN-8,8** in reducing spleen bacterial burdens in *Mtb*-infected mice following 2 weeks of treatment. Unpaired *t* test for the **TUN-8,8** and vehicle control group p value of 0.0017 in lungs and 0.01 in spleens.

Second, microsomal (human and mouse) stability assessment (Figure. 7B) suggested good metabolic survival of **TUN-8,8** to -**11,11**, which was confirmed by *in vivo* pharmacokinetic determination in mice (Figure 7C). This revealed good bioavailability of **TUN-8,8** following intraperitoneal (*ip*) delivery and blood plasma exposures suggesting efficacious daily dosing. Next, tolerance testing of **TUN-8,8** in uninfected mice (n=5) over 10 days at daily doses of 30 mg.kg^−1^ (*ip*) showed no signs of toxicity, in striking contrast to tunicamycin. Finally, antitubercular activity was demonstrated in *Mtb*-infected mice. Consistent with the results found *in cellulo* (see above), treatment of *Mtb*-infected mice (n=10) over two weeks (10 mg.kg^−1^, *ip*) revealed an almost 10-fold reduction in bacterial burdens in lung (Figure 7D) and a 5-fold reduction in spleen (Figure 7E) compared to mice receiving the vehicle control. Notably, despite the tolerability shown even at 30 mg.kg^−1^, up to 10 days in uninfected animals, clinical signs of toxicity in infected mice precluded any longer term testing beyond 2 weeks. The origins of this toxicity seen in only diseased animals are unclear but suggest that further optimization of **TUN-X,X** analogues and/or their formulation, may be critical to disease-weakened animals. **TUN-X,X** analogues are therefore unoptimized but promising, proof-of-principle, leads rather than, as yet, optimized antibiotic drugs.

## Discussion

The structures of DPAGT1 have allowed us to explain the mechanism of this key enzyme in the major eukaryotic pathway of protein N-glycosylation. We show that missense variants in DPAGT1 associated with CMS and CDG-Ij alter DPAGT1 function via diverse mechanisms. For many cases of milder CMS disease, severely reduced activity from one allele is combined with an allele with a partially reduced activity. In two cases, Val264Gly and Gly160Ser, it appears that errors in splicing that reduce the levels of correct mRNA, are partially compensated by 2-fold increases in enzymatic activity. In CDG-Ij, there is either only one allele producing protein with 20% activity or, alternatively, two alleles producing 5-10% activity, leading to much greater disease severity. In all cases there has to be some active protein present, with a threshold of symptoms and increasing disease severity lying between no disease at 50% activity and severe disease with 5-10% of activity. It is also highly significant that DPAGT1 activity can be increased by point mutations at single sites, suggesting that it may be possible to increase enzymatic activity and/or modulate stability with small molecules, e.g. pharmacological chaperones (Convertino et al., 2016; Sanchez-Fernandez et al., 2016).

DPAGT1 represents an ‘off-target’ for the natural bactericidal agent tunicamycin. Comparison of the human PNPT DPAGT1 and bacterial PNPT MraY structures revealed a gating loop (residues Cys299-Arg303) in DPAGT1 next to where the N-2” atom of tunicamycin binds that is absent in the more open structure of MraY. This difference allowed design of analogues **TUN-X,X** with two lipid chains targeted to bind to MraY, but not to DPAGT1 by virtue of a blocking/gated lipid installed at N-2”. This circumvented the toxicity problem normally observed with tunicamycin. Additive modes of action against other carbohydrate-processing enzymes, such as *Mtb* WecA, *Mtb* TagO/TarO or *P. aeruginosa* chitin synthase, may also be important for the effects of the analogues.

*Mtb* is responsible for ~1.3 million deaths per annum and it is estimated that a third of the world’s population is infected. Its resistance to common antibacterial treatments has necessitated new strategies (Young et al., 2008; Zumla et al., 2013) that has led to the development of specialized and innovative candidate medicines (Modlin and Bloom, 2013), the most potent of which are isoniazid (MIC 0.01-0.04 μg/mL) and rifampicin (MIC 0.015-0.4 μg/mL). We have shown that the **TUN-X,X** analogues are effective in killing *Mtb*, they have much lower toxicity than tunicamycin itself and in mice they reduce the *Mtb* burden by 2 orders of magnitude in 2 weeks. The analogues do show some toxicity in mice over periods of more than 2 weeks in diseased animals (although not in healthy animals), they are still much less toxic than tunicamycin. While these lead versions of the **TUN-X,X** lipid analogues are not as effective as the frontline drugs rifampicin and isoniazid in macrophages and in mice *in vivo*, the details of the effects in mice are often not recapitulated in humans. In addition, we do not yet have data on intracellular uptake in animals, this may be affecting the outcome. MICs show that these compounds are excellent leads for the design of novel antibiotics with a new mechanism of action. These analogues are effective antibacterials, with limited toxicity in human cells and in mice (at least with short term dosing), and suggest a novel approach to development of antibiotics against Gram-positive bacteria.

## Author Contributions

Project design: Y.Y.D., H.W., A.C.W.P., D.B., M.J.B., B.G.D. and E.P.C. Pilot expression studies and baculovirus-infected insect cell production: L.S., S.M.M.M., N.A.B.B. Mutagenesis: K.B., S.R.B. Protein production, structure and function studies: Y.Y.D., A.C., A.C.W.P., E.P.C. DPAGT1 in CMS and exon trapping studies, W.-W.L., K.B., D.B. Mass spectrometry, S.M., C.V.R. Tunicamycin lipid role and effects of analogues on bacteria: H.W., D.A.W., A.S., H.I.B, B.G.D, C.E.B., and M.J.B. Tunicamycin **TUN-X,X** development: H.W., S.A.C., S.H., F.J.W., S.F.R., R.L., W.-M.L., S.S.L., T.M., B.G.D. In vivo experiments: A.S., H.I.B., C.E.B. Project supervision and management: B.G.D., E.P.C. Data analysis and manuscript preparation: Y.Y.D., H.W., A.C.W.P., S.R.B., S.A.C., S.H., D.A.W., H.I.B., S.M., M.J.B., C.E.B., C.V.R., D.B., B.G.D., E.P.C.

## Acknowledgements

Y.Y.D., A.C.W.P., S.R.B., L.S., A.C., S.M.M.M., N.A.B.B. and E.P.C. are members of the SGC, (Charity ref: 1097737) funded by AbbVie, Bayer Pharma AG, Boehringer Ingelheim, the Canada Foundation for Innovation, Genome Canada, GlaxoSmithKline, Janssen, Lilly Canada, Merck & Co., Novartis, the Ontario Ministry of Economic Development and Innovation, Pfizer, São Paulo Research Foundation-FAPESP and Takeda, as well as the Innovative Medicines Initiative Joint Undertaking ULTRA-DD grant 115766 and the Wellcome Trust106169/Z/14/Z. B.G.D is funded by The Gates Foundation, BBSRC, EPSRC, Wellcome Trust and Evotec Ltd. S.A.C is funded by the Wellcome Trust 110270/A/15/Z. S.M. and C.V.R.: MRC programme grant MR/N020413/1. DB: MRC Grant MR/M006824, WT Strategic Award WT084655MA. D.W., and M.J.B.: BBSRC Grants BB/J006637/1, BB/J009725/1 and BB/J004561/1. This work was funded, in part, by the Intramural Research Program of NIAID. We thank Diamond Light Source Ltd and their staff for access to the macromolecular crystallography beamlines. We thank all member of the SGC Biotech team, including Claire Strain-Damerell, Kasia Kupinska, Dong Wang, Katie Ellis, Octavia Borkowska and Rod Chalk. We thank all members and ex-members of the SGC IMP1 group, including Annamaria Tessitore, Liang Dong, Berenice Rotty, Andrew Quigley, Mariana Grieben and Chitra Shintre. We thank George Berridge for help developing assays. Brian Marsden, David Damerell, James Bray, James Crowe and Chris Sluman for computing and bioinformatics support, and Frank von Delft and Tobias Krojer for crystallography infrastructure support. We thank Danielle Weiner and Michelle Sutphin for help with treatment of animals and Michael Goodwin for help with PK analysis. We thank Prof. Seok-Yong Lee and his group for initial technical and scientific exchanges.

## Supplemental figures

**Figure S1.**
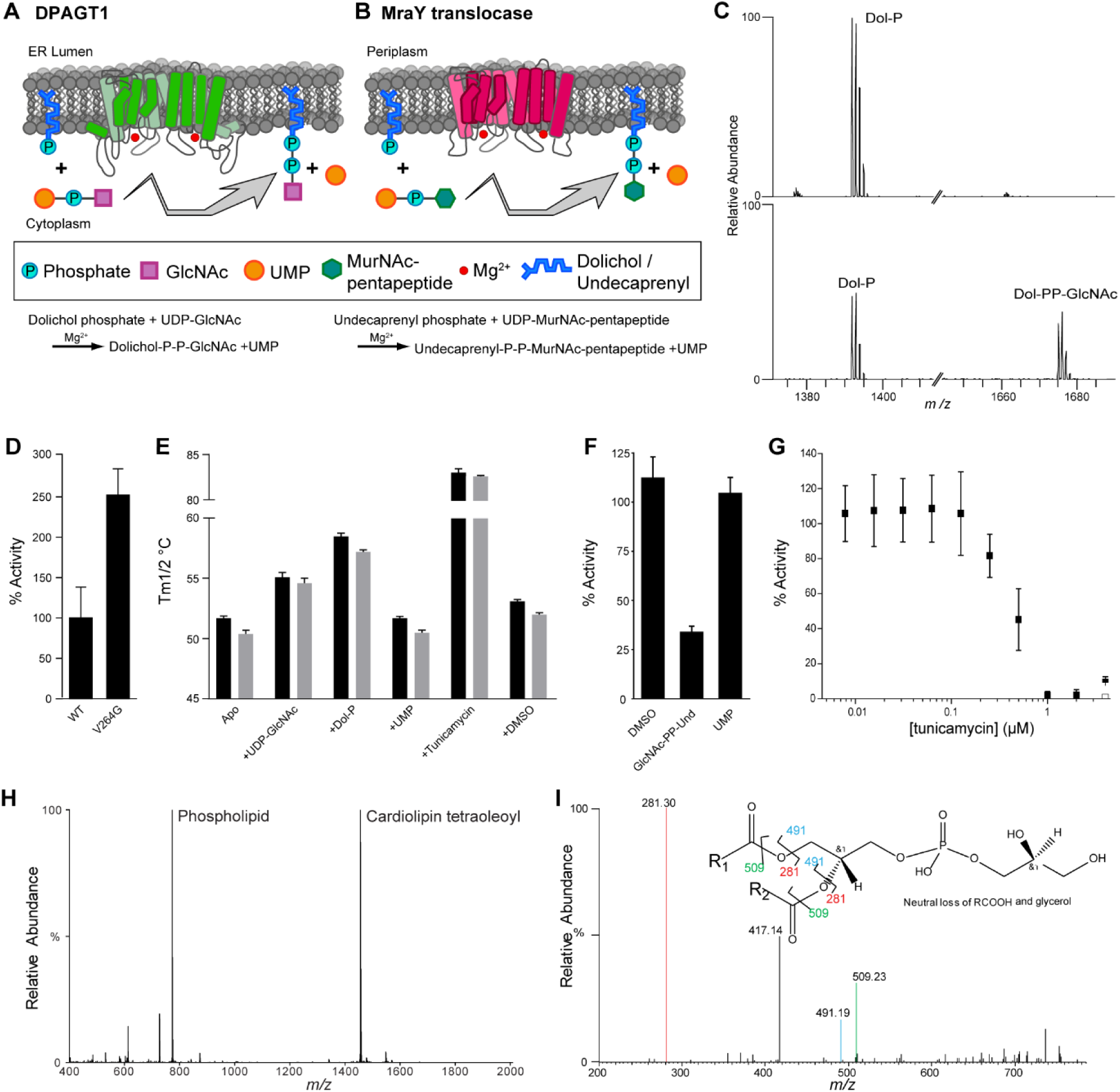
Biochemical and biophysical characterisation of DPAGT1, Related to Figure 1. **(A)** Cartoon of DPAGT1 showing the reaction it performs. **(B)** Cartoon of MraY showing the reaction it performs. **(C)** The identity of the substrate Dol-P and the product, GlcNAc-PP-Dol, was confirmed by mass spectrometry. Top spectra is DPAGT1 incubated with Dol-P only, bottom spectra is DPAGT1 incubated with both Dol-P and UDP-GlcNAc. **(D)** Comparison of the catalytic activity of DPAGT1 WT and Val264Gly mutant protein. **(E)** The thermostability of DPAGT1 WT (black) and Val264Gly (grey) mutant proteins tested using label free differential scanning fluorimetry. The effects of addition of the substrates Dol-P and UDP-GlcNAc, and the inhibitor tunicamycin on thermostability of DPAGT1 were also tested. **(F)** Product inhibition was observed with the product analogue GlcNAc-PP-Und, but not with UMP. **(G)** DPAGT1 is completely inhibited by a 1:1 ratio of tunicamycin:protein. (H) Lipidomics analysis of OGNG purified DPAGT1 showed the presence of co-purified phospholipid in addition to the supplemented cardiolipin associated with the protein. **(I)** The presence of phosphatidylglycerol is confirmed by tandem mass spectrum of the most intense phospholipid in the lipidomics analysis.

**Figure S2.**
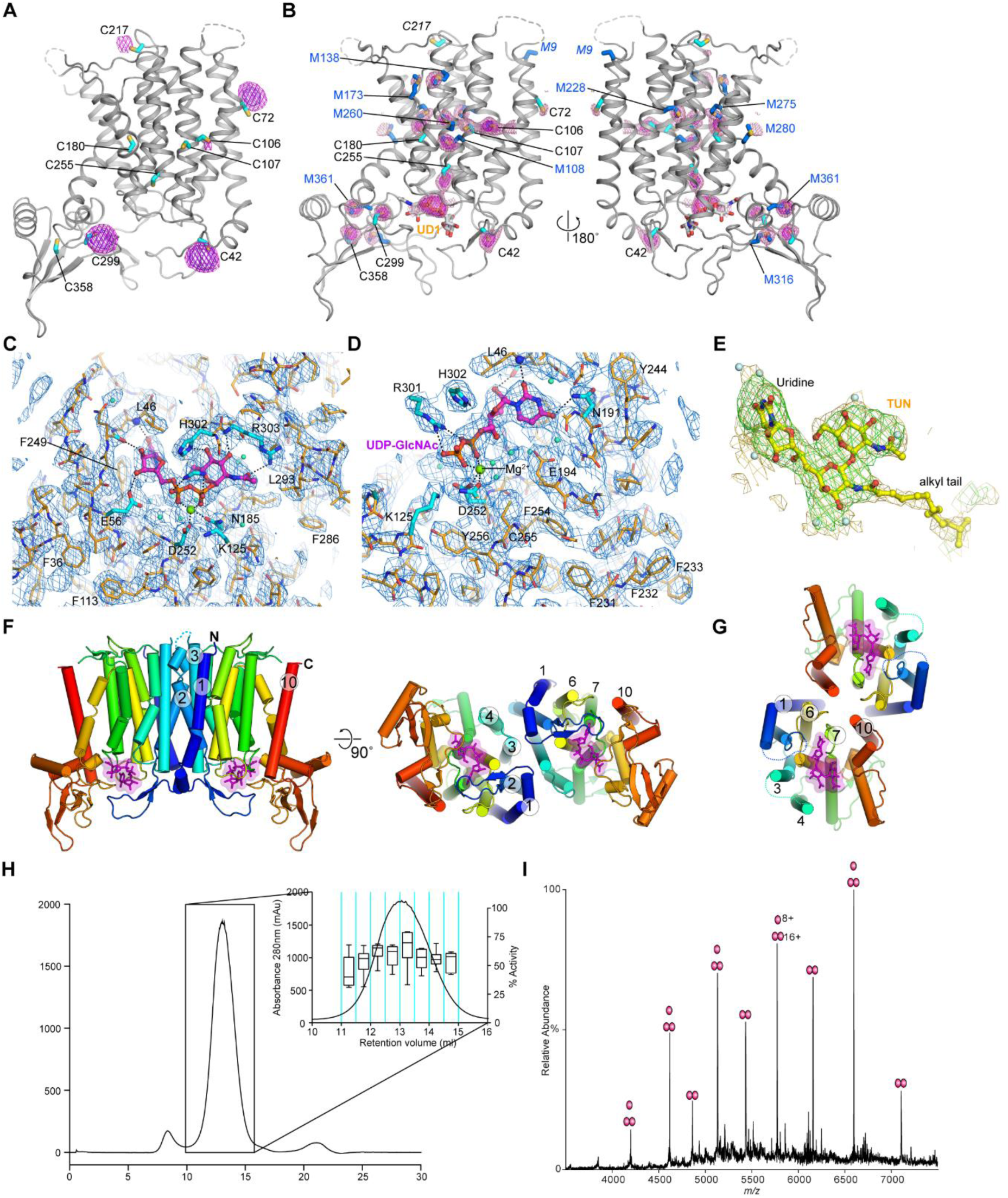
Electron density maps, heavy atom phasing and dimer interface analysis for the structure of dimeric WT and Gly264Val mutant of DPAGT1 solved at resolutions up to 3.2Å. Related to figure 2. (A) Hg bound sites from a soaked crystal. 6 Å anomalous difference Fourier map calculated from a dataset collected from a crystal soaked with EMTS is contoured at 5σ (purple) and 3σ (magenta mesh) and overlaid on the final model. Cysteine positions are highlighted by cyan sticks. Labelling of five of the nine ordered cysteines (Cys42, Cys72, Cys106 (weak), Cys217 and Cys299) is observed under the soaking conditions used. (B) S-SAD peaks. The PHASER-EP log-likelihood map after anomalous model completion with sulphur atoms are shown overlaid on a cartoon representation of the final UDP-GlcNAc complex. The positions of cysteine (cyan sticks) and methionine (blue sticks) residues are highlighted. The map is contoured at 4.5σ (magenta) and 2.5σ (pink mesh). Peaks are observed for 16 sulphur atoms (out of a total of 18 possible ordered sulphurs) and the pyrophosphate of the UDP-GlcNAc is also resolved. (C, D) Two views of final 2F_o_-F_c_ AUTOBUSTER electron density map around the active site in the UDP-GlcNAc complex. The map has been sharpened using a *B*-factor of −100Å^2^ in COOT and is contoured at 1.5σ and overlaid on the final model. UDP-GlcNAc is shown in ball-and-stick form (carbon-magenta, oxygen-red, phosphorus-orange, nitrogen-blue). (E) Omit F_o_-F_c_ electron density map for tunicamycin. The omit difference density is contoured at 2.5σ (green mesh) and a sharpened omit F_o_-F_c_ density map (*B*=-100Å^2^) is contoured at 2σ and overlaid on the final tunicamycin coordinates. Density for tunicamycin’s alkyl tail is not as well resolved as the TUN core. (F-G) Comparison of dimer organisation in DPAGT1 (F) and unbound MraY (PDB: 5JNQ) (G) in crystals, indicating that DPAGT1 and MraY are both ‘head-to’head’ dimers in their respective crystals, but the dimer interfaces are unrelated. (H) Size exclusion chromatography from a DPAGT1 purification with activity data per fraction per unit of protein shown as box plot, indicating that there is no difference in the activity of DPAGT1 across the peak. (I) Native mass spectrometry confirms that DPAGT1 is a mixture of monomers and dimers.

**Figure S3.**
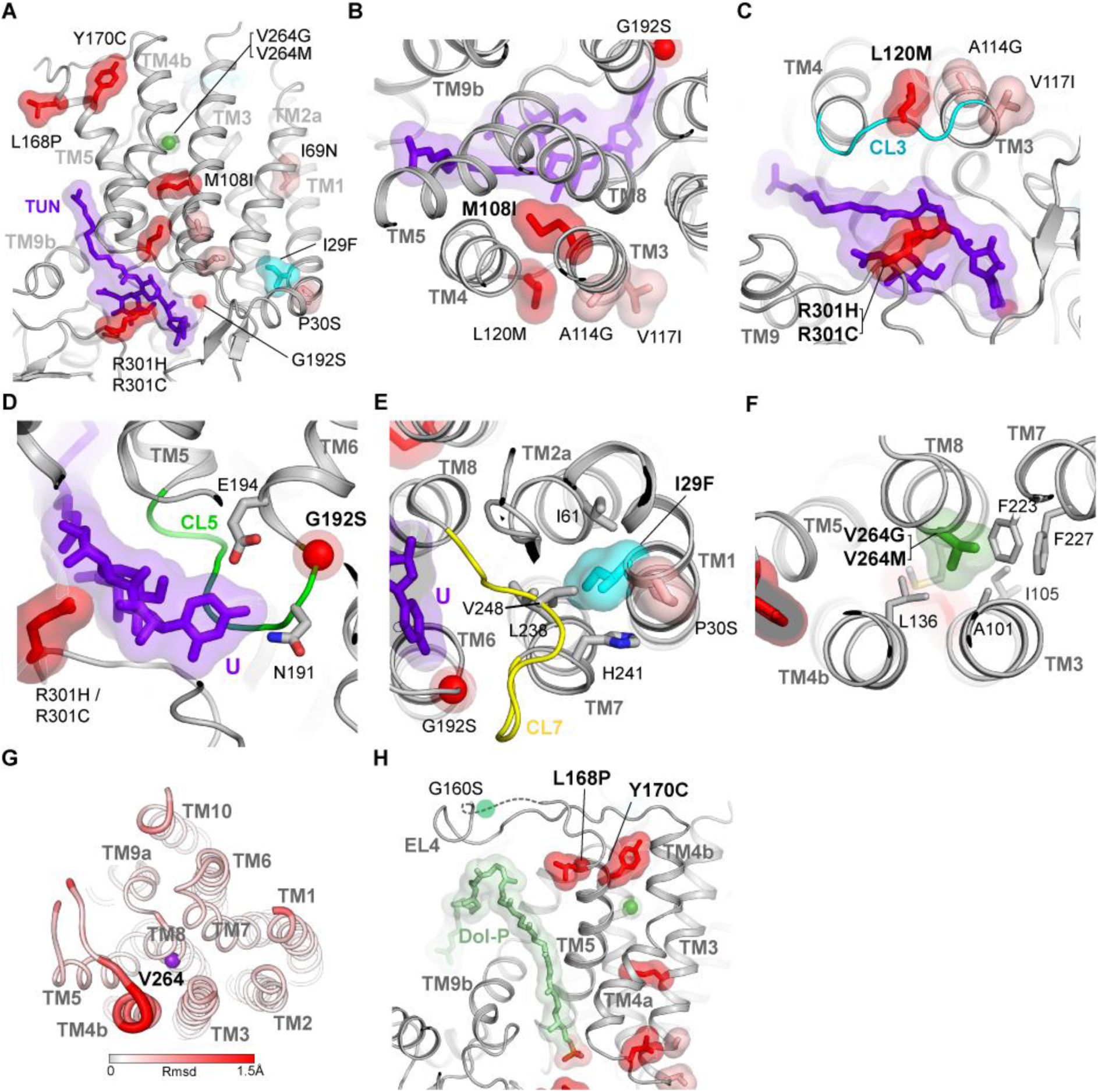
Local structural environment of DPAGT1 CMS and CDG-Ij mutation sites, Related to figure 4. (A) Overview of missense mutation sites mapped onto the DPAGT1 / tunicamycin complex. Colouring of mutation sites is as described in Figure 4. (B) TM3 / TM4 cluster centred on M108. (C) Cytoplasmic face of active site showing CL3 and CL9 loops. (D) G192 lies in CL5 loop adjacent to binding pocket for uridine group of UDP-GlcNAc. (E) Location of a missense site Pro30Ser that impacts on protein stability. (F) Environment of Val264 in wild type unbound structure. TM3 and TM4, along with TM2, form the dimer interface (see Figure S2F). (G) Global conformational changes induced by Val264Gly mutation. Schematic representation of the rms deviation (rmsd) in mainchain atomic positions between the WT and Val264Gly unbound structures. The thickness and colour of the tube reflects the magnitude of the rmsd between the two structures. The main difference is localised at the C-terminal end of TM4b which tilts in towards TM8 in the Val264Gly structure due to closer packing of Leu136 with Gly264. (H) Leu168 & Tyr170 are located at the N-terminus of TM5 at one end of the predicted dolichol phosphate binding groove adjacent to TM5 and TM9. For illustrative purposes, an extended Dol-P lipid (pale green) has been modelled based on the position of the lipid tail of tunicamycin.

**Figure S4.**
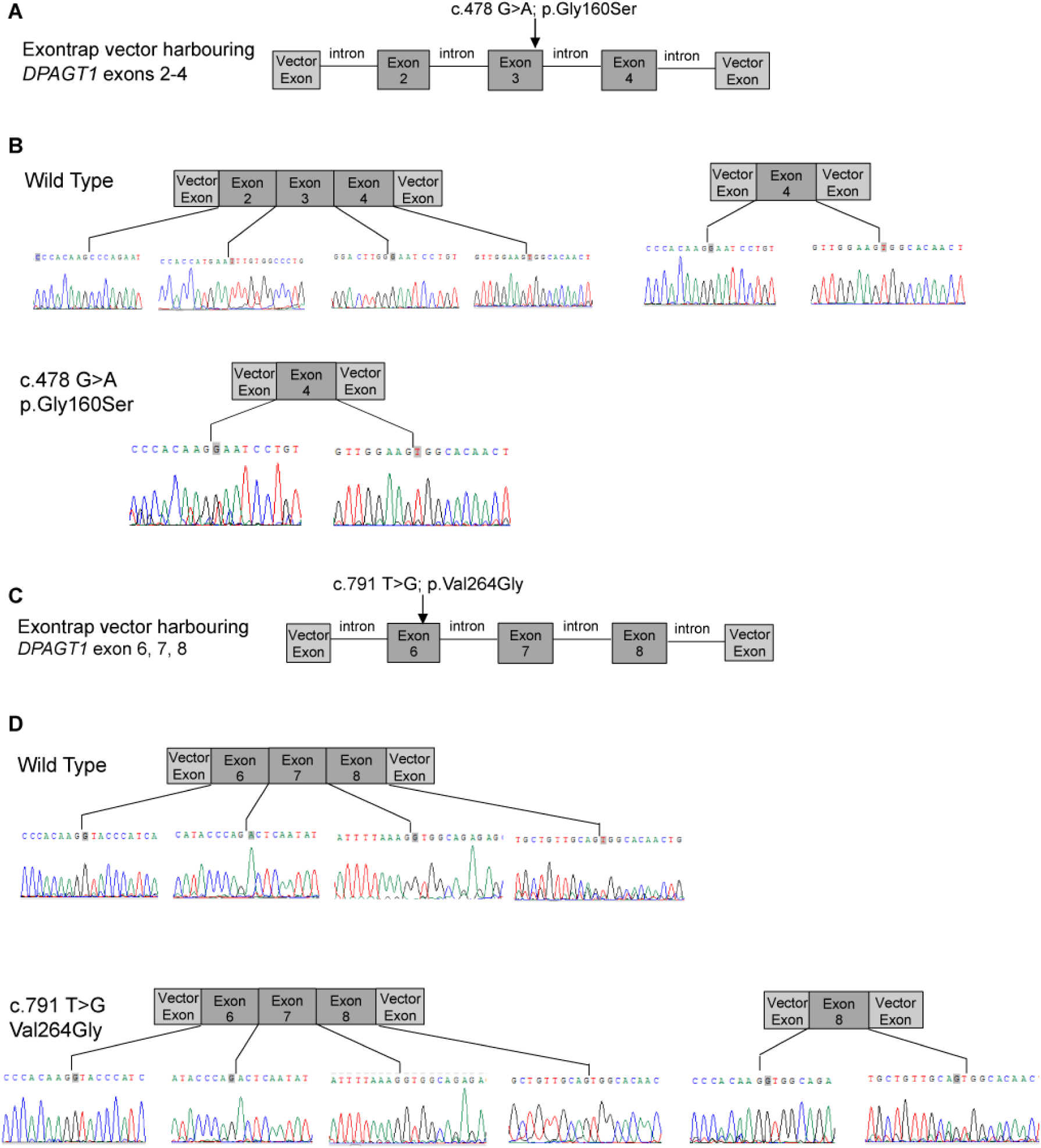
The c.478G>A; Gly160Ser and c.791T>G; Val264Gly mutations are associated with exon splicing errors, resulting in the loss of exons 2 and 3 from the transcript harbouring c.478G>A; Gly160Ser and the loss of exons 6 and 7 from the transcript harbouring c.791T>G; Val264Gly. Relates to figure 4. (A) Schematic of pET01 exon trap vector with DPAGT1 exons 2-4 inserted showing the location within the genomic sequence of c.478G>A; Gly160Ser. (B) Sequencing data and schematic diagrams showing aberrant splicing that results from genomic sequence harbouring the c.478G>A; Gly160Ser variant. RT-PCR on RNA produced in TE671 muscle cell line following transfection with the ‘exon trap’ vector gave wild type RNA sequence, but also some transcripts that excluded exons 2 and 3. When the genomic sequence contained the c.478G>A; Gly160Ser variant was transfected only RNA missing exons 2 and 3 was detected. (C) Schematic of pET01 exon trap vector with DPAGT1 exons 6-8 inserted showing the location within the genomic sequence of the c.791T>G; Val264Gly variant. (D) Sequencing data and schematic diagrams showing sequences obtained following RT-PCR on TE671 cells transfected with the ‘exon trap’ vector containing human genomic DNA that is either wild type or has the c.791T>G; Val264Gly variant. Wild type sequence generated RNA harbouring only exons 6, 7 and 8, as shown. The c.791T>G; Val264Gly variant generated some RNA transcripts containing exons 6, 7 and 8 (as for wild type) but for the majority of transcripts exons 6 and 7 were excluded and only exon 8 was present.

**Figure S5.**
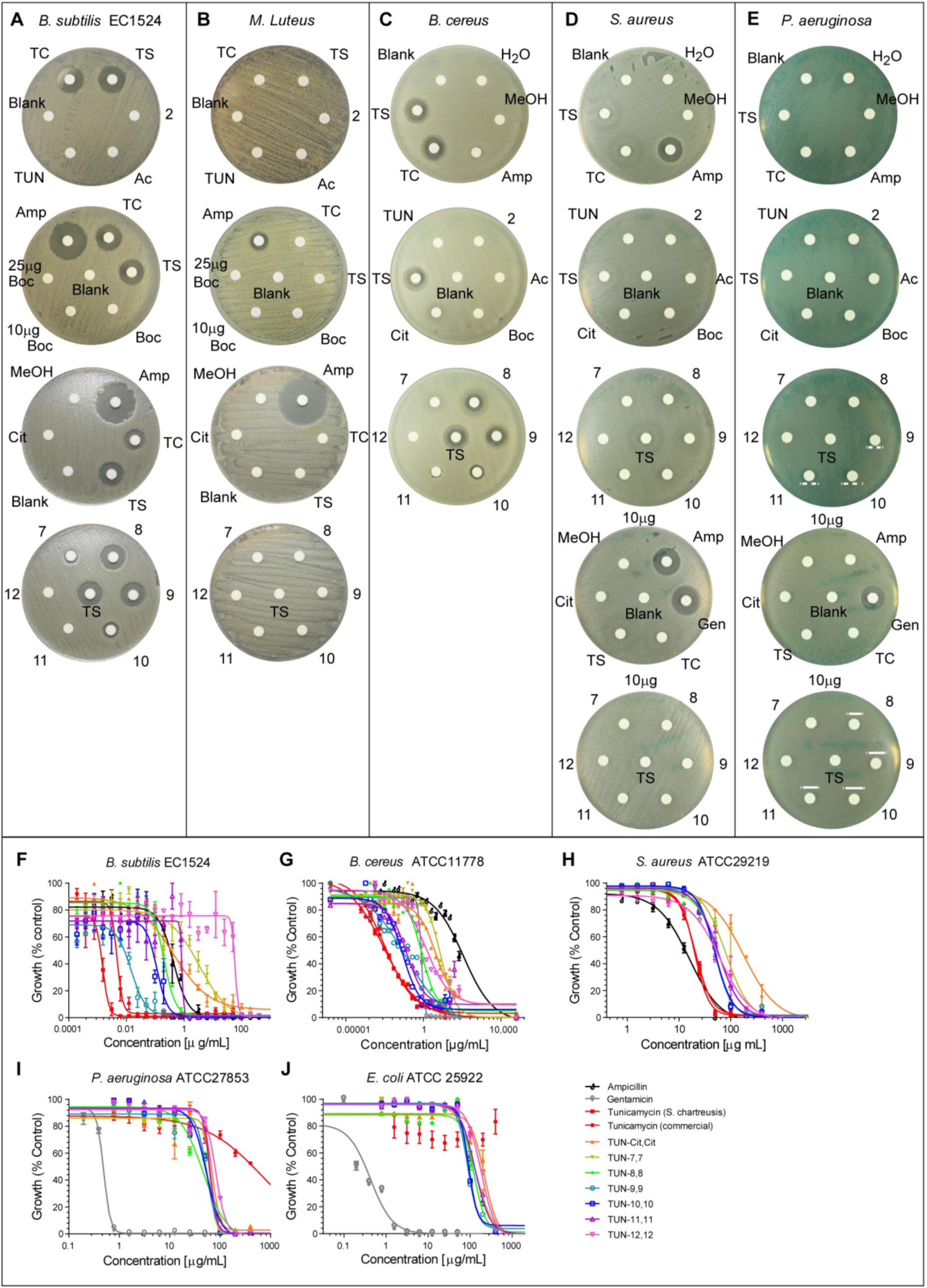
Kirby-Bauer disc diffusion tests and dose response curves used for MIC, MBC and IC_50_ determination for tunicamycin and the TUN-X,X analogues against several bacterial strains. Related to figure 5. Kirby-Bauer disc diffusion tests for the TUN analogues against bacterial strains (A) *B. subtilis* EC 1524, (B) *M. luteus*, (C) *B. cereus* ATCC 11778, (D) *S. aureus* ATCC 29219 and (E) *P. aerugino*sa ATCC 27853. Discs impregnated with 5 µg, unless otherwise indicated, of the compound were laid onto plates with lawns of bacteria. The compounds are labelled: TC: commercial tunicamycin, TS: tunicamycin from *S. chartreusis*, 2: (**2**) N-acetyl tunicamine, Ac: **TUN-Ac,Ac**, Boc: **TUN-Boc,Boc**, 7: **TUN-7,7**, 8: **TUN-8,8**, 9: **TUN-9,9**, 10: **TUN-10,10**, 11: **TUN-11,11**, 12: **TUN-12,12**, Cit: **TUN-Cit,Cit**, Amp: ampicillin, Gen: gentamycin. Dose response curves for tunicamycin and the analogues with (F) *B. subtilis* EC1524. (G) *B. cereus* ATCC11778. (H) *S. aureus* ATCC29219. (I) *P. aerugino*sa ATCC27853 and (J) *E. coli* ATCC25922. These dose-response curves were generated by Prism 6.0 software by plotting percent growth (normalised OD_600_ values) vs. logarithmic scale of the concentrations. The data shown are mean ± SEM errors of three independent experiments. See table S3 for the MIC, MBC and IC_50_ values.

**Figure S6.**
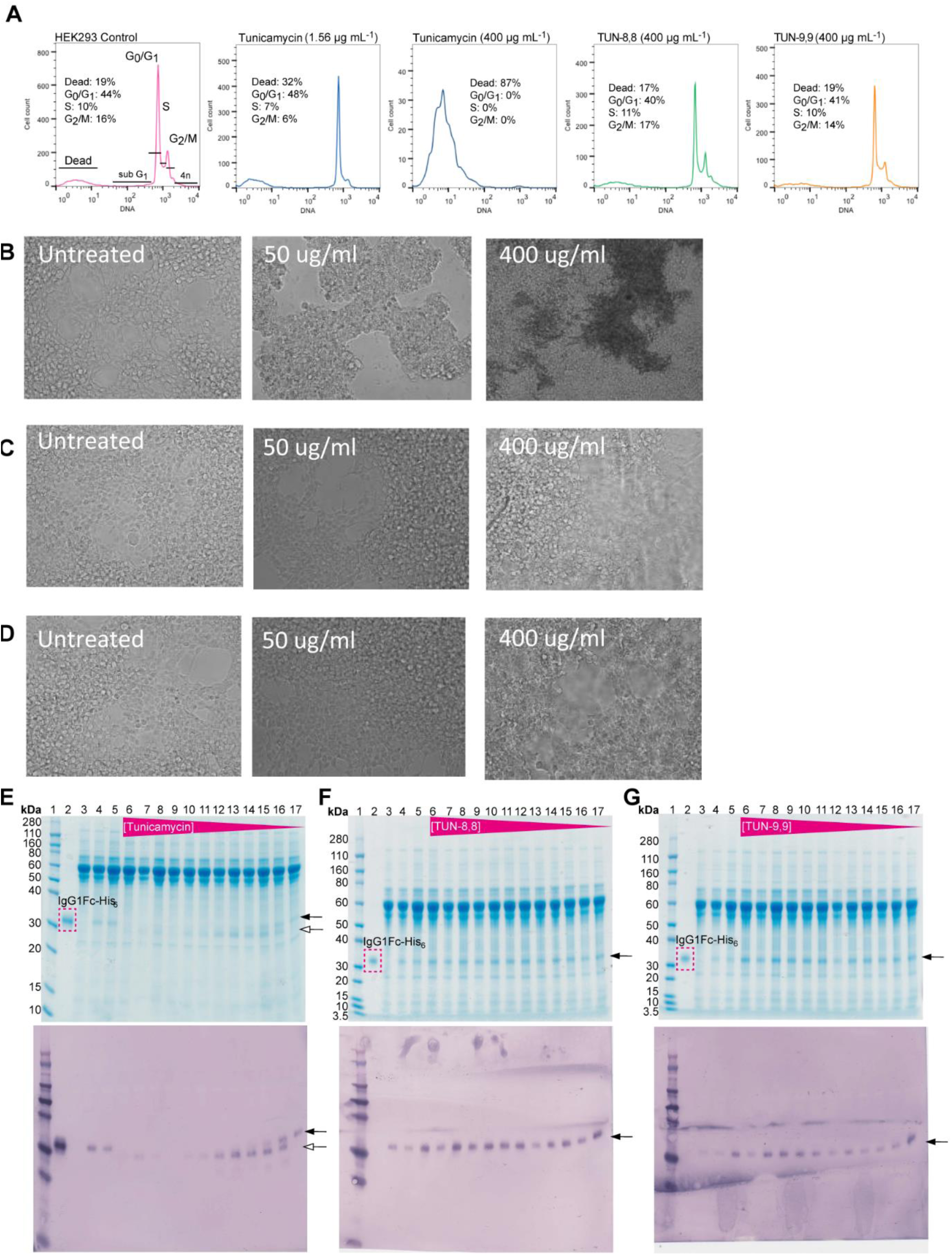
Effect of TUN- and analogues on HEK293 cells, related to figure 6. **(A)** Cell cycle analysis at 24 hours. (B - D) Cell morphology with (B) tunicamycin, (C) **TUN-8,8** and (D) **TUN-9,9**. (E-G) Effects of (E) tunicamycin, (F) **TUN-8,8** and (G) **TUN-9,9** on glycosylation of a model protein – IgG1Fc-His^6^. Black arrow indicates glycosylated protein, white arrow indicates unglycosylated protein.

## Supplemental tables

**Table S1:**
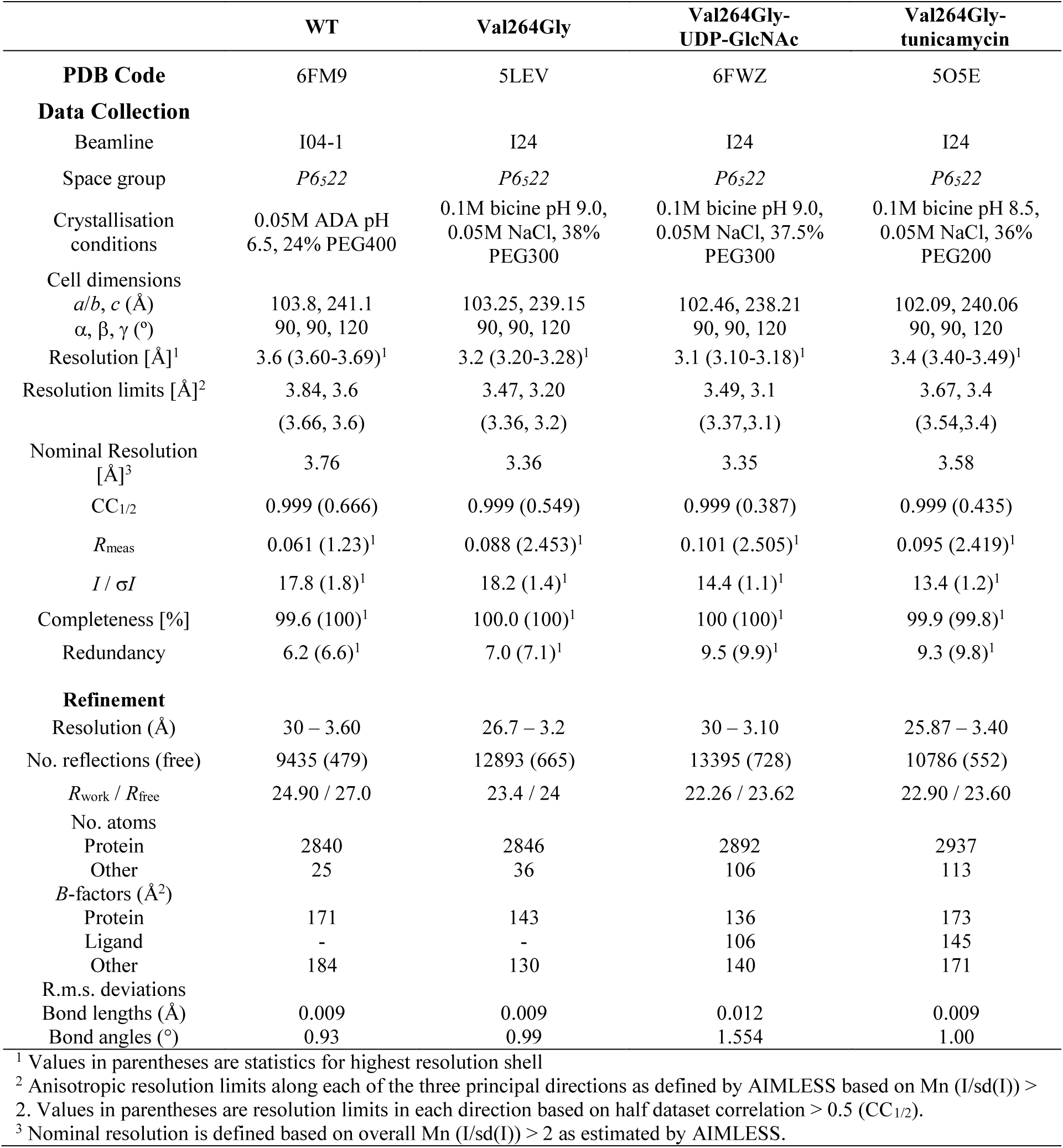
Data collection, phasing and refinement statistics, related to Figures 1 and 2

|  | WT | Val264Gly | Val264Gly-UDP-GlcNAc | Val264Gly-<br>tunicamycin |
| --- | --- | --- | --- | --- |
| **PDB Code** | 6FM9 | 5LEV | 6FWZ | 5O5E |
| **Data Collection** |  |  |  |  |
| Beamline | I04-1 | I24 | I24 | I24 |
| Space group | *P*6<sub>5</sub>22 | *P*6<sub>5</sub>22 | *P*6<sub>5</sub>22 | *P*6<sub>5</sub>22 |
| Crystallisation conditions | 0.05M ADA pH 6.5, 24% PEG400 | 0.1M bicine pH 9.0, 0.05M NaCl, 38% PEG300 | 0.1M bicine pH 9.0, 0.05M NaCl, 37.5% PEG300 | 0.1M bicine pH 8.5, 0.05M NaCl, 36% PEG200 |
| Cell dimensions |  |  |  |  |
| *a/b, c* (Å) | 103.8, 241.1 | 103.25, 239.15 | 102.46, 238.21 | 102.09, 240.06 |
| $\alpha, \beta, \gamma$ (°) | 90, 90, 120 | 90, 90, 120 | 90, 90, 120 | 90, 90, 120 |
| Resolution [Å]<sup>1</sup> | 3.6 (3.60-3.69)<sup>1</sup> | 3.2 (3.20-3.28)<sup>1</sup> | 3.1 (3.10-3.18)<sup>1</sup> | 3.4 (3.40-3.49)<sup>1</sup> |
| Resolution limits [Å]<sup>2</sup> | 3.84, 3.6<br>(3.66, 3.6) | 3.47, 3.20<br>(3.36, 3.2) | 3.49, 3.1<br>(3.37, 3.1) | 3.67, 3.4<br>(3.54, 3.4) |
| Nominal Resolution [Å]<sup>3</sup> | 3.76 | 3.36 | 3.35 | 3.58 |
| CC<sub>1/2</sub> | 0.999 (0.666) | 0.999 (0.549) | 0.999 (0.387) | 0.999 (0.435) |
| *R*<sub>meas</sub> | 0.061 (1.23)<sup>1</sup> | 0.088 (2.453)<sup>1</sup> | 0.101 (2.505)<sup>1</sup> | 0.095 (2.419)<sup>1</sup> |
| *I* / σ*I* | 17.8 (1.8)<sup>1</sup> | 18.2 (1.4)<sup>1</sup> | 14.4 (1.1)<sup>1</sup> | 13.4 (1.2)<sup>1</sup> |
| Completeness [%] | 99.6 (100)<sup>1</sup> | 100.0 (100)<sup>1</sup> | 100 (100)<sup>1</sup> | 99.9 (99.8)<sup>1</sup> |
| Redundancy | 6.2 (6.6)<sup>1</sup> | 7.0 (7.1)<sup>1</sup> | 9.5 (9.9)<sup>1</sup> | 9.3 (9.8)<sup>1</sup> |
| **Refinement** |  |  |  |  |
| Resolution (Å) | 30 – 3.60 | 26.7 – 3.2 | 30 – 3.10 | 25.87 – 3.40 |
| No. reflections (free) | 9435 (479) | 12893 (665) | 13395 (728) | 10786 (552) |
| *R*<sub>work</sub> / *R*<sub>free</sub> | 24.90 / 27.0 | 23.4 / 24 | 22.26 / 23.62 | 22.90 / 23.60 |
| No. atoms |  |  |  |  |
| Protein | 2840 | 2846 | 2892 | 2937 |
| Other | 25 | 36 | 106 | 113 |
| *B*-factors (Å<sup>2</sup>) |  |  |  |  |
| Protein | 171 | 143 | 136 | 173 |
| Ligand | - | - | 106 | 145 |
| Other | 184 | 130 | 140 | 171 |
| R.m.s. deviations |  |  |  |  |
| Bond lengths (Å) | 0.009 | 0.009 | 0.012 | 0.009 |
| Bond angles (°) | 0.93 | 0.99 | 1.554 | 1.00 |

<sup>1</sup> Values in parentheses are statistics for highest resolution shell
<sup>2</sup> Anisotropic resolution limits along each of the three principal directions as defined by AIMLESS based on Mn (I/sd(I)) > 2. Values in parentheses are resolution limits in each direction based on half dataset correlation > 0.5 (CC<sub>1/2</sub>).
<sup>3</sup> Nominal resolution is defined based on overall Mn (I/sd(I)) > 2 as estimated by AIMLESS.

**Table S2.** Table S2. Anti-microbial susceptibility values of tunicamycin and the TUN-X,X analogues against various bacterial strains.

| *MIC (µg/mL)* |  |  |  |  |  |  |  |  |
| --- | --- | --- | --- | --- | --- | --- | --- | --- |
|  | Tunicamycin | TUN-cit,cit | TUN<br>-7,7 | TUN-8,8 | TUN-9,9 | TUN-10,10 | TUN-11,11 | TUN-12,12 |
| *B. subtilis* | 0.0018 ±0.0011 | 5.2 ±1.8 | 5.2 ±1.8 | 0.65 ±0.22 | 0.02 ±0.01 | 0.33 ±0.11 | 1.3 ±0.45 | 83 ±29 |
| *M. luteus* | - | - | - | - | - | - | - | - |
| *B. cereus* | 0.17 ±0.06 | 83 ±29 | 83 ±29 | 5.2 ±1.8 | 0.65 ±0.23 | 0.33 ±0.11 | 0.65 ±0.23 | 42 ±14 |
| *S. aureus* | 42 ±14 | >400 | 170 ±60 | 170 ±60 | 83 ±29 | 83 ±29 | 170 ±60 | 170 ±60 |
| *P. aeruginosa* | >400 | 170 ±60 | 83 ±29 | 83 ±29 | 83 ±29 | 83 ±29 | 83 ±29 | 170 ±6 |
| *E. coli* | >400 | >400 | 330 ±120 | 330 ±120 | 330 ±120 | 330 ±120 | 330 ±120 | 330 ±120 |

| *MBC (µg/mL)* |  |  |  |  |  |  |  |  |
| --- | --- | --- | --- | --- | --- | --- | --- | --- |
|  | Tunicamycin | TUN-cit,cit | TUN-7,7 | TUN<br>-8,8 | TUN-9,9 | TUN-10,10 | TUN-11,11 | TUN-12,12 |
| *B. subtilis* | >0.00.12 | >200 | >50 | >0.78 | >0.05 | >0.39 | >1.56 | >200 |
| *M. luteus* | - | - | - | - | - | - | - | - |
| *B. cereus* | 25 | >200 | >200 | >50 | >25 | 12.5 | >25 | >200 |
| *S. aureus* | >200 | >200 | >400 | 400 | 400 | 400 | 400 | >400 |
| *P. aeruginosa* | >400 | >400 | >400 | >400 | 400 | 400 | 400 | >400 |
| *E. coli* | >400 | 400 | >400 | 400 | 400 | 400 | 400 | >400 |

| *IC<sub>50</sub> (µg/mL)* |  |  |  |  |  |  |  |  |
| --- | --- | --- | --- | --- | --- | --- | --- | --- |
|  | Tunicamycin | TUN-cit,cit | TUN-7,7 | TUN-8,8 | TUN-9,9 | TUN-10,10 | TUN-11,11 | TUN-12,12 |
| *B. subtilis* | 0.00018<br>±0.00006 | 0.36 ±0.12 | 3.12 ±1.04 | 0.20 ±<br>0.07 | 0.015<br>±0.005 | 0.12 ±0.04 | 0.85 ±0.28 | 58.92<br>±19.64 |
| *M. luteus* | - | - | - | - | - | - | - | - |
| *B. cereus* | 0.0029<br>±0.0010 | 6.29 ±2.10 | 13.33<br>±4.45 | 0.78<br>±0.26 | 0.072<br>±0.024 | 0.046<br>±0.015 | 0.088<br>±0.029 | 1.40 ±0.47 |
| *S. aureus* | 20.78 ±6.93 | 176.70<br>±58.90 | 80.49<br>±26.83 | 58.36<br>±19.45 | 48.92<br>±16.31 | 49.29<br>±16.43 | 58.03<br>±19.34 | 61.88<br>±20.63 |
| *P. aeruginosa* | >400 | 70.29<br>±23.43 | 62.44<br>±20.81 | 46.57<br>±15.52 | 54.98<br>±18.32 | 53.46<br>±17.82 | 63.17<br>±21.06 | 81.74<br>±27.31 |
| *E. coli* | >400 | 217.90<br>±72.63 | 131.20<br>±43.73 | 122.70<br>±40.90 | 87.06<br>±29.02 | 91.63<br>±30.54 | 131.10<br>±43.70 | 181.60<br>±60.53 |

**Table S3.** *Mtb* MIC values with average ± Std. Dev.

| *Compound* | *1-week MIC in 7H9/ADC/Tw*<br>*(ug/mL)* | *1-week MIC in GAST/Fe (ug/mL)* |
| --- | --- | --- |
| MilliQ water | No inhibition | No inhibition |
| Methanol | No inhibition | No inhibition |
| Tunicamycin | $0.6 \pm 0.2$ | $0.08 \pm 0.06$ |
| (2) | $>60$ | $42.5 \pm 3.5$ |
| TUN-Ac,Ac | $\geq 60$ | $21.3 \pm 1.8$ |
| TUN | $>60$ | $>60$ |
| TUN-Boc,Boc | $>60$ | $>60$ |
| TUN-Cit,Cit | $16.3 \pm 8.8$ | $3.4 \pm 3.1$ |
| TUN-7,7 | $5.0 \pm 3.5$ | $1.6 \pm 1.8$ |
| TUN-8,8 | $0.7 \pm 3.3$ | $0.14 \pm 0.1$ |
| TUN-9,9 | $0.2 \pm 0.02$ | $0.03 \pm 0.001$ |
| TUN-10,10 | $1.2 \pm 0.3$ | $0.1 \pm 0.1$ |
| TUN-11,11 | $4.1 \pm 2.2$ | $1.6 \pm 1.6$ |
| TUN-12,12 | $18.7 \pm 5.4$ | $6.9 \pm 6.2$ |

**Table S4.** Assessing toxicity of TUN and the analogues in HEK293, HepG2 and Raji cells. LD_50_ calculated from cells cultured in liquid media.

| *Compound* | LD<sub>50</sub> values (µg mL<sup>-1</sup>), Avg. ± Std. Dev. |  |  |
| --- | --- | --- | --- |
|  | HEK293 | HepG2 | Raji |
| MilliQ Water | N/A | N/A | N/A |
| Methanol | N/A | N/A | N/A |
| **Tunicamycin** | 51.25 ±31.27 | 44.74 ±4.73 | 26.82 ±11.46 |
| **(2)** | N/A | N/A | 303.80 ±9.83 |
| **TUN-Ac,Ac** | N/A | N/A | 212.30 ±51.48 |
| **TUN-Boc,Boc** | N/A | N/A | 608.9 ±394.9 |
| **TUN** | N/A | N/A | 698.3 |
| **TUN-Cit,Cit** | N/A | N/A | 177.75 ±75.17 |
| **TUN-7,7** | N/A | N/A | 431.10 ±241.81 |
| **TUN-8,8** | N/A | N/A | 211.05 ±125.94 |
| **TUN-9,9** | N/A | N/A | 355.57 ±194.57 |
| **TUN-10,10** | N/A | N/A | 196.17 ±47.61 |
| **TUN-11,11** | N/A | N/A | 103.65 ±34.34 |
| **TUN-12,12** | N/A | N/A | 81.29 ±30.20 |

The LD<sub>50</sub> values are calculated from MTS cell proliferation assay based on dose response curves. The data shown are mean ± SEM errors of three independent experiments. N/A = not available. No or low cytotoxicity observed at highest concentration tested (400 µg mL<sup>-1</sup>), thus a reliable dose-response curve could not be generated from the experimental data.

**Table S5.** Minimal Lethal Dose, Minimal Inhibitory Concentration (μg/mL) and Relative Therapeutic Index (RTI)

|  | HepG2 | HEK 293 | *Mtb* H37RV <sup>a</sup> | RTI | *Mtb* H37RV <sup>b</sup> | RTI | *B. subtilis* EC1524 | RTI | *B. cereus* NRRL 11778 | RTI |
| --- | --- | --- | --- | --- | --- | --- | --- | --- | --- | --- |
| Tunicamycin | 100 | 100 | 0.6 | 167 | 0.08 | 1250 | 0.0015 | 66700 | 0.195 | 513 |
| TUN<sup>c</sup> | >400 | >400 | >60 |  | >60 |  | 100 | 8 | 100 | 8 |
| TUN-Cit,Cit | >400 | >400 | 16.3 | 49 | 3.4 | 235 | 6.25 | 128 | 100 | 8 |
| TUN-7,7 | >400 | >400 | 5.0 | 160 | 1.6 | 500 | 6.25 | 128 | 100 | 8 |
| TUN-8,8 | >400 | >400 | 0.7 | 1140 | 0.14 | 4714 | 0.78 | 1026 | 6.25 | 128 |
| TUN-9,9 | >400 | >400 | 0.2 | 4000 | 0.03 | 27000 | 0.024 | 33300 | 0.78 | 1026 |
| TUN-10,10 | >400 | >400 | 1.2 | 667 | 0.1 | 8000 | 0.39 | 2050 | 0.39 | 2050 |
| TUN-11,11 | >400 | >400 | 4.1 | 195 | 1.6 | 500 | 1.56 | 513 | 0.78 | 1026 |
| TUN-12,12 | >400 | >400 | 18.7 | 43 | 6.9 | 116 | 100 | 8 | 50 | 16 |

<sup>a</sup> 1-week in 7H9/ADC/Tw
<sup>b</sup> 1-week in GAST/Fe
<sup>c</sup> as *bis*-hydrochloride salt
Relative therapeutic index (RTI) is the ratio between the minimal lethal dose and the minimal inhibition concentration. The micro-broth dilution and the cytotoxicity tests were carried out by serial 2-fold dilutions. For no detectable cytotoxicity at 400 $\mu\text{g mL}^{-1}$ (>400), a minimal lethal dose of 800 $\mu\text{g mL}^{-1}$ was used to calculate the RTI.

## STAR*Methods

### CONTACT FOR REAGENT AND RESOURCE SHARING

Further information and requests for resources and reagents should be directed to and will be fulfilled by the Lead Contacts (Ben Davis, Liz Carpenter).

### Experimental Model and Subject Details

#### Sf9 cell culture

Sf9 cells were cultured in Sf 900 II SFM medium in a 27 °C incubator, rotating at 100 rpm.

#### HepG2 and HEK293 cell culture

HepG2 and HEK293 cells were cultured in DMEM medium supplemented with 10% heat inactivated fetal bovine serum (FBS, v/v). The cultures were maintained in a humidified incubator at 37 °C in 5% CO_2_/95% air. FBS was reduced to 2% for the cell proliferation assay. Raji cell culture.

#### Raji cell culture

Raji cells were cultured in RPMI-1640 medium supplemented with 10% heat inactivated fetal bovine serum (FBS, v/v). The cultures were maintained in a humidified incubator at 37 °C in 5% CO2/95% air. FBS was reduced to 2% for the cell proliferation assay.

### Method Details

#### Cloning and expression

The WT DPAGT1 cDNA sequence was cloned into the pFB-LIC-Bse expression vector (available from the SGC) with an N-terminal purification tag with a tobacco etch virus (TEV) protease cleavage site, and a 6x His purification sequence. Baculoviruses were produced by transformation of DH10Bac cells. Spodoptera frugiperda (Sf9) insect cells in Sf-900 II SFM medium (Life Technologies) were infected with recombinant baculovirus and incubated for 65 h at 27 °C in shaker flasks.

#### Site Directed Mutagenesis

Coding DNA for DPAGT1 point mutations were created via Megaprimer site-directed mutagenesis, as previously described (Xu *et al.* 2003). The mutated DNA strand was subsequently sub-cloned into an appropriately pre-treated pFB-LIC-Bse expression vector via ligase independent cloning (LIC) as described for the wild-type construct.

#### Purification of DPAGT1 protein for structural and functional studies

Cell pellets from 1 litre of insect cell culture were resuspended in 40ml in lysis buffer (50 mM HEPES, pH 7.5, 5 mM MgCl_2_, 200 mM NaCl, 5 mM imidazole, 2 mM TCEP (added fresh), 5% glycerol, Roche protease inhibitors (1 tablet per 40ml buffer, added on day of use) in warm water, mixing constantly to keep the sample cold. Cells were lysed by two passes through an EmulsiFlex-C3 homogenizer (Aventin). Protein was extracted from cell membranes by incubation of the crude cell lysate with 1 % (w/v) OGNG and 0.1 % (w/v) CHS for 1 h at 4 °C on a rotator. Cell debris and unlysed cells were removed by centrifugation at 35,000 g for 45 mins. Immobilized metal affinity chromatography was then used to purify the detergent-solubilized His-tagged protein by batch binding to Co^2+^ charged TALON resin (Clontech) at 4 °C for 1 h. The resin was then washed with wash buffer (WB: 50 mM HEPES (pH 7.5), 5mM MgCl2, 10 mM imidazole (pH 8.0), 200 mM NaCl, 2 mM TCEP (added fresh), 5 % Glycerol, 0.18 % OGNG, 0.01 8% CHS, 0.0036 % cardiolipin) and the protein was eluted with WB supplemented with 250 mM imidazole (pH 8.0). The eluted protein was desalted using PD-10 columns (GE healthcare) pre-equilibrated with gel filtration buffer (GFB: 20 mM HEPES (pH 7.5), 5 mM MgCl2, 200 mM NaCl, 2 mM TCEP, 0.12 % OGNG, 0.012 % CHS, 0.0024 % cardiolipin). Desalted protein was subsequently treated with 10:1 TEV protease (w:w, protein:enzyme) overnight at 4 °C. The TEV protease treated protein was separated from the 6-His tagged enzymes and uncleaved DPAGT1 by incubation for 1 h with Talon resin (prepared as described above) at 4 °C for 1h. The resin was collected in a column, the flowthrough collected and the protein sample was centrifuged at 21,500 rpm in a Beckman TA25.5 rotor for 10 min at 4 °C. The protein was then concentrated to 0.5 ml using a 30 kDa cutoff PES concentrator (Corning), with mixing every 5 mins during concentration. After centrifugation at 20,000 g for 10 min the protein was further purified by size exclusion chromatography (SEC) on a Sepharose S200 column (GE Healthcare) in GFB. The peak fractions were pooled and concentrated using a Sartorius 2ml PES 50 kDa concentrator (pre-equilibrated with GFB without detergent), at 3220 g. The protein was centrifuged at 20,000g for 15 mins, then flash frozen in liquid nitrogen. The final concentration was 20-30 mg/ml.

#### Radiolabelled substrate enzyme activity assay

2 μl of 2 μM DPAGT1 WT or mutant proteins in GFB buffer supplemented with 5mM extra MgCl_2_, 1% OGNG/CHS/cardiolipin and dolichyl monophosphate was combined with 2 μl of UDP-N-acetyl [1-14C] D-glucosamine in the same buffer and incubated at 37 °C on a heat block for 21 min. The reaction was terminated by the addition of 6 μl of 100% methanol and immediately transferred onto ice. 1 μl of sample was spotted onto a silica coated TLC plate in triplicate and run with a mobile phase consisting of chloroform, methanol, and water at a 65:25:4 ratio respectively. After the run, the TLC plate was dried thoroughly, wrapped in cling film, incubated with a phosphor imaging substrate for 4 days, then phosphor imaged using a Biorad. The pixel density of the spots corresponding to the hydrophobic product were divided by combined pixel density of the product and the substrate and multiplied by the known concentration of substrate added to ascertain the amount of product formed.

#### Thermostability assays for DPAGT1 WT and mutant proteins

Samples with a volume of 40 μl were prepared containing 0.5 mg/ml protein and 50 μM compound or 5% DMSO in GFB. A glass capillary was dipped into each sample, with the capillary held horizontally to ensure that the capillary was full of the sample. The capillaries were placed on the capillary holder on the Nanotemper Prometheus. Technical triplicates of each sample were prepared, and each experiment was conducted with biological triplicates of each protein. PR.ThermControl software was used to run the experiment and analyse the data. A melting curve from 20 °C – 95 °C at 5 °C/min was performed. The minima of the first derivative of the 330/350nm ratio was used to determine the Tm_1/2_.

#### Crystallisation of apo DPAGT1 protein

Protein was concentrated to ~20 mg/ml, then diluted to 9–12 mg/ml using GFB without detergent. Initial crystals were grown at 4 °C with WT protein purified with DOPG using sitting drop (150 nl) crystallisation set up in 96-well format using a Mosquito crystallization robot (TTP Labtech) with protein:reservoir ratios of 2:1, 1:1 and 1:2. Crystals of DPAGT1 were initially obtained in MemGold2 HT-96 screen (Molecular Dimensions) (Newstead et al., 2008) condition G10. Reproducibility for this condition was poor, and protein purification was further optimised. Adding cardiolipin instead of DOPG to purification buffers as well as using the Val264Gly mutant construct improved crystal reproducibility. A new crystallisation condition was obtained at 20 °C which was optimised to 32-38 % (v/v) polyethylene glycol (PEG) 300, 50 mM sodium chloride, 0-5 mM sodium tungstate, 0.1 M bicine, pH 9.0. Seeding was used to further improve crystallisation reproducibility.

#### Crystallisation of the Val264Gly mutant DPAGT1 with UDP-GlcNAc and Tunicamycin

To obtain a co-structure of DPAGT1 with UDP-GlcNAc, Val264Gly DPAGT1 was incubated with 10 mM UDP-GlcNAc for 1 hour at 4 °C before setting up crystallisation with seeds at 20 °C in reservoir solution containing 32.5 - 38 % (v/v) PEG 300, 50 mM NaCl, 0.1 M bicine, pH 9.0. To obtain a co-structure of DPAGT1 with tunicamycin, Val264Gly DPAGT1 harvested after SEC was supplemented with 0.1 mM tunicamycin and incubated for 1 hour at 4 °C before concentrating. The concentrated protein was crystallised at 4 °C in sitting drop crystallisation trials with reservoir solution containing 36 % (v/v) PEG 200, 50 mM sodium chloride, 0.1 M bicine, pH 8.5.

#### Data collection and structure determination for the DPAGT1 complexes

All data were collected at Diamond Light Source (beamlines I24, I04-1 and I04) to resolutions between 3.2-3.6Å based on CC_1/2_=0.5 criteria. Data were processed, reduced and scaled using XDS (Kabsch, 2010) and AIMLESS (Evans, 2006) (Table S1). All crystals belong to space group *P*6_5_22 and contain a single DPAGT1 monomer in the asymmetric unit with 70% solvent. Initial phase estimates were obtained using molecular replacement (MR). Briefly, an initial search model was built automatically using the PHYRE2 web server (Kelley et al., 2015) based on the coordinates of the bacterial homolog MraY (19% sequence identity; PDB: 4J72). However, this simple homology model failed to produce any meaningful MR solutions when used in isolation in PHASER (McCoy et al., 2007). The PHYRE model was then subjected to model pre-refinement using the procedures implemented in MR-ROSETTA in PHENIX (Terwilliger et al., 2012) and the resultant five best-scoring output models were trimmed at their termini and in the TMH9/TMH10 cytoplasmic loop region and superposed for use as an ensemble search model in PHASER. A marginal but consistent solution was obtained that exhibited sensible crystal packing in space group *P*6_5_22 but both the initial maps and model refinement were inconclusive. The model positioned using PHASER was converted to a poly-alanine trace and recycled into MR-ROSETTA, using model_already_placed=True option. The resultant MR-ROSETTA output model had an R/Rfree of 42/46 and the electron density maps showed new features not present in the input coordinates that indicated that the structure had been successfully phased.

Using the MR-ROSETTA solution as a starting point, the remaining regions of the DPAGT1 structure could be built manually using COOT (Emsley et al., 2010) using the WT 3.6Å native data. However, the novel 52 amino acid cytoplasmic insertion domain between TMH9 and TMH10 was poorly ordered and proved difficult to trace. This region was primarily traced using the electron density maps for the UDP-GlcNAc complex as substrate binding results in partial stabilisation of the TMH9/10 insertion domain. All electron density maps were sharpened in COOT to aid model building using a *B*-factor of −100Å^2^. Sequence assignment was aided by using both mercury labelling of cysteines (Figure S2A) and the sulphur anomalous signal from a dataset collected from UDP-GlcNAc complexes crystals at a wavelength of 1.7Å (Figure S2B). Anomalous difference maps, combined with anomalous substructure completion using PHASER-EP, clearly revealed the location of 18 of the expected 22 sulphur positions and helped to confirm the sequence register (Figure S2B). Additional experimental phasing information was provided by a Pr^3+^ derivative. The resultant model for the entire chain was then refined against both the unbound Val264Gly (3.2Å), Val264Gly UDP-GlcNAc (3.1Å), Val264Gly tunicamycin (3.4Å) as well as the WT unbound (3.6Å) data using BUSTER v2.10.2 / v2.10.3 and REFMAC (UDP-GlcNAc complex – final cycle only). All data were mildly anisotropic but were used in BUSTER without truncation apart from the unbound Val264Gly dataset which was anisotropically truncated with STARANISO using default cutoffs. Reference model restraints improved the refinement behavior for the 3.6Å unbound WT structure (using the unbound Val264Gly model as the reference model). Ligand restraints were generated using the GRADE webserver (http://grade.globalphasing.org) and a single TLS group encompassing the entire protein chain was used in refinement. The presence of a single magnesium ion in the UDP-GlcNAc complex was verified using the anomalous signal from a dataset collected at 1.82Å from a crystal grown in the presence of MnCl_2_. A single peak (19σ) was observed in an anomalous difference Fourier map calculated at 4Å adjacent to UDP-GlcNAc pyrophosphate; the next highest peak corresponded to various sulphur atoms (5σ). A second putative Mn^2+^/Mg^2+^ site was identified between E94 and H270 on the luminal face (peak height 4.3σ). Elongated lipid–like density on the TMH1/6/7/10 face of DPAGT1 was modelled as a dioleoylphosphoglycerol (DOPG) lipid. This lipid feature was present in both the electron density maps of the WT structure (purified with DOPG added) and the various Val264Gly mutant structures (purified with cardiolipin). The presence of DOPG in the purified protein samples used for crystallization was detected by mass spectrometry (Figure S1I). Lipid-like density was also present in the concave putative Dol-P groove adjacent to the EL4 luminal hairpin and has been modelled as unknown lipid/alkyl chains (UNL) in both the UDP-GlcNAc and TUN complexes. An additional persistent feature in all structures was electron density at the mouth of the active site adjacent to Trp122 (TMH4) and Leu293 (TMH9), presumably arising from a co-purified lipid that mimics the Dol-P substrate binding. However, the density was poorly resolved and no attempt was made to interpret this feature in the final models. The commercial preparation of TUN used for co-crystallisation in this study is a natural product and contains a range of different aliphatic chain lengths (n=8-11). A chain length of n=9 was chosen for the modelled TUN as this appeared to most consistent with the observed electron density (Figure S2E).

The representative final model comprises the entire polypeptide chain between residues Leu7 and Gln400 apart from the flexible EL2 loop connecting TMH2 and TMH3 and part of the poorly-ordered EL4 luminal hairpin (residue 152-161).

#### Native mass spectrometry

For native mass spectrometry analysis purified DPAGT1 protein was diluted to 10 μM protein concentration using 200 mM ammonium acetate supplemented with 0.16% OGNG solution followed by buffer exchanged into 200 mM ammonium acetate, 0.16% OGNG using a micro biospin column (Micro Bio-Spin 6, Bio-Rad). Native MS experiments were conducted using a Q Exactive instrument (Thermo Fisher, Germany) with modifications for high-mass transmission optimisation ((Mehmood et al., 2016)). Typically, 2 µl of buffer exchanged protein solution was electrosprayed from gold-plated borosilicate capillaries prepared in house. The instrument was operated under following parameters: 1.2 kV capillary voltage, 100 V S-lens, 250 ᴼC capillary temperature, 100 cone voltage. The activation voltage in HCD cell was raised from 100-200 V until a nicely resolve charge state pattern was found. Pressure in the HCD cell was raised to 1.2e-9 mbar for efficient transmission of protein. The instrument was operated in positive ion mode and was calibrated using caesium iodide solution.

For protein activity detection, 5 μM DPAGT1 protein was incubated separately with 50 μM dolichol-phosphate and UDP-GlcNAc or together in presence of both substrates at 37 °C for 21 min. Native MS buffer was used to spray the protein in presence of substrates on a modified Q Exactive Plus instrument. For detection of phosphate groups the instrument was operated in negative mode and minimal activation conditions were applied. The instrument was calibrated using cesium iodide. The experiments were repeated three using three different protein preparations.

#### Lipid analysis by tandem mass spectrometry

Lipidomics analysis was performed on DPAGT1 to identify co-purified lipid in a similar manner that has been described previously (Ref: DOI: 10.1074/jbc.M116.732107) after modifications in liquid chromatography gradient. Briefly protein was digested with trypsin (1:50 units) for overnight at 37 °C in a thermomixer (Eppendorf) under continuous shaking. The digest was dried in a SpeedVac until complete dryness and re-dissolved in 68% solution A (ACN:H2O 60:40, 10 mM ammonium formate and 0.1% formic acid) and 32% solution B (IPA:ACN 90:10, 10 mM ammonium formate and 0.1% formic acid). The tryptic digest mixture was loaded onto a pre-equilibrated C18 column (Acclaim PepMap 100, C18, 75 μm × 15 cm; Thermo Scientific) at a flow rate of 300 nl min^−1^. The lipids were separated under following gradient: In 10 min solvent B was ramped from 2% to 65% over 1 min, then 80% over 6 min, before being held at 80% for 10 min, then ramped to 99% over 6 min and held for 7 min. The nano-flow reversed-phase liquid chromatography (Dionex UltiMate 3000 RSLC nano System, Thermo Scientific) was directly coupled to an LTQ-Orbitrap XL hybrid mass spectrometer (Thermo Scientific) via a dynamic nanospray source. Typical MS conditions were spray voltage of 1.6 kV and capillary temperature of 275 °C.

The LTQ-Orbitrap XL was set up in negative ion mode and in data-dependent acquisition mode to perform five MS/MS scans per MS scan. Survey full-scan MS spectra were acquired in the Orbitrap (m/z 350–2,000) with a resolution of 60,000. The chromatogram was manually analysed for presence of different masses followed by lipids identification by manually comparing their experimental and theoretical fragmentation pattern.

#### Exon Trap Analysis

*DPAGT1* exons 2, 3 and 4 and flanking intronic sequences or exons 6, 7 and 8 and flanking sequences were cloned into the pET01 vector (MoBiTec). c.478G>A and c.791T>G were respectively introduced by site-directed mutagenesis using Quikchange kit from Stratagene and confirmed by Sanger sequencing. Control and mutant vector DNA were electroporated into the human rhabdomyosarcoma cell line TE671 using the NEON electroporator (Invitrogen). Total RNA was purified 48hr after transfection, reverse transcribed into cDNA using Retroscript kit (Ambion). cDNA was amplified using primers specific to the vector exons. The amplicons were run on agarose/TBE gels, visualized under UV/ethidium bromide and then gel purified and sequenced.

#### Mueller-Hinton Agar Plate

Muller-Hinton agar plate was prepared according to CLSI standards. Oxoid Mueller-Hinton Agar was prepared according to the manufacture’s protocol. 25 mL of the warm agar solution was transferred to 90mm × 16.2mm plate via sterile pipette. The agar plate was cooled at room temperature for 15 minutes before use or storage at 4 °C up to two weeks.

#### Kirby-Bauer Disc Diffusion Test

Oxoid Blank Disc was impregnated with the desired test substance. A 0.5 McFarland standard inoculum was prepared by adding 3-5 single colonies to 10 mL MH broth in 15 mL-falcon tube and standardised to 0.5-McFarland standard. The inoculum was used within 10 minutes. A sterile cotton swab was dipped in the inoculum, gently pressed against the side of the tube to remove excess liquid, and generously streaked on MH agar plate to fully cover the plate. The impregnated disc was carefully placed on the agar (important: once the disc touches the agar, it should not be moved). The plate was incubated at 35 °C for 20 hrs overnight. A digital calliper was used to measure the zone diameter. The recorded zone diameter is an average of three zone diameters measured of one zone.

#### Micro-dilution culture to determine minimal inhibition and minimal bactericidal concentrations

In a sterile 96-well plate, serial dilutions were made with the test substance to final volume of 50 μL. Inoculum was then prepared in Mueller-Hinton broth to 0.5 McFarland and diluted before adding 50 μL to the well to make ~1×10^5^ CFU/mL. The culture plate was incubated at 35 °C for 20-24 hrs. A positive growth control and sterility wells were also prepared along with the culture wells. Absorbance at OD_600_ was taken using BMG Labtech SPECTROstar Omega spectrophotometer. MIC is determined by the lowest concentration without growth. IC50 is determined from plotting a dose-response curve, see Data Analysis below. To determine the MBC, using a multi-channel pipette, 1 μL of culture broth was taken from the same 96-well microdilution growth plate and carefully inoculated on surface of MH agar plate. The plate is then incubated for additional 20 hrs. at 35 °C. The MBC value is the lowest concentration without observed growth on the agar.

#### Cell proliferation assay

In sterile 96-well plate, each well was seeded with ~1×10^5^ cells. The cells are then grown confluent overnight in 100 μL DMEM with 10% FBS. The medium is replenished with DMEM with 2% FBS and added vehicle control or test substance the next day. For test substance in methanol, stock was added to 25 μL of DMEM with 2% FBS or PBS and placed in the laminar flow hood to let the methanol evaporate (about 1-3 hrs). Once the methanol has evaporated, DMEM with 2% FBS was added to final volume of 100 μL of desired test concentration. A blank methanol control was also made to ensure that any cytotoxicity did not result from methanol contamination. Cells are grown for additional 24 hrs. The cell viability was determined by using Promega CellTiter 96© AQ_ueous_ One Solution Cell Proliferation Assay (3-(4,5-dimethylthiazol-2-yl)-5-(3-carboxymethoxyphenyl)-2-(4-sulfophenyl)-2H-tetrazolium, inner salt; MTS) System following the manufacture protocol.

#### 24 Hr Cell Cycle Test

The cell cycle test is carried out in 96-well tissue culture plate. A T-75 tissue culture flask is first used to culture a stock of HEK293 cells. Once the cells are about 90% confluent, the old medium is decanted and the cells are carefully washed with PBS and resuspended in DMEM with 10% FBS. HEK293 cells are seeded into the 96-well plates (1×10^5^ cells/well). Before placing the culture in the incubator (37 °C with 5% CO_2_), the plate is set aside for 10 minutes for the cells to settle to the bottom of the plate. Once the cells are confluent the next day or when they are ready to be treated, DMEM with 2% FBS containing the test substance is prepared beforehand during the day of the cell treatment.

Once the cell culture is ready for harvesting, the cells are then fixed with cold 70% ethanol after two washes with PBS (w/ 0.1% BSA). The cells were fixed overnight at 4 °C. Propidium Iodide (PI)/RNase Staining Solution from NEB was used for DNA staining.

For test substance in methanol, the methanol from stock solution was added to 50 uL of DMEM or PBS in a round-well plate (8-well plate) and placed in the laminar hood to have the methanol evaporate off in about 1-2 hrs and then adding the needed DMEM with 2% FBS to the desired volume.

#### Time-course Cell Cycle Test

The time-course cell cycle test is carried out in 96-well tissue culture plate. Three sets of plates were prepared as described above with samples prepared in triplicate. A 0hr control cell culture was first collected before setting up the plates for treatment. Each set is used for 24hr, 48hr or 72hr time-point data collection. Due to nature of the experiment, the DMEM used contained 10% Heat-inactivated FBS for the cells to have enough nutrients to the 72hr cell cycle. Once the cell culture is ready for harvesting, the cells are then fixed with cold 70% ethanol after two washes with PBS (w/ 0.1% BSA). The cells were fixed overnight at 4 C. Propidium Iodide (PI)/RNase Staining Solution from NEB was used for DNA staining.

#### IgG1Fc N-Glycosylation Assay

The test can be carried out in 6-or 8-well tissue culture plate. The transfected HEK293T/pHLsec:IgG1Fc cell culture harboring pHLsec:IgG1Fc plasmid with a His_6_-Tag IgG1Fc (Transfection protocol below) is grown for 2-3 days (37 °C with 5% CO_2_). Then the growth medium is collected and analyzed by SDS-PAGE and Western Blot analysis.

#### Transfection Protocol

Transfection reagents and medium are prepared fresh every time. A PEI (Polyethylimine) stock solution is first prepared (25kDa, linear −100ug/mL). PEI is added in MilliQ water and HCl (aq) is then added drop-wise with just enough to help solubilize the PEI. The PEI solution is filtered through 0.2 um membrane in laminar sterile hood. PEI solution is set aside. The DNA stock solution is added to serum-free DMEM medium at room temperature to make final DNA-DMEM concentration of 4 ug/mL. PEI solution is the added to DMEM-DNA solution to make final concentration of PEI of 4.6 ug/mL. The PEI-DNA transfection medium is gently stirred at room temperature for 20 minutes before use. For the transfection step, the DNA-PEI transfection is added first and placed in the incubator (37 °C with 5% CO_2_) for 10 minutes and then DMEM with 2% FBS is added. The DNA-PEI medium and DMEM with 2% FBS medium are added in 1:3 ratio.

#### Statistical Analysis

Data were analysed using Graph Pad PRISM 5.01 software. Dose-response curves were plotted from three independent data sets with SEM error bars.

#### Minimum inhibitory concentrations testing against *Mtb*

^*Mtb*^ H37Rv ATCC27294 was grown to OD_650nm_ of 0.2 in either Glycerol-Alanine salts medium (GAST/Fe) or 7H9/ADC/Tw. GAST/Fe consisted of per liter: 0.3 g Bacto Casitone (Difco), 4.0 g dibasic potassium phosphate, 2.0 g citric acid, 1.0 g L-alanine, 1.2 g magnesium chloride hexahydrate, 0.6 g potassium sulfate, 0.05 g ferric ammonium citrate, 2.0 g ammonium chloride, 1.80 ml of 10N NaOH, and 10.0 ml of glycerol, 0.05% Tween 80 with pH adjusted to 6.6 before sterile filtration. 7H9/ADC/Tw consisted of Middlebrook 7H9 broth base (Becton Dickinson) supplemented with 0.2% glycerol/0.5% BSA fraction V/ 0.2% glucose/ 0.08% NaCl/ 0.05% Tween 80. Cells were diluted 1000-fold in this medium and an equal volume (50µL/well) added to 96-well U-bottom plates (Nunclon, Thermo Scientific) containing compound diluted in the respective growth medium (50µL/well). Dilutions ranged from 0.024 - 50 µg/mL, performed in technical duplicates over two biological repeats. The positive control was isoniazid (tested from 0.024 - 50 µM) and negative control was the solvent. Isoniazid MIC was 0.2 ± 0.1 µM. MIC was determined over 2-3 biological replicates, each time as a technical duplicate.

#### Microsomal stability assay protocol

NADPH-regenerating system was freshly prepared by combining (a) 37.2 mg glucose-6-phosphate into 1 mL of 100 mM potassium phosphate buffer, (b) 39.8 mg NADP into 1 mL of 100 mM potassium phosphate buffer, (c) 11.5 U glucose-6-phosphate dehydrogenase into 1 mL of 100 mM potassium phosphate buffer and (d) 26.8 mg MgCl2.6H2O into 1 mL of water. 271 µl phosphate buffer (100 mM, pH 7.4) and 18 µL of NADPH-regenerating system were added to each tube, which were kept on ice. Pooled mouse/human liver microsomes were removed from the -80°C freezer and thawed. 8 µl of liver microsomes (protein content >20 mg/ml) were added to each tube. The tubes were removed from ice and placed in a 37°C heating block. 3 µl of test compound (0.5 mM stock) was added (final concentration 5 µM). Control incubation was included for each compound where phosphate buffer was added instead of NADPH-regenerating system (minus NADPH). Verapamil was used as control in the assay. The reaction was quenched at selected time-points by the addition of 150 µl methanol containing 5 µM internal standard. The samples were removed from the heating block and centrifuged at 14,000 g at 4°C for 10 min. The supernatants were analyzed on LC/MS under single ionization mode (SIM) and scan mode.

Testing of tunicamycin analogue efficacy against *Mtb* growing in infected macrophages J774A.1 mouse macrophage cells were grown in J774 growth medium consisting of DMEM GlutaMAX (Gibco) supplemented with 10% fetal bovine serum, 20 mM HEPES + 0.5 mM sodium pyruvate, seeded in sterile tissue culture treated 24-well plates (Corning) at 2.5×10^5^ cells/well and allowed to attach for 24 h in J774 growth medium. *Mtb* H37Rv ATCC was grown to OD_650nm_ of 0.2 in 7H9/ADC/Tw, harvested, resuspended in J774 growth medium, filtered through a 5 μm filter to ensure a single cell suspension and diluted to 2.5×10^7^ cells/mL. Cells were infected with 0.1 mL *Mtb* cell suspension at a multiplicity of infection (MOI) of 10:1 and allowed for 24 h. After infection, the medium was aspirated and the monolayer of cells washed twice with Dulbecco’s PBS and subsequently fed with 1 mL of J774 growth medium containing the compounds at the indicated concentrations in triplicate wells for each drug concentration and time point. Rifampicin was used as positive control and methanol and DMSO used for the negative controls. After 3 and 7 days of treatment, cells were lysed by 0.1% SDS, appropriate dilutions made in 7H9/ADC/Tw and plated in duplicate on solid medium consisting of Middlebrook 7H11 (Becton Dickinson) supplemented with OADC [final concentration of 0.5% bovine serum albumin fraction V, 0.08% NaCl, 0.2% glucose, 0.2% glycerol, 0.06% oleic acid]. Colony counts were enumerated after 4 weeks of incubation.

#### Animal care and human ethics assurance

Mouse studies were carried out in accordance with the Guide for the Care and Use of Laboratory Animals of the National Institutes of Health under Animal study protocol numbers LCID 4E.

#### Mouse pharmacokinetics

Test compound was dosed to sixteen female C57BL/6 mice by intraperitoneal injection in a volume of 0.6 mL. To reconstitute **TUN-8,8** for intraperitoneal injections, compound (1mg) was dissolved in 0.1mL ethanol and subsequently diluted with 0.9 mL 1% Tween 80 in Dulbecco’s PBS. The mixture was warmed to 37◦C, vortexed and sonicated followed by sterile filtration through a 0.2 μm filter. Blood samples were taken from the tail vein of the mice at pre-determined time intervals post-dose, and serum collected after 4 h on ice. 30 µL serum samples were prepared with 30µL internal standard and 20µL of water or spiked standard along with 240µL of ACN/MeOH mix (3:1) to precipitate proteins. Samples were centrifuged 13krpm for 5 minutes and 2µL injected into LC-MS/MS system. Calibration standard was prepared 1mg of TM8 to 1mL DMSO and successively diluted by a factor of 3. TM9 IS solution was prepared to 200µg/mL in DMSO.

LC-MS/MS was performed on an Agilent 1290 Infinity HPLC coupled to an Agilent 6460C triple quadrupole mass selective detector with electrospray ionization in positive mode. TM8 and TM9 (as an internal standard) were detected using their M+H precursors ions with 514.1 and 528.2 Da/z product ions produced in a collision cell using collision energy 12V. Capillary voltage was 3000V and ESI used jet stream technology with sheath voltage 2000V. The column was an Agilent C18 Poroshell 120 2.7µm with dimensions 2.1 × 50mm at 40°C. Mobile phase was water (A) and acetonitrile (B) each with 0.1%(v/v) formic acid. With flow rate 0.8 ml/min, a gradient of 25% B progressed to 95% B over 4minutes, washed and re-equilibrated. Water for LC-MS/MS was purified by a Barnstead Diamond system to 18.2 MΩ-cm resistivity. Acetonitrile (ACN) and Methanol were HPLC grade from Fisher (Fairlawn NJ, USA) and formic acid was supplied from EMD Millipore Corp (Darmstadt, Germany). DMSO was ACS grade manufactured by Amresco LLC in Solon Ohio USA.

### Mouse infection

Female C57Bl/6 mice were aerosol-infected with 100-200 colony forming units of *Mtb* H37Rv with implantation dose determined after 24 hours by plating of lung homogenates on Middlebrook 7H11/OADC agar. Nine days after infection, five mice were euthanized and lung and spleens harvested for bacterial burden enumeration by plating of organ homogenates on Middlebrook 7H11/OADC agar. The remaining mice were treated in groups of 10 as follows: (1) daily intraperitoneal injections of 0.2 mL 10% ethanol/0.9% Tween 80/Dulbecco’s PBS; (2) daily intraperitoneal injections of 10 mg/kg **TUN-8,8** in 0.2 mL 10% ethanol/0.9% Tween 80/Dulbecco’s PBS; (3) oral gavage of 10 mg/kg Rifampicin in in 0.1mL water. Mice were treated daily for 2 weeks after which mice were euthanized and organ homogenates in 7H9/ADC/Tw plated on Middlebrook 7H11/OADC agar for colony enumeration.

## QUANTIFICATION AND STATISTICAL ANALYSIS

All values are the mean ± standard deviation. GraphPad Prism v7.02 was used to plot Michaelis-Menten curves using the least squares fitting method, and calculate the v_max_ and K_m_ values.

## DATA AND SOFTWARE AVAILABILITY

Preparation of figures

Figures were prepared using either PyMOL (Schrodinger, 2010) or UCSF-Chimera (Pettersen, 2004). Electrostatic surface potentials (Figure 2B) were calculated using the APBS plugin within PyMOL and the PDB2PQR server (Dolinsky et al., 2004). Hydrogens and missing sidechain atoms were added automatically to the refined X-ray structure using ICM-Pro (Molsoft LLC) prior to electrostatic surface calculations. All electrostatic surface potentials were visualized in UCSF-Chimera and colored between −10 (red) and +10 (blue) kT/e^−^.

## ADDITIONAL RESOURCES

DPAGT1 target enabling package: http://www.thesgc.org/tep/DPAGT1

## KEY RESOURCES TABLE

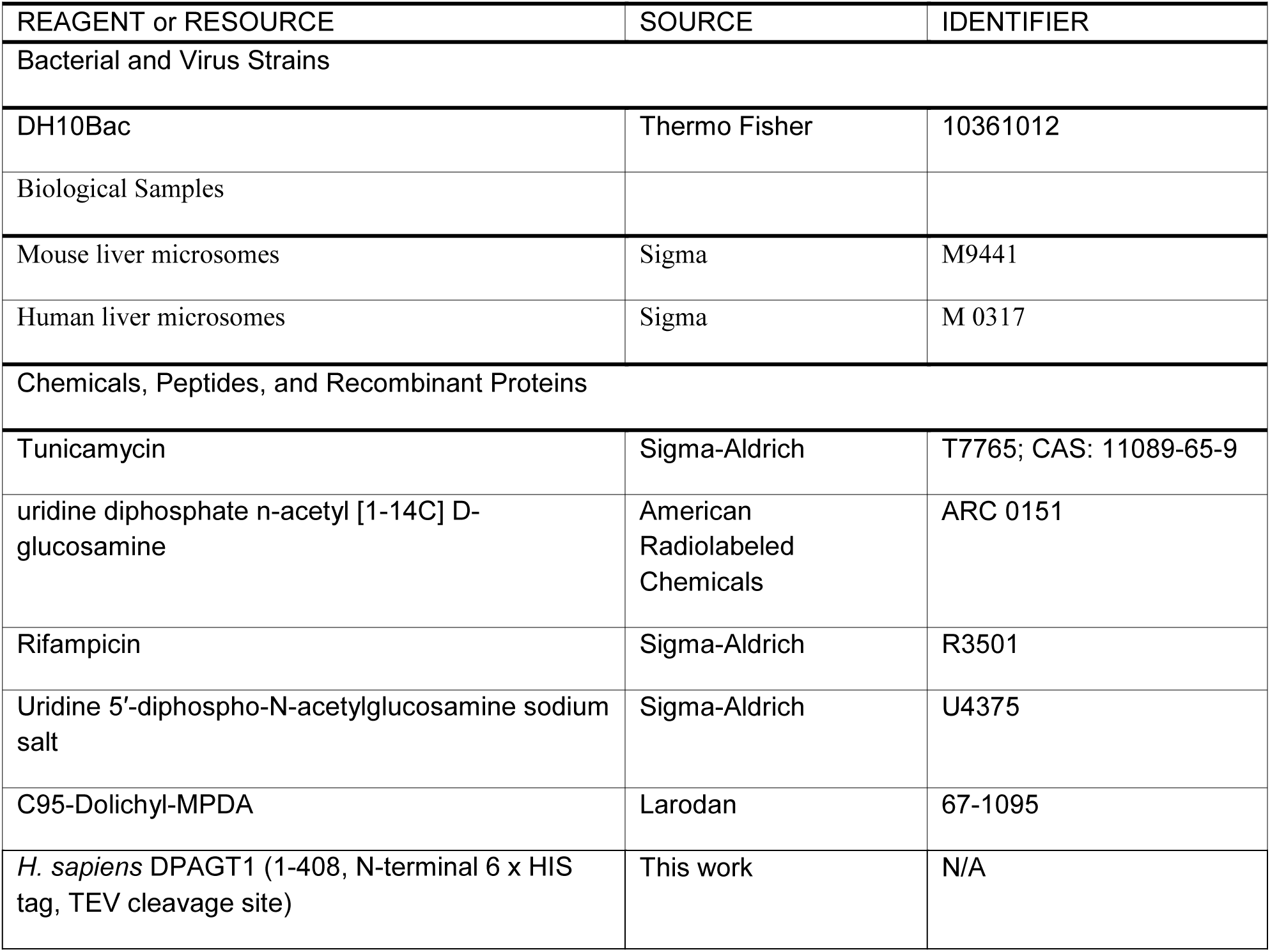
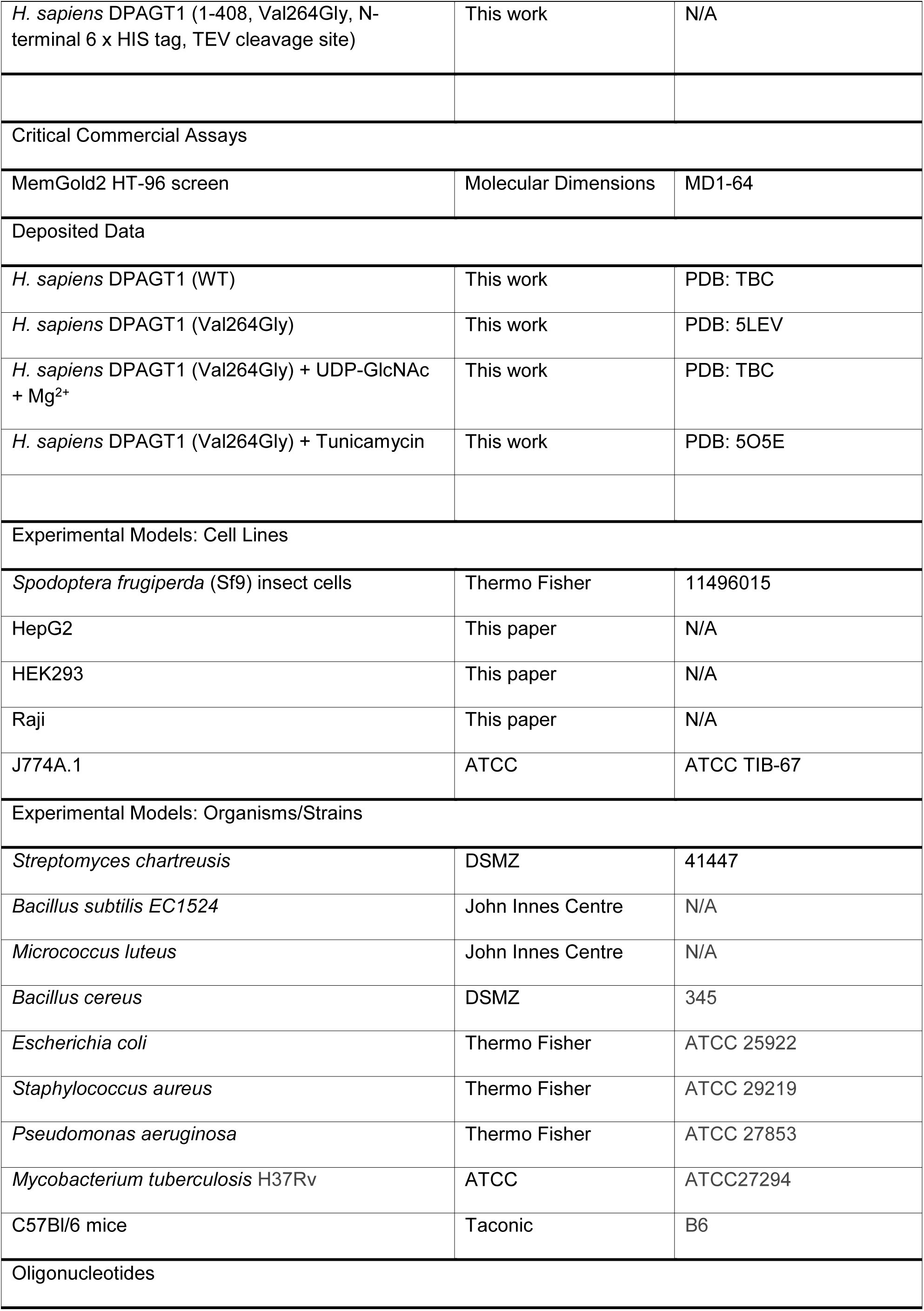
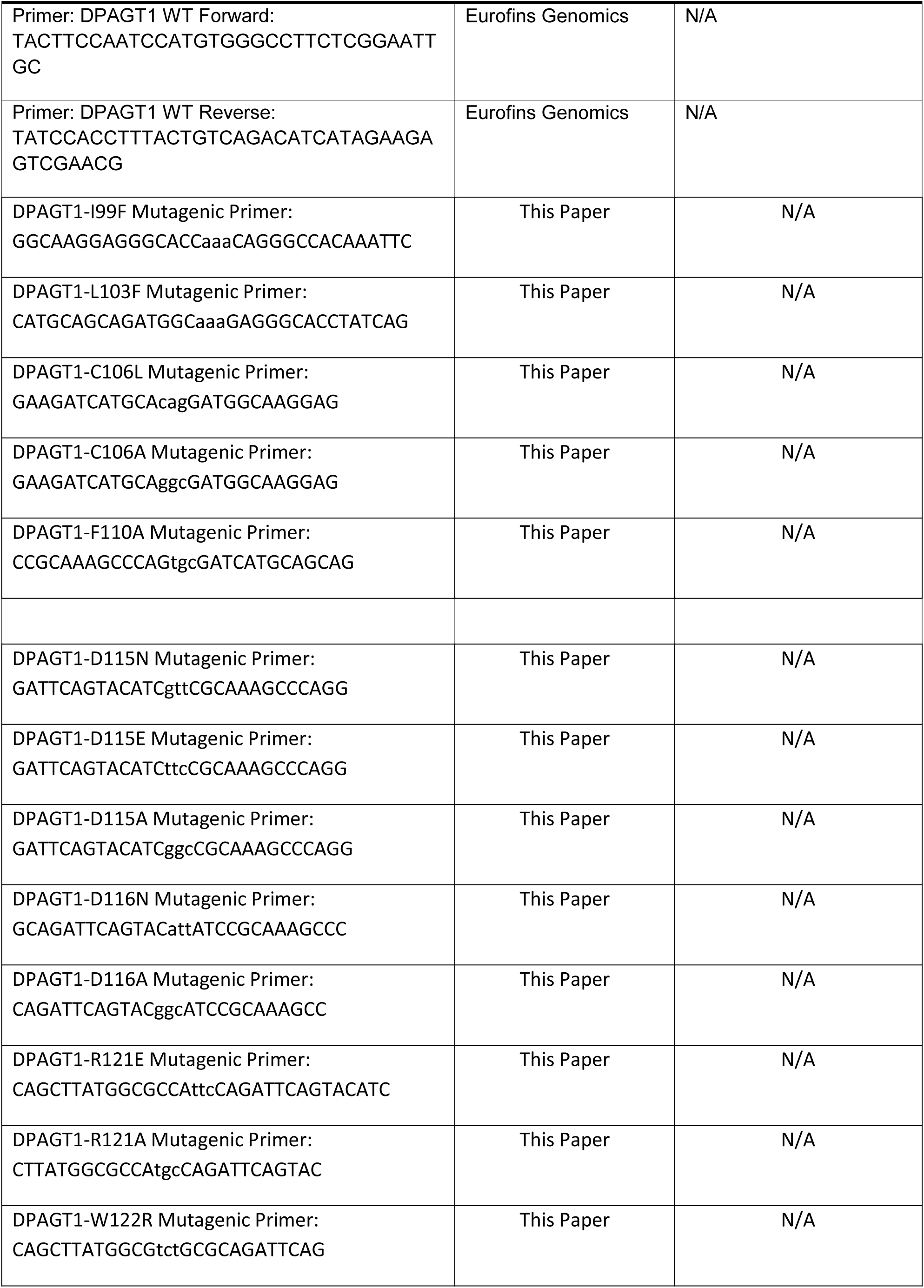
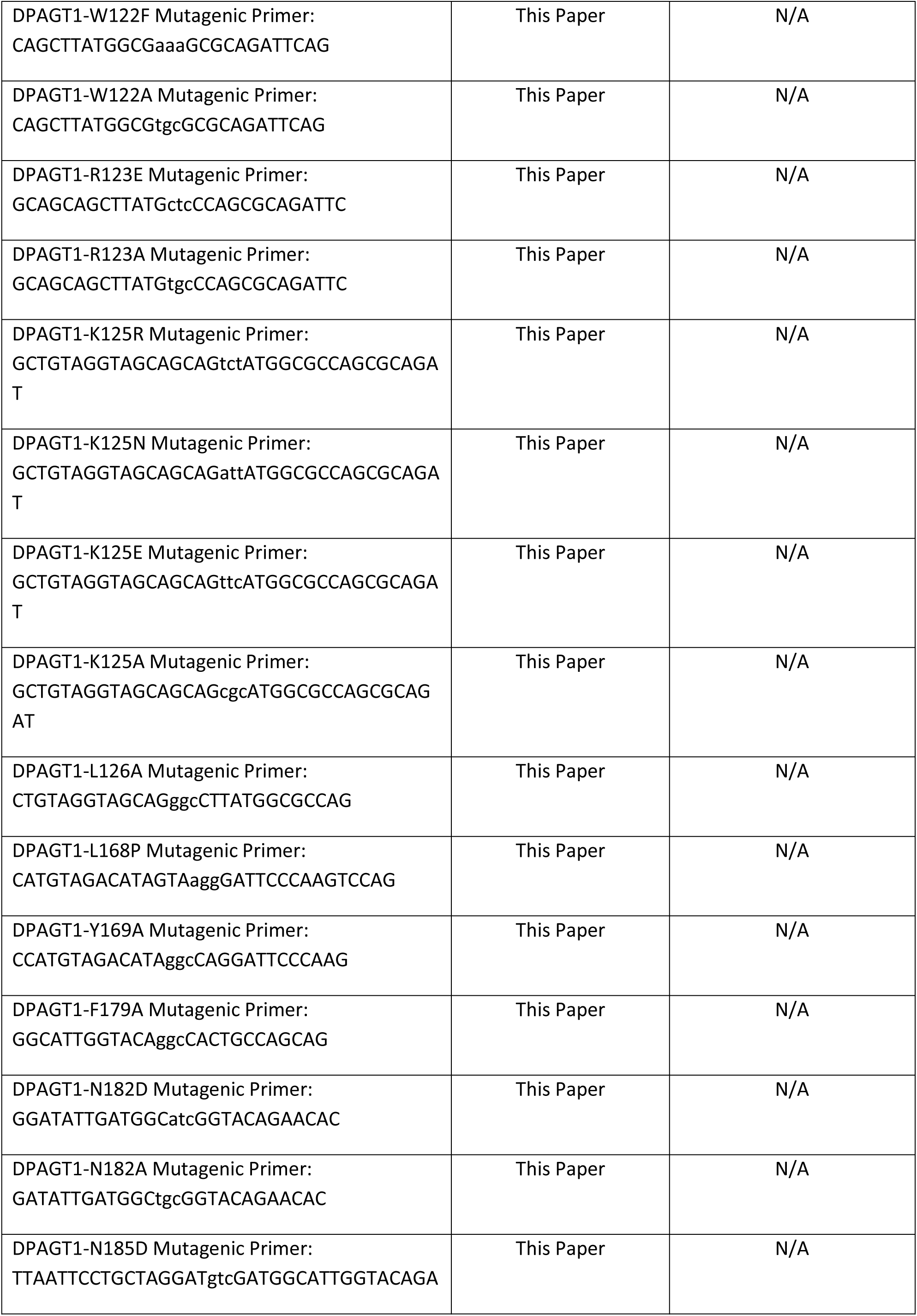
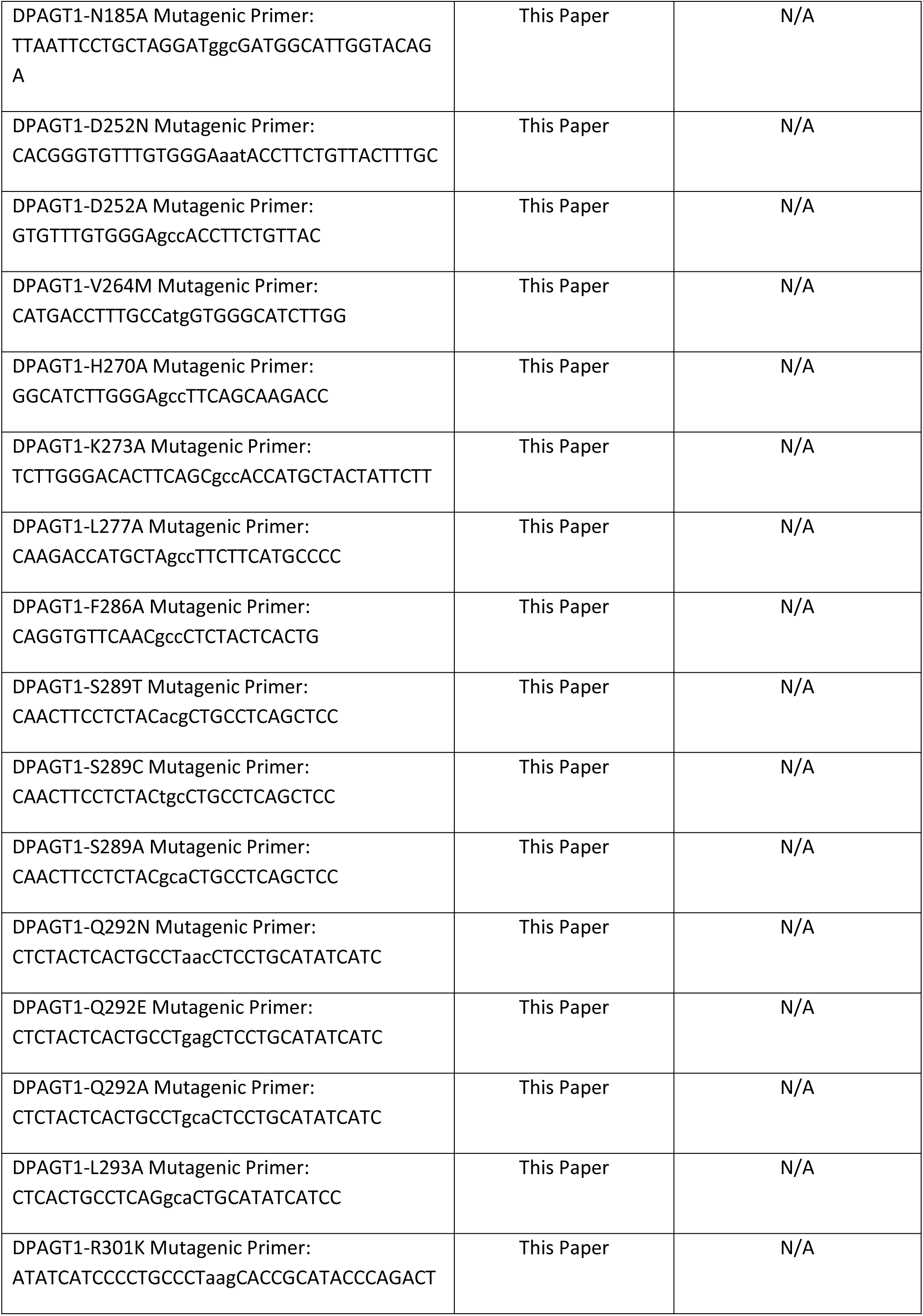
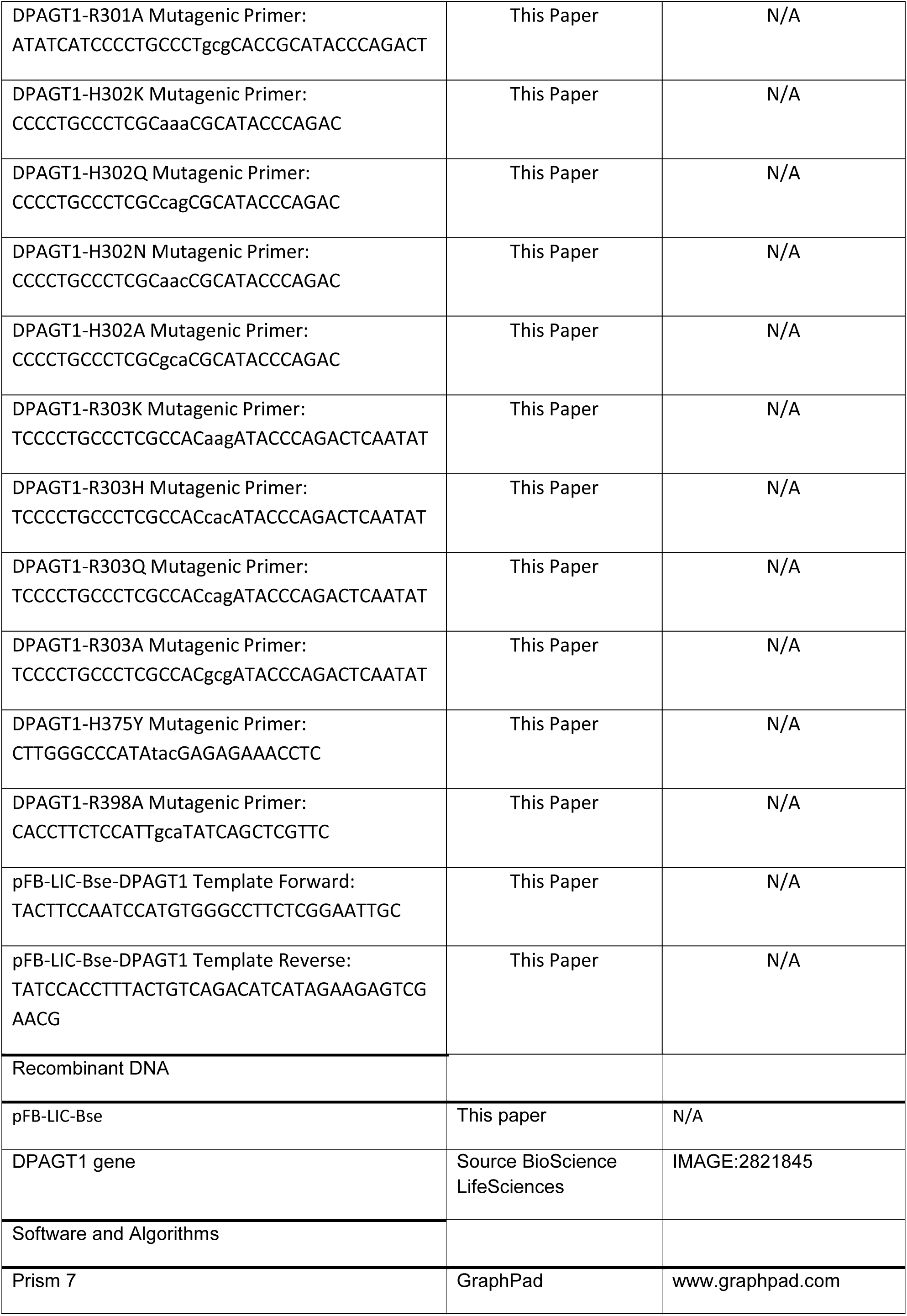
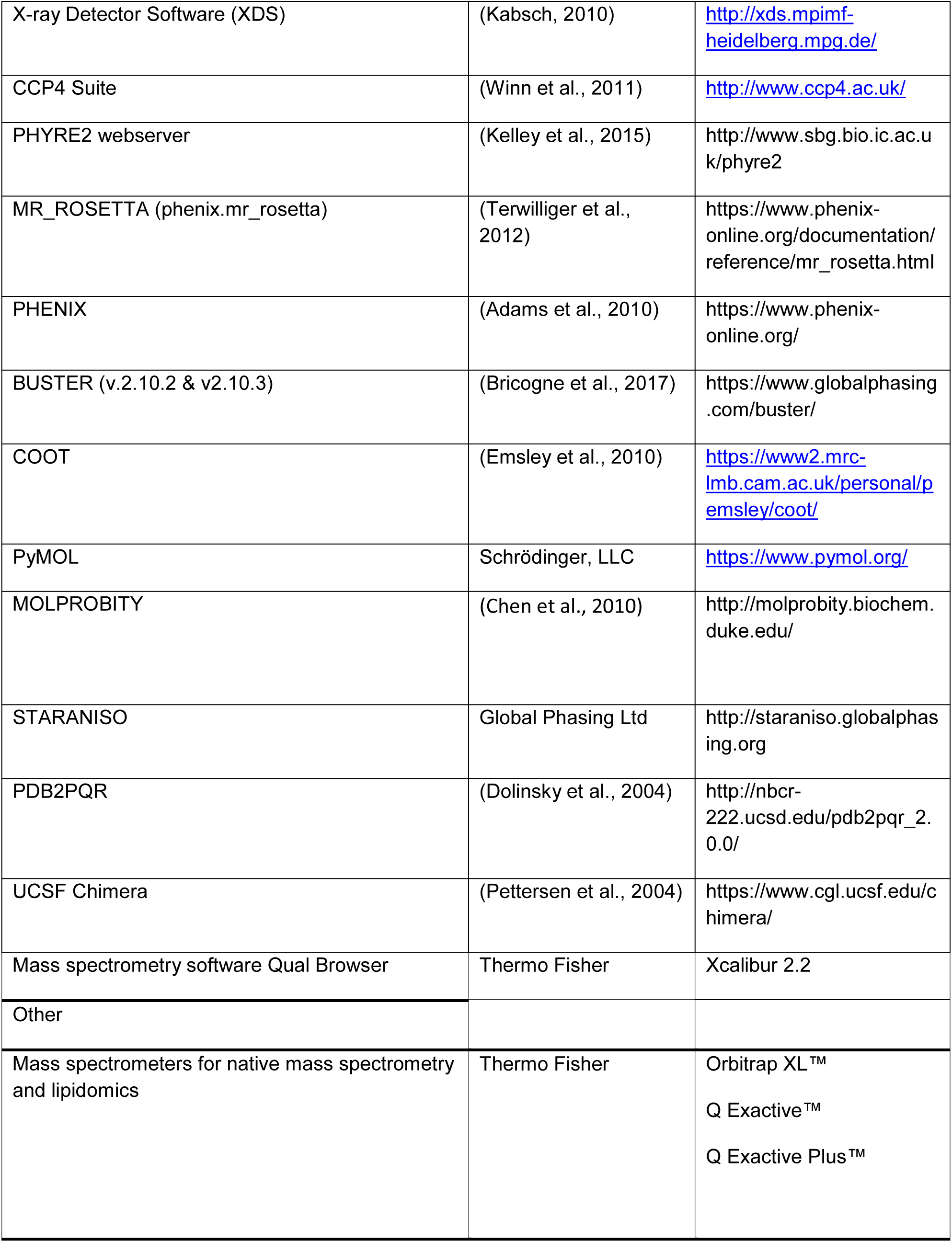

## SUPPLEMENTAL INFORMATION

### Supplemental information SI 1. Excel spreadsheet with mutations in DPAGT1 found in patients with DPAGT1-CMS and CDG-Ij. Related to Figures 4 and S3

This spreadsheet includes information on published DPAGT1 disease variants, together with relative activity and thermostability data for the WT and mutant protein from purified DPAGT1.

### Supplemental information SI 2

PDF file containing methods for semi-synthetic synthesis of the **TUN-X,X** analogues and related compounds. Related to Figure 5. See below.

## SUPPLEMENTARY INFORMATION SEMI SYNTHETIC SYNTHESIS

for Structures of DPAGT1 explain glycosylation disease mechanisms and advance TB antibiotic design

### Chemical Methods

#### General considerations

Proton nuclear magnetic resonance (δ_H_) spectra were recorded on a Bruker DPX 200 (200 MHz), Bruker DPX 400 (400 MHz), Bruker DQX 400 (400 MHz), or Bruker AVC 500 (500 MHz) or Bruker AV 700 (700 MHz) spectrometer. Carbon nuclear magnetic resonance spectra were recorded on a Bruker DQX 400 (100 MHz) or Bruker AVC 500 (125 MHz) with a ^13^C cryobprobe (125 MHz) AV 600 (151 MHz) with a ^13^C cryobprobe (151 MHz) or AV 700 (176 MHz) with a ^13^C cryobprobe (176 MHz). Spectra were assigned using a combination of ^1^H, ^13^C, HSQC, HMBC, COSY, and TOCSY. All chemical shifts were quoted on δ-scale in ppm, with residual solvent as internal standard. Coupling constants (*J*) are reported in hertz (Hz). Infrared spectra were recorded on a Bruker Tensor 27 Fourier Transform spectrophotometer recorded in wavenumbers (cm^−1^). Low-resolution mass spectra were recorded on a LCT Premier XE using electrospray ionization (ES). High-resolution mass spectra were recorded on a Bruker microTOF. Specific rotations were measured on Perkin Elmer 241 polarimeter with pathlength of 1.0 dm and concentration (*c*) in g/100 mL. Thin layer chromatography (TLC) was performed on Merck EMD Kieselgel 60F_254_ precoated aluminum backed plates. Reverse-phase thin layer chromatography (RF-TLC) was performed on Merck EMD Silica Gel RP-18 W F254s precoated glass backed plates. TLC and RF-TLC were visualized in combination of: 254/365 nm UV lamp; sulfuric acid (2 M in EtOH/Water 1:1); ninhydrin (2% ninhydrin in EtOH); aqueous KMnO_4_ (5% KMnO_4_ in 1 M NaOH); aqueous phosphomolybdinc acid/Ce(IV) (2.5% phosphomolybdic acid hydrate, 1% cerium(IV) sulfate hydrate, and 6% H_2_SO_4_); or ammonium molybdate3 (5% in 2M H_2_SO_4_). Flash chromatography was carried out with Fluka Kiegselgel 60 220-440 mesh silica gel. All solvents (analytical or HPLC) used were purchased from Sigma Aldrich, Fisher Scientific, or Rathburn. Anhydrous solvents were purchased from Sigma Aldrich and stored over molecular sieves (<0.005 % H_2_O). Petrol refers to the fraction of petroleum ether boiling point in the range of 40 – 60 °C. Analytical (Synergi™ 4 µm Hydro-RP 80A 100 × 4.60 mm) and preparative (Synergi™ 4 µm Hydro-RP 80A 100 × 21.20 mm) reversed phase C18 column for HPLC were obtained from Phenomenex. Brine refers to saturated solution of NaCl.

#### Analytical and Preparative HPLC Method for tunicamycins-like compound

Analytical-scale HPLC analysis and preparative-scale HPLC purification were performed on an UltiMate 3000, and the resulting data was analysed using Chromeleon software.

*Analytical Scale Analysis*. Column: Phenomenex, Synergi 4u Hydro-RP 80Å 100 × 4.60 mm 4micron; Flow rate: 1mL/min; Solvent A: 5% ACN and 0.1% FA in H_2_O; Solvent B: 0.1% FA in ACN; UV 260 nm.

Eluent gradient

Min. %B 1.000 0.0[%]

25.000 100.0 [%]

27.010 100.0 [%]

29.010 0.0 [%]

35.010 0.0 [%]

*Preparative Scale Purification*. Column: Phenomenex, Synergi 4u Hydro-RP 80Å 100 × 21.20 mm 4micron; Flow rate: 12mL/min; Solvent A: 5% ACN and 0.1% FA in H_2_O; Solvent B: 0.1% FA in ACN; UV 260 nm.

Eluent gradient

Min. %B

1.000 0.0[%]

25.000 100.0 [%]

27.010 100.0 [%]

28.010 0.0 [%]

35.010 0.0 [%]

#### Molecular weight for the extracted tunicamycin homologues used in this work

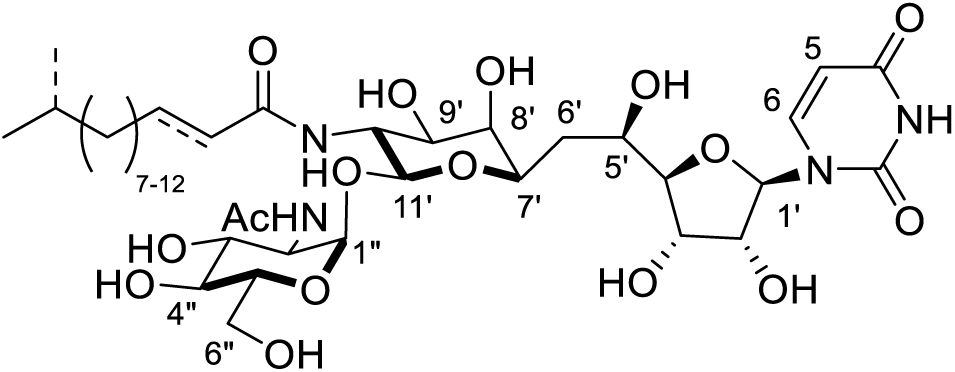

An average molecular weight has been used in molar calculations involving the extracted tunicamycins. Naturally produced tunicamycin is a mixture of homologues (see above). An average molecular weight of 838 g mol^−1^ is used in calculations based on the common homologue carbon chain lengths of *n* = 8, 9, 10, 11 unless otherwise specified.

#### Extraction of tunicamycins

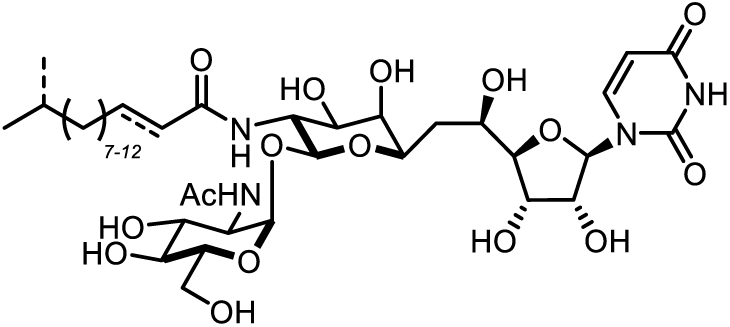

Crude tunicamycin was isolated from a *S. chartreusis* NRRL3882 fermentation culture by methanol extraction (Hamill, 1980; Takatsuki et al., 1971). *S. chartreusis* spore stock (2 µL) was added to 50 mL of TYD media in a 250 mL spring coiled flask, and incubated at 28°C and 200 rpm in a New Brunswick Series 25 shaker. After 36 h, aliquots of this culture (12 × 2 mL) was added to 12 × 1 L of TYD media including 6 g of glucose and 0.3 g of MgCl_2_ in unbaffled 2 L conical flasks, which were subsequently incubated at 28°C and 200 rpm in a New Brunswick Series 25 shaker. After 7 days, cells and supernatant were separated via decantation and centrifugation at 8500 rpm (Beckman Coulter Avanti J-25). Tunicamycin was extracted from both the centrifuged cells and supernatant. Tunicamycin in the supernatant was isolated by hydrophobic interaction chromatography. Amberlite XAD-16 was first preconditioned by washing with MeOH (x 3) and then distilled water (x 2). This preconditioned resin (15 g/L) was then added to the resulting supernatant and stirred for 2 h. The magnetic stirrer was then turned off and the XAD-16 resin was allowed to settle to the bottom of the flask, after which the majority of the supernatant was decanted and the remaining supernatant was removed by filtration. The collected resin was washed with water (200 mL) for 15 min and filtered through filter paper, and then stirred sequentially in MeOH (600 mL, 15 min), iPrOH (600 mL, 15 min) and MeOH (600 mL, overnight). The organic fractions were combined and concentrated *in vacuo*. The concentrated tunicamycin solution was aliquoted into four Falcon tubes and the volume adjusted to 40 mL with 1M HCl to precipitate tunicamycin. The insoluble precipitate was collected via centrifugation, re-dissolved in MeOH and then diluted with 400 mL of acetone. The acetone solution was kept at −20 °C overnight and the precipitated crude tunicamycin collected by filtration. Tunicamycin was also extracted from the cell pellet. The pellet was stirred in 1M aq. HCl (800 mL) for 30 min, after which the cells were collected by centrifugation at 9000 rpm (Beckman Coulter Avanti J-25). This process was repeated, after which the cell pellet was stirred in MeOH (400 mL) overnight. The cells were collected by filtration, resuspended in MeOH (400 mL) and stirred for a further 4 h. The MeOH fractions were combined, concentrated *in vacuo*, and tunicamycin precipitated with acetone (400 mL). Crude tunicamycin: TLC: R_*f*_ 0.3 in water/isopropanol/ethyl acetate (W/_*i*_POH/EtOAc, 1:3:6); ^1^H NMR (400 MHz, CD_3_OD) δ ppm 0.89, 0.91 (2 × s, 2 × 3 H, -CH(*CH*_*3*_)_2_), 1.14 – 1.66 (m, n × CH_2_^fatty acid^), 1.95 (s, 3 H, -CH_3_^NHAc^), 3.36 – 4.05 (m, -CH_2_^sugar^, CH^sugar^), 4.10 (t, *J* = 9.30 Hz, 1 H, H-10’), 4.20 (t, *J*_2’,1’_ = 5.80 Hz, 1 H, H-2’), 4.59 (d, *J*_11’,10’_ = 8.9 Hz, 1 H, H-11’), 4.94 (d, *J* = 3.6 Hz, 1 H, H-1’’), 5.77 (d, *J*_5,6_ = 8.2 Hz, 1 H, H-5^uracil^), 5.95 (d, *J*_1’,2’_ = 5.5 Hz, 1 H, H-1’), 5.96 (d, *J*_HC=CH trans_ = 15.4 Hz, 1 H, = C*H*C(O)-), 6.84 (dt, *J*_HC=CH trans_ = 14.5 Hz, *J* = 7.85 Hz, 1 H, -CH_2_*H*C=), 7.94 (d, *J*_6,5_ = 8.2 Hz, 1 H, H-6^uracil^); LRMS *m/z* (ESI^+^): [(M + Na)^+^] = 839 (18%), 853 (100%), 867 (92%), 881 (30%); (ESI^−^): [(M + Cl)^−^] = 851 (20%), 865 (100%), 879 (94%), 893 (34%). Flanking peaks with mass ± 14 corresponded to 8 × CH_2_, 9 × CH_2_, 10 × CH_2_, and 11 × CH_2_. IR ν: 3325, 2925, 2360, 2342, 1665, 1376, 1234, 1093, 1025; LC/MS *m/z* (TOF MS ES^+^): 761, 775, 789, 803, 817, 831, 845, 859, 873, 887, 901.

**SI, Figure 1.**
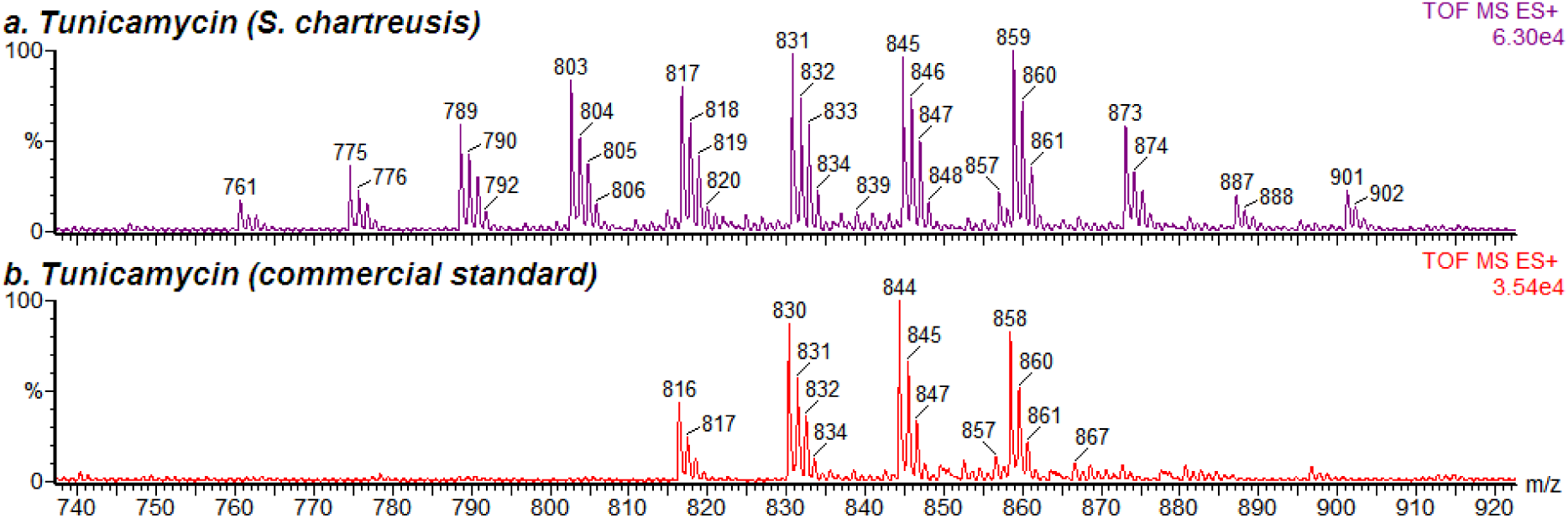
LC-MS Analysis of Tunicamycin Production by *S. chartreusis* NRRL 3882 by TOF-MS. (a) crude tunicamycin extracted from the *S. chartreusis* culture. (b) commercial tunicamycin standard (Sigma Aldrich, retention time 14-19 min.).

**SI, Table 1.** Tunicamycin extraction yield and purity.

| *S. chartreusis* strain | Culture Vol. (L) | tunicamycin isolated<sup>a</sup> (mg) | Sample<sup>a</sup> (mg/mL) | HPLC<sup>b</sup> (mg/mL) | Purity<sup>c</sup> (%) | tunicamycin /Liter<sup>d</sup> (mg/L) |
| --- | --- | --- | --- | --- | --- | --- |
| NRRL3882 | 12 | 687.3 | 1.25 | 1.0296 | 82.4 | 47.2 |
| NRRL3882 | 24 | 1066.4 | 1.30 | 1.0201 | 78.5 | 34.9 |
| NRRL3882 | 12 | 1483.1 | 1.40 | 0.4718 | 33.7 | 41.7 |
| NRRL3882 | 11 | 891.1 | 1.40 | 0.7899 | 56.4 | 45.7 |

<sup>a</sup>Crude sample; <sup>b</sup>Crude sample concentration injected into HPLC for analysis. tunicamycins dissolved in methanol; <sup>c</sup>Determined by HPLC, based on a standardised curve; <sup>d</sup>Purity and Culture Vol. were taken in consideration into the initial tunicamycins isolated. Average tunicamycins yield per liter of culture: $42 \pm 5$ mg

#### Large scale fermentation of *Streptomyces chartreusis* cells

Sterile TYD media (2 g Tryptone, 2 g yeast extract, 6 g glucose and 30 mg MgCl_2_.6H_2_O per litre) was added to 4 × 500 mL conical spring flasks. Each flask was inoculated with 50μl of the *Streptomyces chartreusis* spore stock (~ 5×10^7^ spores) and incubated at 28 °C with shaking on a rotary shaker (250 RPM) for 4 – 5 days. The flasks were then used to inoculate 90 L of TYD media in a Bioflow5000 fermenter at the University of East Anglia Fermentation Suite. Cells were fermented at 32 °C with an air flow rate of 0.25 L/L/min for 5 – 7 days before being harvested. Tunicamycin was extracted from resulting mycelial cake as described above.

#### Octa-*O*-acetyl-tunicamycin (tunicamycin-8OAc)

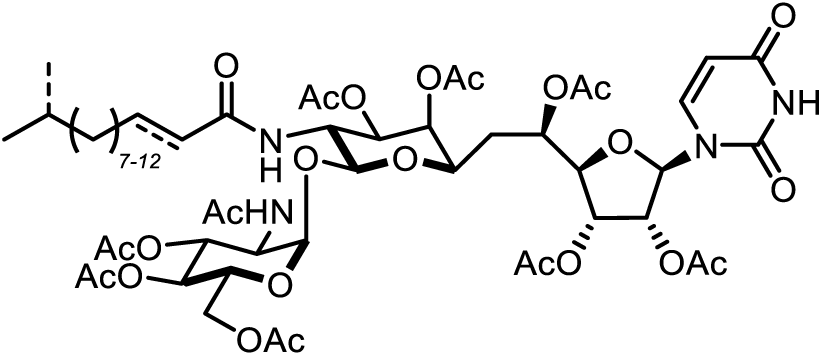

Crude tunicamycin (682 mg, 0.814 mmol) was dissolved in dry pyridine (5 mL) and Ac_2_O (3 mL). The reaction mixture was stirred for 18 h, concentrated *in vacuo* and purified by flash column chromatography (MeOH/DCM, 3:97) to afford the product as clear glass (782 mg, 0.667 mmol, 82 %); TLC: R_*f*_ 0.4 in methanol/dichloromethane (MeOH/DCM, 3:97); ^1^H NMR (500 MHz, CD_3_OD) δ ppm 7.48 (d, *J*_6,5_ = 8.0 Hz, 1 H, H-6^uracil^), 6.83 (dt, *J*_HC=CH trans_ = 14.2 Hz, *J* = 7.3 Hz, 1 H, C=C*H*-CH_2_), 5.87 (d, *J*_HC=CH trans_ = 15.5 Hz, 1 H, C=C*H*-CO), 5.81 (d, *J*_1’,2’_ = 5.1 Hz, 1 H, H-1’), 5.75 (d, *J*_5,6_ = 8.0 Hz, 1 H, H-5uracil), 5.56 (dd, *J*_3’,2’_ = 6.1 Hz, *J*_3’,4’_ = 5.5 Hz, 1 H, H-3’), 5.51 (dd, *J*_2’,1’_ = *J*_2’,3’_ = 5.3 Hz, 1 H, H-2’), 5.26 (dd, *J*_3”,2”_ = 10.6 Hz, *J*_3”,4”_ = 9.9 Hz, 1 H, H-3”), 5.27 (ddd, *J*_5’,6’_ = 9.7 Hz, *J*_5’,4’_ = 6.8 Hz, *J* = 3.6 Hz, 1 H, H-5’), 5.11 (dd, *J*_8’,9’_ = 9.7 Hz, *J*_8’,7’_ = 7.3 Hz, 1 H, H-8’), 5.07 (app t, *J*_4”,5”_ = 11.3 Hz, *J*_4”,3”_ = 3.2 Hz, 1 H, H-4”), 5.03 (dd, *J*_9’,10’_ = 3.6 Hz, *J*_9’,8’_ = 3.2 Hz, 1 H, H-9’), 4.98 (d, *J*_1”,2”_ = 4.9 Hz, 1 H, H-1”), 4.75 (d, *J*_11’,10’_ = 8.4 Hz, 1 H, H-11’), 4.33 (dd, *J*_6a”,6b”_ =11.1 Hz, *J*_6”,5”_ = 3.9 Hz, 1 H, H-6”), 4.34 (dd, *J*_10’,9’_ = 4.6 Hz, *J*_10’,11’_ = 3.6 Hz, 1 H, H-10’), 4.32 (ddd, *J*_5”,4”_ = 10.4, *J*_5”,6”_ = 2.9 Hz, *J*_5”,6”_ = 2.2 Hz, 1 H, H-5”), 4.20 (dd, *J*_4’,3’_ = 7.7 Hz, *J*_4’,5’_ = 3.6 Hz, 1 H, H-4’), 4.19 (dd, *J*_2”,3”_ = 7.2 Hz, *J*_2”,1”_ = 3.1 Hz, 1 H, H-2”), 4.19 (dd, *J*_6a”,6b”_ = 14.2 Hz, *J*_6”,5”_ = 2.6 Hz, 1 H, H-6”), 3.92 (ddd, *J* = 9.4 Hz, *J*_7’,6’_ = 3.8 Hz, *J*_7’,8’_ = 3.1 Hz, 1 H, H-7’), 2.22 (s, 3 H, C*H*_3_^Ac^), 2.17 (m, 2 H, -C*H*_2_CH=C), 2.14 (s, 6 H, 2 × C*H*_3_^Ac^), 2.10 (s, 3 H, C*H*_3_^Ac^), 2.06 (m, 2 H, H-6’), 2.04, 2.03, 1.98, 1.95, 1.89 (5 × s, 5 × 3 H, 5 × C*H*_3_^Ac^), 1.78 (ddd, *J*_6b’,a’_ = 14.8 Hz, *J*_6’,5’_ = 8 Hz, *J*_6’,7’_ = 3.3 Hz, 1 H), 1.55 (spt, *J* = 6.7 Hz, 1 H, -C*H*(CH_3_)_2_), 1.46 (quin, *J* = 6.8 Hz, 2 H, -C*H*_2_CH_2_CH=C), 1.23 - 1.37 (m, 14 H, -C*H*_2_^acyl^), 1.18 (dt, *J* = 13.1, 7.0 Hz, 2 H, C*H*_2_CH(CH_3_)_2_), 0.91, 0.89 (2 × s, 2 × 3 H, -CH(C*H*_3_)_2_); ^13^C NMR (126 MHz, CD_3_OD) δ ppm 173.2, 172.4, 172.3, 172.3, 172.0, 171.7, 171.5, 171.3, 171.2 (C=O^Ac^, C=O^NHAc^), 169.5 (C=O^acyl^), 165.9 (C-4 C=O), 151.8 (C-2 C=O), 147.7 (C=*C*H-CH_2_), 143.4 (C-6^uracil^), 124.2 (C=*C*H-CO), 103.5 (C-5^uracil^), 101.6 (C-11’), 100.0 (C-1’’), 91.1 (C-1’), 84.1 (C-4’), 73.5 (C-2’), 72.2 (C-3”), 72.2 (C-9’), 71.7 (C-7’), 70.9 (C-3’), 70.8, 70.3 (C-5’, C-8’), 69.8 (C-5”), 69.7 (C-4”), 63.0 (C-6”), 52.6 (C-2”), 51.8 (C-10’), 40.3 (-*C*H_2_CH(CH_3_)_2_), 33.2 (-*C*H_2_CH=C), 33.1 (C-6’), 30.3 - 31.1 (5x-*C*H_2_^acyl^), 29.4 (-*C*H_2_CH_2_CH=C), 29.2 (-*C*H(CH_3_)_2_), 28.6 (-*C*H_2_^acyl^), 23.0, 23.1 (-CH(*C*H_3_)_2_), 22.9 (-*C*H_3_^NHAc^), 21.1 (-*C*H_3_^Ac^), 20.7 (2 × -*C*H_3_^Ac^), 20.6, 20.6, 20.6, 20.6, 20.3 (5 × -*C*H_3_^Ac^); IR ν: 2927, 2361, 2341, 1745, 1696, 1540, 1369, 1219, 1031; MS *m/z* (ESI^+^): 1203 [(M+Na)^+^, 100%]; (ESI^−^): 1179 [(M+Cl)^−^, 100%]. Flanking peaks with mass ± 14 corresponded to 8 × CH_2_, 9 × CH_2_, 10 × CH_2_, and 11 × CH_2_. Full assignment was not possible due to the presence homologues with mass ± 14.

#### Tri-*N*-(tert-butoxylcarbonyl)-octa-*O*-acetyl-tunicamycin (tunicamycin-8OAc-3Boc)

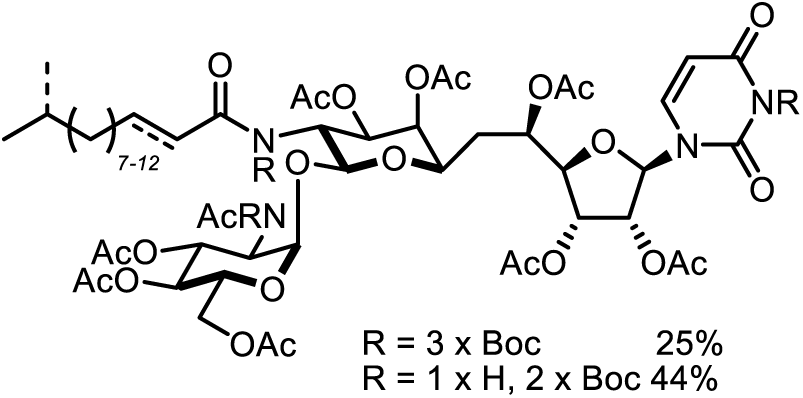

In order to cleave the lipid chain, the *tert*-butoxylcarbonyl (Boc) protecting group was added to the secondary amides at positions 3, 10’, and 2” to afford the tri-N-Boc-octa-*O*-acetylated tunicamycins. Amide cleavage usually involves the use of a strong acid or base and high temperatures, but these harsh conditions would be unsuitable to use in the presence of the uridine moiety as they would degrade the tunicamycins. Several methodologies have been published on how to remove the highly stable and unreactive acetyl group. One of them is Kunieda’s mild *N*-bocylation methodology. Attachment of Boc group to the secondary amide increases the electrophillicity of carbonyl, allowing the acetyl group to be readily cleaved in the presence of a base.

**Tunicamycin-8OAc** (101 mg, 0.086 mmol) was dissolved in dry THF (1.5 mL) with the addition of 4-(dimethylamino)pyridine (10.5 mg, 0.086 mmol) and di-*tert*-butyl dicarbonate (187.7 mg, 0.86 mmol). The reaction mixture was heated to 60 °C with stirring for 4 h, and subsequently another portion of di-*tert*-butyl dicarbonate (187.7 mg, 0.86 mmol) was added to the reaction mixture, with stirring continued for an additional 2 h. After a total of 6 h, the reaction mixture was checked by TLC (EtOAc/Petrol, 6:4). This which showed the formation of two products, **tunicamycin-8OAc-3Boc** (R_*f*_ 0.3) and (**tunicamycin-8OAc-2Boc**) (R_*f*_ 0.1). The reaction mixture was concentrated *in vacuo* and purified by flash column chromatography (EtOAc/Petrol, 6:4). **tunicamycin-8OAc-3Boc** (31.1 mg, 0.021 mmol, 25 %) was obtained as a yellow glass and **tunicamycin-8OAc-2Boc** (52.4 mg, 0.038 mmol, 44%) as a yellow oil; **tunicamycin-8OAc-3Boc**: TLC: R_*f*_ 0.5 in ethyl acetate/petrol (EtOAc/Petrol, 6:4); 1H NMR (500 MHz, CDCl_3_) δ ppm 7.48 (d, *J*_6,5_ = 8.0 Hz, 1 H, H-6uracil), 6.89 (dt, *J*_HC=CH trans_ = 15.1 Hz, *J* = 6.9 Hz, 1 H, C=C*H*-CH_2_), 6.82 (dt, *J*_HC=CH trans_ = 14.5 Hz, *J* = 7.6 Hz, 1 H, C=C*H*-CH_2_), 6.39 (d, *J*_HC=CH trans_ = 15.1 Hz, 1 H, C=C*H*CO), 6.27 (d, *J*_HC=CH trans_ = 15.4 Hz, 1 H, C=C*H*-CO), 6.11 (m, 1 H, C-H_anomeric_), 5.85 (m, 1 H, CH_anomeric_, H-5_uracil_), 5.83 (d, *J* = 8.2 Hz, 1 H, H-5_uracil_), 5.61 (dd, *J* = 11.5 Hz, *J* = 3.3 Hz, 1 H), 5.53 (dd, *J* = 11.3 Hz, *J* = 3.5 Hz, 1 H), 5.49 (d, *J* = 8.2 Hz, 1 H), 5.43 (m, 1 H), 5.29 – 5.35 (m, 1 H), 5.09 – 5.25 (m, 4 H), 5.01 – 4.10 (m, 1 H), 4.99 (d, *J* = 9.1 Hz), 4.94 (dd, *J* = 11.5 Hz, *J* = 3.3 Hz, 1 H), 4.91 (s, 1 H), 4.52 – 4.60 (m, 1 H), 4.30 – 4.39 (m, 1 H), 4.17 (d, *J* = 10.4 Hz, *J* = 2.2 Hz, 1 H), 4.05 – 4.10 (m, 1 H), 3.76 (dd, *J* = 8.8 Hz, *J* = 3.5 Hz, 1 H), 3.68 (dd, *J* = 9.9 Hz, *J* = 2.0 Hz, 1 H), 2.34 (s, 1 H), 2.29 (s, 2 H, C*H*_3NHAc_), 2.27 (s, 1 H, C*H*_3_^NHAc^), 1.87 – 2.22 (m, 24H, 8 × CH_3_^Ac^), 1.59, 1.56, 1.55, 1.53, 1.52 (5 × s, 27 H, 9 × CH_3_^Boc^), 1.40 (m, 13 H), 1.08 – 1.18 (m, 2 H, CH_2_^acyl^), 0.86, 0.85 (2 × s, 2 × 3 H, C*H*_3_^acyl^); ^13^C NMR (126 MHz, CD_3_OD) δ ppm 177.4, 177.5, 170.8,170.7, 170.1, 169.9, 169.7, 169.6, 169.5, 169.4, 169.1 (C=O), 168.2 (C-4 C=O), 159.8 (C-2 C=O), 153.0, 152.9, 157.7, 152.7, 152.1, 148.3, 148.1, 147.3, 139.1, 139.0, 138.9 (C=*C*H-CH_2_, C-6_uracil_), 124.4, 123.9 (C=*C*H-CO), 103.5, 103.2 (C-1’’), 97.8, 87.7, 86.9, 87.0, 86.9, 86.3, 82.4, 82.3, 72.1, 70.4, 70.2, 70.1, 69.6, 69.5, 69.4, 69.2, 69.1, 68.8, 68.0, 67.9, 61.5, 61.4, 57.657.0, 54.8, 39.0, 38.5, 36.6, 34.3, 32.7, 32.5, 32.4, 31.9, 29.9, 29.6, 29.5, 29.4, 29.3, 29.2, 28.2, 28.0, 27.9, 27.8, 27.6, 27.4, 22.6, 20.9, 20.9, 20.7, 20.6, 20.5, 20.4 (C-1’), 84.1 (C-4’), 73.5 (C-2’), 72.2 (C-3”), 72.2 (C-9’), 71.7 (C-7’), 70.9 (C-3’), 70.8, 70.3 (C-5’, C-8’), 69.8 (C-5”), 69.7 (C-4”), 63.0 (C-6”), 52.6 (C-2”), 51.8 (C-10’), 40.3 (-*C*H_2_CH(CH_3_)_2_), 33.2 (-*C*H_2_CH=C), 33.1 (C-6’), 30.3 - 31.1 (5 × C, 5 × - *C*H_2_^acyl^), 29.4 (-*C*H_2_CH_2_CH=C), 29.2 (-*C*H(CH_3_)_2_), 28.6 (-*C*H_2_^acyl^), 23.0, 23.1 (2 × C, - CH(*C*H_3_)_2_), 22.9 (-*C*H_3_^NHAc^), 21.1 (-*C*H_3_^Ac^), 20.7 (2 × C, 2 × -*C*H_3_^Ac^), 20.6, 20.6, 20.6, 20.6, 20.3 (5 × C, 5 × -*C*H3^Ac^) IR ν: 2928, 2361, 2341, 1743, 1686, 1369, 1218, 1143, 1029; LRMS *m/z* (ESI^+^): 1503 [(M+Na)^+^, 100%]; (ESI^−^): 1515 [(M+Cl)^−^, 100%]. Flanking peaks with mass ± 14 corresponded to 8 × CH_2_, 9 × CH_2_, 10 × CH_2_, and 11 × CH_2_.

#### 10’,2′-Di-*N*-Boc-α-D-glucosamine-(1″-11′)-tunicamyl uracil (TUN-Boc,Boc)

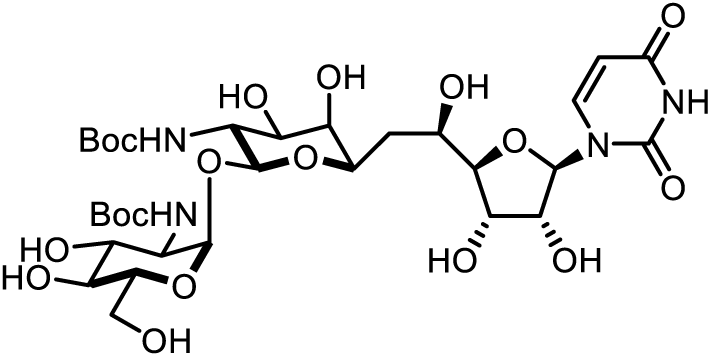

**Tunicamycin-8OAc-3Boc** (134 mg, 0.091 mmol) was dissolved in MeOH:H_2_O (v/v, 3:1) with the addition of TEA (25 equiv. 2.27 mmol, 317 µl). The reaction mixture was heated to 71 °C and reaction progress monitored by TLC (1:2:6, W/*i*PrOH/EtOAc). After 43 h the mixture was directly purified by preparative scale HPLC (retention time 9.5 min). Product containing fractions were pooled and lyophilized to afford **TUN-Boc,Boc** (36.4 mg, 0.047 mmol, 52%) as white amorphous powder; TLC: R_*f*_ 0.3 in water/isopropanol/ethyl acetate (W/*i*POH/EtOAc, 1:2:6); R_*f*_ = 0.3 (H_2_O/iPrOH/EtOAc, 1/2/7); [α]_D_^20^ = +54.9 ± 0.3 (c 1, MeOH); Mp (amorphous) 177.4−181.2 °C; ^1^H NMR (500 MHz, CD_3_OD) δ ppm 7.91 (d, *J* = 8.1 Hz, 1H, H-6), 5.93 (d, *J* = 5.9 Hz, 1H, H-1’), 5.75 (d, *J* = 8.1 Hz, 1H, H-5), 4.99 (s, 1H, H-1”), 4.70 (d, *J* = 7.9 Hz, 1H, H-11’), 4.24 – 4.16 (m, 2H, H-2’, H-3’), 4.05 – 3.97 (m, 2H, H-5’, H-5”), 3.86 (t, *J* = 3.3 Hz, 1H, H-4’), 3.81 (dd, *J* = 11.7, 1.8 Hz, 1H, H-6”), 3.77 – 3.65 (m, 3H, H-7’, H-9’, H-6”), 3.64 (d, *J* = 3.1 Hz, 1H, H-8’), 3.62 (d, *J* = 4.9 Hz, 2H, H-2”, H- 3”), 3.49 (t, *J* = 9.6 Hz, 1H, H-10’), 3.37 – 3.33 (m, 1H, H-4”), 2.12 – 2.05 (m, 1H, H-6’), 1.57 – 1.49 (m, 1H, H-6’), 1.47 (s, 9H, C*H*_3_), 1.45 (s, 9H, C*H*_3_); ^13^C NMR (126 MHz, CD_3_OD) δ ppm 166.2 (C-4), 158.7 (C=O^Boc^), 158.5 (C=O^Boc^), 152.6 (C-2), 142.8 (C-6), 103.1 (C-5), 101.4 (C-11’), 100.6 (C-1”), 89.7 (C-1’), 89.5 (C-4’), 80.7 (*C*-(CH3)3), 80.3 (*C*-(CH3)3), 75.5 (C-2’), 74.5 (C-5”), 73.6 (C-3”), 72.7, 72.6, 72.4 (C-7’, C-9’, C-4”), 72.3 (C-8’), 70.9 (C-3’), 68.4 (C-5’), 63.2 (C-6”), 56.6 (C-4”), 55.8 (C-10’), 35.9 (C-6’), 29.1((*C*H_3_)_3_), 28.8 ((*C*H_3_)_3_); IR (neat) ν: 3367 (N-H, O-H), 2979 (=C-H), 2930 (-C-H), 1684 (C=O); LRMS m/z (ESI^+^): 789 [(M+Na)^+^, 100%]; HRMS m/z (ESI^+^): calc. C_31_H_50_N_4_O_18_Na (M+Na)^+^ = 789.3012, found 789.3017.

#### α-D-*N*-acetylglucosamine-(1”-11’)-N-acetyl tunicamyl uracil (TUN-Ac,Ac)

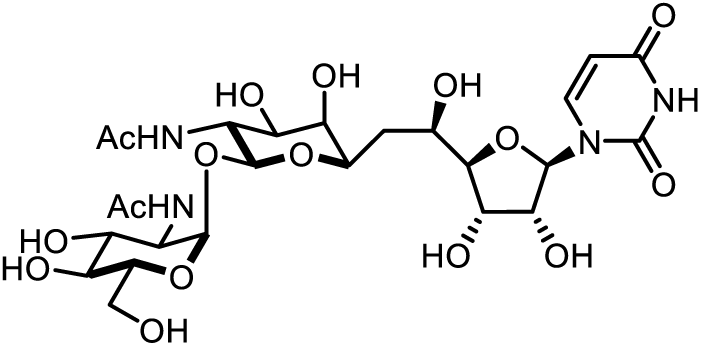

**Tunicamycin-8OAc-3Boc** (127 mg, 0.086 mmol) was dissolved in dry MeOH (5 mL) and cooled to 0° C. NaOMe was added to a final concentration of 0.01 M and reaction progress monitored by TLC (1:3:6, W/*i*POH/EtOAc). The reaction was neutralized after 4 h by addition of Dowex 50W X8 H^+^ resin in parts until pH 7. The mixture was then filtered, the resin washed with methanol and the combined organics and concentrated *in vacuo*. The resulting solid was dissolved in TFA (1 mL) and stirred at RT for 1 h. The TFA was coevaporated with toluene and the crude material then redissolved in MeOH (5 mL) and Ac_2_O (1 mL). The reaction mixture was stirred at RT for 12 h, neutralized to pH 6-7 with Dowex Marathon A - OH resin and stirred for an additional 1 h. The reaction mixture was filtered, concentrated *in vacuo* and purified by flash column chromatography (W/*i*POH/EtOAc, 1:2:2) to afford **TUN-Ac,Ac** (13.1 mg, 0.020 mmol, 23%) as yellow glass; TLC: R_*f*_ 0.3 (W/*i*POH/EtOAc, 1:2:2); [α]_D23_ = +50.7 (c = 0.7, H_2_O); ^1^H NMR (500 MHz, CD_3_OD) δ ppm 7.76 (d, *J*_6,5_ = 7.9 Hz, 1 H, H-6^uracil^), 5.84 (d, *J*_1’,2’_ = 7.9 Hz, 1 H, H-1’), 5.83 (d, *J*_5,6_ = 5.4 Hz, 1 H, H5^uracil^), 4.98 (d, *J*_1”,2”_ = 3.5 Hz, 1 H, H-1”), 4.58 (d, *J*_11’,10’_ = 8.5 Hz, 1 H, H-6^uracil^), 4.25 (dd, *J*_2’,1’_ = 5.4 Hz, *J*_2’,3’_ = 9.1 Hz, 1 H, H-2’), 4.22 (dd, *J*_3’,4’_ = 3.5 Hz, *J*_3’,2’_ = 5.7 Hz, 1 H, H-3’), 4.08 – 4.03 (m, 2 H, H-4’, H-5’), 3.87 (dd, *J*_10’,9’_ = 10.7 Hz, *J*_10’,11’_ = 8.5 Hz, 1 H, H-10’), 3.82 – 3.82 (m, 1 H, H-4”), 3.80 (dd, *J*_2”,1”_ = 3.8 Hz, *J*_2”,3”_ = 10.7, 1 H, H-2”), 3.77 (d, *J* = 10.1 Hz, 1 H, H-7’), 3.73 – 3.67 (m, 2 × 1 H, H-8’, H-6b”), 3.70 (dd, *J*_3”,2”_ = 10.7 Hz, *J*_3”,4”_ = 3.2 Hz, 1 H, H-3”), 3.68 – 3.41 (m, 2 H, H-6”a, H-9’), 3.44 (app t, *J*_5”,6a”_ = 9.8 Hz, 1 H, H-5”), 1.98, 1.94 (2 × s, 2 × 3H, 2 × -C*H*_3_^NHAc^), 1.94 (dd, *J*_6b’,6a’_ = 6.6 Hz, *J*_6b’,5’_ = 3.2 Hz, 1 H, H-6b’), 1.57 (app t, *J*_6a’,6b’_ = *J*_6a’,5’_ = 13.2 Hz, 1 H, H- 6a’); ^13^C NMR (126 MHz, CD_3_OD) δ ppm 174.5, 174.1 (C=O^NHAc^), 166.2 (C-4, C=O), 151.8 (C-2, C=O), 141.9 (C-6^uracil^), 102.5 (C-5^uracil^), 99.8 (C-11’), 98.3 (C-1”), 88.5 (C-1’), 87.2 (C-4’), 73.4 (C-2’), 72.6 (C-4”), 71.3 (C-7’), 71.1, 70.4, 69.8 (C-3”, C-8’, C-9’), 69.6 (C-5”), 68.9 (C-3’), 67.0 (C-5’), 60.4 (C-6”), 53.4 (C-2”), 52.8 (C-10’), 33.5 (C-6’), 22.2, 22.1 (2 × -C*H*_*3*_^NHAc^); IR (neat) ν: 3344, 2362, 2341, 2110, 1636, 1371, 1216; LRMS *m/z* (ESI^+^): 673.26 [(M+Na)^+^, 23%]; (ESI^−^): 649.23 [(M-H)^−^, 100%]; HRMS *m/z* (ESI^+^): calc. for C_25_H_38_N_4_NaO_16_ (M+Na)^+^ = 673.2175, found 673.2195.

#### α-D-glucosamine-(1”-11’)-tunicamyl uracil dihydrochloride (TUN)

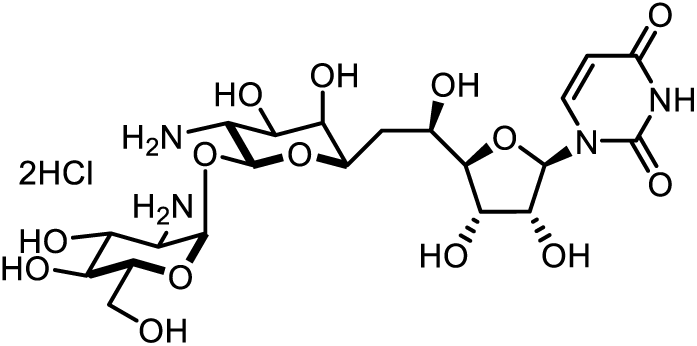

To a solution of **TUN-Boc,Boc** (1.50 mg, 0.002 mmol) in DCM (120 μL) was added TFA (0.393 mmol 30 µL). The reaction mixture was stirred at room temperature for 1 h, with reaction progress monitored by TLC (1:2:2, W/*i*POH/EtOAc). When the reaction was complete, the reaction mixture was concentrated *in vacuo*. The crude product was washed twice with H_2_O and DCM, the aqueous fraction collected and concentrated *in vacuo.* The dried crude product was then redissoved in 1 M HCl (1 mL), stirred for 1 h at room temperature and lyophilized to yield the product **TUN** (1.20 mg, 99%). [α]_D_^20^ = +60.1 ± 0.2 (c 1, H_2_O); ^1^H NMR (700 MHz, D_2_O) δ ppm 7.82 (d, *J* = 8.2 Hz, 1 H, H-6), 5.87 (d, *J* = 8.2 Hz, 1 H, H-5) 5.86 (d, *J* = 5.3 Hz, 1 H, H-1’), 5.53 (d, *J* = 3.4 Hz, 1 H, H-1”), 5.00 (d, *J* = 8.3 Hz, 1 H, H-11’), 4.31 - 4.26 (m, 2 H, H-2’, H-3’), 4.06 (td, *J* = 2.6, 11.1 Hz, 1 H, H-5’), 3.94 (dd, *J* = 3.3, 11.0 Hz, 1 H, H-9’), 3.92 - 3.87 (m, 3 H, H-7’, H-3”, H-5”), 3.84 (d, *J* = 3.2 Hz, 1 H, H-8’), 3.79 (dd, *J* = 3.8, 12.5 Hz, 1 H, H-6”), 3.70 (dd, *J* = 2.2, 12.4 Hz, 1 H, H6”), 3.57 (t, *J* = 9.6 Hz, 1 H, H-4”), 3.39 (dd, *J* = 3.5, 10.8 Hz, 1 H, H-2”), 3.31 (dd, *J* = 8.4, 11.0 Hz, 1 H, H-10’), 1.97 (ddd, *J* = 2.0, 10.4, 14.6 Hz, 1 H, H-6’), 1.70 - 1.64 (dtd, *J* = 2.8, 11.2 Hz, 1 H, H-6’); ^13^C NMR (176 MHz, CD_3_OD) δ ppm 166.17 (C-4), 151.8 (C-2), 142.0 (C-6), 102.4 (C-5), 99.4 (C-11’), 97.0 (C-1”), 88.7 (C-1’), 86.9 (C-4’), 73.4 (C-2’), 73.2 (C-5”), 71.8 (C-7”), 69.5 (C-8’), 69.3 (C- 3”), 69.2 (C-9’), 68.9 (C-4’), 68.7 (C-3’), 66.9 (C-5’), 59.8 (C-6”), 53.8 (C-2”), 53.0 (C-10’), 33.3 (C-6’); IR (neat) ν: 3295 (N-H, O-H), 3057 (=C-H), 2922 (-C-H), 1673 (C=O), 1263 (C-N), 1109 (C-O), 1064 (C-O); LRMS m/z (ESI^+^): 567 [(M+H)^+^, 100%]; HRMS m/z (ESI^+^): calc. C_21_H_35_N_4_O_14_ (M+H)^+^ = 567.2144, found 567.2136.

#### General protocol for preparing tunicamycin analogues

HATU (2.5 equiv) was added to a solution of the appropriate carboxylic acid (2.5 equiv), EDC (2.5 equiv) and DIPEA (2.5 equiv) in dry DMF. The reaction mixture was stirred at RT for 10 min, followed by the addition of **TUN** (1 equiv) and DIPEA (2.5 equiv). The reaction mixture was stirred at RT for 2 ~ 4 h, diluted with a mixture of ACN/*i*POH/Water (1:1:1) and purified by preparative HPLC.

#### Di-*N*-citronoyl-tunicamycin (TUN-Cit,Cit)

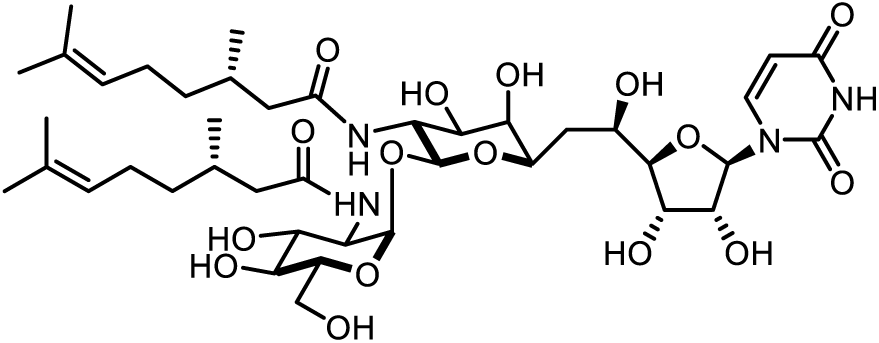

The product was purified by HPLC (0.1% FA and 5% - 100% acetonitrile gradient in 24 mins on C18 preparative column) and the desired product was eluted at 14 min. The lyophilised product was washed with DCM and MilliQ water and resulted in 5.1mg of the final product, 54% yield. R_*f*_ = 0.4 (1/3/6, H_2_O/*i*PrOH/EtOAc); [α]_D_^20^ = +45.2 ± 0.2 (c 0.4, MeOH); ^1^H NMR (500 MHz, CD_3_OD) δ ppm 7.91 (d, *J* = 8.1 Hz, 1H, H-6), 5.92 (d, *J* = 5.9 Hz, 1H, H-1’), 5.75 (d, *J* = 8.1 Hz, 1H, H-5), 5.11 (td, *J* = 7.0, 1.0 Hz, 2H, H-5”’), 4.96 (d, *J* = 3.4 Hz, 1H, H-1”), 4.58 (d, *J* = 8.5 Hz, 1H, H-11’), 4.24 – 4.15 (m, 2H, H-2’, H-3’), 4.06 – 3.95 (m, 3H, H-5’, H- 10’, H-5”), 3.91 (dd, *J* = 10.6, 3.5 Hz, 1H, H-2”), 3.87 – 3.80 (m, 2H, H-4’, H-6”), 3.76 (appt dd, *J* = 9.5, 1.9 Hz, 1H, H-7’), 3.71 – 3.60 (m, 4H, H-8’, H-9’, H-3”, H-6”), 3.33 (appt d, *J* = 9.4 Hz, 1H, H-4”), 2.24 (m, 2H, H-1”’), 2.17 – 1.88 (m, 9H, H-6’, H-1”’, H-2”’, H-4”’), 1.67 (s, 6H, H-7”’), 1.61 (s, 6H, H-8”’), 1.52 (m, 1H, H-6’), 1.37 (m, 2H, H-3”’), 1.23 (m, 2H, H- 3”’), 0.96 (d, *J* = 6.4 Hz, 6H, H-9”’); ^13^C NMR (126 MHz, CD_3_OD) δ ppm 176.6, 176.0 (N- C=O^aliphatic chain^), 166.2 (C-4), 152.6 (C-2), 142.8 (C-6), 132.3, 132.2 (C-6”’), 125.6, 125.5 (C- 5”’), 103.0 (C-5), 101.6 (C-11’), 99.9 (C-1”), 89.8 (C-1’), 89.6 (C-4’), 75.5 (C-2’), 74.4 (C- 5”), 73.1, 73.0 (C-8’, C-9’), 72.7 (C-4”), 72.5 (C-7’), 72.2 (C-3”), 70.8 (C-3’), 68.3 (C-5’), 63.2 (C-6”), 54.7 (C-2”), 54.3 (C-10’), 45.4, 44.9 (C-1”’), 38.6, 38.5 (C-3”’), 35.9 (C-6’), 31.8, 31.7 (C-2”’), 26.7, 26.6 (C-4”’), 25.9 (C-7”’), 19.6 (C-9”’), 17.8 (C-8”’); IR (neat) ν: 3291 (O-H), 2966 (C-H), 2928 (C-H), 1700 (C=O), 1638 (C=O),1541 (C=C), 1092 (C-N); LRMS m/z (ESI^−^): 915 [(M+FA-H)^−^, 100%]; HRMS m/z (ESI^−^): calc. C_41_H_65_N_4_O_16_ (M-H)^−^ = 869.4401, found 869.4407.

#### Di-*N*-heptanoyl tunicamycin (TUN-7,7)

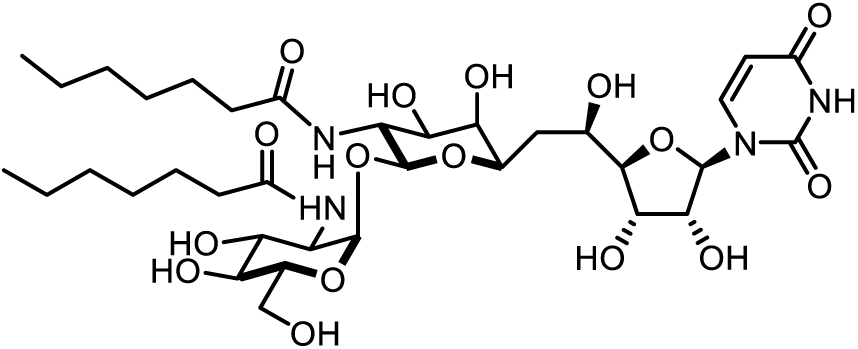

The product was purified by HPLC (0.1% FA and 5% - 100% acetonitrile gradient in 24 mins on C18 preparative column) and the desired product was eluted at 12 min. The lyophilised product was washed with DCM and MilliQ water and resulted in 4.8 mg of the final product, 39% yield. R_*f*_ = 0.4 (1/3/6, H_2_O/*i*PrOH/EtOAc); [α]_D20_ = +26.8 ± 0.7 (c 0.2, MeOH); ^1^H NMR (500 MHz, CD_3_OD) δ ppm 7.92 (d, *J* = 8.1 Hz, 1H, H-6), 5.93 (d, *J* = 6.0 Hz, 1H, H-1’), 5.75 (d, *J* = 8.1 Hz, 1H, H-5), 4.94 (d, *J* = 3.4 Hz, 1H, H-1”), 4.61 (d, *J* = 8.5 Hz, 1H, H-11’), 4.24 – 4.16 (m, 2H, H-2’, H-3’), 4.06 – 3.99 (m, 2H, H-5’, H-5”), 3.95 (dd, *J* = 10.2, 8.6 Hz, 1H, H-10’), 3.90 (dd, *J* = 10.6, 3.5 Hz, 1H, H-2”), 3.87 – 3.81 (m, 2H, H-4’, H-6”), 3.77 (appt br d, *J* = 9.1 Hz, 1H, H-7’), 3.71 – 3.62 (m, 4H, H-8’, H9’, H-3”, H-6”), 3.34 (appt s, 1H, H-4”), 2.38 – 2.02 (m, 5H, 2 × CH_2_^fatty acyl^, H-6’), 1.69 – 1.49 (m, 5H, 2 × CH_2_^fatty acyl^, H-6’), 1.41 – 1.28 (m, 15H, C*H*_*2*_^fatty acyl^), 0.92 (t, *J* = 6.8 Hz, 6H, C*H*_*3*_^fatty acyl^); ^13^C NMR (126 MHz, CD_3_OD) δ ppm 177.2, 176.6 (N-C=O^fatty acyl^), 166.2, (C-4), 152.6, (C-2), 142.8, (C-6), 103.0, (C-5), 101.3, (C-11’), 99.9, (C-1”), 89.8, (C-1’), 89.6, (C-4’), 75.5 (C-2’), 74.4 (C-5”), 73.0 (C-8’, C-9’), 72.6 (C-4”), 72.5 (C-7’), 72.1 (C-3”), 70.9 (C-3’), 68.3 (C-5’), 63.3 (C-6”), 54.8 (C-2”), 54.5 (C-10’), 37.8, 37.2 (CO*C*H_2_-^fatty acyl^), 35.9 (C-6’), 32.9, 32.8, 30.2, 27.01, 26.8, 23.7 (*C*H_2_-^fatty acyl^), 14.4 (*C*H_3_^fatty acyl^); IR (neat) ν: 3305 (O-H), 2927 (C-H), 2856 (C-H), 1682 (C=O), 1645 (C=O),1552 (C=C), 1467 (CH2), 1376 (CH3), 1259 (C-O), 1094 (C-N); LRMS m/z (ESI^+^): 813 [(M+Na)^+^, 100%]; HRMS m/z (ESI^+^): calc. C_35_H_58_N_4_O_16_ (M+Na)^+^ = 813.3740, found 813.3708.

#### Di-*N*-octanoyl-tunicamycin (TUN-8,8)

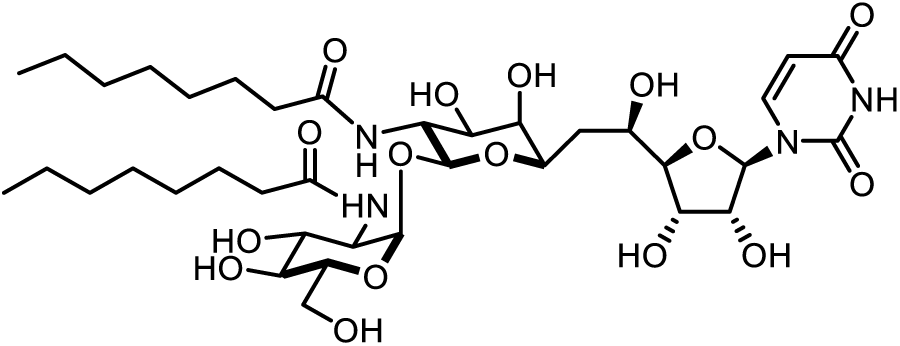

The product was purified by HPLC (0.1% FA and 5% - 100% acetonitrile gradient in 24 mins on C18 preparative column) and the desired product was eluted at 13.5 min. The lyophilised product was washed with DCM and MilliQ water and resulted in 2.5 mg of the final product, 63% yield. R_*f*_ = 0.3 (1/3/6, H_2_O/*i*PrOH/EtOAc); [α]_D_^20^ = +57.4 ± 0.4 (c 0.2, MeOH); ^1^H NMR (500 MHz, CD_3_OD) δ ppm 7.91 (d, *J* = 8.1 Hz, 1H, H-6), 5.92 (d, *J* = 6.0 Hz, 1H, H-1’), 5.75 (d, *J* = 8.1 Hz, 1H, H-5), 4.94 (d, *J* = 3.4 Hz, 1H, H-1”), 4.60 (d, *J* = 8.5 Hz, 1H, H-11’), 4.24 – 4.15 (m, 2H, H-2’, H-3’), 4.06 – 3.98 (m, 2H, H-5’, H-5”), 3.95 (dd, *J* = 10.0, 8.6 Hz, 1H, H-10’), 3.90 (dd, *J* = 10.6, 3.4 Hz, 1H, H-2”), 3.87 – 3.80 (m, 2H, H-4’, H-6”), 3.76 (dd, *J* = 10.6, 1.6 Hz, 1H, H-7’), 3.70 – 3.61 (m, 4H, H-8’, H-9’, H-3”, H-6”), 3.34 (appt d, *J* = 4.0 Hz, 1H, H-4”), 2.38 – 2.14 (m, 4H, 2 × CH_2_^fatty acyl^), 2.10 (m, 1H, H-6’), 1.70 – 1.57 (m, 4H, 2 × CH_2_^fatty acyl^), 1.53 (ddd, *J* = 13.9, 11.4, 2.2 Hz, 1H, H-6’), 1.41 – 1.24 (appt br m, 16H, C*H2*fatty acyl), 0.91 (t, *J* = 6.9 Hz, 6H, C*H*_*3*_^fatty acyl^); ^13^C NMR (126 MHz, CD_3_OD) δ ppm 177.2, 176.6 (N-C=O^fatty acyl^), 166.1 (C-4), 152.6 (C-2), 142.8 (C-6), 103.1 (C-5), 101.3 (C-11’), 99.9 (C-1”), 89.8 (C-1’), 89.6 (C-4’), 75.5 (C-2’), 74.4 (C-5”), 73.0 (C-8’, C-9’), 72.6 (C-4”), 72.5 (C-7’), 72.1 (C-3”), 70.9 (C-3’), 68.3 (C-5’), 63.3 (C-6”), 54.8 (C-2”), 54.5 (C-10’), 37.8, 37.2 (CO*C*H_2_-^fatty acyl^), 35.9 (C-6’), 33.0, 33.0, 30.5, 30.3, 30.3, 27.0, 26.8, 23.7 (*C*H_2_-^fatty acyl^), 14.4 (*C*H_3_^fatty acyl^); IR (neat) ν: 3297 (O-H), 2957 (C-H), 2925 (C-H), 2853 (C-H), 1684 (C=O), 1644 (C=O), 1556 (C=C), 1469 (CH2), 1258 (C-O), 1091 (C-N), 1016 (=C-H); LRMS m/z (ESI^−^): 931 [(M+TFA-H)-, 100%]; HRMS m/z (ESI^+^): calc. C_37_H_62_N_4_O_16_Na (M+Na)^+^ = 841.4053, found 841.4045.

#### Di-*N*-nonanoyl-tunicamycin (TUN-9,9)

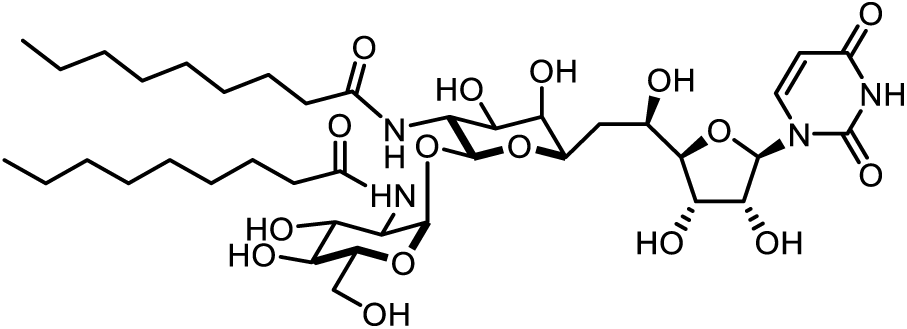

The product was purified by HPLC (0.1% FA and 5% - 100% acetonitrile gradient in 24 mins on C18 preparative column) and the desired product was eluted at 15 min. The lyophilised product was washed with DCM and MilliQ water and resulted in 3.4 mg of the final product, 85% yield. R_*f*_ = 0.4 (1/3/6, H_2_O/*i*PrOH/EtOAc); [α]_D20_ = +53.8 ± 1.2 (c 0.3, MeOH); ^1^H NMR (500 MHz, CD_3_OD) δ ppm 7.93 (d, *J* = 8.1 Hz, 1H, H-6), 5.95 (d, *J* = 5.9 Hz, 1H, H-1’), 5.77 (d, *J* = 8.1 Hz, 1H, H-5), 4.96 (d, *J* = 3.0 Hz, 1H, H-1”), 4.62 (d, *J* = 8.6 Hz, 1H, H-11’), 4.26 – 4.18 (m, 2H, H-2’, H-3’), 4.08 – 4.00 (m, 2H, H-5’, H-5”), 3.97 (t, *J* = 9.1 Hz, 1H, H-10’), 3.92 (dd, *J* = 10.7, 3.2 Hz, 1H, H-2”), 3.89 – 3.81 (m, 2H, H-4’, H-6”), 3.78 (appt br d, *J* = 9.8 Hz, 1H, H-7’), 3.73 – 3.63 (m, 4H, H-8’, H-9’, H-3”, H-6”), 3.36 (appt d, *J* = 4.2 Hz, 1H, H-4”), 2.41 – 2.16 (m, 4H, 2 × CH_2_^fatty acyl^), 2.12 (appt br t, *J* = 12.1 Hz, 1H, H-6’), 1.71 – 1.59 (m, *J* = 6.7 Hz, 4H, 2 × CH_2_^fatty acyl^), 1.55 (appt br t, *J* = 12.6 Hz, 1H, H-6’), 1.34 (s, 20H, C*H*_*2*_^fatty acyl^), 0.93 (t, *J* = 6.5 Hz, 6H, C*H*_*3*_^fatty acyl^); ^13^C NMR (126 MHz, CD_3_OD) δ ppm 177.2, 176.6 (N-C=O^fatty acyl^), 166.1 (C-4), 152.6 (C-2), 142.8 (C-6), 103.0 (C-5), 101.3 (C-11’), 99.9 (C-1”), 89.8 (C-1’), 89.6 (C-4’), 75.5 (C-2’), 74.4 (C-5”), 73.1, 73.0 (C-8’, C-9’), 72.6 (C-4”), 72.5 (C-7’), 72.1 (C-3”), 70.9 (C-3’), 68.3 (C-5’), 63.3 (C-6”), 54.7 (C-2”), 54.5 (C-10’), 37.8, 37.2 (CO*C*H_2_-^fatty acyl^), 35.9 (C-6’), 33.1, 33.0, 30.6, 30.5, 30.4, 30.3, 27.0, 26.8, 23.8 (*C*H_2_-^fatty acyl^), 14.5 (*C*H_3_^fatty acyl^); IR (neat) ν: 3301 (O-H), 2923 (C-H), 2852 (CH), 1738 (C=O), 1646 (C=O), 1544 (C=C), 1420 (CH_2_), 1366 (CH_3_), 1229 (C-O), 1092 (CN), 1015 (=C-H); LRMS m/z (ESI^−^): 891 [(M+FA-H)^−^, 100%]; HRMS m/z (ESI^−^): calc. C_39_H_65_N_4_O_16_ (M-H)^−^ = 845.4401, found 845.4412.

#### Di-*N*-decanoyl-tunicamycin (TUN-10,10)

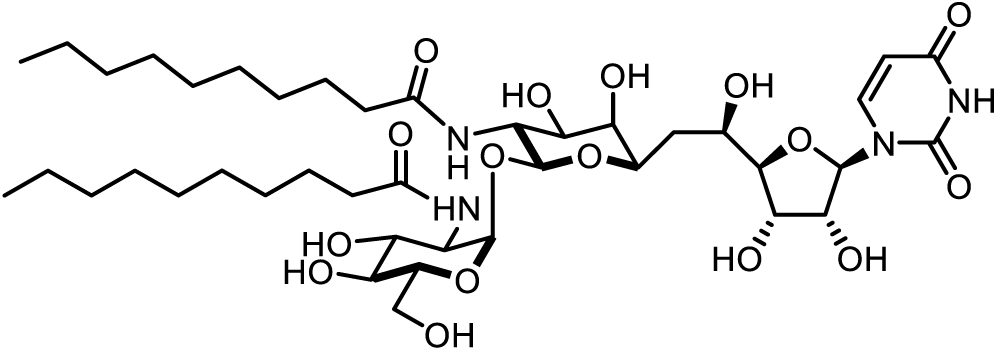

The product was purified by HPLC (0.1% FA and 5% - 100% acetonitrile gradient in 24 mins on C18 preparative column) and the desired product was eluted at 16.5 min. The lyophilised product was washed with DCM and MilliQ water and resulted in 3 mg of the final product, 31% yield. R_*f*_ = 0.4 (1/3/6, H_2_O/*i*PrOH/EtOAc); [α]_D20_ = +38.0 ± 0.6 (c 0.3, MeOH); ^1^H NMR (500 MHz, CD_3_OD) δ ppm 7.91 (d, *J* = 8.1 Hz, 1H, H-6), 5.92 (d, *J* = 6.0 Hz, 1H, H-1’), 5.75 (d, *J* = 8.1 Hz, 1H, H-5), 4.93 (d, *J* = 3.4 Hz, 1H, H-1”), 4.59 (d, *J* = 8.5 Hz, 1H, H-11’), 4.23 – 4.16 (m, 2H, H-2’, H-3’), 3.97 – 3.92 (m, 2H, H-5’, H-5”), 3.90 (appt t, *J* = 8.5 Hz, 1H, H-10’), 3.84 (dd, *J* = 10.6, 3.4 Hz, 1H, H-2”), 3.87 – 3.80 (m, 2H, H-4’, H-6”), 3.76 (appt br dd, *J* = 10.7, 1.7 Hz, 1H, H-7’), 3.71 – 3.61 (m, 4H, H-8’, H-9’, H-3”, H-6”), 3.33 (appt d, *J* = 5.8 Hz, 1H, H-4”), 2.38 – 2.02 (m, 4H, 2 × CH_2_^fatty acyl^), 2.10 (m, 1H, H-6’), 1.69 – 1.49 (m, 4H, 2 × CH_2_^fatty acyl^), 1.55 (m, 1H, H-6’), 1.30 (s, 24H, C*H*_*2*_^fatty acyl^), 0.90 (t, *J* = 6.9 Hz, 6H, C*H*_*3*_^fatty acyl^); ^13^C NMR (126 MHz, CD_3_OD) δ ppm 177.17, 176.6 (NC=O^fatty acyl^), 166.2 (C-4), 152.6 (C-2), 142.8 (C-6), 103.1 (C-5), 101.3 (C-11’), 100.0 (C-1”), 89.8 (C-1’), 89.6 (C-4’), 75.5 (C-2’), 74.4 (C-5”), 73.1, 73.0 (C-8’, C-9’), 72.6 (C-4”), 72.5 (C-7’), 72.1 (C-3”), 70.9 (C-3’), 68.3 (C-5’), 63.3 (C-6”), 54.7 (C-2”), 54.5 (C-10’), 37.8, 37.2 (CO*C*H_2_-^fatty acyl^), 35.9 (C-6’), 33.1, 30.9, 30.8, 30.7, 30.6, 30.5, 27.0, 26.8, 23.8 (*C*H_2_-^fatty acyl^), 14.5 (*C*H_3_^fatty acyl^); IR (neat) ν: 3305 (O-H), 2922 (C-H), 2851 (C-H), 1683 (C=O), 1645 (C=O),1551 (C=C), 1468 (CH_2_), 1260 (C-O), 1094 (C-N), 1017 (=C-H); LRMS m/z (ESI^−^): 919 [(M+FA-H)^−^, 100%]; HRMS m/z (ESI^+^): calc. C_41_H_70_N_4_O_16_Na (M+Na)^+^ = 897.4679, found 897.4666.

### Di-*N*-undecanoyl-tunicamycin (TUN-11,11)

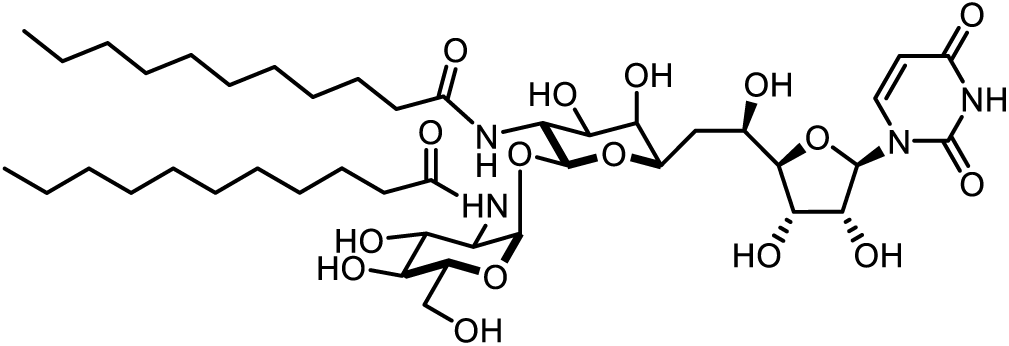

The product was purified by HPLC (0.1% FA and 5% - 100% acetonitrile gradient in 24 mins on C18 preparative column) and the desired product was eluted at 18 min. The lyophilised product was washed with DCM and MilliQ water and resulted in 3mg of the final product, 30% yield. R_*f*_ = 0.4 (1/3/6, H_2_O/*i*PrOH/EtOAc); [α]_D_^20^ = +30.9 ± 0.4 (c 0.25, MeOH); ^1^H NMR (500 MHz, CD_3_OD) δ ppm 7.91 (d, *J* = 8.1 Hz, 1H, H-6), 5.92 (d, *J* = 6.0 Hz, 1H, H-1’), 5.75 (d, *J* = 8.1 Hz, 1H, H-5), 4.93 (d, *J* = 3.4 Hz, 1H, H-1”), 4.60 (d, *J* = 8.5 Hz, 1H, H-11’), 4.23 – 4.15 (m, 2H, H-2’, H-3’), 4.05 – 3.98 (m, 2H, H-5’, H-5”), 3.95 (m, 1H, H-10’), 3.90 (dd, *J* = 10.6, 3.4 Hz, 1H, H-2”), 3.86 – 3.80 (m, 2H, H-4’, H-6”), 3.76 (appt br dd, *J* = 10.7, 1.8 Hz, 1H, H-7’), 3.71 – 3.61 (m, 4H, H-8’, H-9’, H-3”, H-6”), 3.34 (m, 1H, H-4”), 2.37 – 2.14 (m, 4H, 2 × CH_2_^fatty acyl^), 2.13 – 2.05 (m, 1H, H-6’), 1.68 – 1.56 (m, 4H, 2 × CH_2_^fatty acyl^), 1.57 – 1.49 (m, 1H, H-6’), 1.30 (appt br s, 32H, C*H*_*2*_^fatty acyl^), 0.90 (t, *J* = 6.9 Hz, 6H, C*H*_*3*_^fatty acyl^); ^13^C NMR (126 MHz, CD_3_OD) δ ppm 177.2, 176.6, (NC=O^fatty acyl^), 166.2 (C-4), 152.7 (C-2), 142.8 (C-6), 103.0 (C-5), 101.3 (C-11’), 100.0 (C-1”), 89.8 (C-1’), 89.6 (C-4’), 75.5 (C-2’), 74.4 (C-5”), 73.1, 73.0 (C-8’, C-9’), 72.6 (C-4”), 72.5 (C-7’), 72.1 (C-3”), 70.9 (C-3’), 68.3 (C-5’), 63.3 (C-6”), 54.8 (C-2”), 54.5 (C-10’), 37.8, 37.2 (CO*C*H_2_-^fatty acyl^), 35.9 (C-6’), 33.1, 30.9, 30.8, 30.7, 30.6, 30.5, 30.4, 27.0, 26.9, 23.8 (*C*H_2_-^fatty acyl^), 14.5 (*C*H_3_^fatty acyl^); IR (neat) ν: 3297 (O-H), 2956 (C-H), 2921 (C-H), 2852 (C-H), 1738 (C=O), 1719 (C=O), 1680 (C=C), 1645 (C=O), 1550 (N-H), 1468 (CH_2_), 1366 (CH_3_), 1229 (C-O-C), 1217 (C-OH), 1260 (C-O), 1092 (C-N), 1017 (=C-H); LRMS m/z (ESI^−^): 947 [(M+FA-H)-, 100%]; HRMS m/z (ESI^−^): calc. C_43_H_73_N_4_O_16_ (M-H)^−^ = 901.5027, found 901.5015.

#### Di-*N*-dodecanoyl-tunicamycin (TUN-12,12)

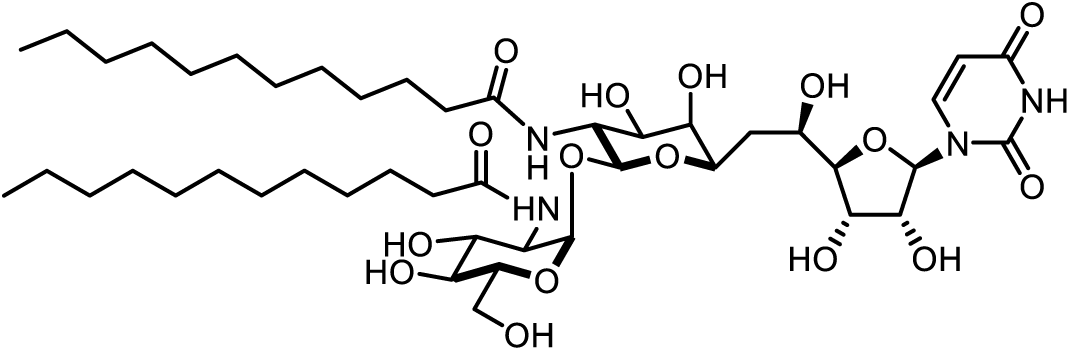

The product was purified by HPLC (0.1% FA and 5% - 100% acetonitrile gradient in 24 mins on C18 preparative column) and the desired product was eluted at 20.5 min. The lyophilised product was washed with DCM and MilliQ water and resulted in 3mg of the final product, 29% yield. R_*f*_ = 0.4 (1/3/6, H_2_O/*i*PrOH/EtOAc); [α]_D_^20^ = +15.9 ± 0.4 (c 0.25, MeOH); ^1^H NMR (500 MHz, CD_3_OD) δ 7.92 (d, *J* = 8.1 Hz, 1H, H-6), 5.93 (d, *J* = 5.9 Hz, 1H, H-1’), 5.76 (d, *J* = 8.1 Hz, 1H, H-5), 4.94 (d, *J* = 3.5 Hz, 1H, H-1”), 4.60 (d, *J* = 8.5 Hz, 1H, H-11’), 4.24 – 4.17 (m, 2H, H-2’, H-3’), 4.08 – 3.99 (m, 2H, H-5’, H-5”), 3.96 (m, *J* = 8.6 Hz,1H, H-10’), 3.91 (dd, *J* = 10.6, 3.4 Hz, 1H, H-2”), 3.88 – 3.80 (m, 2H, H-4’, H-6”), 3.77 (appt br dd, *J* = 11.1, 1.8 Hz, 1H, H-7’), 3.72 – 3.61 (m, 4H, H-8’, H-9’, H-3”, H-6”), 2.39 – 2.15 (m, 4H, 2 × CH_2_^fatty acyl^), 2.11 (m, 1H, H-6’), 1.70 – 1.57 (m, 4H, 2 × CH_2_^fatty acyl^), 1.54 (m, 1H, H-6’), 1.40-1.28 (appt broad m, 32H, C*H*_*2*_^fatty acyl^), 0.91 (t, *J* = 6.9 Hz, 6H, C*H*_*3*_^fatty acyl^); ^13^C NMR (126 MHz, CD_3_OD) δ ppm 177.17, 176.57 (N-C=O^fatty acyl^), 166.16 (C-4), 152.63 (C-2), 142.76 (C-6), 103.06 (C-5), 101.35 (C-11’), 100.04 (C-1”), 89.77 (C-1’), 89.62 (C-4’), 75.52 (C-2’), 74.37 (C-5”), 73.08, 73.04 (C-8’, C-9’), 72.60 (C-4”), 72.49 (C-7’), 72.10 (C-3”), 70.90 (C-3’), 68.33 (C-5’), 63.27 (C-6”), 54.76 (C-2”), 54.45 (C-10’), 37.82, 37.19 (CO*C*H_2_-^fatty acyl^), 35.94 (C-6’), 30.87, 30.83, 30.79, 30.77, 30.68, 30.65, 30.63, 30.55, 27.05, 26.83, 23.79 (*C*H_2_-^fatty acyl^), 14.48 (*C*H_3_^fatty acyl^); IR (neat) ν: 3297 (O-H), 2956 (C-H), 2921 (C-H), 2851 (C-H), 1682 (C=O), 1646 (C=O), 1556 (C=C), 1468 (CH_2_), 1260 (C-O), 1092 (C-N), 1016 (=C-H); LRMS m/z (ESI^−^): 976 [(M+FA-H)-, 100%]; HRMS m/z (ESI^+^): calc. C_45_H_78_N_4_O_16_Na (M+Na)^+^ = 953.5305, found 953.5334.

#### Heptaacetyl-tunicamyl-uracil (3)

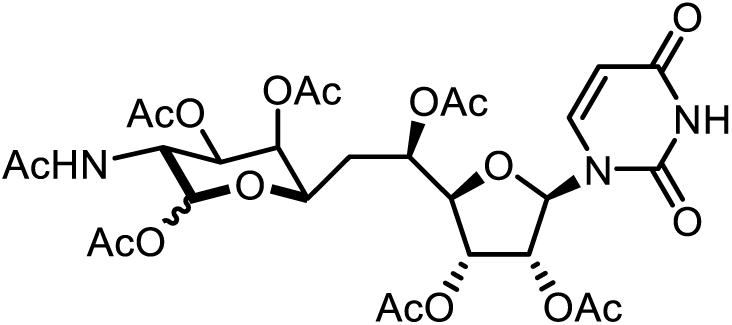

Crude tunicamycin (183 mg, 0.218 mmol) was suspended in 3 M aq. HCl (2 mL) and stirred under reflux at 105 °C for 135 min. The solvent was then co-evaporated with toluene *in vacuo*. The resulting residue was re-dissolved in dry pyridine (3 mL) and Ac_2_O (2 mL) and stirred for 18 h at RT. The reaction mixture was then concentrated *in vacuo* and purified by flash column chromatography (MeOH/EtOAc, 1:19) to afford Heptaacetyl-tunicamyl-uracil **3** (98.4 mg, 0.141 mmol, 64 %); TLC: R_*f*_ 0.3 in methanol/ethyl acetate (MeOH/EtOAc, 1:19); ^1^H NMR (500 MHz, CDCl_3_) δ ppm 8.53 (d, *J* = 1.00 Hz, 1 H, N-H^uracil,β^), 8.43 (d, *J* = 0.95 Hz, 1 H, N-H^uracil,α^), 7.22 (d, *J*_6,5_ = 8.2 Hz, 1 H, H-6^uracil,α^), 7.19 (d, *J*_6,5_ = 8.2 Hz, 1 H, H-6^uracil,β^), 6.13 (d, *J*_11’,10’_ = 3.5 Hz, 1 H, H-11^’α^), 5.90 (d, *J*_1’,2’_ = 5.4 Hz, 1 H, H-1^’α^), 5.83 (d, *J*_1’,2’_ = 3.8 Hz, 1 H, H-1^’β^), 5.80 (d, *J* = 2.2 Hz, 1 H, H-5^uracil,α^), 5.78 (dd, *J* = 2.1 Hz, *J*_5,6_ = 8.0, 1 H, H-5^uracil,β^), 5.64 (d, *J*_11’,10’_ = 8.8 Hz, 1 H, H-11^’β^), 5.55 (d, *J*_N-H,10’_ = 9.5 Hz, 1 H, N-H^Ac,β^), 5.44 (d, *J*_N-H,10’_ = 9.1 Hz, 1 H, N-H^Ac,α^), 5.42 (dd, *J*_3’,4’_ = 5.4 Hz, *J*_3’,2’_ = 10.4 Hz, 1 H, H-3^’β^), 5.36 (dd, *J*_3’,4’_ = 5.0 Hz, *J*_3’,2’_ = 5.9 Hz, 1 H, H-3^’α^), 5.33 (app t, *J*_2’,1’_ = *J*_2’,3’_ = 5.7 Hz, 1H, H-2^’α^), 5.30 (app t, *J*_2’,1’_ = *J*_2’,3’_ = 6.0 Hz, 1 H, H-2^’β^), 5.25 (d, *J*_9’,8’_ = *J*_9’,10’_ = 2.8 Hz, 1 H, H-9^’α^), 5.22 (dd, *J*_8’,7’_ = 3.2 Hz, *J*_8’,9’_ = 6.6 Hz, 1 H, H-8^’β^), 5.19 (dd, *J*_5’,4’_ = 1.9 Hz, *J*_5’,6’_ = 3.5 Hz, 1H, H-5^’β^), 5.19 (dd, *J*_8’,7’_ = 1.5 Hz, *J*_8’,9_ = 3.5 Hz, 1 H, H-8^’α^), 5.12 (ddd, *J* = 2.8 Hz, *J*_5’,4’_ = 5.0 Hz, *J*_5’,6’_ = 7.9 Hz, 1 H, H-5^’α^), 5.08 (dd, *J*_9’,8’_ = 3.50 Hz, *J*_9’,10’_ = 11.3 Hz, 1 H, H-9^’β^), 4.72 (ddd, *J*_10’,11’_ = 4.1 Hz, *J*_10’,N-H_ = 9.5 Hz, *J*_10’,9’_ = 11.4 Hz, 1 H, H-10^’α^), 4.42 (ddd, *J*_10’9’_ = 7.6 Hz, *J*_10’,NH’_ = 9.5 Hz, *J*_10’,9’_ = 11.3 Hz, 1 H, H-10^’β^), 4.13 (app t, *J*_4’,5’_ = *J*_4’,3’_ = 4.7 Hz, 1 H, H-4^’β^), 4.09 (app t, *J*_4’,5’_ = *J*_4’,3’_ = 4.7 Hz, 1 H, H-4^’α^), 4.05 - 4.07 (m, 1 H, H-7^’α^), 3.90 (dd, *J*_7’,8’_ = 2.5 Hz, *J*_7’,6’_ = 9.7 Hz, 1 H, H-7^’β^), 2.20 (s, 3 H, -CH_3_^Ac,α^), 2.20 (s, 3 H, -CH_3_^Ac,β^), 2.18, 2.13 (2 × s, 2 × 3H, 2 × - CH_3_^Ac,α^), 2.13, 2.12, 2.11, 2.11 (4 × s, 4 × 3 H, 4 × -CH_3_^Ac,β^), 2.09 (2 × s, 2 × 3 H, 2 × -CH_3_^Ac,α^), 2.04 (s, 3 H, -CH_3_^Ac,α^), 2.02 (s, 3 H, -CH_3_^Ac,β^), 1.99 - 2.01 (m, 1 H, H-6^’β^), 1.97 - 1.99 (m, 1 H, H-6^’α^), 1.96 (s, 3 H, -CH_3_NH^Ac,α^), 1.94 (s, 3 H, -CH_3_NH^Ac,β^), 1.74 (ddd, *J*_6’,7’_ = 3.5 Hz, *J*_6’,5’_ = 6.9 Hz, *J*_6’a,6b’_ = 14.9 Hz, 1 H, H-6^’β^), 1.57 (ddd, *J*_6’,7’_ = 1.9 Hz, *J*_6’,5’_ = 8.2 Hz, *J*_6’a,6’b_ = 16.7 Hz, 1 H, H-6^’α^); ^13^C NMR (126 MHz, CDCl_3_) δ ppm 173.6, 171.2, 170.7, 170.6, 170.6, 170.3, 170.1, 170.0, 169.7, 169.6, 169.4, 169.3, 169.3, 169.1 (C=O^NHAc,α,β^, C=O^Ac,α,β^), 162.5 (C-4 C=O^β^), 162.5 (C-4 C=O^α^), 149.8 (C-2 C=O^α^), 149.8 (C-2 C=O^β^), 140.1 (C-6^uracil,β^), 139.8 (C-6^uracil,α^), 103.4 (C-5^uracil,α^), 103.3 (C-5^uracil,β^), 93.0 (C-11’^β^), 91.1 (C-11’^α^), 88.7 (C-1’^β^), 88.1 (C-1’^α^), 82.5 (C-4’^β^), 82.5 (C-4’^α^), 72.4 (C-2’^α^), 72.4 (C-2’^β^), 71.0 (C-7’^β^), 70.5 (C-9’^β^), 69.6, 69.6, 69.3, 68.7, 68.1 (5 × s, 7 × C, C-3’^α^, C-3’^β^, C-5’^α^, C-5’^β^, C-8’^α^, C-8’^β^, C-9’^α^), 49.6 (C-10’^β^), 46.7 (C-10^’α^), 32.5 (C-6^’α^), 31.5 (C-6^’β^), 23.3 (-CH_3_NH^Ac,β^), 23.2 (-CH_3_^NHAc,α^), 20.3 – 21.0 (CH_3_^Ac,α,β^); IR: 3370, 1736, 1710, 1697, 1651, 1635, 1540, 1520, 1370; LRMS *m/z* (ESI^+^): 722 [(M+Na)^+^, 100%]; (ESI^−^): 734 [(M+Cl)^−^, 100%]. HRMS *m/z* (ESI^+^): calc. for C_29_H_37_N_3_NaO_17_ (M+Na)^+^ = 722.2015, found 722.2023.

#### N-acetyl-tunicamyl-uracil (2)

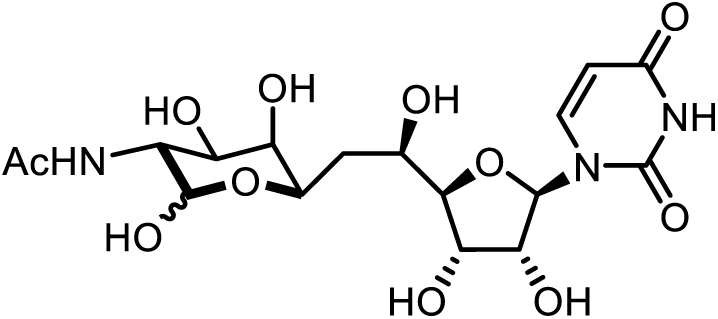

Heptaacetyl-tunicamyl-uracil **3** (41.4 mg, 0.059 mmol) was dissolved in dry MeOH (5 mL) and cooled to 0° C. NaOMe was added to a final concentration of 0.01 M and the reaction mixture stirred for 3 h. The reaction mixture was then neutralized with Dowex 50W X8 H^+^ resin, filtered and concentrated *in vacuo*. Purification by flash column chromatography (W/*i*POH/EtOAc, 1:2:2) afforded the product **2** (25.9 mg, 0.058 mmol, 98 %); TLC: R_*f*_ 0.5, 0.6 (W/*i*POH/EtOAc, 1:2:2); [α]_D_^23^ = +12 (c 1, H_2_O); ^1^H NMR (500 MHz, D_2_O) δ ppm 7.78 (d, *J*_6,5_ = 8.2 Hz, 1 Hα, H-6^uracil,α^), 7.76 (d, *J*_6,5_ = 8.2 Hz, 1 H^β^, H-6^uracil,β^), 5.87 (m, 1 H^α^ + 1 H^β^, H-1^’α^, H-1^’β^), 5.84 (d, *J*_5,6_ = 8.2 Hz, 1 H^α^, H-5^uracil,α^), 5.82 (d, *J*_5,6_ = 8.2 Hz, 1 H^β^, H-5^uracil,β^), 5.13 (d, *J*_11’,10’_ = 3.8 Hz, 1 H^α^, H-11^’α^), 4.58 (d, *J*_11’,10’_ = 8.5 Hz, 1 H^β^, H-11^’β^), 4.23 - 4.27 (m, 2 H^α^ + 2H^β^, H-2^’α^, H-2^’β^, H-3^’α^, H-3^’β^), 4.19 (d, *J* = 9.1 Hz, 1 H^α^, H-7^’α^), 4.04 (dd, *J*_10’,9’_ = 11.0 Hz, *J*_10’,11_ = 3.8 Hz, 1 H^α^, H-10^’α^), 4.00 (dt, *J* = 10.7 Hz, *J* = 3.2 Hz, 1 H^β^, H-5^’β^), 3.92 - 3.97 (m, 2 H^α^ + 1 H^β^, H-4^’α^, H-4^’β^, H-5^’α^), 3.88 (dd, *J*_9’,10’_ = 11.4 Hz, *J*_9’,8’_ = 3.2 Hz, 1 H^α^, H-9^’α^), 3.79 (d, *J*_8’,9’_ = 3.5 Hz, 1 H^α^, H-8^’α^), 3.78 (dd, *J*_10’,9’_ = 10.7 Hz, *J*_10’,9’_ = 8.2 Hz, 1 H^β^, H- 10^’β^), 3.75 (dd, *J*_7’,6’_ = 8.5 Hz, *J*_7’,8_ = 1.0 Hz, 1 H^β^, H-7^’β^), 3.73 (d, *J*_8’,9’_ = 3.5 Hz, 1 H^β^, H-8^’β^), 3.69 (dd, *J*_9’,10’_ =10.7 Hz, *J*_9’.8’_ = 3.2 Hz, 1 H^β^, H-9^’β^), 1.98 (s, 3 H^α^ + 3 H^β^, -CH_3_^NHAc,β^, - CH_3_^NHAc,α^), 1.86 - 1.96 (m, 1 H^α^ + 1 H^β^, H-6a^’α^, H-6a^’β^), 1.54 - 1.63 (m, 1 H^α^ + 1 H^β^, H-6b^’α^, H-6b^’β^); ^13^C NMR (126 MHz, D_2_O) δ ppm 175.0, 174.7 (C=O^NHAc,β,α^), 166.22 (C-4 C=O^α+β^), 151.9 (C-2 C=O^α+β^), 141.8 (C-6^uracil,α+β^), 102.5 (C-5^uracil,α+β^), 95.3 (C-11’^β^), 90.9 (C-11’^α^), 88.1 (C-1’^α+β^), 87.1, 87.1 (C-4’^α^, C-4’^β^), 73.4, 73.4 (C-2’^α^, C-2’^β^), 71.2, 71.1, 70.8, 70.4 (C-7’^β^, C- 8’^β^, C-9’^β^, C-8’^α^), 68.9, 68.9 (C-3’^α^, C-3’^β^), 67.5 (C-9’^α^), 67.1, 67.0 (C-5’^α^, C-5’^β^), 66.3 (C-7^’β^), 53.6 (C-10^’β^), 50.2 (C-10^’α^), 33.7, 33.6 (C-6^’α^, C-6^’β^), 23.3, 22.2 (-CH_3_^NHAc,α^, -CH_3_^NHAc,β^); IR ν: 3362, 1638, 1410, 1264, 1072; LRMS *m/z* (ESI^+^): 470 [(M+Na)^+^, 100%]; (ESI^−^): 482 [(M+Cl)^−^, 100%]; HRMS *m/z* (ESI^+^): calc. for C17H25N3NaO11 (M+Na)^+^ = 470.1381, found 470.1367.

#### *N*-Octanoyl-*N’*-acetyl tunicamycin (TUN-8,Ac)

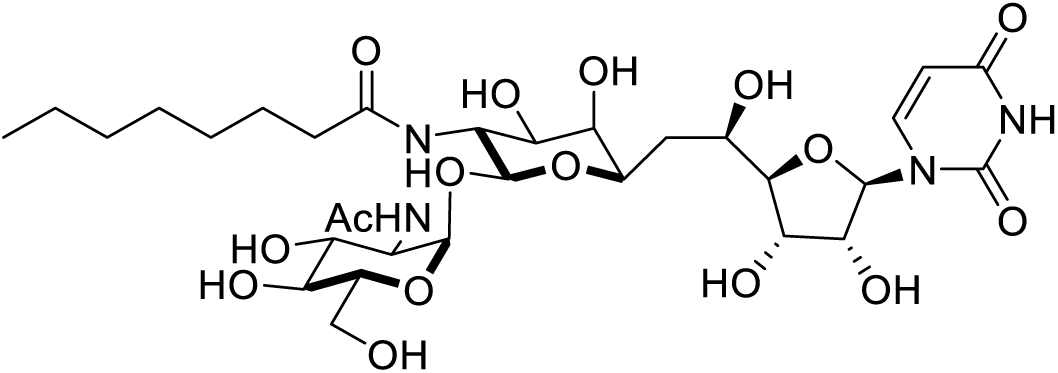

A 25% solution of NaOMe in MeOH (100 uL) was added to anhydrous MeOH (1 mL). An aliquot of the resulting NaOMe solution (100 uL) was then added the mixture of **tunicamycin- 8OAc-2Boc** (130 mg, 0.095 mmol) in a MeOH (4 mL) under argon and the resulting orange solution stirred for 4 h. The reaction mixture was carefully quenched with DOWEX 50WX8 H^+^ form resin and the resin was then removed by filtration and the filtrate concentrated *in vacuo*. The resulting yellow solid was dissolved in TFA (2 mL) and stirred at ambient temperature for 2 h. The reaction mixture was then concentrated *in vacuo* and azeotroped with toluene (2 × 1 mL) and MeOH (2 × 1 mL), followed by drying under high vacuum overnight. In a separate flask, octanoic acid (0.204 mmol) and HATU (70.8 mg, 0.186 mmol) were dissolved in dry DMF (1 mL) and cooled to 0 °C. DIPEA (62 uL, 0.354 mmol) was added and the resulting yellow solution stirred at 0 °C for 10 min. A solution of the crude diamine in DMF (1 mL) was added and the resulting yellow solution stirred at ambient temperature for 18 h. The reaction mixture was concentrated *in vacuo* and purified by column chromatography (SiO_2_, 1:3:6 H_2_O:IPA:EtOAc). This was further purified by HPLC and product containing fractions were lyophilized to yield pure **TUN-8,Ac** (3 mg, 0.004 mmol, 4%). ^1^H NMR (400 MHz, CD_3_OD) δ ppm 7.94 (d, 1H, *J* = 8.1 Hz, H6), 5.95 (d, 1H, *J* = 6.0 Hz, H1’), 5.78 (d, 1H, *J* = 8.1 Hz, H5), 4.96 (d, 1H, *J* = 3.4 Hz, H1”), 4.63 (d, 1H, *J* = 8.5 Hz, H11’), 4.26-4.21 (m, 2H, H2’ + H3’), 4.04-3.97 (m, 2H, H5’ + H5”), 3.96-3.86 (m, 4H, H10’ + H2” + H4’ + H6”), 3.73- 3.64 (m, 5H, H7’ + H8’ + H9’ + H-3” + H6”), 3.34 (obscured by solvent, 1H, H4”), 2.47-2.17 (m, 2H, Oct-Hα), 2.13-2.05 (m, 1H, H6’), 2.03 (s, 1H, NHAc), 1.63-1.50 (m, 3H, Lipid-Hβ + H6’), 1.40-1.20 (m, 8H, Lipid-Hγ + Hδ + Hε + Hζ + Hη), 0.92-0.89 (m, 3H, Lipid Hθ); ^13^C NMR (151 MHz, CD_3_OD) δ 177.22, 173.57 (N-C=O^fatty acyl^), 166.12 (C-4), 152.62 (C-2), 142.74 (C-6), 103.05 (C-5), 101.21 (C-11’), 99.78 (C-1”), 89.82 (C-1’), 89.60 (C-4’), 75.49 (C-2’), 74.30 (C-5”), 73.06, 72.96 (C-8’, C-9’), 72.51 (C-4”), 72.49 (C-7’), 72.10 (C-3”), 70.90 (C-3’), 68.34 (C-5’), 63.27 (C-6”), 54.95 (C-2”), 54.49 (C-10’), 37.75, 35.92 (CO*C*H_2_-^fatty acyl^), 32.90 (C-6’), 30.43, 30.18, 27.00, 23.68, 23.10 (*C*H_2_-^fatty acyl^), 14.40 (*C*H_3_^fatty acyl^);); IR ν: 3367, 3192, 1667, 1588, 1368, 1318, 1098, 1019; HRMS (ESI^+^) Calcd for C_31_H_50_O_16_N_4_Na [M+Na]^+^ 757.31140, found 757.31085.

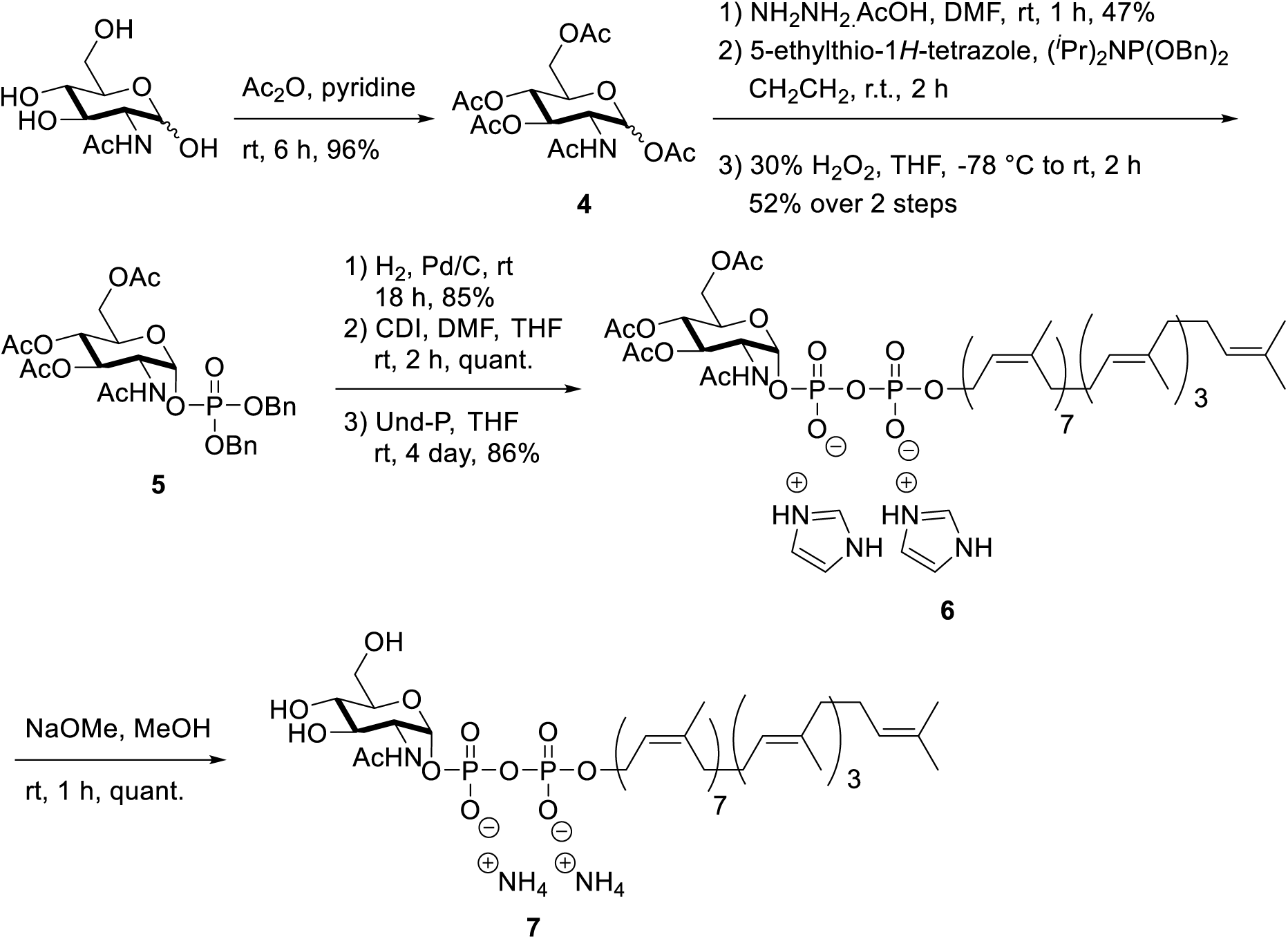

#### SI, Scheme 1. The Synthesis of GlcNAc-PP-Und (7) (the protocol was followed from(Gale et al., 2014; Liu et al., 2014)). 1,3,4,6-Tetra-*O*-acetyl-*N*-acetyl-D-glucosamine (4)

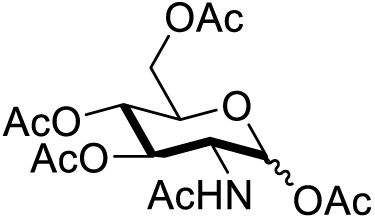

*N*-Acetyl-D-glucosamine (5.0 g, 22.6 mmol) was suspended in pyridine (50 mL) and Ac_2_O (25 mL) and stirred for 6 h at ambient temperature. The reaction mixture was then concentrated *in vacuo* and azeotroped with toluene (3 × 20 mL). The resulting oil was dissolved in CH_2_Cl_2_ (100 mL), washed with 1 M HCl (50 mL) and brine (50 mL), dried over anhydrous Na2SO4 and concentrated *in vacuo* to yield product **4** as a white foam (8.47 g, 96%). ^1^H NMR (CDCl_3_, 400 MHz) δ 6.15 (d, 1H, *J* = 3.7 Hz, H1), 5.65 (d, 1H, *J* = 9.1 Hz, NH), 5.25-5.16 (m, 2H, H3 + H4), 4.50-4.44 (m, 1H, H2), 4.23 (dd, 1H, *J* = 12.5, 4.1 Hz, H6), 4.05 (dd, 1H, *J* = 12.5, 2.4 Hz, H6’), 3.98 (ddd, 1H, *J* = 9.7, 4.0, 2.3, H5), 2.18 (s, 3H), 2.07 (s, 3H), 2.04 (s, 3H), 2.03 (s, 3H), 1.92 (s, 3H); ^13^C NMR (125 MHz, CDCl_3_) δ 171.9, 170.9, 170.2, 169.3, 168.8, 90.6, 70.9, 69.9, 67.7, 61.7, 51.2, 23.2, 21.1, 20.9, 20.7; LRMS (ES) Calcd for C_16_H_23_NNaO_10_ [M+Na]^+^ 412.12, found 412.12.

#### *N*-acetyl-3,4,6-Tris-*O*-acetyl-1-(dibenzyl phosphate)-α-D-glucosamine (5)

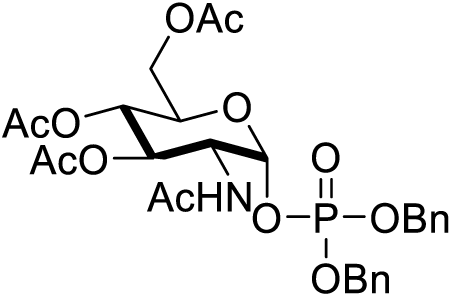

Acetate **4** (3.0 g, 7.71 mmol) was dissolved in dry DMF (50 mL). Hydrazine acetate (1.05 g, 11.4 mmol) was added and the resulting solution stirred at ambient temperature for 2 h. The reaction mixture was diluted with EtOAc (100 mL) and washed with H_2_O (100 mL) and saturated aqueous NaHCO3 (100 mL). The combined aqueous washings were back extracted with EtOAc (2 × 50 mL) and the combined organic extracts dried over anhydrous Na_2_SO_4_ and concentrated *in vacuo* to yieldthe anomeric lactol as a colourless oil (1.25 g, 47%). The lactol (1.25 g, 3.6 mmol) was then dissolved in anhydrous CH_2_Cl^2^ (30 mL) and added rapidly via syringe to a vigorously stirred suspension of 5-ethylthio-1*H*-tetrazole (2.20 g, 16.9 mmol) and dibenzyl-*N*,*N*’-diisopropylphosphoramidite (3.73 g, 10.8 mmol) in anhydrous CH_2_Cl_2_ (30 mL) under argon at ambient temperature. The reaction mixture became homogeneous within a few min. After 2 h, the mixture was diluted with CH_2_Cl_2_ (40 mL) and washed with saturated sodium bicarbonate (50 mL), water (50mL) and brine (50 mL). The organic solution was dried over anhydrous sodium sulfate and concentrated *in vacuo* to yield the phosphite as a colorless oil. The product was dissolved in THF (60 mL) and cooled to −78 °C. Hydrogen peroxide (30%, 6 mL) was added dropwise via syringe to the vigorously stirred solution. After the addition was complete, the ice bath was removed and the mixture was allowed to warm to ambient temperature over 1.5 h. The reaction mixture was then diluted with ice-cold saturated sodium sulfite (15 mL), followed by EtOAc (30 mL), and stirred for 5 min. The organic layer was concentrated *in vacuo* and the crude redissolved in EtOAc (100 mL). This was washed with saturated NaHCO_3_ (50 mL), H_2_O (50 mL) and brine (50 mL), dried over anhydrous sodium sulfate and concentrated *in vacuo*. The crude product was purified by column chromatography (SiO_2_, 2:98 to 5:95 MeOH:CH_2_Cl_2_) to yield phosphate **5** as a clear oil (1.14 g, 52%). ^1^H NMR (CDCl_3_, 400 MHz) δ 7.38-7.32 (m, 10H, ArH), 5.84 (d, 1H, *J* = 9.2 Hz, NH), 5.66 (dd, 1H, *J* = 6.1, 3.4 Hz, H1), 5.18-5.00 (m, 6H, 2 × PhCH_2_ + H3 + H4), 4.37 (app. ddt, 1H, *J* = 10.7, 9.3, 3.2 Hz, H2), 4.12 (dd, 1H, *J* = 12.5, 3.9 Hz, H6), 4.00 (ddd, 1H, *J* = 9.6, 3.8, 2.2 Hz, H5), 3.91 (dd, 1H, *J* = 12.5, 2.4 Hz, H6’), 2.01 (s, 3H), 2.00 (s, 6H), 1.70 (s, 3H); ^13^C NMR (125 MHz, CDCl_3_) δ 171.2, 170.6, 170.3, 169.2, 129.0, 128.9, 128.9, 128.8, 128.2, 128.2, 128.1, 96.3, 96.2, 70.1, 70.1, 70.0, 70.0, 69.7, 67.4, 61.3, 51.9, 51.8, 22.8, 20.7, 20.7; LRMS (ES) Calcd for C_28_H_34_NNaO_12_P [M+Na]^+^ 630.2, found 630.2.

#### Undecaprenol

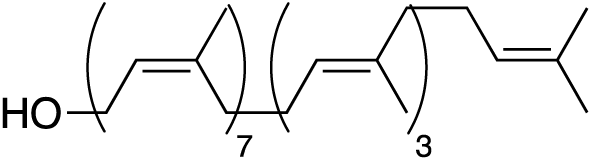

Ground bay leaves (100 g, *Laurus nobilis*) were extracted with a refluxing mixture of 9:1 acetone:*n-*hexanes (1500 mL) by soxhlet extraction for 3 days. The resulting green solution was concentrated *in vacuo* and resuspended (not all solids dissolve) in a mixture of *n*-hexanes (150 mL), EtOH (750 mL) and 15% KOH(aq) (100 mL) and refluxed for 1 h. The resulting mixture was cooled to ambient temperature, followed by addition of H_2_O (500 mL) and Et2O (500 mL). The ether extract was separated, dried over anhydrous Na_2_SO_4_ and concentrated *in vacuo*. The resulting orange solid was purified by column chromatography (SiO_2_ (900 g), 100:0 to 95:5 petrol:EtOAc) using authentic undecaprenol (from American Radiolabelled Chemicals) as a TLC standard, to yield undecaprenol as a yellow oil (950 mg). 1H NMR (CDCl3, 400 MHz) δ 5.46-5.43 (m, 1H, C<u>H</u>CH_2_OH), 5.15-5.08 (m, 10H); 4.09 (dd, 2H, *J* = 7.2, 0.9 Hz, C<u>H</u>_2_OH), 2.09-1.05 (m, 40H), 1.75-1.74 (m, 3H), 1.69-1.67 (m, 21H), 1.61-1.59 (m, 12H); LRMS (ES) Calcd for C_55_H_90_NaO [M+Na]^+^ 789.7, found 789.6.

#### Undecaprenyl phosphate bisammonium salt (Und-P)

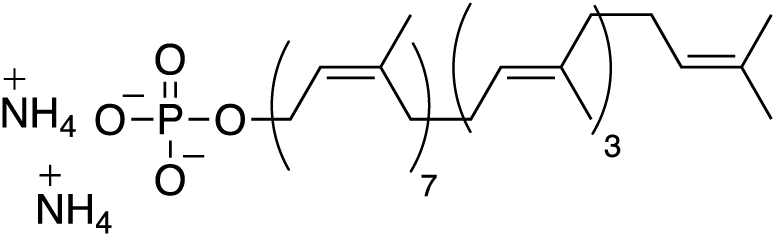

Undecaprenol (710 mg, 1.0 mmol) was dissolved in anhydrous CH_2_Cl_2_ (10 mL) and added rapidly via syringe to a vigorously stirred suspension of 5-ethylthio-1*H*-tetrazole (570 mg, 4.41 mmol) and bis(2-cyanoethyl)-*N*,*N*’-diisopropylphosphoramidite (0.73 mL, 2.85 mmol) in anhydrous CH_2_Cl_2_ (10 mL) under argon at ambient temperature. The reaction mixture became homogeneous within a few min. After 3 h, the mixture was diluted with CH_2_Cl_2_ (80 mL) and washed with saturated sodium bicarbonate (50 mL), water (50mL) and brine (50 mL). The organic solution was dried over anhydrous sodium sulfate and concentrated *in vacuo* to yield the phosphite as a yellow oil. The product was dissolved in THF (20 mL) and cooled to −78 °C. Hydrogen peroxide (30%, 1.9 mL) was added dropwise via syringe to the vigorously stirred solution. After the addition was complete, the ice bath was removed and the mixture was warmed to ambient temperature over 2 h. The reaction mixture was then diluted with ice-cold saturated sodium sulfite (5 mL) and stirred at 0 °C for 5 min. The reaction mixture was then extracted with EtOAc (80 mL) and the organic layer was washed with saturated NaHCO_3_ (50 mL), water (50 mL) and brine (50 mL), dried over anhydrous sodium sulfate and concentrated *in vacuo* to yield the phosphate as a yellow oil. The crude phosphate was suspended in anhydrous MeOH (17 mL) and a 25% NaOMe in MeOH solution (0.7 mL) was added. The resulting suspension was stirred at ambient temperature for 16 h. The reaction mixture was diluted with MeOH (30 mL) and CHCl_3_ (30 mL) and carefully neutralized with DOWEX 50WX8 H^+^ form resin. The resin was removed by filtration and the filtrate concentrated *in vacuo*. The resulting yellow oil was purified by column chromatography (SiO_2_, 90:10:0:0.1 to 65:25:5:0.1 CHCl_3_:MeOH:H_2_O:NH_4_OH) to yield **Und-P** as an off-white foam. Aggregation of this compound prevented acquisition of good NMR spectra. LRMS (ES) Calcd for C_55_H_90_O_4_P [M-H]^−^ 845.6, found 845.6.

#### *N*-Acetyl-3,4,6-tris-*O*-acetyl-1-[*P*’-(3Z,7Z,11Z,15Z,19Z,23Z,27Z,31E,35E,39E,43- undecamethyl-2,6,10,14,18,22,26,30,34,38,42-tetratetracontaundecaenyl) *P,P*’- dihydrogen diphosphate]-α-D-glucosamine diimidazolium salt (6)

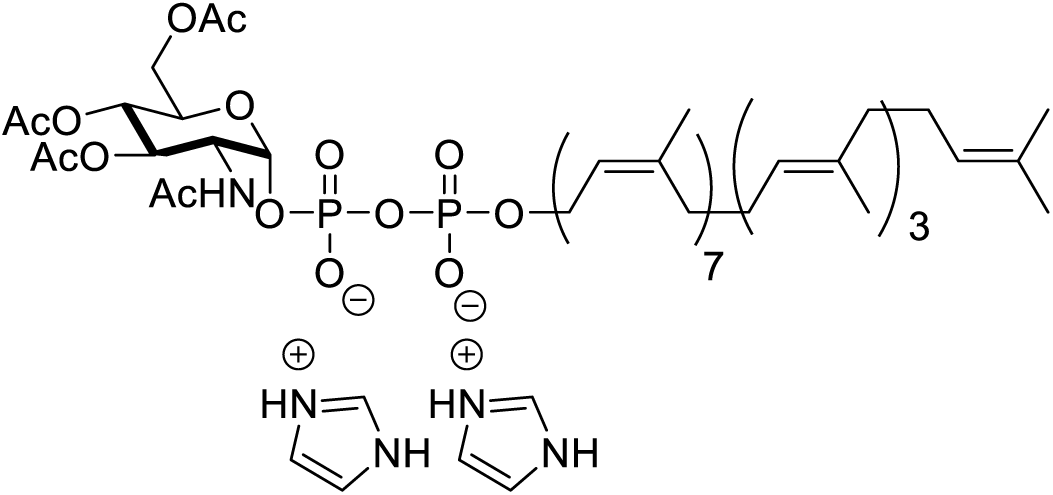

Phosphate **5** (1.00 g, 1.65 mmol) was dissolved in MeOH and the resulting solution degassed with an Ar balloon. A 10% dispersion of palladium on carbon (1.21 g, 1.14 mmol) was added and a H_2_ balloon bubbled through the resulting suspension. The reaction mixture was then stirred under a H_2_ atmosphere for 18 h at ambient temperature. The reaction mixture was then filtered through celite, which was washed with MeOH. Pyridine (1 mL) was added to the filtrate and the resulting solution concentrated *in vacuo* to yield the phosphate dipyridine salt as a white solid (822 mg, 85%). A portion of this product (59 mg, 0.101 mmol) was dissolved in dry THF (1 mL) and DMF (1 mL) under an Ar atmosphere. CDI (76 mg, 0.468 mmol) was added and the resulting solution stirred at RT for 2 h. Reaction monitoring by LRMS (ESI) indicated complete product formation ([M-H]^−^ = 476.1) at this point. Anhydrous MeOH (14 uL, 0.368 mmol) was added and the reaction mixture stirred for a further 45 min. The reaction mixture was concentrated to remove MeOH and THF. To the resulting DMF solution of CDI activated phosphate was added a solution of **Und-P** (89 mg, 0.101 mmol) in THF (2 mL). 5-ethylthio- 1*H*-tetrazole (13 mg, 0.101 mmol) was added to the resulting solution and the reaction mixture stirred for 4 days at ambient temperature. The solution was then concentrated *in vacuo* and purified by column chromatography (SiO_2_, 1:2:4 water:*i*PrOH:EtOAc) to yield the product **6** as a white solid (120 mg, 86%). ^1^H NMR (400 MHz, 1:1 CDCl_3_:CD_3_OD) δ 7.80 (s, Imidazole), 7.10 (s, Imidazole), 5.64 (dd, 1H, *J* = 6.9, 3.1 Hz, H1), 5.45-5.41 (m, 1H, =C<u>H</u>CH_2_OP), 5.32 (dd, 1H, *J* = 10.7, 9.5 Hz, H3), 5.15-5.07 (m, 12H, H4 + NH + 10 × sp^2^ C-H), 4.53 (t, 2H, 7.0 Hz, CH_2_OP), 4.36-4.30 (m, 3H, H2 + H5 + H6), 4.41-4.11 (m, 1H, H6’), 2.10-1.95 (m, 52H, 20 × CH_2_, 4 × Ac), 1.73 (s, 3H), 1.68-1.66 (m, 21H), 1.61 (s, 3H), 1.59 (s, 9H); ^32^P NMR (162 MHz, 1:1 CDCl_3_:CD_3_OD) δ −9.7, −12.7; LRMS (ES) Calcd for C_69_H_110_NO_15_P_2_ [M-H]^−^ 1254.7, found 1254.7.

#### *N*-Acetyl-1-[*P*’-(3Z,7Z,11Z,15Z,19Z,23Z,27Z,31E,35E,39E,43-undecamethyl-2,6,10,14, 18,22,26,30,34,38,42-tetratetracontaundecaenyl)*P,P*’-dihydrogen diphosphate]-α-D-gluc - osamine diammonium salt (7)

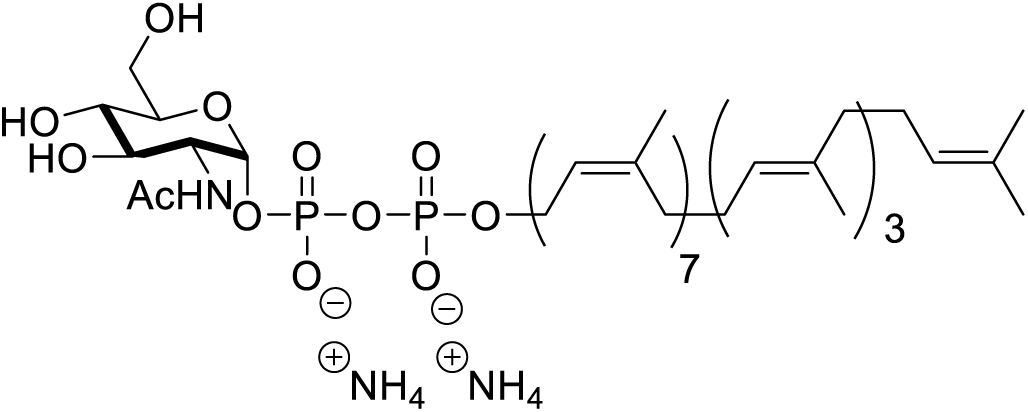

Acetate **6** (120 mg, 0.086 mmol) was suspended in dry MeOH (5 mL) and stirred under an Ar atmosphere. A 25% solution of NaOMe in MeOH (20 uL, 0.093 mmol) was added and the mixture stirred at ambient temperature for 1 h. The reaction mixture was then quenched with excess DOWEX 50WX8 ammonium form and the resin removed by filtration and concentrated *in vacuo* to yield the product **7** as a white solid (100 mg, quant.). LRMS (ES) Calcd for C_69_H_110_NO_15_P_2_ [M-H]^−^ 1254.7, found 1254.7. NMR acquisition proved difficult due to the challenging solubility parameters of this product.

#### SI, Scheme 2. Synthesis of glycosyl donor (9) for lipid II synthesis (the protocol was followed from (Cochrane et al., 2016)

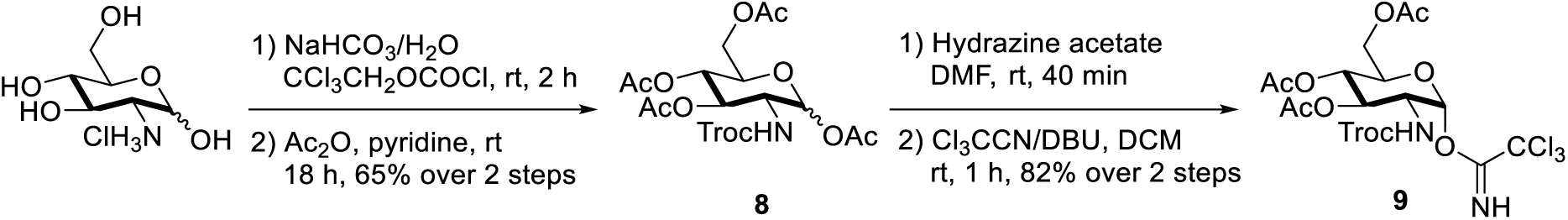

#### 2-Deoxy-2-[[(2,2,2-trichloroethoxy)carbonyl]amino]-3,4,6-triacetyl-1-(2,2,2- trichloroethanimidate)-α-D-glucopyranose (9)

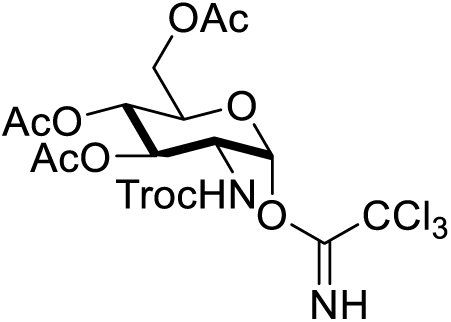

D-Glucosamine (20.00 g, 92.8 mmol) and sodium bicarbonate (15.6 g, 185.6 mmol) were dissolved in water (240 mL) and stirred vigorously for 5 min. 2,2,2-Trichloroethoxycarbonyl chloride (15.3 mL, 111.2 mmol) was added dropwise and the solution stirred at ambient temperature for 2 h, over which time a white precipitate forms. The suspension was filtered, washed with water and coevaporated with toluene (3 × 50 mL). The resulting white powder was dissolved in dry pyridine (200 mL) and acetic anhydride (100 mL) and stirred at ambient temperature for 18 h. The reaction mixture was concentrated *in vacuo* and co-evaporated with toluene (3 × 50 mL). The resulting oily residue was dissolved in CHCl_3_ (200 mL) and washed with 1M HCl (150 mL). The aqueous phase was back extracted with CHCl_3_ (200 mL) and the combined organic extracts washed with brine (100 mL). The organic phase was then dried over anhydrous sodium sulfate and concentrated *in vacuo* to yield 1,3,4,6-tetra-*O*-acetyl-2-troc-D-glucosamine **8** (31.8 g, 65%) as a white solid. **8** (31.8 g, 60.8 mmol) was dissolved in dry DMF (300 mL) and hydrazine acetate (6.72 g, 23.0 mmol) was added. The reaction mixture was stirred at ambient temperature for 40 min, diluted with EtOAc (300 mL) and washed with water (300 mL), saturated sodium bicarbonate (200 mL) and water (200 mL). The combined aqueous phases were then back extracted with EtOAc (300 mL) and the combined organic extracts washed with brine (200 mL), dried over anhydrous sodium sulfate and concentrated *in vacuo*. The resulting red oil was dissolved in CH_2_Cl_2_ (300 mL) and trichloroacetonitrile (61 mL, 608 mmol). 1,8-Diazabicycloundec-7-ene (1.81 mL, 12.16 mmol) was added and the reaction mixture stirred at ambient temperature for 90 min and concentrated *in vacuo*. The crude reaction mixture was purified by flash column chromatography (silica, 2:1 hexanes:EtOAc + 0.1% triethylamine) to yield the product **9** as a white foam (31 g, 82%). [α]_D_^25^ = 76.1 (*c* = 1.1 g/100mL, CH_2_Cl_2_); ^1^H NMR (CDCl_3_, 400 MHz) δ 8.80 (s, 1H, acetimidate-NH), 6.43 (d, *J* = 3.6 Hz, 1H, H1), 5.35 (dd, *J* = 10.8, 9.5 Hz, H3), 5.25 (t, *J* = 9.9 Hz, H4), 5.19 (d, *J* = 9.3 Hz, Troc-NH), 4.75 – 4.67 (m, 2H, Troc-CH_2_), 4.29 (m, 2H, H_2_ + H6), 4.16 – 4.09 (m, 2H, H5 + H6), 2.07 (m, 9H, 3 × OCH_3_); ^13^C NMR (CDCl_3_, 125 MHz) δ 171.0, 170.4, 169.1, 160.3, 154.0, 95.1, 94.4, 90.5, 74.6, 70.2, 67.3, 61.3, 53.8, 20.6, 20.5.

#### SI, Scheme 3. Synthesis of alanine ester (10) for lipid II synthesis (the protocol was followed from (Cochrane et al., 2016)

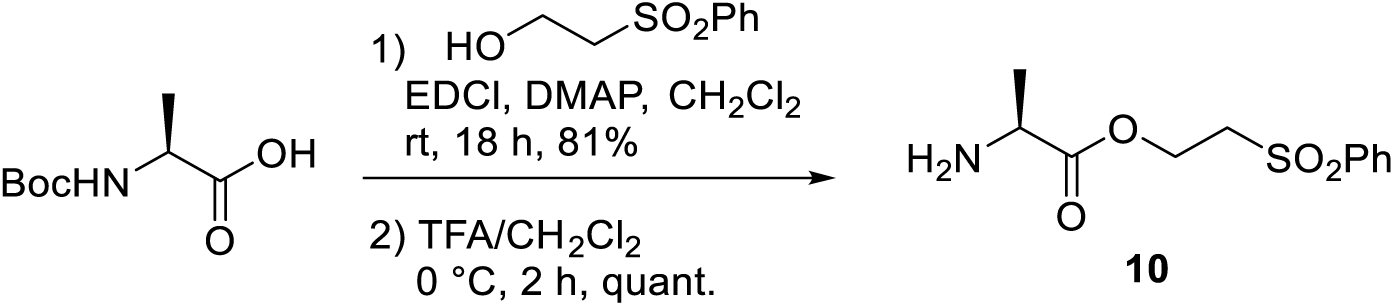

L-Alanine-2-(phenylsulfonyl)ethyl ester (10)

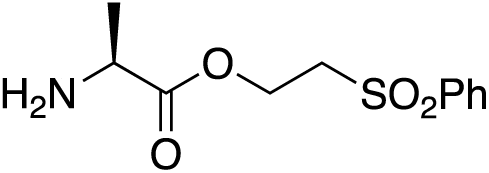

Boc-L-alanine (5.08 g, 26.8 mmol), 2-phenylsulfonylethanol (5.00 g, 26.8 mmol), 1-(3-Dimethylaminopropyl)-3-ethylcarbodiimide hydrochloride (5.20 g, 26.8 mmol) and 4- dimethylaminopyridine (0.33 g, 2.68 mmol) were dissolved in dry CH_2_Cl_2_ (150 mL) and stirred at ambient temperature under an argon atmosphere for 18 h. The reaction mixture was then washed with 0.5 M HCl (100 mL) and saturated NaHCO_3_ (100 mL). Each aqueous phase was back extracted with CH_2_Cl_2_ (50 mL) and the combined organic extracts washed with brine (100 mL). The organic extracts were then dried over anhydrous Na_2_SO_4_ and concentrated *in vacuo*. The resulting orange oil was dissolved in CHCl_3_ (30 mL) and passed through a silica plug, with CHCl_3_ washing until no further product fractions eluted. The solution was concentrated *in vacuo*, redissolved in CH_2_Cl_2_ (40 mL) and cooled to 0 °C. Trifluoroacetic acid (40 mL) was added slowly and the solution warmed to ambient temperature and stirred for 3 hours. The resulting orange solution was then concentrated *in vacuo*, azeotroped with toluene (2 × 10 mL) and purified by column chromatography (SiO_2_, 1:9 MeOH:CH_2_Cl_2_) to yield the product **10** as a colourless oil (quant.). ^1^H NMR (CDCl_3_, 400 MHz) δ 7.89 – 7.86 (m, 2H, *o*-ArH), 7.69 – 7.65 (m, 1H, *p*-ArH), 7.59 – 7.55 (m, 2H, *m*-ArH), 4.53 (t, 2H, *J* = 5.8 Hz, O-CH_2_), 3.91 (app. q, 1H, *J* = 7.2 Hz, Ha); 3.49 – 3.46 (m, 2H, S-CH_2_), 1.30 (d, *J* = 7.3 Hz, 3H, Hβ); ^13^C NMR (CDCl_3_, 125 MHz) δ 175.7, 139.5, 134.2, 129.6, 128.3, 58.1, 55.2, 49.9, 20.2.

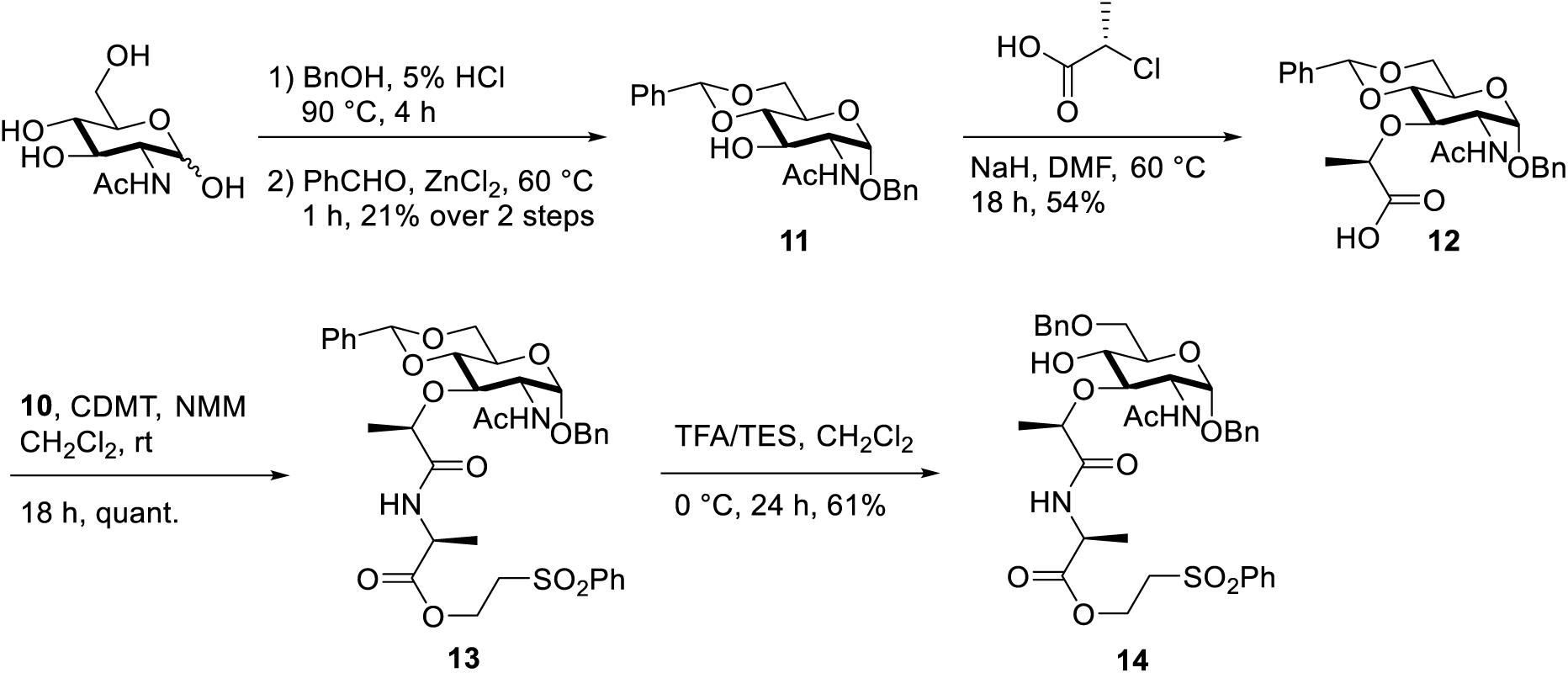

#### SI, Scheme 4. Synthesis of glycosyl acceptor (14) for lipid II synthesis (the protocol was followed from (Cochrane et al., 2016)

Phenylmethyl-2-(acetylamino)-2-deoxy-4,6-*O*-(phenylmethylene)-α-D-glucopyranoside (11)

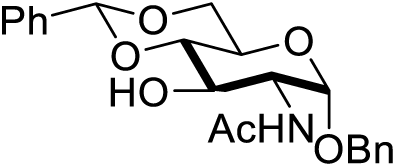

*N*-Acetyl-D-glucosamine (30 g, 136 mmol) was suspended in benzyl alcohol (300 mL) and 37% HCl(aq) (7.5 mL) and heated at 95 °C for 4 h. During this time the white suspension became a purple/brown solution. The reaction mixture was concentrated to ~50 mL and precipitated by adding to ice-cold Et_2_O (600 mL) and stirring for 1 h. The off-white precipitate was filtered, was with Et_2_O (3 × 100 mL) and dried under high vacuum overnight. The resulting white solid (38 g) was suspended in benzaldehyde (140 mL) and finely ground anhydrous ZnCl_2_ (38 g) was added. The resulting suspension was stirred at 60 °C until for 1 h, with most solids dissolving after 5 min. The mixture was then poured into a stirring ice-water mixture (400 mL) and the resulting precipitate collected by filtration. The pinkish solid was washed with H_2_O (2 × 200 mL), ice-cold EtOH (100 mL) and Et_2_O (3 × 200 mL). The resulting off-white solid (20 g) was dissolved in boiling pyridine (100 mL) and boiling H_2_O was added until to solution turned cloudy. The solution was then cooled to ambient temperature, followed by ice-bath for 1 h. The resulting precipitate was collected by filtration, washed with H_2_O (2 × 100 mL), azeotroped with toluene (500 mL) and dried by high vacuum overnight to yield the product **11** as a white fluffy solid (11.4 g, 21% over 2 steps). [α]_D_^25^ = +124 (*c* = 1.1 g/100mL, pyridine); ^1^H NMR (DMSO-*d*_*6*_, 400 MHz) δ 8.00 (d, 1H, *J* = 8.3 Hz, N<u>H</u>Ac), 7.46-7.28 (m, 10H, Ar-H), 5.62 (s, 1H, PhC<u>H</u>), 5.19 (d, 1H, *J* = 5.8 Hz, 3-OH), 4.80 (d, 1H, *J* = 3.5 Hz, H1), 4.70 (d, 1H, *J* = 12.6 Hz, PhC<u>H</u>H), 4.49 (d, 1H, *J* = 12.6 Hz, PhC<u>H</u>H), 4.16-4.14 (m, 1H, H6), 3.88-3.82 (m, 1H, H6’), 3.77-3.67 (m, 3H, H2 + H3 + H5), 3.53-3.49 (m, 1H, H4), 1.85 (s, 3H, NH<u>Ac</u>).

#### *N*-Acetyl-1-*O*-(phenylmethyl)-4,6-*O*-(phenylmethylene)-α-D-muramic acid (12)

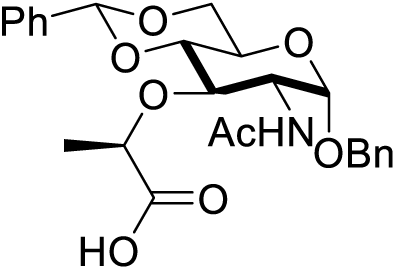

Glycol **11** (15.0 g, 37.6 mmol) was suspended in dry dioxane (700 mL) under an argon atmosphere and stirred at 60 °C. A 60% dispersion of NaH in mineral oil (3.0 g, 75 mmol) was added and the mixture stirred for 5 min. (2*S*)-chloropropionic acid (13 mL, 75 mmol) was added and the mixture stirred at 60 °C for a further 10 min. Another portion of a 60% dispersion of NaH in mineral oil (7.5 g g, 188 mmol) was added and the mixture stirred for at 60 °C for 16 h. The reaction mixture was cooled to ambient temperature and H_2_O (400 mL) added slowly with stirring. The dioxane was then removed from the resulting yellow solution by concentration with rotary evaporator. The resulting yellow solution was extracted with CHCl_3_ (200 mL), cooled on ice, acidified to pH 1 with 6M HCl and extracted with CHCl_3_ (3 × 200 mL). The combined organic extracts were washed with H_2_O (200 mL), dried over anhydrous Na_2_SO_4_ and concentrated *in vacuo*. The resulting yellow solid was recrystallized from boiling CHCl_3_ (~250 mL) to yield the product **12** as a white crystalline solid (11.0 g, 54%). ^1^H NMR (DMSO-*d*_*6*_, 400 MHz) δ 7.97 (d, 1H, *J* = 8.3 Hz, N<u>H</u>Ac), 7.44-7.28 (m, 10H, Ar-H), 5.70 (s, 1H, PhC<u>H</u>), 5.04 (d, 1H, *J* = 3.5 Hz, H1), 4.70 (d, 1H, *J* = 12.4 Hz, PhC<u>H</u>H), 4.49 (d, 1H, *J* = 12.4 Hz, PhC<u>H</u>H), 4.29 (q, 1H, *J* = 6.9 Hz, Ala1-Hα), 4.17-4.13 (m, 1H, H6), 3.83-3.68 (m, 4H, H6’ + H_2_ + H3 + H5), 1.85 (s, 3H, NH<u>Ac</u>), 1.28 (d, *J* = 7.2 Hz, 3H, Ala1-Hβ).

#### Phenylmethyl-2-(acetylamino)-2-deoxy-3-*O*-[(1*R*)-1-methyl-2-[[(1*S*)-1-methyl-2-oxo-2-[2-(phenylsulfonyl)ethoxy]ethyl]amino]-2-oxoethyl]-4,6-*O*-[(*R*)-phenylmethylene]-α-D-glucopyranoside (13)

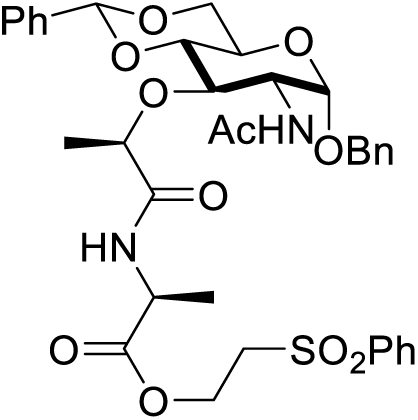

Acid **12** (7.85 g, 16.7 mmol) was suspended in dry CH_2_Cl_2_ (120 mL) and cooled to 0 °C. NMM (1.83 mL, 16.7 mmol) and CDMT (3.51 g, 20.0 mmol) were added and the resulting cloudy suspension stirred at 0 °C for 45 min. A solution of amine **10** (20.0 mmol) and NMM (1.83 mL, 16.7 mmol) in CH_2_Cl_2_ (120 mL) were then added and the resulting solution stirred at ambient temperature overnight. The reaction mixture was then filtered and the filtrate was washed with 1M HCl (100 mL) and brine (100 mL), dried over anhydrous Na_2_SO_4_ and concentrated *in vacuo*. The resulting solid was azeotroped with toluene (2 × 50 mL) and CHCl_3_ (2 × 50 mL) to yield the product **13** as a white solid (11.8 g, 99%). ^1^H NMR (CDCl_3_, 400 MHz) δ 7.91-7.90 (m, 2H, ArH), 7.68-7.64 (m, 1H, ArH), 7.58-7.54 (m, 2H, ArH), 7.50-7.46 (m, 2H, ArH), 7.38-7.27 (m, 8H, ArH), 6.93 (d, *J* = 7.1 Hz, 1H, Ala1-NH), 6.22 (d, *J* = 8.9 Hz, 1H, AcNH), 5.57 (s, 1H, O_2_C<u>H</u>), 4.96 (d, *J* = 3.8 Hz, 1H, H1), 4.71 (d, *J* = 11.8 Hz, 1H, PhC<u>H</u>H), 4.51 – 4.39 (m, 3H, PhCHH + OC<u>H</u>_2_), 4.32-4.23 (m, 2H, H_2_ + H6), 4.17 (t, *J* = 7.2 Hz, 1H, Ala1-Hα), 4.07 (q, *J* = 6.7 Hz, 1H, OC<u>H</u>), 3.91-3.85 (m, 1H, H5), 3.80-3.64 (m, 3H, H6’ + H3 + H4), 3.48- 3.53 (m, 2H, SC<u>H</u>_2_), 1.95 (s, 3H, Ac), 1.38 (d, *J* = 6.8 Hz, 3H, MurNAc-CH_3_), 1.30 (d, *J* = 7.2 Hz, 3H, Ala1-Hβ); ^13^C NMR (CDCl_3_, 125 MHz) δ 173.2, 170.9, 170.65, 137.20, 136.8, 134.2, 129.6, 128.8, 128.4, 128.3, 128.2, 126.1, 101.60, 97.6, 81.7, 78.5, 78.2, 70.3, 69.0, 63.3, 58.2, 55.0, 53.2, 48.10, 23.5, 19.5, 17.3.

#### Phenylmethyl-2-(acetylamino)-2-deoxy-3-*O*-[(1*R*)-1-methyl-2-[[(1*S*)-1-methyl-2-oxo-2-[2-(phenylsulfonyl)ethoxy]ethyl]amino]-2-oxoethyl]-6-*O*-(phenylmethyl)-α-D-glucopyranoside (14)

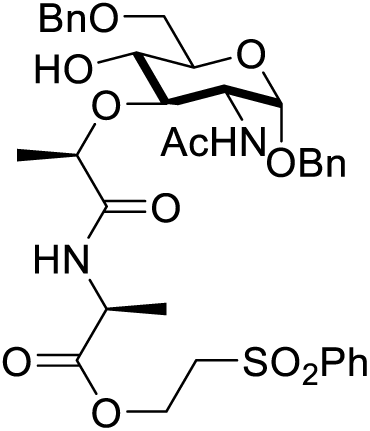

Benzylidene **13** (6.00 g, 8.44 mmol) was dissolved in dry CH_2_Cl_2_ (75 mL) and cooled to 0 °C. Triethylsilane (6.72 mL, 42.2 mmol) was added and the solution stirred for 5 min. TFA (3.23 mL, 42.2 mmol) was then added over 5 min and the reaction mixture stirred for 6 h at 0 °C. Another portion of TFA (1.94 mL, 25.3 mmol) was added at once and the reaction mixture stirred for a further 18 h at 0 °C. The reaction mixture was then diluted with CH_2_Cl_2_ (100 mL) and washed with saturated aqueous NaHCO_3_ (100 mL). The aqueous phase was back extracted with CH_2_Cl_2_ (2 × 100 mL) and the combined organic extracts washed with brine (100 mL), dried over anhydrous Na_2_SO_4_ and concentrated *in vacuo*. The resulting crude product was purified by column chromatography (SiO_2_, 8:2 to 10:0 EtOAc:petrol) to yield glycosyl acceptor **14** was a white foam (3.65 g, 61%). ^1^H NMR (CDCl_3_, 400 MHz) δ 7.94-7.85 (m, 2H, ArH), 7.69-7.63 (m, 1H, ArH), 7.57 (t, *J* = 7.7 Hz, 2H, ArH), 7.41-7.22 (m, 9H, ArH), 6.92 (d, *J* = 7.2 Hz, 1H, Ala1-NH), 6.10 (d, *J* = 9.0 Hz, 1H, MurNAc-NH), 4.92 (d, *J* = 3.6 Hz, 1H, H1), 4.71 (d, *J* = 11.8 Hz, 1H, OC*H*H), 4.65-4.54 (m, 2H, OC*H*H + OCH*H*), 4.49-4.34 (m, 3H, OCH*H* + OCH_2_), 4.29-4.17 (m, 2H, H_2_ + Ala1-Hα), 4.13 (t, *J* = 6.7 Hz, 1H, MurNAc-OCH), 3.80 (dd, *J* = 9.5, 4.6 Hz, 1H, H5), 3.74 (dd, *J* = 10.3, 4.5 Hz, 1H, H6), 3.72-3.65 (m, 2H, H3 + H4), 3.53 (dd, *J* = 10.5, 8.7 Hz, 1H, H6), 3.39 (ddd, *J* = 9.2, 6.6, 5.5 Hz, 2H, S-CH_2_), 3.01 (s, 1H, OH), 1.90 (s, 3H, NHAc), 1.40 (d, *J* = 6.7 Hz, 3H, MurNAc-CH_3_), 1.30 (d, *J* = 7.2 Hz, 3H, Ala1-Hβ); ^13^C NMR (CDCl_3_, 125 MHz) δ 173.1, 171.91, 170.4, 139.2, 137.9, 137.1, 134.2, 129.6, 128.7, 128.6, 128.3, 128.3, 128.2, 127.9, 127.8, 97.2, 80.6, 77.9, 73.8, 71.7, 70.5, 70.3, 69.9, 58.1, 55.0, 52.6, 48.0, 23.5, 19.1, 17.2.

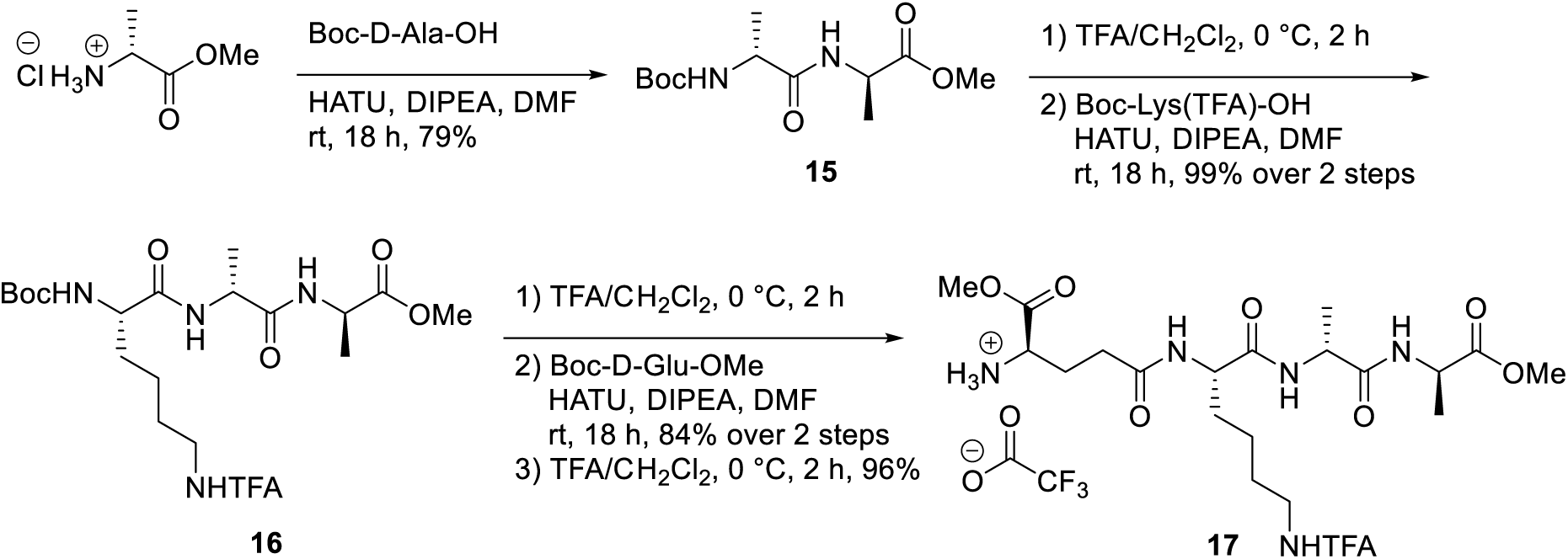

#### SI, Scheme 5. Synthesis of tetrapeptide (17) for lipid II synthesis (the protocol was followed from (Cochrane et al., 2016)

**Boc-D-Ala-D-Ala-OMe (15)**

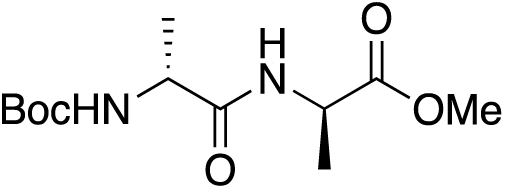

H-D-Ala-OMe.HCl (5.00 g, 35.8 mmol), Boc-D-Ala-OH (6.78 g, 35.8 mmol) and HATU (13.60 g, 35.8 mmol) were dissolved in dry DMF (175 mL) and cooled to 0 °C. DIPEA (6.25 mL, 107.4 mmol) was added and the reaction stirred at ambient temperature for 18 h. The solution was then concentrated *in vacuo*, re-dissolved in EtOAc (200 mL) and washed with 0.5 M HCl (100 mL), saturated sodium bicarbonate (100 mL) and brine (100 mL). The organic phase was then dried over anhydrous sodium sulfate and concentrated *in vacuo*. The crude material was dissolved in CHCl_3_ (100 mL), filtered through celite and concentrated *in vacuo* to yield Boc-dipeptide **15** as a white foam (7.75 g, 79%). ^1^H NMR (CDCl_3_, 400 MHz) δ 6.67 (m, 1H, D- Ala5NH), 5.03 (m, 1H, D-Ala4NH), 4.56 (app. pentet, *J* = 7.2 Hz, 1H, D-Ala5Hα), 4.17 (m, 1H, D-Ala4Hα), 3.74 (s, 3H, D-Ala5-OMe), 1.44 (m, 9H, Boc), 1.39 (d, *J* = 7.1 Hz, 3H, D-Ala5Hβ), 1.35 (d, *J* = 7.1 Hz, 3H, _D_-Ala4Hβ). ^13^C NMR (CDCl_3_, 125 MHz) δ 173.3, 172.3, 52.6, 48.1, 28.4, 18.5, 18.4.

#### Boc-Lys-D-Ala-D-Ala-OMe (16)

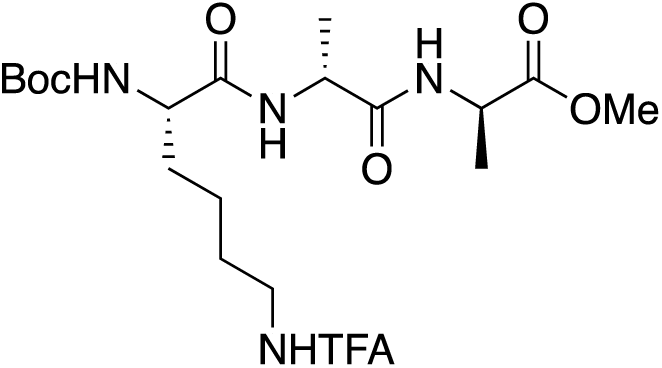

Boc-dipeptide **15** (3.00 g, 10.9 mmol) was dissolved in CH_2_Cl_2_ (30 mL) and cooled to 0 °C. TFA (30 mL) was added and the resulting solution stirred at ambient temperature for 3 h. The reaction mixture was then concentrated *in vacuo*, azeotroped with toluene and dried under high vacuum for 1 h. During this time, in a separate flask, Boc-Lys(TFA)-OH (3.73 g, 10.9 mmol) and HATU (4.14 g, 10.9 mmol) were dissolved in dry DMF (50 mL) and cooled to 0 °C. DIPEA (5.70 mL, 32.7 mmol) was added and the resulting yellow solution stirred at 0 °C for 15 min. The deprotected dipeptide (10.9 mmol) was dissolved in DMF (10 mL) and added to the activated acid solution. The resulting reaction mixture was stirred at ambient temperature overnight and concentrated *in vacuo*. The resulting oil was redissolved in EtOAc (100 mL), washed with 1M HCl (100 mL), saturated aqueous NaHCO_3_ (100 mL) and brine (100 mL), dried over anhydrous Na_2_SO_4_ and concentrated *in vacuo* to yield the product **16** as a white foam (5.4 g, 99%). ^1^H NMR (DMSO-*d*_*6*_, 400 MHz) δ 9.34 (t, 1H, *J* = 5.5 Hz, Lys3-NHTFA), 8.20 (d, 1H, *J* = 7.1 Hz, D-Ala4-NH), 8.01 (d, 1H, *J* = 7.9 Hz, D-Ala5-NH), 6.93 (d, 1H, *J* = 7.5 Hz, Lys3-NH), 4.35-4.23 (m, 2H, Lys3-Hα + D-Ala4-Hα), 3.89-3.84 (m, 1H, D-Ala5-Hα), 3.60 (s, 3H, OMe), 3.15 (app. q, 2H, *J* = 6.5 Hz, Lys3-Hε), 1.60-1.40 (m, 4H, Lys3-Hβ + Lys3- Hδ), 1.36 (s, 9H, *t*Bu), 1.31-1.17 (m, 8H, Lys3-Hγ + D-Ala4-Hβ + D-Ala5-Hβ); ^13^C NMR (CDCl_3_, 125 MHz) δ 172.8, 172.0, 171.7, 78.2 51.9, 47.6, 47.5, 28.1, 27.9, 22.7, 18.2, 16.8; LRMS (ESI) Calcd for C_20_H_33_F_3_N_4_NaO_7_ [M+Na]+ 521.2, found 521.2.

#### H-γ-D-Glu(α-OMe)-Lys(TFA)-D-Ala-D-Ala-OMe trifluoroacetate salt (17)

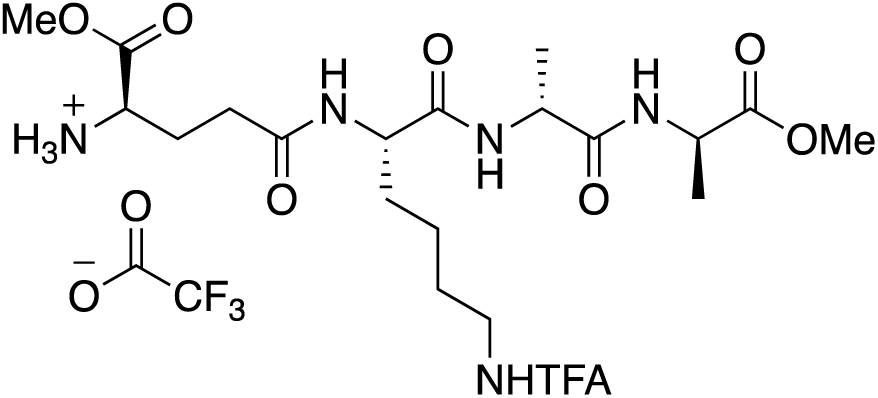

Boc-Tripeptide **16** (5.4 g, 10.8 mmol) was dissolved in CH_2_Cl_2_ (30 mL) and cooled to 0 °C. TFA (30 mL) was added and the resulting solution stirred at ambient temperature for 2 h. The reaction mixture was then concentrated *in vacuo*, azeotroped with toluene and dried under high vacuum for 1 h. During this time, in a separate flask, Boc-γ-D-Glu(α-OMe)-OH (2.92 g, 10.8 mmol) and HATU (4.10 g, 10.8 mmol) were dissolved in dry DMF (50 mL) and cooled to 0 °C. DIPEA (5.70 mL, 32.4 mmol) was added and the resulting yellow solution stirred at 0 °C for 15 min. The deprotected tripeptide (10.8 mmol) was dissolved in DMF (15 mL) and added to the activated acid solution. The resulting reaction mixture was stirred at ambient temperature overnight and concentrated *in vacuo*. The resulting oil was redissolved in EtOAc (100 mL) and DMF (5 mL), washed with 1M HCl (100 mL), saturated aqueous NaHCO_3_ (100 mL) and brine (100 mL), dried over anhydrous Na_2_SO_4_ and concentrated *in vacuo* to yield the Boc-tetrapeptide as a white powder (5.85 g, 84%). A portion of the crude Boc-tetrapeptide (2.00 g, 3.11 mmol) was suspended in CH_2_Cl_2_ (10 mL). TFA (10 mL) was added, at which point all solids dissolved. The reaction mixture was stirred for 2 h at ambient temperature, concentrated *in vacuo* and azeotroped with MeOH (2 × 8 mL), CH_2_Cl_2_ (2 × 10 mL) to yield the product **17** was an off-white foam (1.91 g, 96%). ^1^H NMR (DMSO-*d*_*6*_, 400 MHz) δ 9.42 (t, 1H, *J* = 5.5 Hz, Lys3-NHTFA), 8.41 (br. s, 3H, D-Glu2-H_3_N^+^), 8.27 (d, 1H, *J* = 7.0 Hz, _D_-Ala4-NH), 8.22 (d, 1H, *J* = 7.9 Hz, _D_-Ala5-NH), 8.13 (d, 1H, *J* = 7.8 Hz, Lys3-NH), 4.34-4.20 (m, 4H, _D_-Glu2- Hα + Lys3-Hα + _D_-Ala4-Hα + _D_-Ala5-Hα), 3.73 (s, 3H, _D_-Glu2-OMe), 3.60 (s, 3H, _D_-Ala5-OMe), 3.15 (app. q, 2H, *J* = 6.4 Hz, Lys3-Hε), 2.38-2.24 (m, 2H, _D_-Glu-Hγ), 2.04-1.92 (m, 2H _D_-Glu-Hβ), 1.64-1.43 (m, 4H, Lys3-Hβ + Lys3-Hδ), 1.30-1.18 (m, 8H, Lys3-Hγ + Ala4-Hβ + Ala5-Hβ); ^13^C NMR (DMSO-*d*_*6*_, 400 MHz) δ 172.9, 172.1, 171.2, 170.7, 169.8, 52.8, 52.5, 51.9, 51.6, 47.6, 31.7, 30.2, 28.0, 25.9, 22.5, 18.2, 16.8; LRMS (ESI) Calcd for C_20_H_33_F_3_N_4_NaO_7_ [M+Na]^+^ 521.2, found 521.2.

#### SI, Scheme 6. Synthesis of lipid II (the protocol was followed from (Cochrane et al., 2016; VanNieuwenhze et al., 2002).

**Phenylmethyl-2-(acetylamino)-2-deoxy-3-*O*-[(1*R*)-1-methyl-2-[[(1*S*)-1-methyl-2-oxo-2-[2-(phenylsulfonyl)ethoxy]ethyl]amino]-2-oxoethyl]-6-*O*-(phenylmethyl)-4-*O*-[3,4,6-tri-*O*-acetyl-2-deoxy-2-[[(2,2,2-trichloroethoxy)carbonyl]amino]-β-D-glucopyranosyl]-α-D-glucopyranoside (18)**

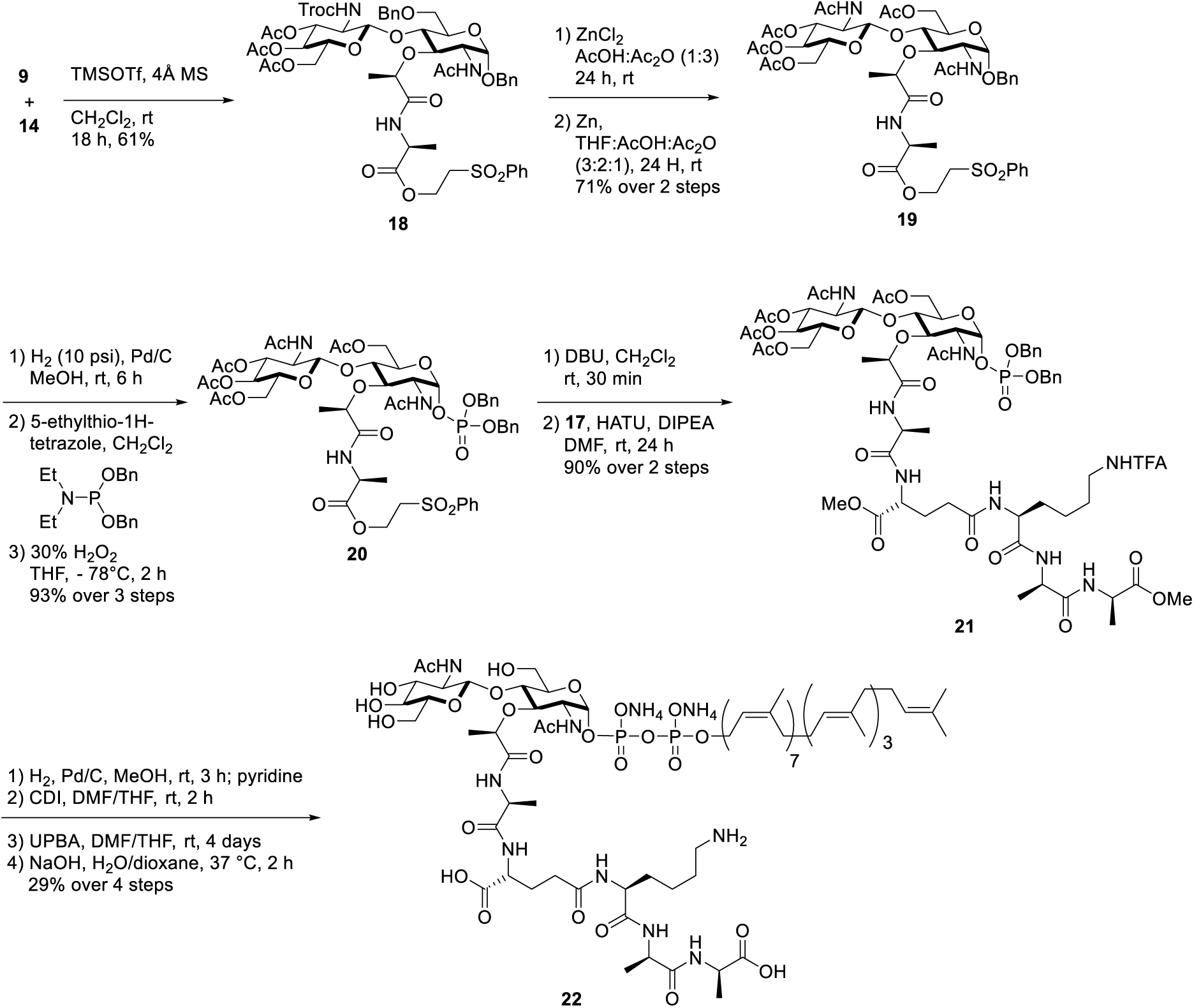

4 Å molecular sieves (25 g) in a round-bottomed flask were heated under vacuum with a heat gun for approximately 5 min and left to cool at ambient temperature. The flask was depressurized with argon and directly used in the reaction. A solution of glycol **14** (3.2g, 4.49 mmol) in alcohol-free CH_2_Cl_2_ (50 mL) was added to this flask under argon and the suspension gently stirred. TMSOTf (0.81 mL, 4.49 mmol) was added, followed by a solution of acetimidate **9** (8.42 g, 13.47 mmol) in dry CH_2_Cl_2_ (50 mL). The resulting suspension was stirred at ambient temperature overnight. Another portion of acetimidate **9** (5.61 g, 8.98 mmol) and TMSOTf (0.4 mL, 2.25 mmol) were added and the reaction stirred for a further 24 h. The reaction mixture was decanted and the solution diluted with CH_2_Cl_2_ (100 mL). The organic solution was washed with saturated sodium bicarbonate (100 mL) and brine (100 mL) and dried over anhydrous sodium sulfate. The solution was concentrated *in vacuo* and purified by column chromatography (SiO_2_, gradient: 1:1 EtOAc:petrol to EtOAc) to yield the product **18** as a white foam (3.2 g, 61%). ^1^H NMR (CDCl_3_, 400 MHz) δ 7.91-7.90 (m, 2H, ArH), 7.67-7.63 (m, 1H, ArH), 7.58-7.43 (m, 6H, ArH), 7.35-7.25 (m, 6H, ArH), 6.85 (d, *J* = 7.5 Hz, 1H, Ala1NH), 6.53 (d, *J* = 7.5 Hz, 1H, MurNAc-NH), 5.08 (d, *J* = 3.6 Hz, 1H, MurNAc-H1), 4.97 (t, *J* = 9.6 Hz, 1H, GlcNAc-H4), 4.87 (d, *J* = 12.0 Hz, 1H, MurNAc-1-CHHPh), 4.79-4.73 (m, 2H, GlcNAc-H3 + Troc-CHH), 4.62-4.56 (m, 2H, Troc-CHH + MurNAc-6-CHHPh), 4.47-4.32 (m, 4H, OCHH + MurNAc-6-CHHPh + OCHH + MurNAc-1-CHHPh), 4.25 – 4.07 (m, 5H, MurNAc-H_2_ + MurNAc-CHO + GlcNAc-H1 + GlcNAc-H6 + Ala1Hα), 3.98 (dd, *J* = 12.3, 1.9 Hz, 1H, GlcNAc-H6), 3.91 (t, *J* = 9.5 Hz, 1H, MurNAc-H3), 3.69-3.51 (m, 3H, MurNAc-H6 + MurNAc-H4 + MurNAc-H5), 3.44-3.38 (m, 4H, CH_2_S + GlcNAc-H_2_ + GlcNAc-H5), 2.05 – 1.98 (m, 9H, 3 × Ac), 1.90 (s, 3H, Ac), 1.34 (d, *J* = 6.7 Hz, 3H, Ala1Hβ), 1.23 (dd, *J* = 9.7, 7.2 Hz, 3H, MurNAc-CH_3_). ^13^C NMR (CDCl_3_, 125 MHz) δ 173.4, 171.9, 170.7, 170.5, 170.4, 169.5, 154.2, 139.3, 137.4, 137.2, 134.1, 129.5, 129.2, 128.6, 128.2, 128.2, 100.1, 97.2, 95.7, 77.7, 75.7, 74.6, 73.8, 72.2, 71.3, 70.5, 70.4, 68.4, 67.2, 61.5, 58.2, 56.3, 55.0, 53.7, 47.8, 23.3, 20.7, 18.4, 17.6; LRMS (ESI) Calcd for C_51_H_62_Cl_3_N_3_NaO_20_S [M+Na]^+^ 1196.2, found 1196.2.

#### Phenylmethyl 2-(acetylamino)-2-deoxy-3-*O*-[(1*R*)-1-methyl-2-[[(1*S*)-1-methyl-2-oxo-2-[2-(phenylsulfonyl)ethoxy]ethyl]amino]-2-oxoethyl]-4-*O*-[3,4,6-tri-*O*-acetyl-2-(acetylamino)-2-deoxy-β-D-glucopyranosyl]-6-*O*-acetyl-α-D-glucopyranoside (19)

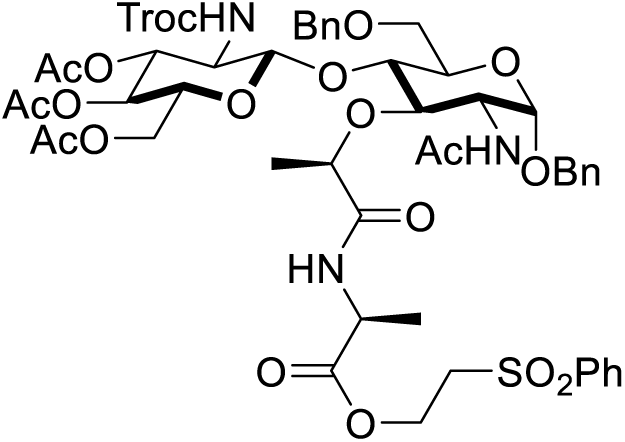

The Troc-disaccharide **18** (3.0 g, 2.55 mmol) was dissolved in Ac_2_O (12 mL) and AcOH (6 mL) and to this solution was added a solution of anhydrous ZnCl_2_ (3.48 g, 25.5 mmol) in Ac_2_O (5.5 mL) and AcOH (2.5 mL). The reaction mixture was stirred for 24 h at ambient temperature, at which point zinc dust (6.67 g, 102.0 mmol) and a mixture of THF (20 mL), Ac_2_O (13 mL) and AcOH (7 mL) were added. The reaction was stirred for a further 24 h at ambient temperature and filtered through celite, washed with EtOAc (300 mL), and concentrated *in vacuo*. The resulting residue was co-evaporated with toluene (2 × 50 mL) and re-dissolved in EtOAc (200 mL). The organic layer was washed with saturated sodium bicarbonate (2 × 10 mL), which was then back-extracted with EtOAc (100 mL). The combined organics were washed with water (100 mL) and brine (100 mL) and dried over anhydrous Na_2_SO_4_ and concentrated *in vacuo*. The crude product was purified by column chromotography (SiO_2_, EtOAc) to yield the product **19** as a white foam (1.8 g, 71%). ^1^H NMR (CDCl_3_, 400 MHz) δ 7.90-7.88 (m, 2H, *ortho*-ArH), 7.69-7.65 (m, 1H, *para*-ArH), 7.59-7.55 (m, 2H, *meta*-ArH), 7.34-7.25 (m, 5H, ArH), 7.14 (d, *J* = 7.6 Hz, 1H, MurNAc-NH), 6.81 (d, *J* = 6.9 Hz, 1H, Ala1NH), 6.04 (d, *J* = 9.5 Hz, 1H, GlcNAc-NH), 5.12-5.10 (m, 2H, MurNAc-H1 + GlcNAc- H3), 4.63 (d, *J* = 12.1 Hz, 1H, MurNAc-1-CHHPh), 4.49 (d, *J* = 12.1 Hz, 1H, MurNAc-1-CHHPh), 4.40-4.22 (m, 6H, OCH_2_ + MurNAc-CHO + GlcNAc-H1 + GlcNAc-H6 + MurNAc-H6), 4.16-4.08 (m, 2H, MurNAc-H61H + GlcNAc-H6), 4.06-3.98 (m, 3H, GlcNAc-H2 + MurNAc-H2 + MurNAc-H3), 3.78 (d, *J* = 5.2 Hz, MurNAc-H5), 3.62-3.50 (m, 2H, GlcNAc-H5 + MurNAc-H4), 3.40-3.30 (m, 2H, CH_2_S), 2.14 (s, 3H), 2.02 (s, 3H), 2.01 (s, 3H), 2.00 (s, 3H), 1.95 (m, 3H), 1.92 (s, 3H), 1.37 (d, *J* = 6.7 Hz, 3H, MurNAc-CH_3_), 1.29 (d, *J* = 7.2 Hz, 3H, Ala1Hβ); ^13^C NMR (CDCl_3_, 400 MHz) δ 173.8, 172.0, 171.3, 171.0, 170.9, 170.7, 170.7, 169.4, 139.2, 137.4, 134.2, 129.5, 128.6, 128.2, 128.1, 128.0, 100.4, 97.0, 76.1, 75.7, 72.6, 71.9, 70.4, 69.6, 68.2, 62.4, 61.7, 60.5, 58.1, 55.0, 54.7, 53.7, 47.9, 23.3, 23.3, 21.1, 20.7, 20.7, 20.7, 18.4, 17.4. LRMS (ES) Calcd for C_45_H_59_N_3_NaO_20_S [M+Na]^+^ 1016.3, found 1016.3.

#### ***N***-[*N*-Acetyl-6-*O*-acetyl-1-*O*-[bis(phenylmethoxy)phosphinyl]-4-*O*-[3,4,6-tri-*O*-acetyl-2-(acetylamino)-2-deoxy-β-D-glucopyranosyl]-α-muramoyl]-L-alanine-2-(phenylsulfonyl)ethyl ester (20)

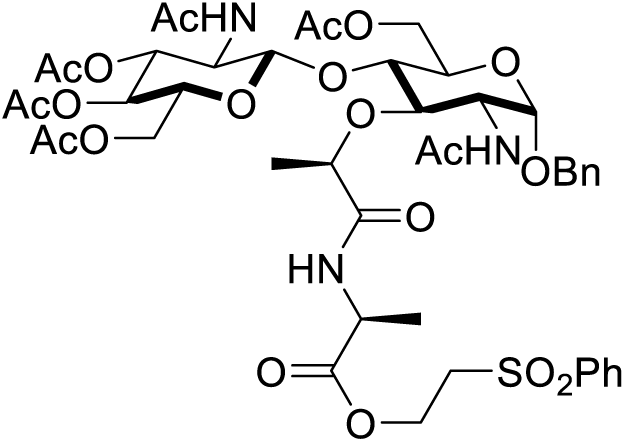

Benzyl ether **19** (1.0 g, 1.0 mmol) was dissolved in THF and MeOH (4:1, 40 mL) and degassed with an Ar balloon. A suspension of 10% palladium on charcoal (1.8 g, 1.7 mmol) was added to this solution and a H_2_ balloon bubbled through the resulting mixture. The reaction mixture was then stirred under hydrogen pressure (10 psi) for 6 h and filtered through a thin layer of celite. The celite was washed with MeOH (2 × 50 mL), the filtrate concentrated *in vacuo* and the resulting oil precipitated from ether and hexanes. The precipitate was filtered and dried to yield the lactol as a white solid (850 mg, 94%). The lactol was then dissolved in anhydrous CH_2_Cl_2_ (10 mL) and added rapidly via syringe to a vigorously stirred suspension of 5-ethylthio- 1*H*-tetrazole (575 mg, 4.41 mmol) and dibenzyl-*N*,*N*’-diisopropylphosphoramidite (0.96 mL, 2.85 mmol) in anhydrous CH_2_Cl_2_ (10 mL) under argon at ambient temperature. The reaction mixture became homogeneous within a few min. After 2 h, the mixture was diluted with CH_2_Cl_2_ (80 mL) and washed with saturated sodium bicarbonate (50 mL), water (50mL) and brine (50 mL). The organic solution was dried over anhydrous sodium sulfate and concentrated *in vacuo* to yield a colourless oil, which was precipitated from ether and hexanes (1:1) to yield the phosphite as a white solid. The product was dissolved in THF (20 mL) and cooled to −78 °C. Hydrogen peroxide (30%, 1.9 mL) was added dropwise via syringe to the vigorously stirred solution. After the addition was complete, the ice bath was removed and the mixture was allowed to warm to ambient temperature over 2 h. The reaction mixture was then diluted with ice-cold saturated sodium sulfite (5 mL), followed by EtOAc (50 mL), and stirred for 5 min. The organic layer was washed with saturated NaHCO_3_ (20 mL) and brine (20 mL), dried over anhydrous sodium sulfate and concentrated *in vacuo* to yield phosphate **20** as a white solid (1.02 g, 93%). 1H NMR (DMSO-*d*_*6*_, 400 MHz) δ 8.70 (d, *J* = 4.6 Hz, ^1^H, NHAc), 8.43 (d, *J* = 6.9 Hz, 1H, NHAc), 8.09 (d, *J* = 9.0 Hz, 1H, NHAc), 7.87-7.85 (m, 2H, *ortho*-ArH), 7.76-7.72 (m, 1H, *para*-ArH), 7.65-7.62 (m, 2H, *meta*-ArH), 7.39-7.30 (m, 10H, 2 × Bn-ArH), 5.81 (dd, *J* = 6.3, 3.1 Hz, 1H, MurNAc-H1), 5.24 (t, *J* = 9.9 Hz, 1H, GlcNAc-H3), 5.08-4.96 (m, 4H, 2 × C*H*_*2*_Ph), 4.91 (t, *J* = 9.8 Hz, 1H, GlcNAc-H4), 4.73 (d, *J* = 8.3 Hz, 1H, GlcNAc-H1), 4.60 (d, *J* = 6.7 Hz, 1H, MurNAc-C*H*O), 4.33-4.18 (m, 3H, MurNAc-H6 + GlcNAc-H6 + OC*H*H), 4.08-3.95 (m, 4H, MurNAc-H6 + GlcNAc-H6 + OC*H*H + Ala1Hα), 3.87-3.73 (m, 4H, GlcNAc-H_2_ + GlcNAc-H5 + MurNAc-H3 + MurNAc-H5), 3.64-3.55 (m, 3H, MurNAc-H_2_ + SC*H*_*2*_), 3.42 (dd, *J* = 10.9, 8.7 Hz, 1H, MurNAc-H4), 1.97 (s, 3H), 1.96 (s, 3H), 1.95 (s, 3H), 1.92 (s, 3H), 1.75 (s, 3H), 1.69 (s, 3H), 1.29 (d, *J* = 6.7 Hz, 3H, MurNAc-C*H3*), 1.11 (d, *J* = 7.3 Hz, 3H, AlaHβ); ^13^C NMR (DMSO-*d*_*6*_, 125 MHz) δ 174.5, 171.3, 169.9, 169.8, 169.5, 169.3, 139.2, 135.7, 133.9, 129.3, 128.4, 128.3, 128.3, 128.3, 127.9, 127.8, 127.7, 127.6, 127.6, 99.6, 75.9, 75.7, 73.8, 72.3, 70.7, 70.4, 68.6, 68.4, 68.4, 68.3, 66.3, 61.6, 57.9, 53.6, 47.3, 40.0, 39.9, 39.8, 39.7, 39.6, 39.6, 39.5, 39.4, 39.3, 39.1, 39.0, 22.6, 22.3, 20.5, 20.3, 20.2, 18.9, 16.5; RMS (ES) Calcd for C_52_H_66_N_3_NaO_23_PS [M+Na]^+^ 1186.3, found 1186.3.

#### *N*-[*N*-Acetyl-6-*O*-acetyl-1-*O*-[bis(phenylmethoxy)phosphinyl]-4-*O*-[3,4,6-tri-*O*-acetyl-2- (acetylamino)-2-deoxy-β-D-glucopyranosyl]-α-muramoyl]-L-alanyl-L-γ-glutamyl-*N*6- (2,2,2-trifluoroacetyl)-L-lysyl-D-alanyl-2,5-dimethyl ester (21)

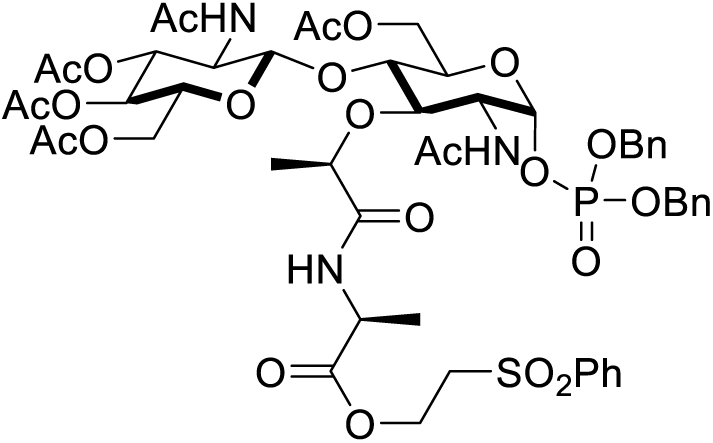

Disaccharidyl ester **20** (350 mg, 0.3 mmol) was dissolved in dry CH_2_Cl_2_ (3 mL) and stirred at ambient temperature under argon. A solution of Diazabicycloundec-7-ene (45 μL, 0.3 mmol) was added and the resulting solution stirred for 1 h. The reaction mixture was diluted with CH_2_Cl_2_ (15 mL), washed with 1 M HCl (5 mL) and brine (5 mL), dried over anhydrous Na_2_SO_4_ and concentrated *in vacuo*. The oil was precipitated with Et_2_O dried under high vacuum for 2 h to yield the acid as a white solid. The acid was dissolved in dry DMF (5 mL) and cooled to 0 °C with an ice-bath. HATU (114 mg, 0.3 mmol), followed by DIPEA (157 μL, 0.9 mmol) were added and the resulting yellow solution stirred for 15 min. Tetrapeptide **13** (191 mg, 0.3 mmol) was added and the resulting solution stirred at ambient temperature for 24 h. The reaction mixture was then concentrated *in vacuo* and re-dissolved in CHCl_3_ and IPA (9:1, 10 mL) and washed with 1 M HCl (5 mL) and saturated sodium bicarbonate (5 mL). Both aqueous washes were back-extracted with CHCl_3_ (5 mL) and the combined organic extracts washed with brine (2 × 5 mL), dried over anhydrous sodium sulfate and concentrated *in vacuo*. The crude product with precipitated from Et_2_O to yield the pentapeptidyl disaccharide **21** as an off-white solid (410 mg, 90%), which was used directly in the next step without further purification.

#### Lipid II diammonium salt (22)

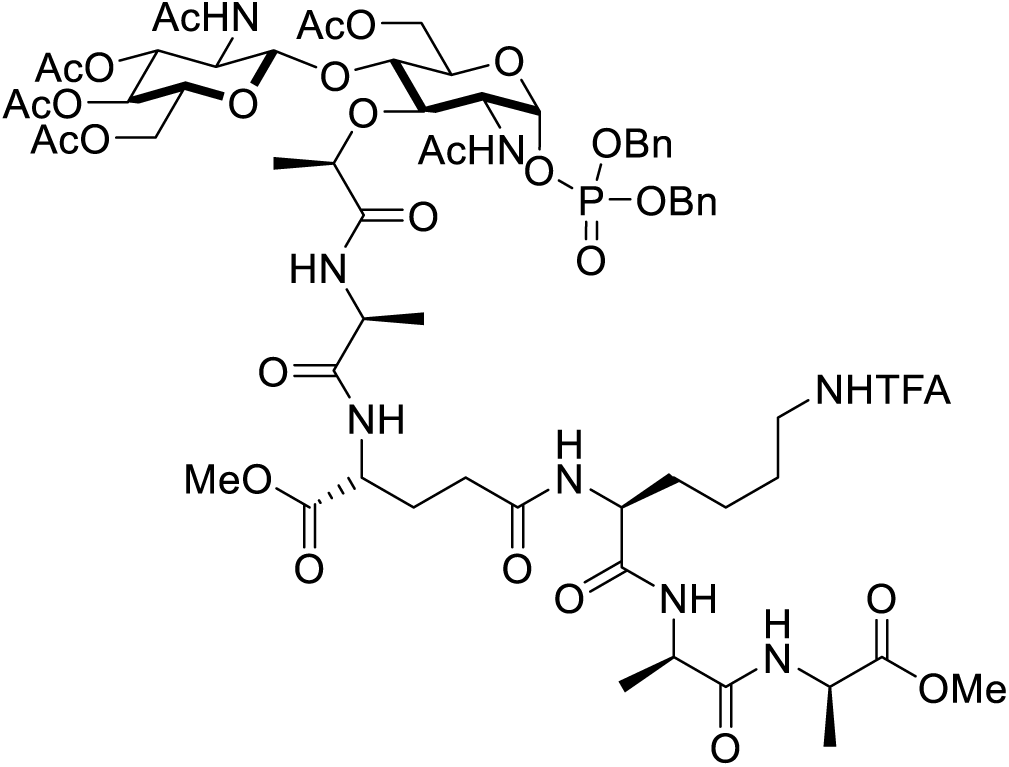

Dibenzyl phosphate **21** (50 mg, 33 μmol) was dissolved in anhydrous MeOH (6 mL) and the flask flushed with an argon baloon. Pd/C (10 % w/w, 106 mg, 99 μmol) was added and the resulting suspension stirred under a H_2_ atmosphere for 3 h. The suspension was then filtered through celite, which was washed with MeOH (2 × 3 mL). Pyridine (1 mL) was added to the filtrate, which was then concentrated *in vacuo* and dried by high vacuum for 1 h to yield the sugar phosphate salt as a white solid. This salt was dissolved in dry DMF (1 mL) and dry THF (1 mL) and carbonyl diimidazole (26.8 mg, 165 μmol) was added. The resulting clear solution was stirred at ambient temperature for 2 h, at which point analysis by ESI showed complete product formation ([M-H]^−^ = 1388.4). Excess carbonyl diimidazole was destroyed by the addition of dry MeOH (5.34 μL, 132 μmol) and stirring continued for 45 min. The reaction mixture was then concentrated *in vacuo* and dried under high vac for 1 h. To resulting activated phosphate was added a solution of UPBA (29 mg, 33 μmol) in THF (2 mL) and 5-ethylthio- 1*H*-tetrazole (4.3 mg, 33 μmol). The resulting solution was stirred for 96 h under argon at ambient temperature and concentrated *in vacuo*. To this crude mixture was added 1,4-dioxane (1 mL) and a solution of sodium hydroxide (40 mg, 1 mmol) in water (1 mL). The resulting mixture was stirred at 37 °C for 2 h and filtered through an aqueous filter disc, which was washed with 1:1 H_2_O/1,4-dioxane (2 mL). Lipid II was then purified by HPLC: column = Phenomenex Luna C_18_(2) 100 Å prep-scale column; flow-rate = 20 mL/min, UV = 220 nm, method: solvent A = 50 mM NH_4_HCO_3_(aq), solvent B = MeOH, gradient = 2 to 98 % B over 30 min, 98 % B for 10 min, 98 to 2 % B over 1 min and 2 % B for 4 min. Lipid II eluted between 33.7 – 34.4 min. Product containing fractions were concentrated by rotary evaporator and diluted with H_2_O, frozen and lyophilized to yield Gram-positive lipid II **22** as a fluffy white powder (17.7 mg, 29%). HRMS (ESI) Calcd for C_94_H_154_N_8_O_26_P_2_ [M-2H]^−^ 936.52302, found 936.52667.

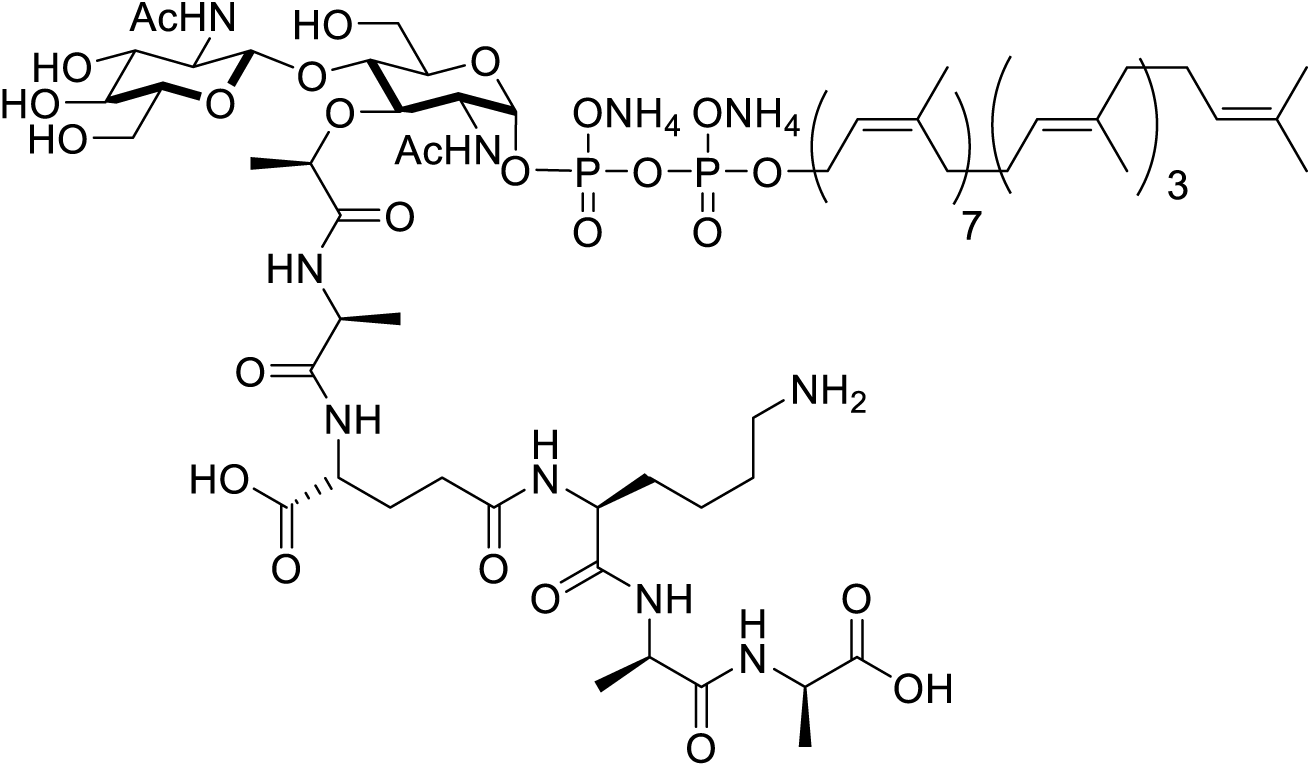

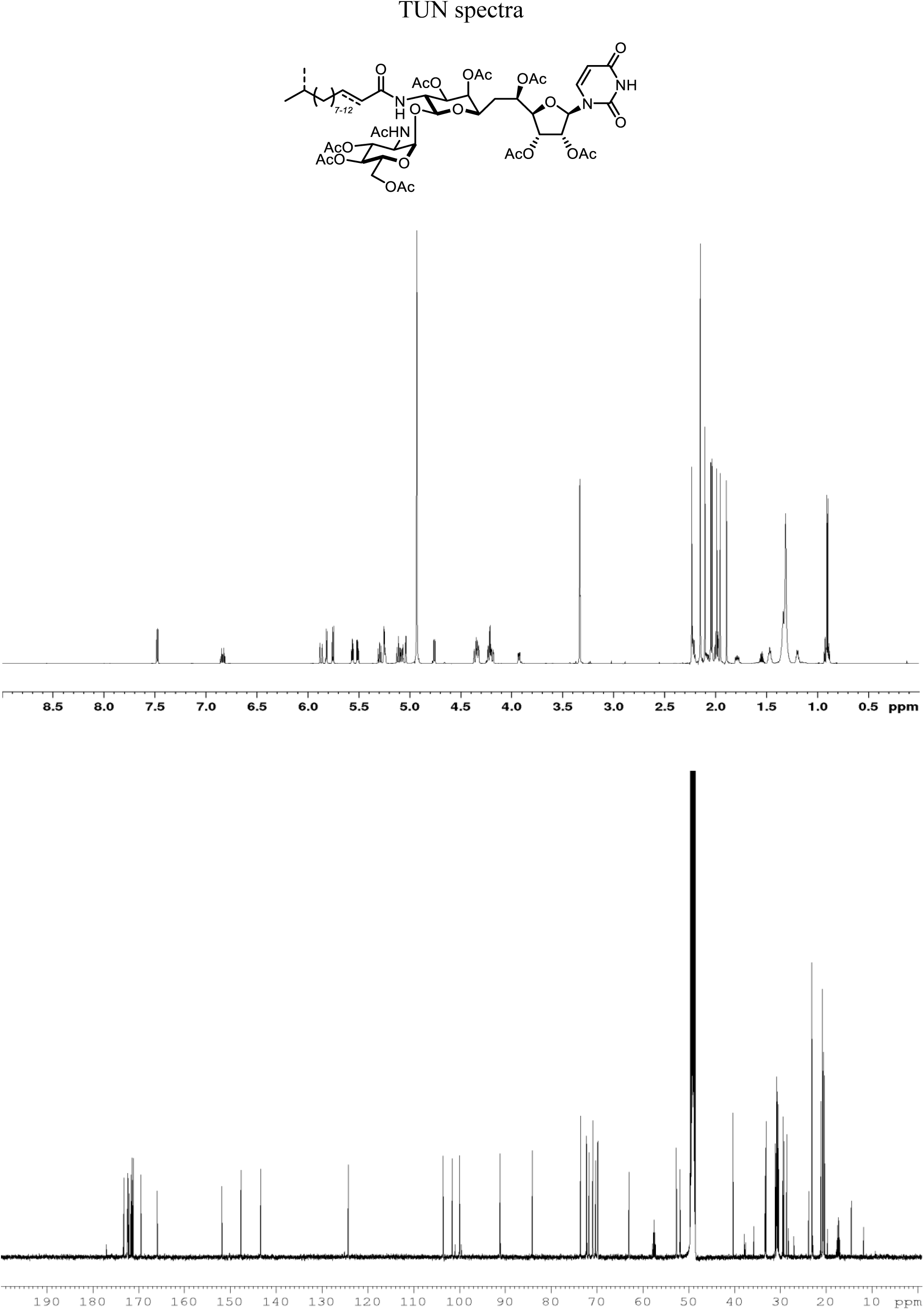

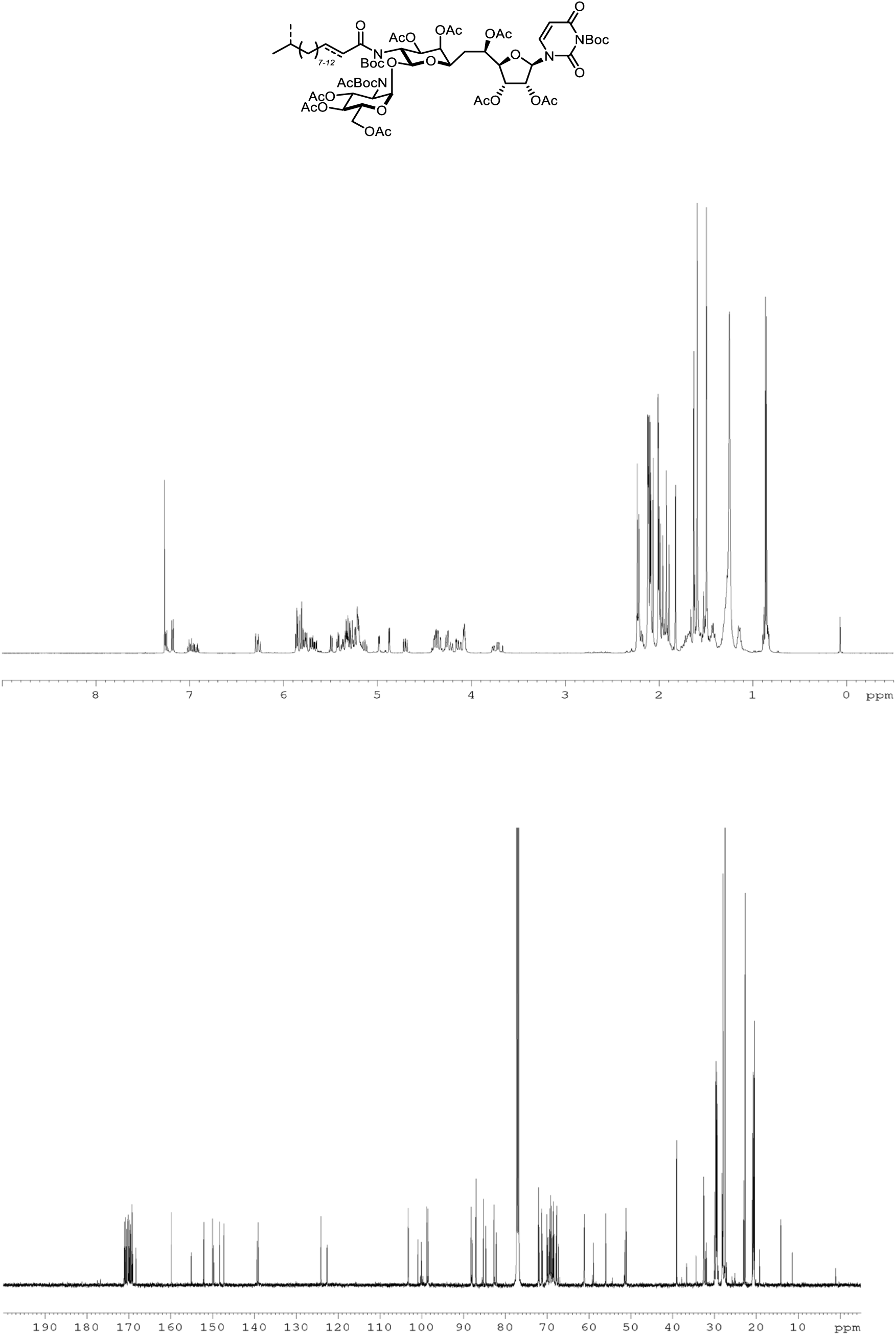

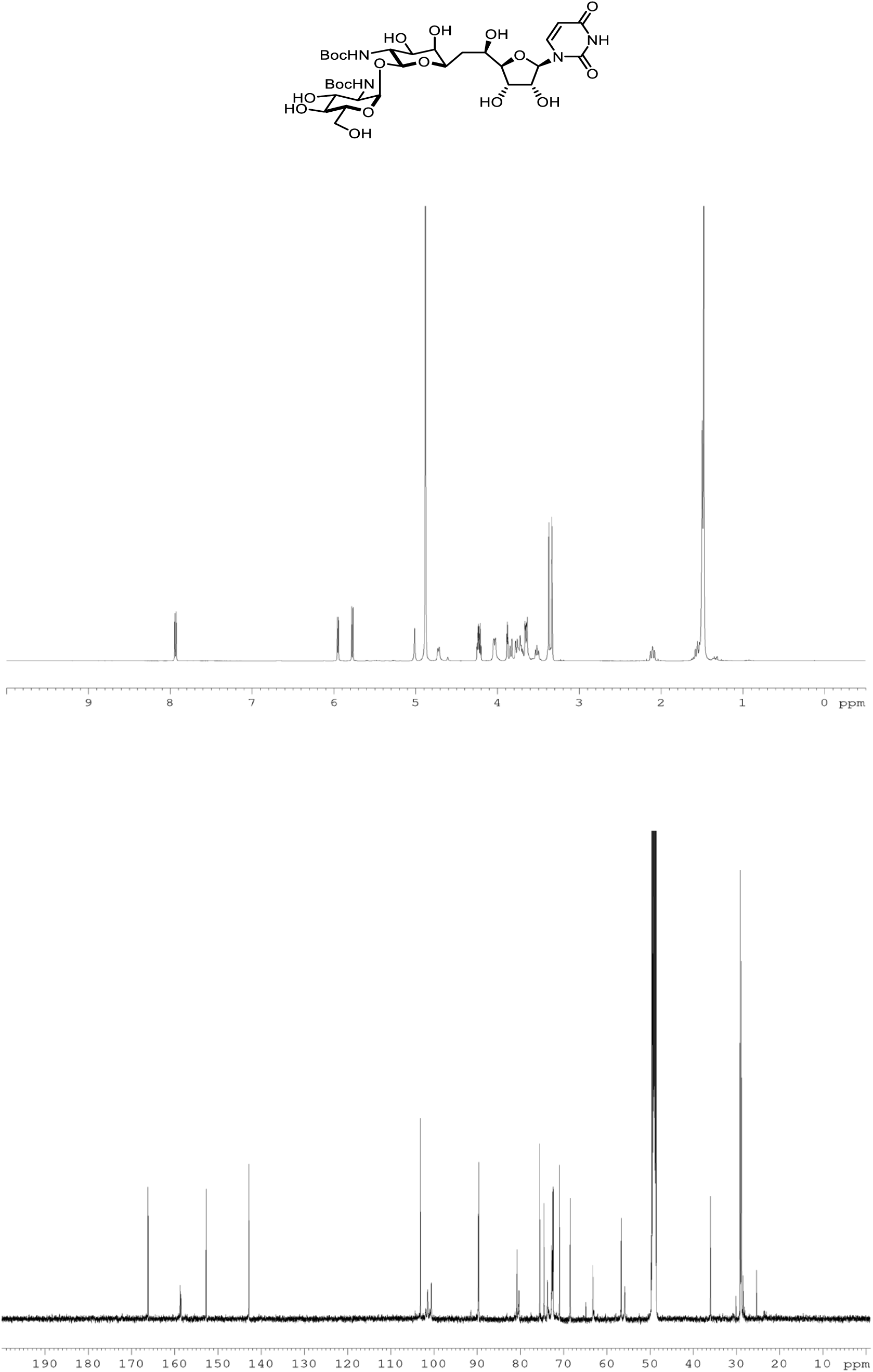

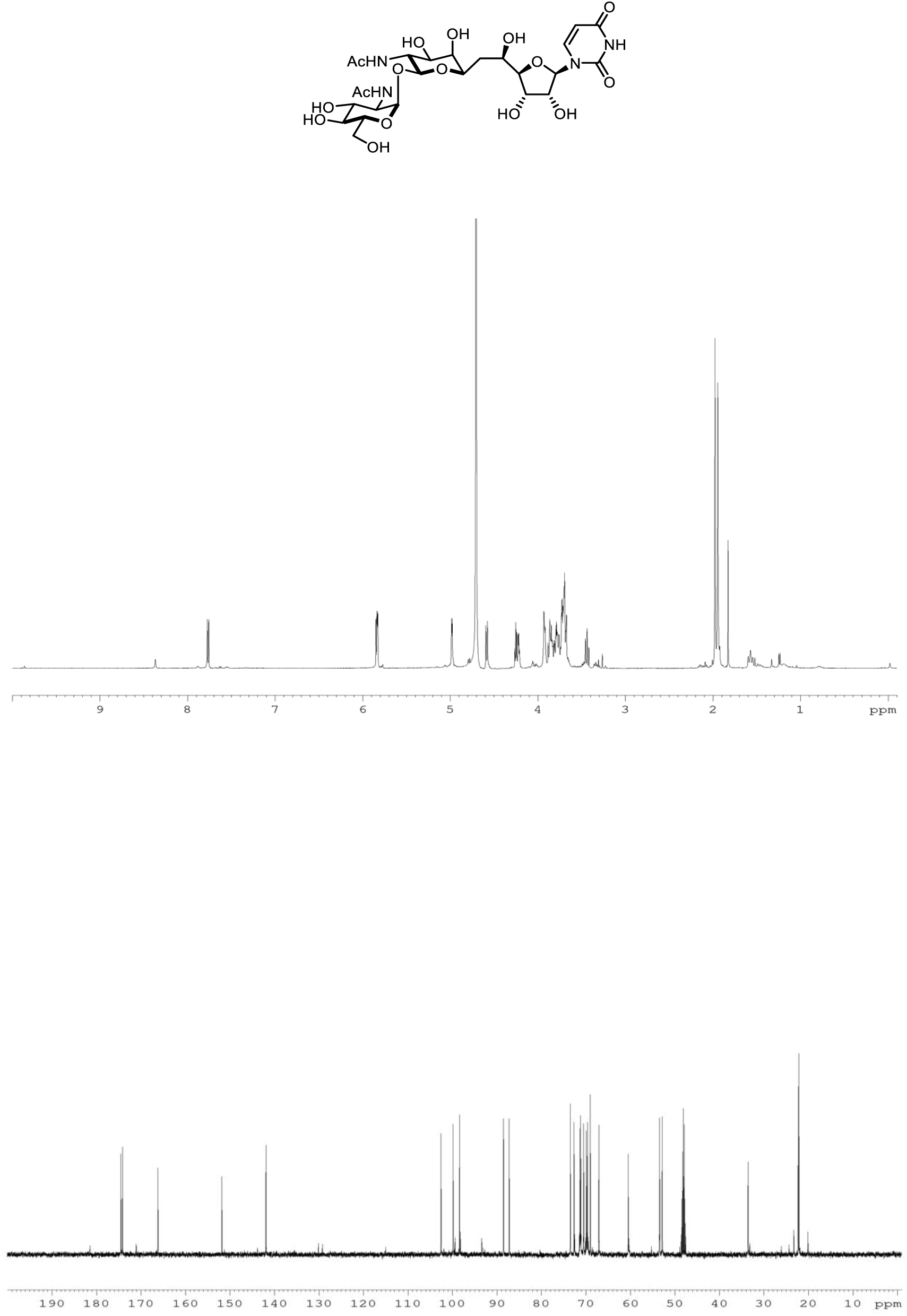

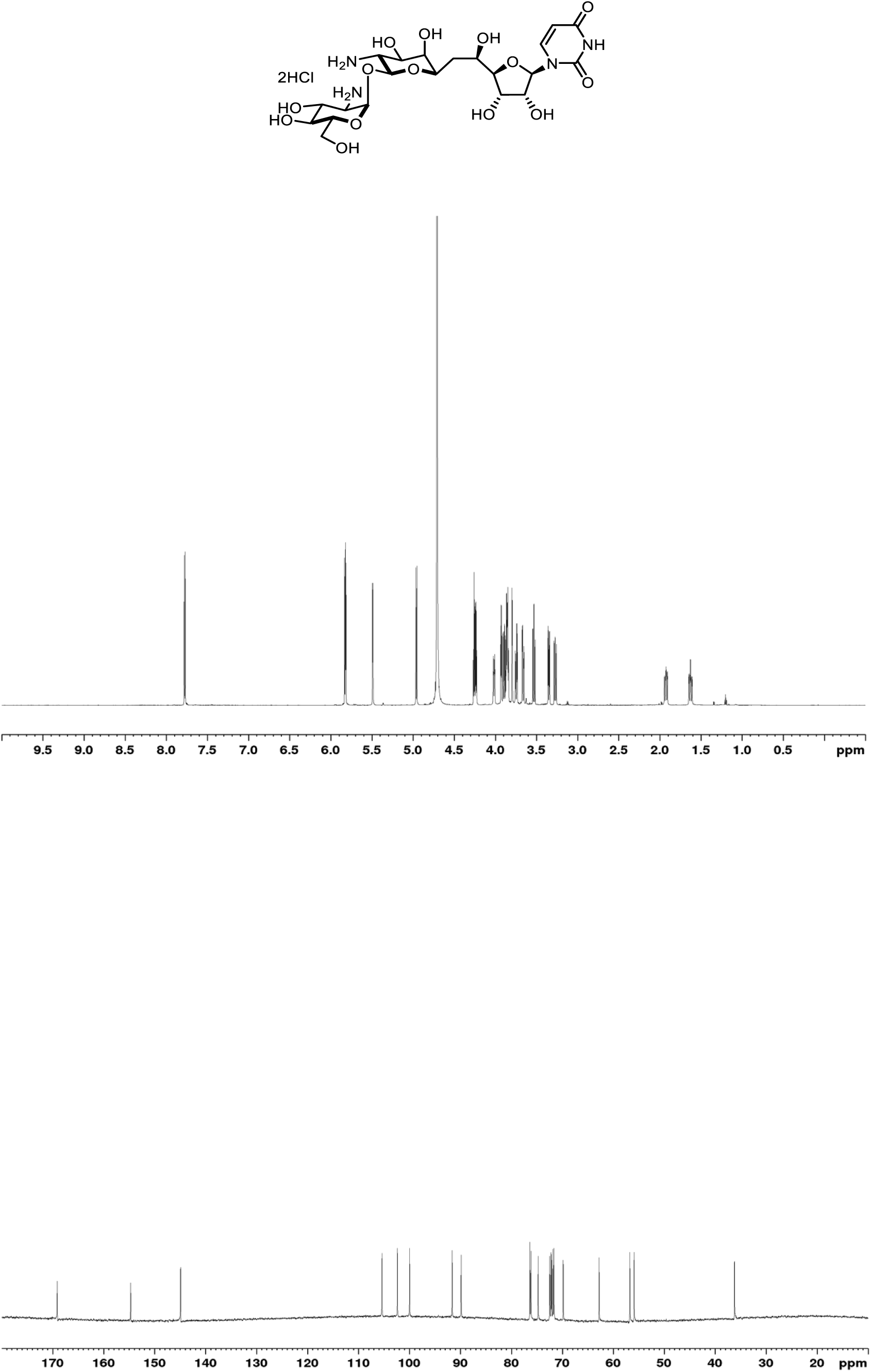

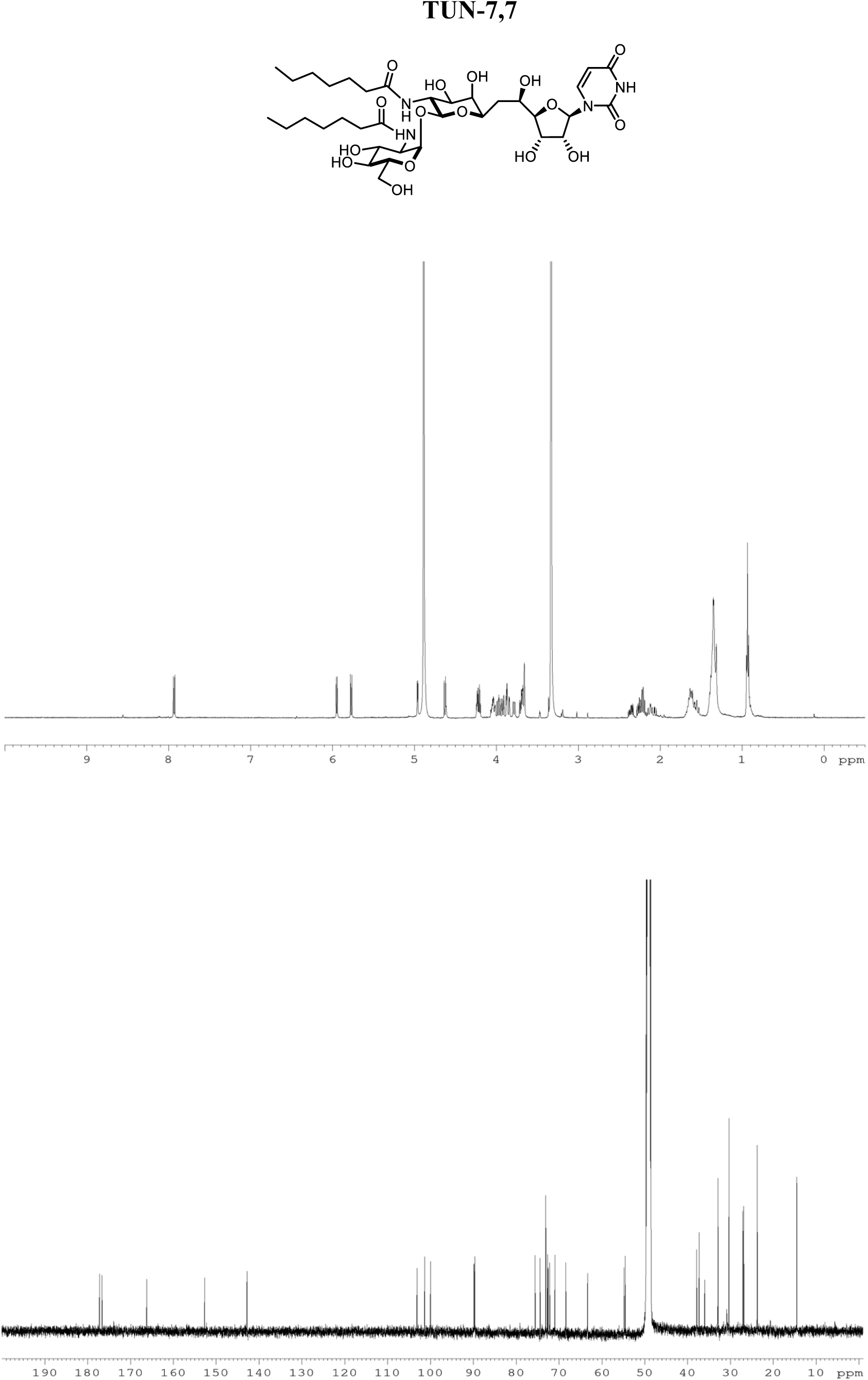

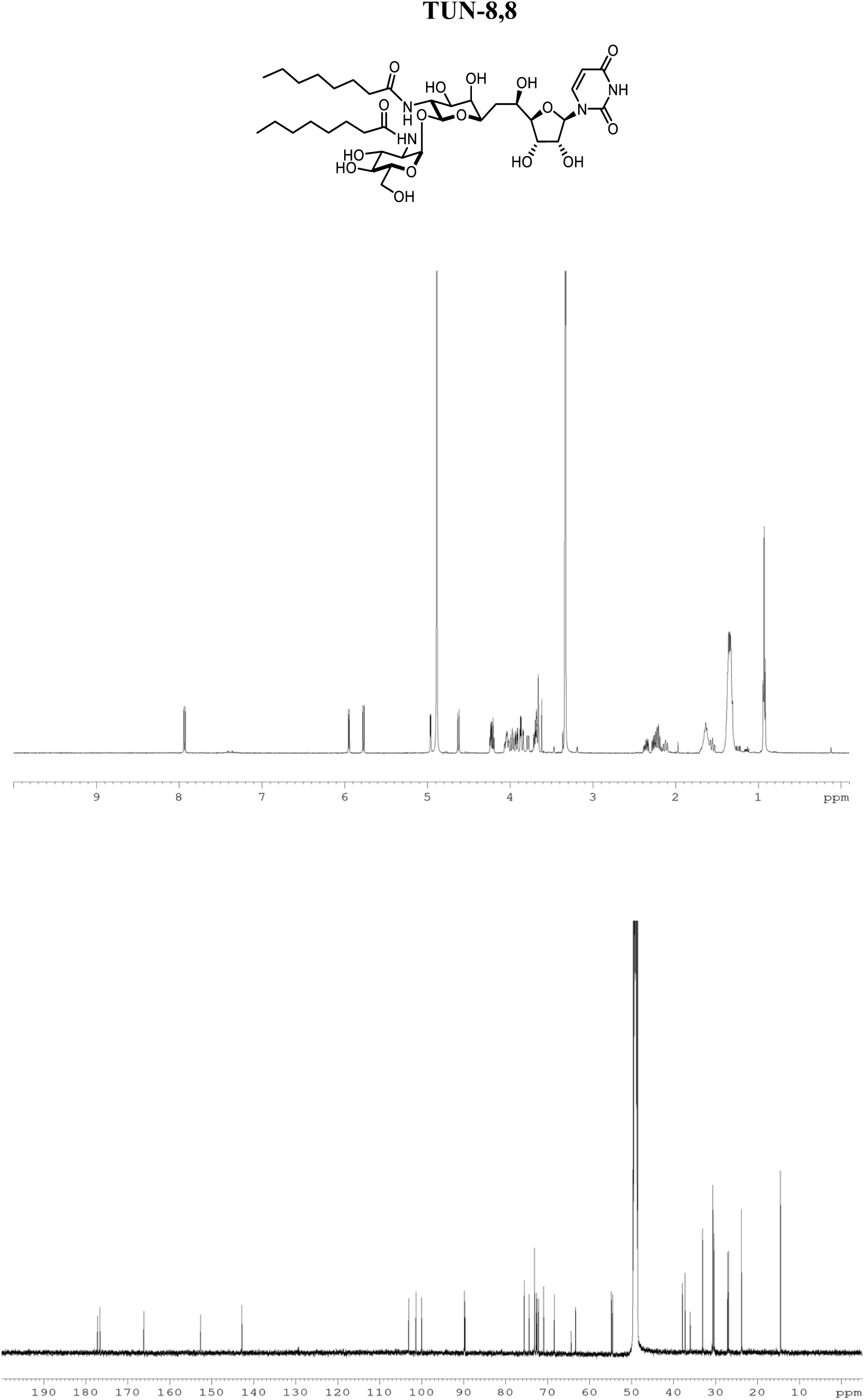

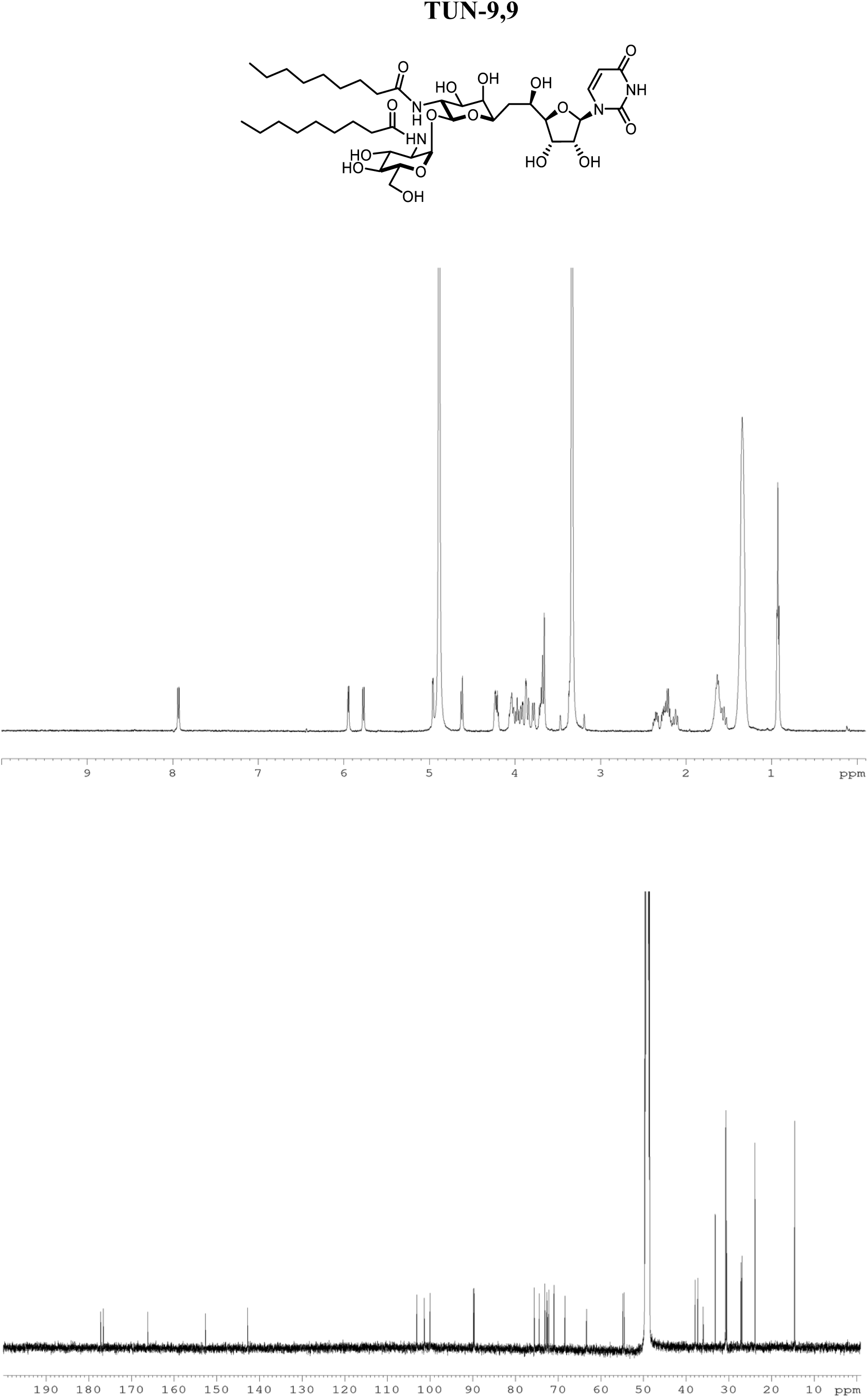

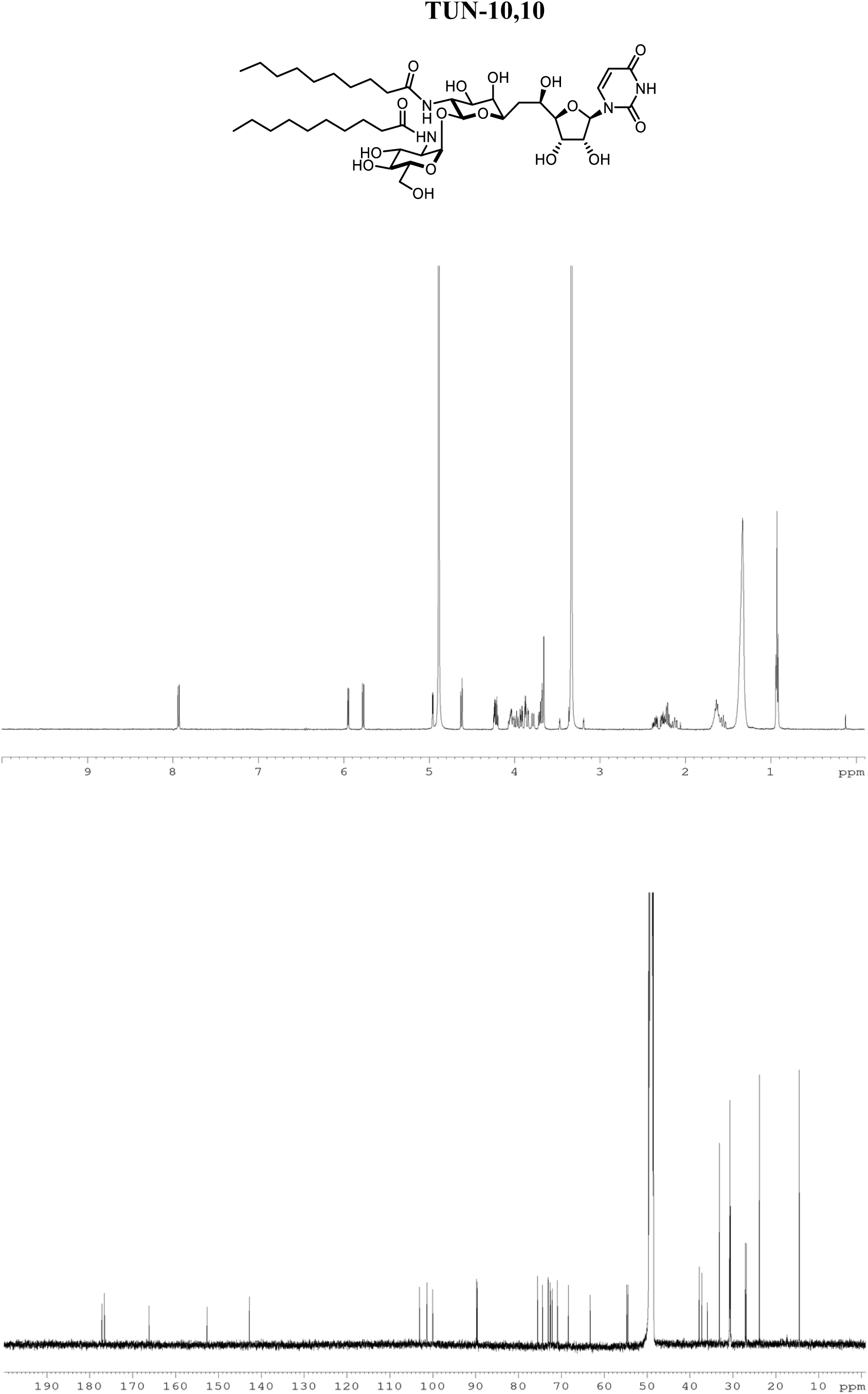

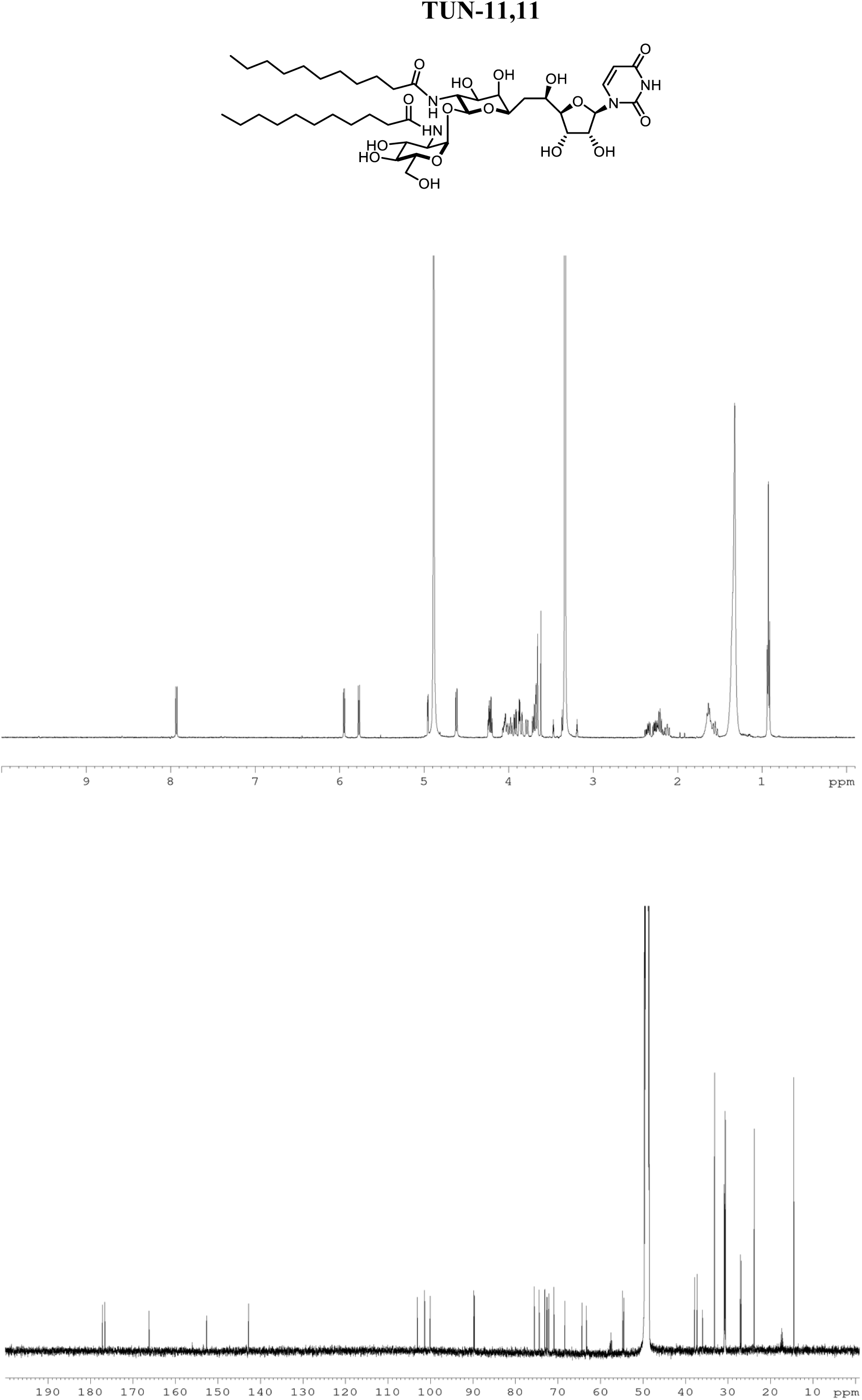

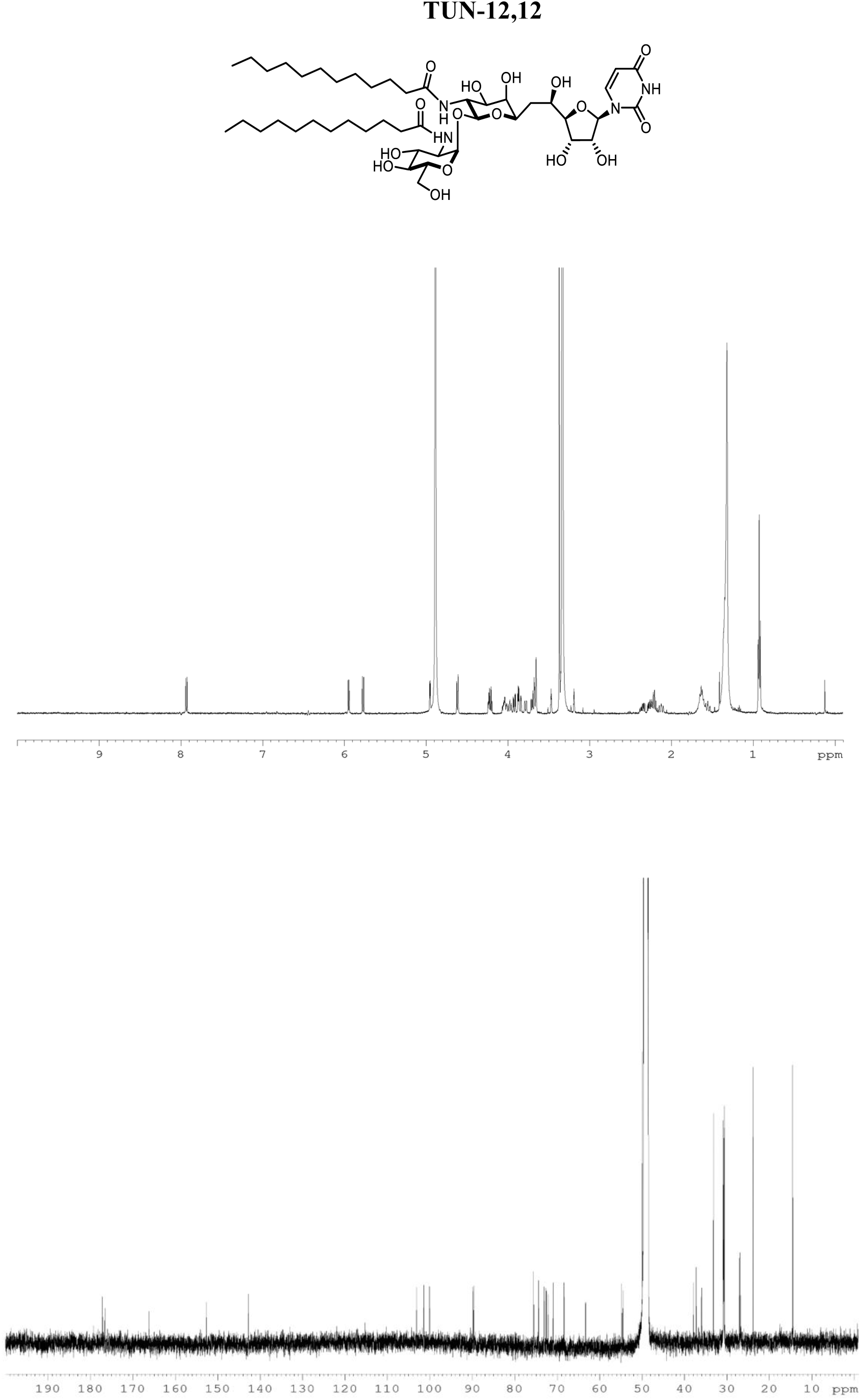

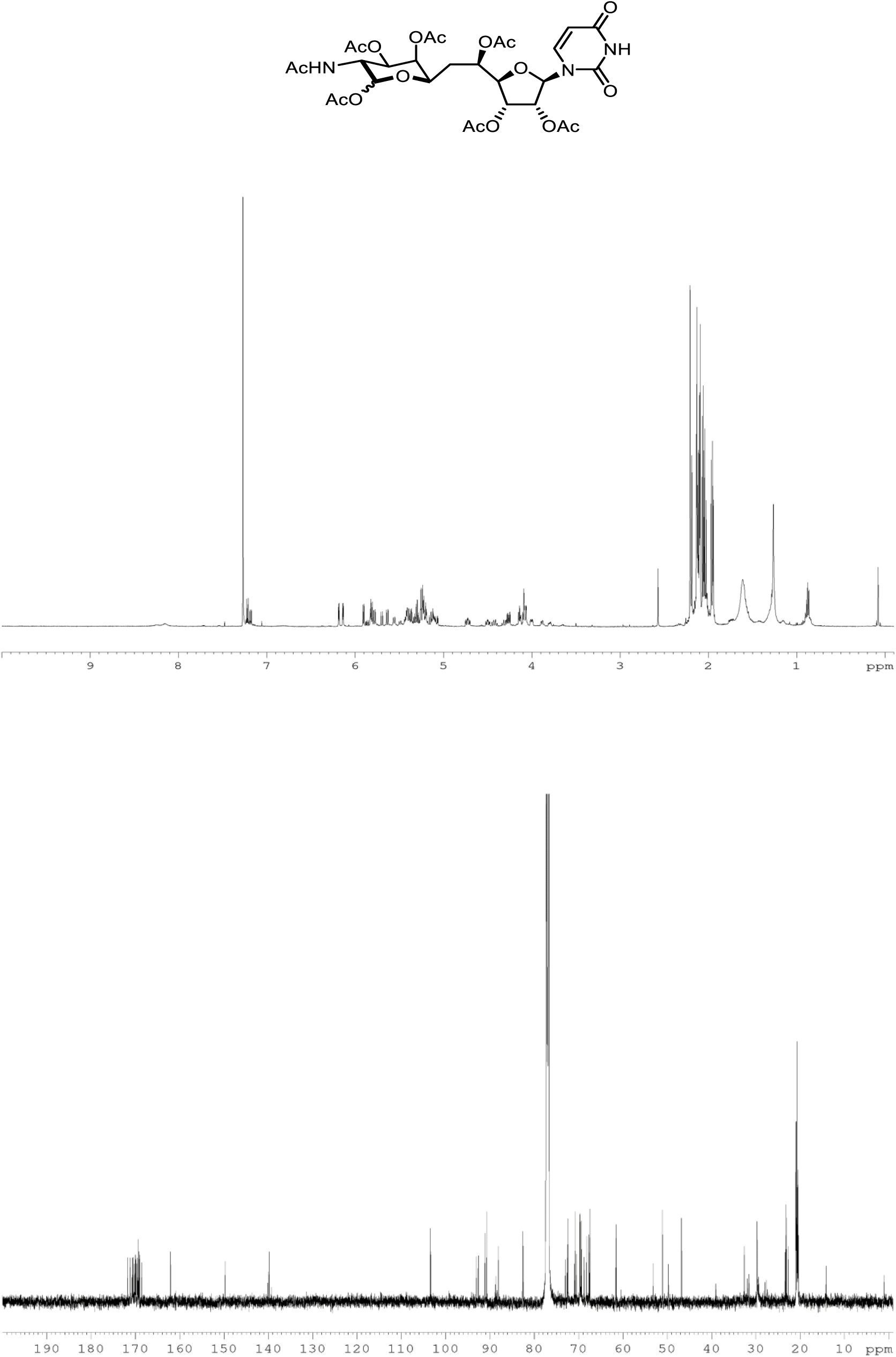

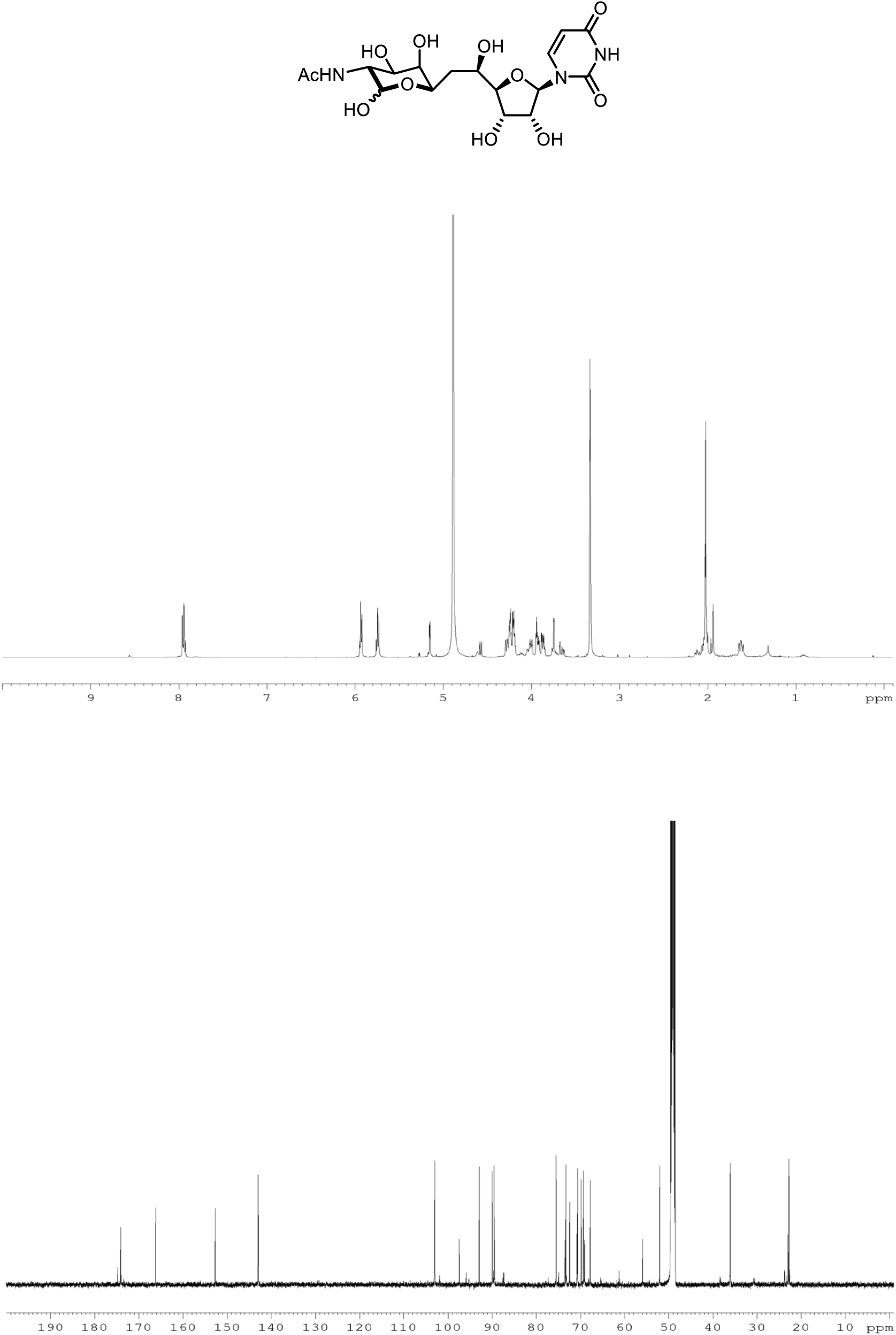

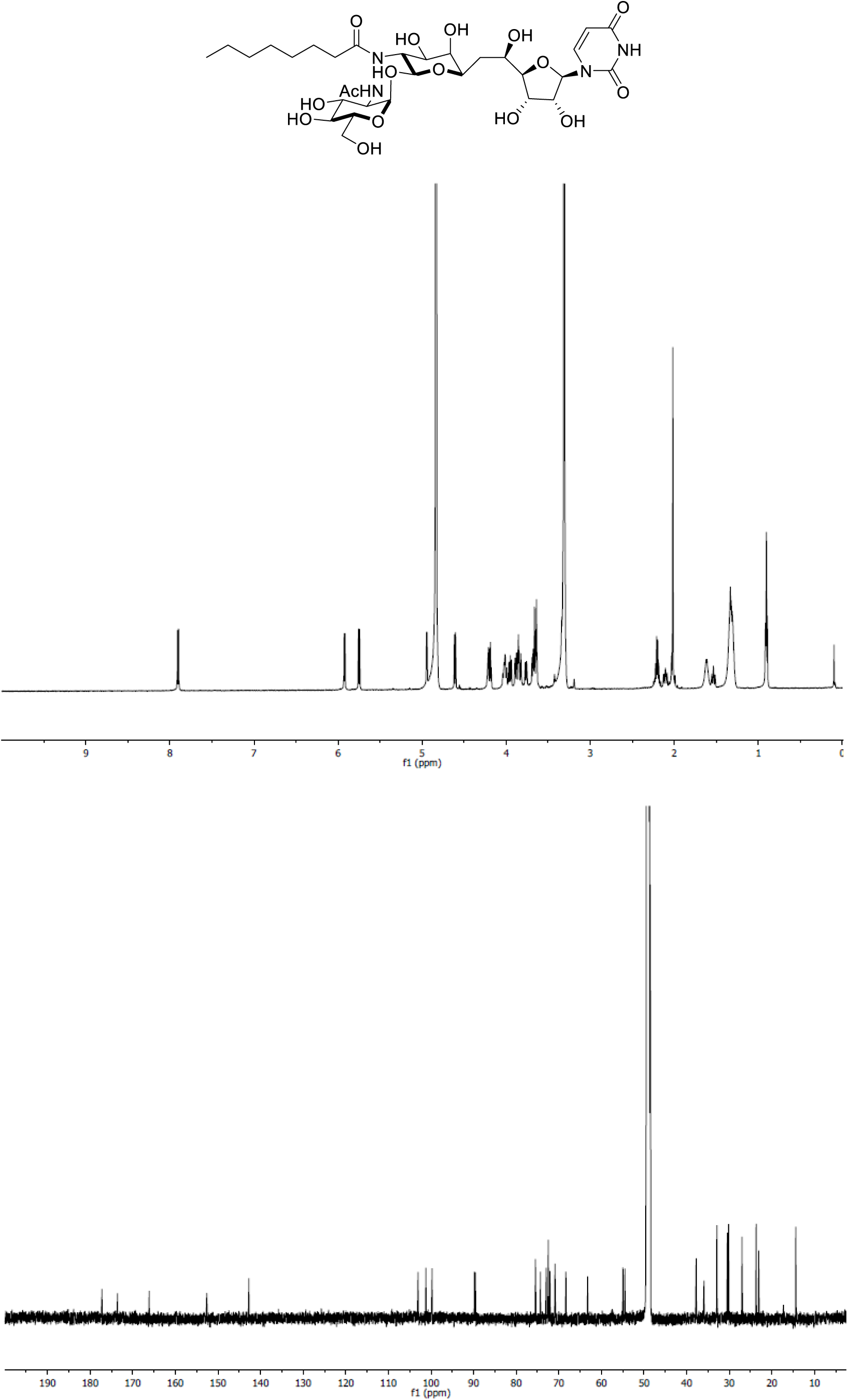

